# Pangenome alignment reveals global diversity and evolution of human centromeric regions

**DOI:** 10.64898/2026.09.03.749043

**Authors:** Jordan Eizenga, Mira Mastoras, Julian K. Lucas, Julian Menendez, Faith Okamoto, Glenn Hickey, Prajna Hebbar, Sasha A. Langley, Hailey Loucks, Fedor Ryabov, Yulia Zybina, Mobin Asri, J. Matthew Franklin, Nicolas Altemose, Human Pangenome Reference Consortium, Ivan A. Alexandrov, Charles H. Langley, Benedict Paten, Karen H. Miga

**Author notes:** Shared first authorship (Jordan Eizenga, Mira Mastoras, Julian K. Lucas). Shared co-corresponding authorship (Benedict Paten, Karen H. Miga).

## Abstract

Centromeres play essential roles in chromosome segregation and genome stability, yet they remain among the least characterized regions of the human genome. Despite advances in long-read sequencing and complete genome assembly, the extreme repetitiveness and structural complexity of these regions still challenge population-scale analysis, obscuring their mutational dynamics. The Human Pangenome Reference Consortium has now accurately assembled over 6,000 centromeres, providing an opportunity to catalog global centromere variation. However, centromeric regions have been systematically excluded from pangenome alignments due to the technical challenge of aligning their highly repetitive tandem arrays and extreme structural variability. Here we introduce Centrolign, a graph-based multiple sequence alignment tool that combines a uniqueness-driven objective function with partial-order partial-order alignment to accurately align alpha satellite higher-order repeats. By prioritizing rare matches within tandem arrays and leveraging extended centromere-spanning haplotypes formed by suppressed recombination, Centrolign produces progressive multiple sequence alignments that preserve ancestral repeat organization. Applied across human centromeres, these alignments reveal the phylogenetic structure of similar satellite array haplotypes and enable precise estimation of variation rates, structural variant frequencies, and spatial patterns of mutation within satellite arrays. Integrating Centrolign graphs with repeat annotation tools and pangenome mapping algorithms allows accurate variant calling and genotyping from long reads without prior assembly. Moreover, we show that centromere haplotypes can be accurately subtyped with k-mers alone. Together, these advances establish a robust framework for incorporating centromeres into broader pangenomes, and population genomics in general, advancing our understanding of human genome evolution and diversity.

## Introduction

Human centromeres are specialized chromosomal regions that serve as the sites of kinetochore assembly, ensuring faithful chromosome segregation during mitosis and meiosis^1^. In humans, centromeres almost invariably coincide with α-satellite (αSat) DNA ^2–4^, which forms highly repetitive arrays spanning several megabases^5^. Within these arrays, 171-bp αSat monomers are organized into chromosome-specific higher-order repeats (HORs) that are themselves repeated in tandem^6^. The αSat arrays that recruit the kinetochore, termed active arrays, are embedded within broader satellite-rich regions containing inactive αSat, other satellite families, segmental duplications, as well as nonrepetitive sequences^7^. For decades, the immense size of these arrays and the near-identical sequence of neighboring HOR copies prevented complete assembly of human centromeres, leaving one of the most fundamental regions of the genome largely inaccessible to genetic and genomic investigation. Complete telomere-to-telomere (T2T) assemblies, beginning with the CHM13 reference genome, provided the first complete assembly of every centromeric region in a human genome^7,8^. Combined with long-read methylation profiling, they also revealed the first genome-wide maps of functional centromeres by localizing the kinetochore to regions of hypomethylation within the otherwise hypermethylated active αSat array, termed centromere dip regions (CDRs)^9–11^. Initial studies showed that CDRs tend to occupy young, more homogeneous portions of active αSat arrays^7,12,13^, linking centromere function to the evolutionary organization of centromeric DNA.

Building on advances in complete human genome assembly, the Human Pangenome Reference Consortium (HPRC)^14^, the Human Genome Structural Variation Consortium (HGSVC)^13,15^, and other global initiatives^16^ are now generating complete centromere assemblies from hundreds of diverse human genomes. The next challenge is to determine how these sequences are related across individuals. This will allow the variation that distinguishes human centromeric satellites to be traced through its evolutionary history and related to centromere function. Doing so requires establishing homology across centromeres whose sequence and structure have greatly diverged among individuals.

Across most of the genome, a linear reference provides a stable coordinate system for relating homologous sequences and genetic variation across individuals. Within centromeric regions, however, megabase-scale inversions, expansions, contractions, and changes in satellite composition continually remodel the organization of satellite arrays^7,13,17^. Homologous active αSat arrays often differ not only in sequence but also in their underlying architecture^7,15,18^. Suppressed meiotic recombination across the centromere preserves long centromere-spanning haplotypes, or cenhaps, upon which detailed phylogenetic trees can be constructed. Such trees provide a biologically meaningful basis upon which to investigate the unique structural diversity of centromeres^19^. Assuming clock-like divergence in alignable, unique-sequence regions, the descendants of distinct cenhap lineages extant at an arbitrary time in the past can be clustered into evolutionarily related genomes. In some cases, large blocks of sequence present in one cenhap are entirely absent from another^7^. This creates a fundamental reference problem in which no single haplotype can represent the full sequence and structural diversity of the region, limiting consistent comparison, read mapping, and representation of genetic variation.

Pangenome approaches, which jointly represent sequences from multiple genomes rather than relative to a single reference^20^, provide a natural framework for representing centromere diversity by establishing homology across many structurally distinct assemblies rather than forcing every sequence against a single reference. However, to date, pangenome alignments have either excluded the centromeres^21^ or collapsed repeat copies into very high degree tangles that obfuscate information about recent evolutionary events ^22^. Here, we combine highly accurate complete centromere assemblies from the HPRC2 release^14^, state-of-the-art satellite annotation^7,23^, and Centrolign, a new method for accurately aligning homologous satellite arrays, to construct a centromere pangenome that establishes homology across structurally diverse human centromeres. Using this resource, we characterize how satellite arrays evolve through sequence mutational processes and structural rearrangement. We further investigate how these evolutionary processes shape the position and organization of functional centromeres, as defined by the CDR. Finally, we establish alignment strategies that enable sequencing reads to be mapped accurately to the centromere pangenome, allowing genetic variation to be identified and represented independently of any single reference genome. Together, this work provides the foundation for the integration of complete human centromeres into routine genetic and genomic analyses, enabling them to be studied with the same rigor and resolution as the rest of the human genome.

## Results

### Accurate assembly and annotation of thousands of peri/centromeric satellite arrays

To systematically characterize centromeric and pericentromeric satellites across the HPRC2 assemblies^14^, we used an automated pipeline based on the CenSat annotation developed for the T2T-CHM13 reference^7,25^. These annotations include αSats, classical human satellites (HSats 1-3), gamma satellites (γSats), and beta satellites (βSats). αSat annotations further distinguish active, inactive, and divergent HOR arrays, with active arrays defined as those that typically encompass the site of the functional centromere^7,26^. To define a high-confidence subset for analysis, we evaluated centromeric and pericentromeric assembly quality using multiple complementary quality metrics, including Flagger^27^ with HiFi and ONT as well as NucFlag^14,28^ (Methods). After quality filtering, the dataset retained broad centromere representation across every chromosome, with more than 100 well-assembled αSat arrays per chromosome except chromosome Y, which has fewer (43 unique haplotypes) due, in part, to its presence in only one in four haplotypes. The resulting high-confidence dataset comprises 6,946 active αSat arrays and 4,147 complete pericentromeric regions (Supplementary Figs. 1 and 2), providing a foundation for systematic analysis of satellite organization and evolution.

### Human pericentromeres exhibit extensive structural variation

Human centromeres and pericentromeres vary extensively in satellite content and structure (**Supplementary Figs. 3 and 4**), but the extent to which these differences resolve into recurrent structural configurations remains poorly understood. We compared satellite array content, order, and orientation across complete pericentromeres, revealing pervasive large-scale variation and recurrent structural configurations (Methods). Analyses were restricted to non-acrocentric autosomes and chromosome X, where assemblies were sufficiently reliable for these comparisons. Array-associated structural variation exceeding 20 kbp occurs on all chromosomes analyzed. Restricting to variation defined by events over 500 kbp found structural types different from CHM13 on all chromosomes analyzed except chromosomes 6, 8, 18, and X (**Supplementary Table 1**). Across chromosomes, these large-scale (>500 kb) structural differences define 81 distinct pericentromeric configurations (**Supplementary Table 1**), revealing multiple recurrent architectures within the human population.

Many common pericentromeric structures are not represented by the CHM13 reference (**Supplementary Fig. 5**). On chromosomes 1, 4 and 7, most haplotypes differ in gross structure from CHM13, including 86% of chromosome 1 haplotypes. For example, a 1.2 Mbp insertion on chromosome 1 is absent from T2T-CHM13 yet present in 68.8% of haplotypes (66/96). The insertion contains approximately 500 kbp of HSat3 and 360 kbp of βSat sequence, giving rise to bimodal distributions of HSat3 and βSat content among haplotypes (**Fig. 1a**). K-mer analysis identified the ∼500 kbp HSat3 array as the distinct HSat3B2 subfamily^7,29^, which is not represented elsewhere in the T2T-CHM13 genome (**Supplementary Fig. 6**). This reveals that entire, sequence-distinct satellite arrays spanning hundreds of kilobases can vary in presence or absence among human haplotypes.

**Figure 1.**
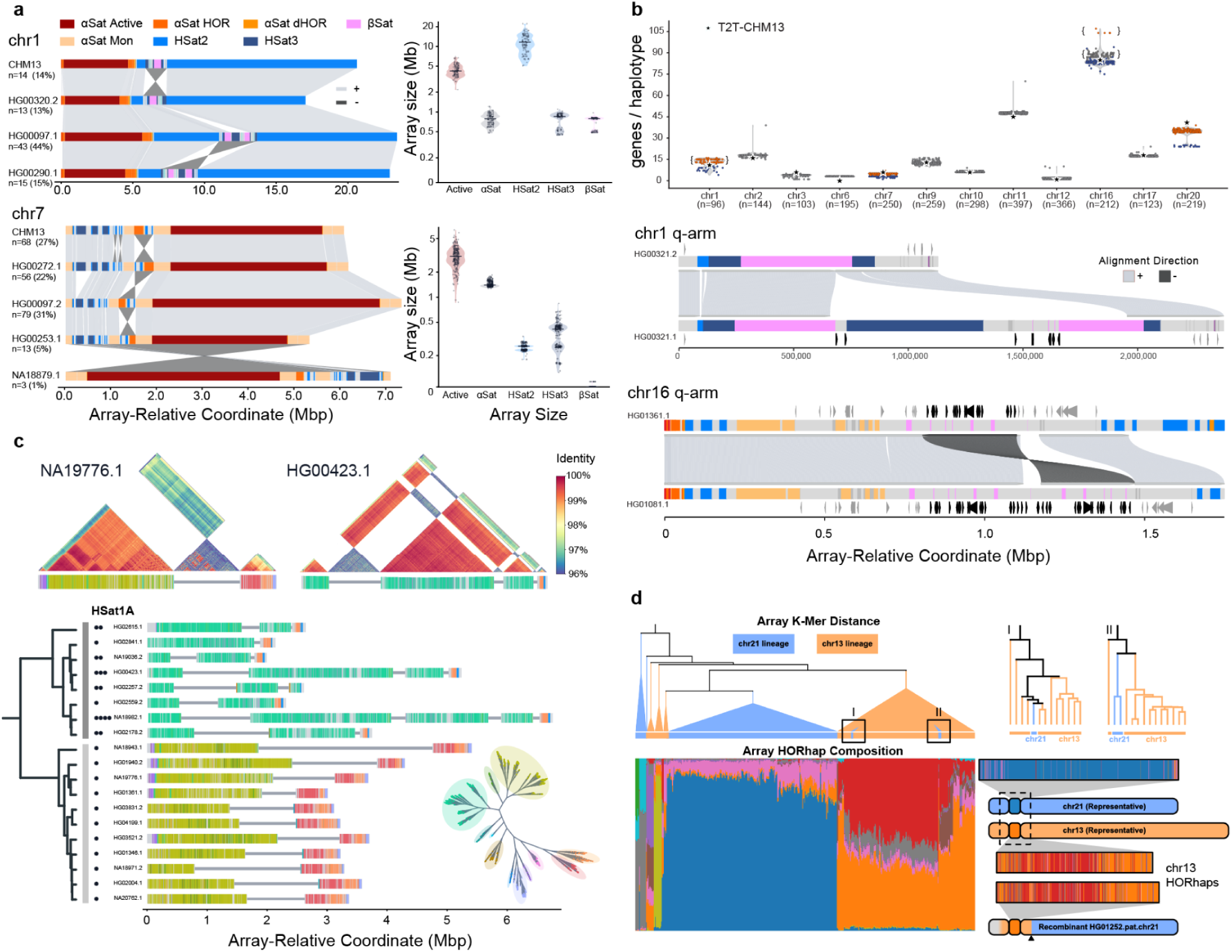
Large-scale structural diversity of human centromeric and pericentromeric regions. **a,** Representative satellite organization of chromosome 1 (top) and chromosome 7 (bottom) haplotypes. Satellite annotations are shown in array-relative coordinates, with ribbons indicating corresponding sequence and orientation among structurally distinct haplotypes. For each, n indicates the number of sampled haplotypes represented, and the percentage indicates its frequency among all sampled haplotypes for that chromosome. Distributions at right show the total sequence (Mbp) assigned to each satellite class across all haplotypes. Alpha satellites are broken down into “Active” for active arrays and “αSat” for all other types. **b,** Number of genes within pericentromeric regions for each haplotype, grouped by chromosome (n is the number of haplotypes represented); CHM13 is indicated by a star (top). Colored points indicate haplotypes carrying structural events associated with differences in gene copy number. Examples below show pairs of haplotypes with distinct structural arrangements and their CenSat annotation for chromosome 1q and chromosome 16q pericentromeric regions. Ribbons connect aligned sequences with light and dark grey indicating forward (+) and reverse (-) alignment orientations. Centromeric satellite annotations are colored according to legend in (a). Genes are shown along each haplotype, with genes that differ in copy number between the two haplotypes highlighted in black and other genes shown in grey. **c,** HSat1A organization within chromosome 4 active αSat arrays. Haplotypes are ordered by the cenhap lineages (left), and black circles indicate the number of HSat1A insertions within each array. HORs are colored according to phylogenetic clustering (HORhap tree inset, with HORhap grouping noted with color-shading). Two major clades differ in HSat1A, HORhap and αSat HOR organization (as indicated dark and light grey). ModDotPlot^24^ comparisons above show representative arrays and the HORhap arrays flanking the interspersed HSat1A. In HG00423.1 (right), αSat arrays with interspersed HSat1A retain the same HORhap assignment (teal, with shared high identity across blocks), whereas a distinct HORhap array (red) is observed q-arm proximal for NA19776.1 and typically observed with a single interspersed HSat1A array. **d,** Active arrays of chromosomes 13 and 21 are clustered by array-wide k-mer distance (top left), with chromosome-associated lineages indicated in orange (chromosome 13) and blue (chromosome 21). Enlarged portions of the tree (top right) illustrate local phylogenetic mixing between chromosome 13 and 21 haplotypes. The Admixture plot shows the composition of HOR haplotypes (bottom left), ordered according to the tree shown above. Representative arrays below show HORhap composition across chromosome-associated and recombinant haplotypes (bottom right), showing exchange on αSat between chromosome 13 and 21 arrays, with site of recombination noted with a black arrow.

Pericentromeric diversity includes widespread inversions, ranging from intra-satellite strand switching to multi-megabase inversions encompassing multiple satellites. Of the chromosomes analyzed for pericentromeric structural rearrangements, inversions were found on 6 (**Supplementary Fig. 5**). In some cases, pericentromeric inversions reorganized groups of satellite arrays and intervening sequence (**Fig 1a, Supplementary Fig. 5**). At the largest scale, entire pericentromeric regions are inverted, including chromosome 7 (**Fig. 1a**) (3 haplotypes) and chromosome 9 (1 haplotype). Chromosome 7 showed extensive inversion diversity, with multiple inversion sites and a broad range of inversion sizes. Overall, 232 of 252 haplotypes (92%) carried at least one inversion relative to T2T-CHM13, from recurrent 20–70 kbp HSat2-associated events and 550–660 kbp inversions spanning HSat2 and HSat3 to rare ∼6.9 Mbp inversions encompassing nearly the entire pericentromeric region (**Fig. 1a**). In the chromosomes analyzed, many inversion boundaries coincide with palindromic repeats, implicating local sequence architecture in recurrent rearrangement (**Supplementary Figs. 7 and 8**). Independently, array orientation, inferred from the k-mer composition of annotated satellite sequence (Methods), confirmed inversions^7^ within the chromosome 1 active array (93 of 385 haplotypes, **Supplementary Fig. 9**) and the large HSat3 array on chromosome 9. On chromosome 9, strand annotations identified approximately 129 orientation switches per haplotype (**Supplementary Fig. 10**). These findings show that pericentromeric inversions range from local satellite reorientations to megabase-scale rearrangements of the entire region.

Pericentromeric structural variation also altered the non-satellite sequence islands embedded within and flanking satellite arrays, which include protein-coding genes, lncRNAs and pseudogenes^8^. We identified 1,143 named genes within high-confidence pericentromeric regions across the genome and across all haplotypes, largely consistent with previously reported CenSat-associated genes in T2T-CHM13 (**Fig. 1b, top**). Gene content varies extensively among chromosome-matched haplotypes: 91.8% of pericentromeric genes are absent from at least one haplotype. Gene content is strongly associated with structural configuration and local satellite organization (**Fig. 1b, top**). The chromosome 1 polymorphic insertion described above includes gene-containing sequence, with structural configurations differing in the copy number of a segmentally duplicated region containing *FRG2C* (**Fig. 1b, middle**). Copy-number variation is associated with expression differences for a subset of pericentromeric genes. Approximately two-thirds of pericentromeric genes were detected in donor-matched Kinnex long-read RNA-sequencing data. On chromosome 16p, an interchromosomal segmental duplication introduced a second copy of *DUSP22*, and duplication carriers showed increased *DUSP22* transcript abundance (**Supplementary Figs. 11 and 12**). Additional copy-number-associated expression differences occur on chromosomes 1, 7 and 16q, including an inverted tandem duplication on 16q encompassing a segmental duplication containing 44 genes (**Fig. 1b, bottom**). Therefore, the recurrent structural configurations that distinguish human pericentromeres encompass not only megabase-scale differences in satellite organization, but also substantial variation in gene content and, in some cases, gene expression.

### Alpha satellite variation reflects distinct evolutionary histories

Having characterized large-scale structural diversity across pericentromeres, we next asked how variation within active αSat arrays relates to cenhap lineage. Active arrays evolve rapidly and have variation in both HOR structure and sequence^7^, raising the question of whether this diversity remains organized by shared ancestry. We inferred cenhap lineages from the alignable sequences flanking active arrays^19^. The five acrocentrics, which lack two chromosome-specific flanks, and chromosome Y, whose major haplogroups are already well characterized^30,31^, were excluded, yielding cenhap assignments for 18 chromosomes.

We first examined HOR structural variation (HOR-StVs), defined by differences in monomer organization within higher-order repeat units. Previous studies established that active αSat arrays contain HOR-StVs, but were limited to few haplotypes^7,32^. We therefore examined how common HOR-StVs are distributed across the HPRC2 assemblies. We found that HOR-StV composition varied substantially among chromosomes: some arrays contained one dominant HOR-StV, others carried a mixture of StVs shared in similar proportions across haplotypes, whereas on some chromosomes particular StVs were concentrated in subsets of haplotypes (**Supplementary Fig. 13**). Among 12 chromosomes with both cenhap assignments and common HOR-StVs, 8 showed lineage-dependent StV composition: particular StVs were either enriched within or restricted to specific cenhap lineages, including five chromosomes with clearly lineage-specific StVs. (**Supplementary Fig. 14**).

We next asked whether cenhap ancestry is detectable in HOR sequence composition. To identify related HOR sequences across active arrays, we aligned individual HORs and clustered them by sequence similarity into HORhaps^19,23,33^. In contrast to HOR-StVs (differences in monomer organization), HORhaps capture sequence-level variation among HORs including both single nucleotide differences as well as differences in monomer organization. We found that HORhap composition shows strong cenhap association on multiple chromosomes and that association is not driven primarily by HOR-StVs (**Supplementary Figs. 13, 15, and 16**). Among 16 chromosomes with cenhap assignments and interpretable HORhap patterns, 10 showed lineage-dependent HORhap composition, with particular HORhaps enriched in or restricted to specific cenhap lineages; seven chromosomes showed clear lineage-specific patterns and three showed weaker partitioning. These patterns show that arrays harbor extensive variation in HOR structure and sequence while still remaining associated with deeper cenhap ancestry.

Larger structural changes also show cenhap specificity. The chromosome 1 active αSat inversion described above is restricted to a single cenhap lineage (**Supplementary Fig. 9**), consistent with a single ancestral rearrangement. On chromosome 4, the two major cenhap lineages differ in HSat1A organization: one lineage consistently contains a single HSat1A insertion, whereas the other contains one to six insertions (**Fig 1c**). The surrounding HOR sequence also retains lineage structure: the two cenhap lineages contain distinct HORhap populations, and lineage-associated HORhaps persist between the variable HSat1A insertions. Active arrays therefore retain deep lineage relationships even as their internal sequence and structure continue to diversify.

### Active α-satellite arrays are exchanged between nonhomologous chromosomes

Active αSat arrays are generally composed of chromosome-specific HORs ^6,34^. Across the complete assemblies, however, we identified active αSat arrays with HOR sequences characteristic of a different chromosome (**Fig. 1d, Supplementary Figure 17, Supplementary Note 1, Fig. SN3**). Most prominently, HOR switching was seen between two pairs of acrocentric chromosomes: 13 and 21, and 14 and 22. These pairs are known to have closely related active αSat arrays and undergo recurrent interchromosomal exchange within their rDNA-bearing short arms^7,35–38^. We therefore investigated whether interchromosomal exchange among acrocentrics is restricted to their short arms or extends through the centromeric region.

In a tree inferred from k-mer-based sequence similarity, chromosome 13 and 21 active arrays are not consistently grouped by chromosome, with several sequence groups containing arrays from both chromosomes (**Fig. 1d, Supplementary Fig. 18, Methods**). This pattern could reflect either interchromosomal exchange or ancestral variation shared between the two chromosomes. Complete assemblies provided direct evidence for exchange. Three chromosome 21 haplotypes carry active arrays characteristic of chromosome 13. These haplotypes’ HORhap composition also supports a chromosome 13-associated ancestry (**Fig. 1d**, Methods). In one of these haplotypes (HG01252.pat.chr21), chromosome 13-associated sequence extends through the active array and into the q-arm, where the transition to chromosome 21-associated sequence is localized to approximately 128 kbp from the array, within a region of high sequence homology between the two chromosomes, which constitutes an inter-chromosomal segmental duplication (**Supplementary Note 1**, Methods). A corresponding transition in the p-arm could not be confidently resolved because of the complexity of the acrocentric short arms. Exchange was also observed in the reciprocal direction, with eight chromosome 13 haplotypes carrying chromosome 21-associated active arrays (**Supplementary Note 1**). We also identified a hybrid array (HG01960.mat.chr13) containing both chromosome 13- and chromosome 21-associated αSat sequence. The q-arm and adjacent half of the active array are chromosome 13-associated, whereas the remainder of the array is chromosome 21-associated, placing an interchromosomal recombination within the active αSat array itself (**Supplementary Note 1**).

Interchromosomal αSat exchange between chromosomes 14 and 22 in both directions was also evident (**Supplementary Note 1**). Separately, a previously reported chromosome 13/22 recombinant array, involving arrays with different HORs^13^, was confirmed in two HPRC samples (**Supplementary Note 1**). Together, these findings show that interchromosomal exchange among acrocentrics is not restricted to their rDNA-bearing short arms but can extend through active αSat and into the q-arm segmental duplications flanking the centromere. Ancestry transitions within individual arrays further show that exchange can occur within active αSat itself.

Additionally, in three samples, we identified and verified in short-read datasets by specific kmers an approximately 250 kbp insertion of chromosome 10 active αSat sequence and neighboring HORs into the chromosome 19 active array (**Supplementary Note 1**). This event may represent an example of inter-array seeding, previously proposed as a mechanism of αSat evolution in which sequence originating from one array is introduced into another and subsequently expands^32,36,39^. Collectively, these observations point to multiple routes of sequence exchange between distinct αSat arrays of nonhomologous chromosomes.

### Sequence context of functional centromeres

Although centromere identity is epigenetically specified, the underlying αSat sequence may influence where centromeric chromatin is organized within the array^7,23,40^. The histone H3 variant CENP-A specifies the site of kinetochore assembly and is a defining feature of functional centromeric chromatin, whereas CENP-B is a sequence-specific centromeric protein that binds 17-bp CENP-B-box motifs within αSat DNA. Centromere dip regions (CDRs) are hypomethylated regions that closely correspond to CENP-A occupancy, providing a proxy for functional centromere position. We therefore used CDRs to ask whether functional centromeres preferentially occupy particular sequence contexts within active αSat arrays. CDRs often contain multiple closely spaced hypomethylated subdomains (sub-CDRs; **Fig. 2a**). We mapped CDRs from ONT methylation profiles and their underlying sequence compared with the remainder of each active array (<u>Methods</u>, **Supplementary Fig. 19**). To characterize the sequence context surrounding CDRs, we annotated αSat HOR organization, CENP-B boxes and local sequence identity across each active array (**Fig. 2a**). Local sequence identity was calculated from pairwise sequence identities between neighboring 5 kbp windows, providing a measure of local αSat sequence homogeneity (<u>Methods</u>). Across the majority of haplotypes (6,766 centromeres; 97.4% of QC-pass centromeres), CDRs occur in regions of higher local sequence identity significantly more often than the surrounding active array (99.3% versus 99.0%, one-sided Wilcoxon signed-rank test, P < 10⁻⁴; **Fig. 2b**), supporting previous observations that CENP-A preferentially occupies younger, more homogeneous regions of αSat arrays^7,23,40^. Across haplotypes, CDRs are also significantly enriched for CENP-B boxes (one-sided Wilcoxon signed-rank test, P < 10⁻⁴), 17-bp motifs that bind CENP-B and contribute to centromeric nucleosome organization, with densities of 2.60 motifs per kbp within CDRs compared with 2.52 motifs per kbp elsewhere in the active array (**Fig. 2b**).

**Figure 2.**
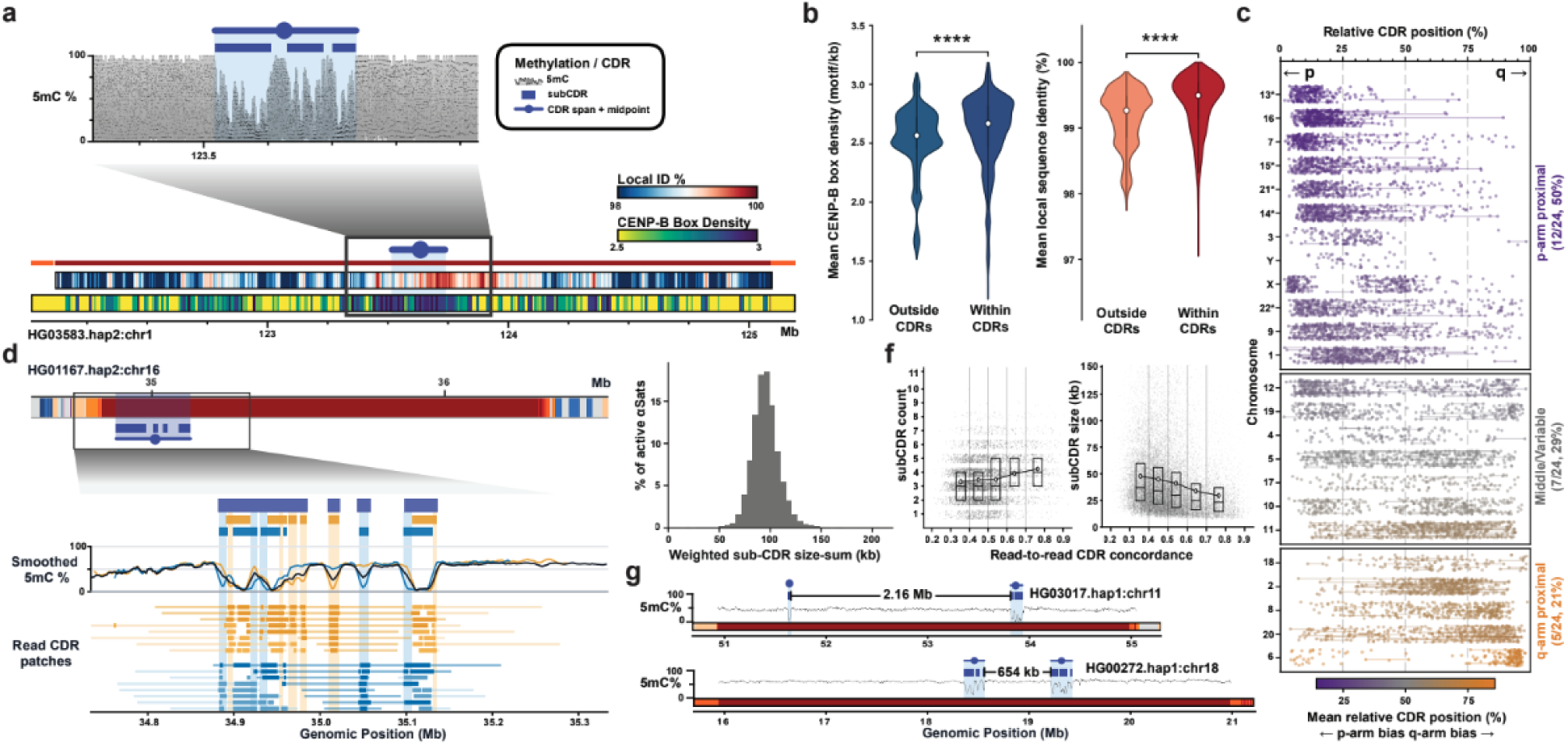
Sequence context, position and organization of human centromere dip regions. **a**, Example active αSat array showing ONT-derived 5mC methylation, CDR and subCDR sites, together with CenSat annotation, local sequence identity and CENP-B-box density. The CDR midpoint used to determine relative CDR position is indicated. **b**, Distributions of mean local identity and CENP-B-box motif density within CDRs compared with regions outside of CDRs in the same active-αSat, with noted p-value (****) < 10⁻⁴, one-sided Wilcoxon signed-rank test. **c,** Relative CDR positions across active αSat arrays. Chromosomes are grouped according to p-proximal, middle/variable, or q-proximal CDR positioning and colored according to mean relative CDR position. **d,** Example centromere array (chromosome 16; HG01167.hap2) showing read-to-read-heterogeneity in CDR positioning. Individual ONT reads separate into two clusters supporting distinct hypomethylated configurations; aggregate methylation profiles and sub-CDR calls are shown for all reads and for each read cluster. **e,** Distribution of read-agreement-weighted summed sub-CDR length across all active arrays. **f,** Number (left) and size (right) of sub-CDRs as a function of read-to-read CDR concordance. **g,** Examples of sub-CDR annotated active arrays with unusually wide sub-CDRs on chromosomes 11 and 18. Distances indicate the separation between sub-CDR domains.

While increased sequence identity at CDRs was observed across all 24 chromosomes, CENP-B-box enrichment is more chromosome-dependent; chromosome Y, which lacks CENP-B boxes, was excluded from this analysis (**Supplementary Figs. 20 and 21**). Notably, chromosomes X and 11 have comparatively low CENP-B-box densities overall but retain higher densities within CDRs than in the surrounding active array. CENP-B-box density also differs substantially between chromosomes but is relatively consistent among haplotypes of the same chromosome (**Supplementary Fig. 22**). Because individual monomers within an αSat HOR differ in whether they contain a CENP-B box, changes in StV HOR composition through rearrangement, expansion and contraction can reshape local CENP-B-box density without altering the HOR monomers themselves. High local identity and CENP-B box density are also correlated within active αSat arrays, linking CDR position to a broader sequence context rather than to either feature independently (**Supplementary Fig. 23**).

Beyond their local sequence context, we asked whether CDRs occupy characteristic positions within active αSat arrays. For CDRs containing multiple sub-CDRs, we defined the CDR position as the midpoint of the sub-CDR cluster. To compare arrays of different sizes, this position was normalized from 0% (p-proximal) to 100% (q-proximal). CDR position within active αSat arrays varies across chromosomes but shows chromosome-specific patterns (**Fig. 2c**). Half of the chromosomes (12 of 24) show a p-proximal bias, compared with five showing a q-proximal bias and seven with CDRs predominantly near the center of the array. Notably, all five acrocentric chromosomes show a p-proximal bias. This shared positioning may reflect the unusual evolutionary dynamics of acrocentric chromosomes. Acrocentric short arms contain homologous rDNA and satellite sequences that undergo interchromosomal recombination, and their centromeric arrays can contain closely related αSat HORs^7,37^.

### Organization and variability of centromere dip regions

We next examined CDR organization across human centromeres and used single-molecule methylation profiles from individual sequencing reads to assess how consistently CDRs are positioned within the αSat array. CDR positioning varies among individual reads, such that discrete aggregate sub-CDRs reflect consistent positioning, whereas broader, shallower domains reflect greater positional variability at the single-molecule level (**Fig. 2d**; **Methods**; **Supplementary Fig. 24 and 25**). CDR positional variability tends to be similar among centromeres from the same individual, with approximately 74% of the variance attributable to differences between individuals (**Methods**; **Supplementary Fig. 26**). Concordance is only weakly associated with array methylation, mapping quality and sequencing coverage (**Supplementary Fig. 27**). Using a read agreement weighted sum to account for positional variability, the total length of sub-CDRs followed a unimodal distribution centred at approximately 90 kbp per centromere, even as sub-CDR organization varied (**Fig. 2e**; **Methods**). Centromere haplotypes from samples with more consistent inter-read CDR positioning contain more numerous, smaller, and more discrete sub-CDRs, (**Fig. 2f**), suggesting a source of technical noise that limits the resolution of sub-CDR calls in some samples. Sub-CDRs are typically clustered, although nine centromeres contain domains separated by more than 500 kbp, including one with a separation of 2.16 Mbp (**Fig. 2g**; **Supplementary Fig. 28**). Despite this extreme variation in sub-CDR spacing, nearly all of these centromeres maintained a similar summed sub-CDR length (**Supplementary Fig. 28**). Together, these findings reveal that the total extent of centromere-associated hypomethylation, measured as summed sub-CDR length, is remarkably consistent across αSat arrays and chromosomes and is maintained across individuals and across diverse human haplotypes despite variation in sub-CDR organization.

### Identifying and representing active array variation with Centrolign

Fine grained analysis of active array variation requires different methodologies than macro-scale analysis of structural haplotypes. Assembled arrays must be aligned to uncover such variation, but this poses significant technical challenges. Conventional alignment methods that maximize sequence identity are unsuited to these sequences. The existence of many near-identical copies of the repeat provides bait for misalignment^41^, and this problem is compounded by prevalent large structural variants, which can make orthologous sequences have lower identity with each other than with nearby paralogous repeats. To address these limitations, we developed Centrolign, an accurate multiple sequence alignment (MSA) algorithm for active arrays. Centrolign uses the strategy of uniqueness alignment pioneered by UniAligner^42^ in which matching sequences are scored by their rarity across the input sequences (**Fig 3a**). We believe that, in effect, this prioritizes the alignments that include the most confident orthology.

**Figure 3.**
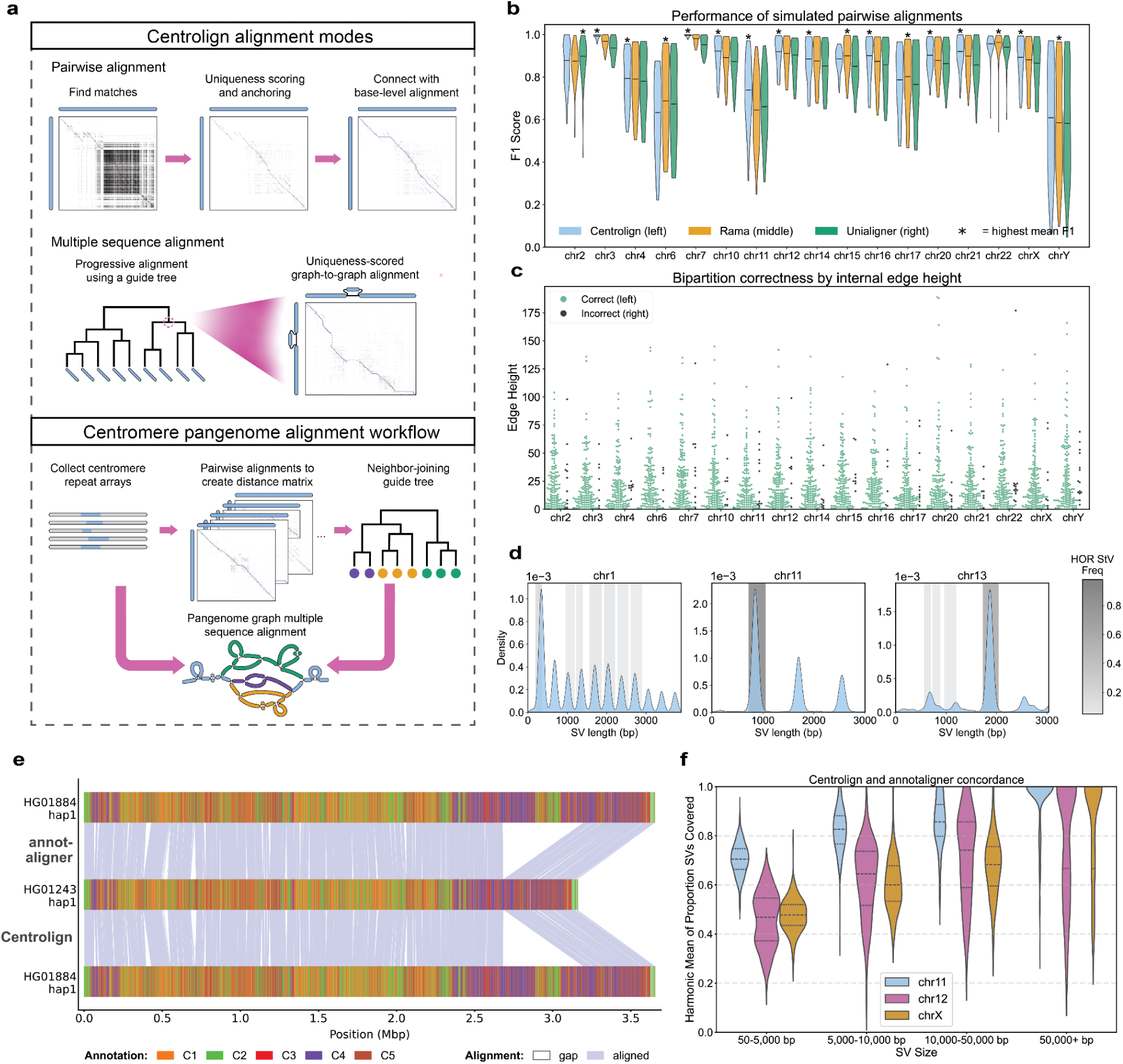
Identifying and representing active array variation with Centrolign. **a,** Top subpanel shows a schematic of the two alignment modes of Centrolign (pairwise and multiple sequence alignment). Bottom subpanel describes how both modes are used to generate a pangenome for human centromeres. **b,** F1 score of pairwise alignments on simulated data from Centrolign (left,blue), Rama (middle,orange), and Unialigner (right,green). A star indicates the aligner with the highest mean F1 score for each chromosome. **c,** Concordance between the tree inferred by the MSA of simulated sequences and the ground truth tree. Correctness of a bipartition indicates whether a bipartition of leaves induced by removing an internal edge in the tree is shared with the ground truth tree. Each chromosome has two swarm plots: left/green indicate correct bipartitions, right/black indicate incorrect bipartitions. Y-axis shows the height of the associated internal edge. **d,** SV length distributions (light blue bars) for pairwise alignments induced from the HPRC2 Centrolign graphs for chromosomes 1, 11, and 13. Grey shaded bars indicate the size range of observed HOR StV counts from CenSat annotations in HPRC release 2, with darker shading indicating higher frequency. Only StVs observed at > 5% frequency were included. **e,** Visualization of the alignment concordance for a pairwise alignment produced by annotaligner (top) and Centrolign (bottom) between two chromosome 11 active arrays (HG01884 haplotype 1 and HG01245 haplotype 1). Colored regions indicate the HORhap annotations for each array, with purple shaded lines between them representing aligned sequence. White spaces indicate gaps. **f,** Harmonic mean of the proportion of Centrolign SVs covered by annotaligner and vice versa. SVs are broken by size category on the x-axis, with the left-most (blue) violin showing the distribution for chromosome 11, middle (magenta) showing chromosome 12, and right (orange) chromosome X.

Centrolign also uses the graph-based progressive MSA framework of partial order-partial order alignment (POPOA)^43^ (**Fig 3a**; **Methods**). Pairwise alignment is also available as a special case. The POPOA strategy has two main benefits. First, it can gracefully handle large indels without penalizing them excessively by representing them as alternate paths in the graph. This promotes accurate alignment in the presence of large structural variants, such as those discussed above. Second, the final alignment is structured as a pangenome graph, and it can therefore feed directly into downstream pangenome graph methodologies. Accordingly, these alignments could also address the lack of centromere representation in the HPRC pangenome alignments.

### Centrolign validation on simulated data

To be confident in the variation identified by Centrolign, we validated it using multiple approaches. First, we developed a multiple active array sequence simulation in which the ground-truth alignment is known. This tool starts with a real active array sequence and modifies it into multiple sequences, according to a simulated genealogical tree (**Methods**). The simulation is not intended to accurately reflect processes of sequence change in active arrays. However, it loosely matches qualitative features that we have observed when comparing real active arrays, such as maintenance of the HOR register, occasional HOR StVs, frequent indels of HOR units, and occasional massive deletions or duplications.

Using the simulation, we benchmarked pairwise alignments from Centrolign to the comparable alignment tools for tandem repeats: UniAligner and Rama. We simulated 60 repetitions of 6 sequences for different levels of sequence divergence. Each tool’s alignments were compared to the ground-truth alignment to measure accuracy (Methods). While all three aligners show comparable accuracy, Centrolign has the highest mean F1 score for 11/17 chromosomes tested **(Fig. 3b and Supplementary Table 5).**

For multiple sequence alignments, there are no comparable algorithms to benchmark against. Instead, we assessed the ability of Centrolign’s MSAs to capture information on evolutionary relationships. We aligned 30 repetitions of 8 simulated sequences. From the MSA, we inferred a tree using the neighbor-joining algorithm ^44^ and compared it to the true tree the sequences were simulated from. The distance matrix for the neighbor-joining algorithm was constructed by inducing pairwise alignments and using them to compute a metric that we term “pairwise match distance” (2 * mismatches / sum of sequence lengths). We assessed concordance to the true simulated tree by comparing the set of bipartitions of leaves induced by removing each internal edge in the trees (similarly to the Robinson-Foulds distance^45^). 94% of internal edges’ bipartitions were correctly recapitulated, and errors were disproportionately in deep nodes of the tree **(Fig. 3c, and Supplementary Table 6.)**.

### Construction and validation of active array MSAs for HPRC2 assemblies

For each chromosome, we built MSA graphs for all of the HPRC2 assemblies that passed quality control (Methods). The guide trees for progressive MSA were constructed in a similar manner as in the simulation. We computed pairwise match distance matrices from direct pairwise Centrolign alignments and then applied the neighbor-joining algorithm. **(Fig. 3a)**.

We assessed the consistency of pairwise alignments induced from the HPRC2 graphs and direct pairwise alignments. We found that Centrolign’s induced and direct alignments are tightly concordant where pairwise match distances are less than 20% **(Supplementary Fig. 29).** As pairwise match distance increases, Centrolign’s induced alignments have larger indels and a higher distance relative to the direct alignments. This may reflect the progressive alignment strategy Centrolign uses, where suboptimal indel placements introduced early in the MSA can propagate through subsequent alignments **(Supplementary Fig. 30)**.

To establish the biological plausibility of Centrolign alignments, we contextualized the alignments with the HOR structure of the sequences. From the HPRC2 CenSat annotations, we extracted all HOR-StVs above 5% frequency for each chromosome and visualized them alongside the size distribution of Centrolign structural variants (SVs) from induced pairwise alignments. We found that the majority of Centrolign SV sizes coincided strongly with multiples of the observed HOR StV sizes **(Fig. 3d, Supplementary Fig. 31)**. This finding aligns with previously published work highlighting expansions and contractions of entire HOR units in the active arrays ^7^.

To further validate our Centrolign alignments, we checked whether large scale structural variations (SVs) reported by Centrolign were consistent with those found in an orthogonal approach, Annotaligner, which implements a global alignment between HORhap annotations **(Fig. 3e**; **Methods).** Because SV placement and representation can be uncertain among nearly identical HORs (especially for the lower-resolution strategy of Annotaligner), we looked for approximate matches between SVs in the two tools, allowing for some discrepancy in the size, placement, and fragmentation (**Methods**). Because SV representations are not one-to-one, we refer to an SV being “covered” as opposed to matched. We observed that Centrolign calls more events than Annotaligner **(Supplementary Fig. 32 and 33)**, which is expected due to the increased precision of a base-level alignment. Large events are consistent between both methods: 89% of SVs > 10 kbp are reciprocally covered **(Fig. 3f, Supplementary Table 8).** Although concordance decreases with size, 57% of SVs are still covered for events < 5 kbp, supporting the accuracy of Centrolign’s SV callset.

### Characterizing alpha satellite variation

The HPRC2 Centrolign pangenome graphs we constructed provide a powerful dataset for studying the mutational properties of αSat arrays. However, there is a limit in the evolutionary distances over which individual mutational events can be reliably resolved. Arrays sharing recent ancestry within a cenhap lineage also share similarities in sequence composition and structure, but extensive turnover between distant lineages, reflected in the differences in their HORhap composition and HOR-StVs, make direct comparison uncertain or impossible. This pattern is reflected in the Centrolign graphs, in which cenhaps occupy largely separate “lobes” that converge primarily at the edges of the active array, where older αSat layers retain sequence shared across deeper evolutionary timescales (**Fig. 4a, b**). This places limits on our ability to resolve variants, many of which are likely lost to evolutionary turnover and/or population bottlenecks. We therefore restricted subsequent analyses, unless otherwise stated, to pairwise alignments from the Centrolign MSA with a match distance < 0.2. Notably, only 42.5% of arrays had more than one other array with a pairwise match distance < 0.2 (**Fig. 4b**; **Supplementary Fig. 34)**.

**Figure 4.**
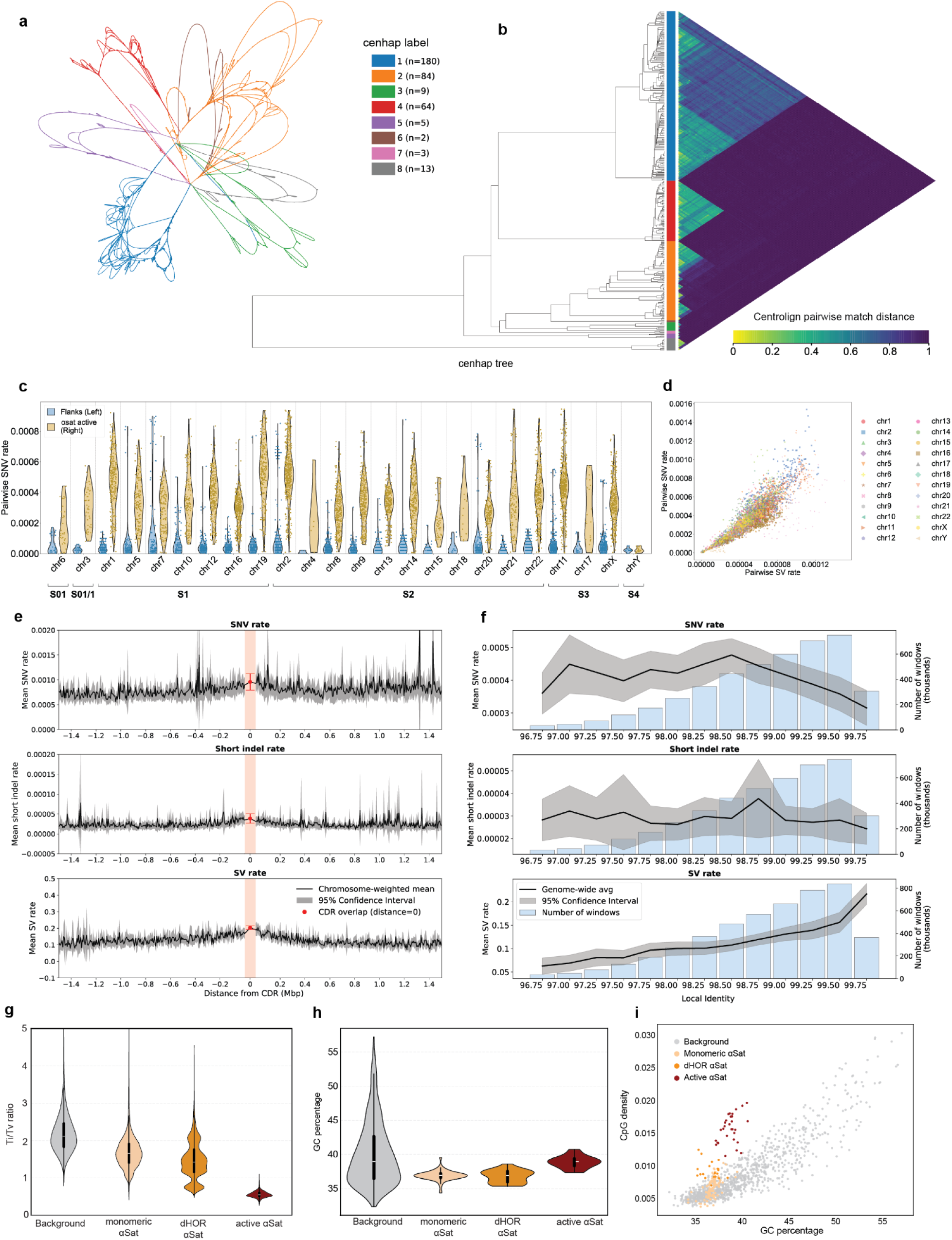
Centrolign enables characterization of alpha satellite variation. **a,** Centrolign MSA of HPRC2 chromosome 12, subset to up to 20 samples per cenhap for clarity (93 haplotypes were included in total), with paths colored by cenhap (Methods). Visualized with Bandage ^46^. **b,** Cenhap tree for all of HPRC2 chromosome 12, with colors indicating cenhap for each sample. Heatmap to the right of the tree indicates the Centrolign pairwise match distance between each sample pair. **c,** Pairwise SNV rate for 50 kbp up- and down-stream of the active αSat arrays (flanks) is shown in blue, in the lefthand violins and was derived from HPRC2 Minigraph-Cactus pangenome graphs, while SNV rates for the αSat (active) arrays are shown in yellow, in the righthand violins, for each chromosome (grouped by suprachromosomal family). **d,** Pairwise SNV rate (y-axis) versus pairwise SV rate (x-axis) in the αSats, for each chromosome. **e,** Variation rates calculated across each array in 10 kbp windows for SNVs and short indels, and in 100 kbp windows for SVs, then binned into 500 bins up to 1.5 Mbp away from the CDR. Black line indicates chromosome-weighted mean, and grey indicates 95% confidence intervals. Mean rates for windows overlapping the CDR are shown in the 0 distance bin in red. **f,** Active array variation rates across all chromosomes calculated within 5 kbp windows from local identity annotations are binned and plotted by local identity in black, with 95% confidence interval in grey. Blue histograms represent the number of windows in each bin. **g**, Ti/Tv ratio, **h.** Percentage GC sequence content and **i.** GC content (x axis) vs CpG density (y axis) for all chromosomes in background genomic regions (non-centromere), αSat monomeric, αSat diverged, and αSat active.

Using Centrolign, we observed approximately sixfold more single nucleotide variants (SNVs) within active αSat arrays than in the immediately flanking 50 kbp sequence (one per 2,609 bp versus one per 15,817 bp; **Fig. 4c, Supplementary Fig. 36**), consistent with previous studies^7,13,40,47^. The low magnitude of these rates is expected given that we restricted our study to closely related pairs; on more distantly related samples the flank SNV rate approaches the genome-wide expectation of ∼1 in 1000 bp **(Supplementary Fig. 37)**. The elevated diversity highlights the active array as a distinct evolutionary environment, where tandem-repeat organization and homogenizing processes may shape the rapid accumulation of sequence variation. SNV variation differs by more than threefold among chromosomes, with mean SNV density ranging from one per 5,488 bp on chromosome 6 to one per 1,740 bp on chromosome 2 (**Supplementary Fig. 38**; **Supplementary Table 9)**. These chromosome-specific differences persisted after accounting for evolutionary distance, measured as the patristic distance separating them in the guide tree. This suggests that αSat arrays evolve at different rates across chromosomes. SNVs, short indels, and SVs each increase with evolutionary distance between arrays **(Fig 4d, Supplementary Fig. 39 and 40)**. However, the relative accumulation of SNVs and SVs differ among some cenhap lineages **(Supplementary Fig. 41**). Despite differences in overall variation, active arrays maintain a broadly similar balance of variant classes across chromosomes, averaging six SNVs per SV **(Supplementary Fig. 42**). This ratio differs markedly from the rest of the genome, which has 513 SNVs per SV, corresponding to an ∼85-fold enrichment of structural variation relative to SNVs in active αSat (**Supplementary Table 9**). The strong bias toward structural variation highlights repeated remodeling of HOR organization as a major mode of active αSat evolution.

We next asked whether variation is distributed uniformly across active arrays or associated with the position of the functional centromere. In aggregate across chromosomes, all three variant classes are elevated within CDRs, and elevated variation was also observed in the 250 kbp regions flanking CDRs (**Fig. 4e**). SVs are significantly enriched within CDRs on every chromosome, while SNVs and short indels are enriched on 14 and 13 chromosomes, respectively (permutation test, α = 0.01, 10,000 iterations) (**Supplementary Fig. 43 and 44**). Because CDRs coincide with more homogeneous regions of the active array (**Fig. 2b**), we also stratified variation by local sequence identity, a measure of HOR homogenization. In the most homogeneous regions, SNV rates decrease while SV rates increase, and short indel rates remain relatively constant (**Fig. 4f**). This relationship is consistent with recent structural turnover of αSat, including expansions and contractions that generate homogeneous repeat blocks with less time to accumulate SNVs. SV enrichment in the most homogeneous regions is significant on every chromosome, while SNV depletion is significant on 17 chromosomes (permutation test, α = 0.01, 10,000 iterations). Chromosome 11 is a notable exception, with all three variant classes enriched in highly homogeneous regions (**Supplementary Fig. 43)**. Restricting the analysis to closely related arrays representing the most confident alignments (pairwise match distance < 0.1) supported the enrichment of SVs and depletion of SNVs in highly homogeneous regions (**Supplementary Fig. 45**). The significance of the enrichments of both CDR and high-identity regions was maintained after excluding overlaps with the other, showing that the two associations are partially separable (**Supplementary Fig. 43**). This separation suggests that elevated variation at CDRs cannot be explained solely by their occurrence in highly homogeneous αSat, which instead shows increased structural variation and reduced SNV accumulation.

SNVs in the euchromatic genome show a characteristic transition bias, with approximately two transitions per transversion^48^. In contrast, active αSat arrays had a Ti/Tv ratio of only 0.55, close to the 0.5 expected without transition bias (**Fig. 4g**). We repeated these comparisons using UniAligner, Rama, and minimap2, and found all three methods recovered similarly low Ti/Tv ratios across chromosomes, indicating that the distinct substitution spectrum was not specific to Centrolign (**Supplementary Fig. 46**). In addition, the low Ti/Tv ratios hold when excluding SNVs within 20 bp of an indel, providing confidence these results are not caused by alignment artifacts^49^. They also remain consistent when restricting to the most closely related arrays (pairwise match distance < 0.1), where alignments are more confident (**Supplementary Fig. 46**). The markedly reduced transition bias suggests that rapid sequence turnover in αSat is shaped by processes that differ from those underlying the characteristic genome-wide substitution spectrum. Analysis of non-αSat embedded within the Chromosome 3 active array show similarly low Ti/Tv ratios for HSat1A (**Supplementary Fig. 47**). Comparisons across additional satellite families will be important for determining which features of this substitution spectrum reflect αSat evolution and which are shared more broadly across other centromeric satellites.

Substitution spectra reflect both the processes acting on DNA and the underlying sequence composition. Active αSat arrays are known to differ in sequence composition from the euchromatic genome, raising the possibility that sequence context contributes to their low Ti/Tv ratio. When we compared Centrolign calls in active arrays of alpha satellites against variant calls from other alpha satellite arrays (Methods) we found that Ti/Tv increases progressively from active arrays through divergent and monomeric αSat, a trend observed across chromosomes (**Fig. 4g** and **Supplementary Fig. 48**). The transversion-rich spectrum is therefore strongest in active αSat and diminishes in older, more divergent αSat sequences. One possible compositional explanation is variation in CpG abundance, because CpG deamination is a major source of C→T transitions in the human genome^50^. The repeated CENP-B boxes within active αSat arrays provide a rich source of CpGs, resulting in higher CpG density than genomic regions with comparable GC content (Fig. 4h–i). CpG scarcity therefore cannot explain the low Ti/Tv ratio. To account more broadly for sequence composition, we estimated the Ti/Tv expected by weighting genome-wide trinucleotide-specific substitution frequencies by the trinucleotide composition of active αSat (**Supplementary Fig. 49)**. This analysis predicted a Ti/Tv ratio of 2.3, nearly fourfold higher than the observed ratio of 0.55. Sequence composition cannot explain the low Ti/Tv ratio of active αSat. This distinct substitution spectrum may reflect other properties of these arrays, including higher methylation levels^9^ and late replication timing^51^.

### Using Centrolign graphs as a centromere pangenome reference

The extensive diversity of active αSat presents a challenge for using any single centromere sequence as a reference. Centrolign pangenome graphs provide a potential alternative for genotyping and read alignment across diverse centromeres. Notably, cenhap assignments depend both on the haplotypes sampled and on the resolution at which lineages are delineated. We therefore asked whether the lineages identified in our current sample could be reliably recovered from unaligned HiFi reads. We used the vg haplotypes tool^52^ to identify graph haplotypes with k-mer content similar to those in each sample’s HiFi reads. We found that the highest-scoring haplotypes from vg haplotypes frequently corresponded to the sample’s nearest neighbor haplotype (based on Centrolign pairwise match distance) (**Fig. 5a and Supplementary Fig. 50**). Assigning each sample the cenhap of its top-scoring haplotype correctly classified ∼95% of haploid read sets (**Fig. 5b and Supplementary Table 10**). For diploid read sets, cenhap genotyping correctly typed ∼90% of individual haplotypes and ∼80% of diplotypes on most chromosomes (**Supplementary Table 10 and Supplementary Fig. 52**).

**Figure 5.**
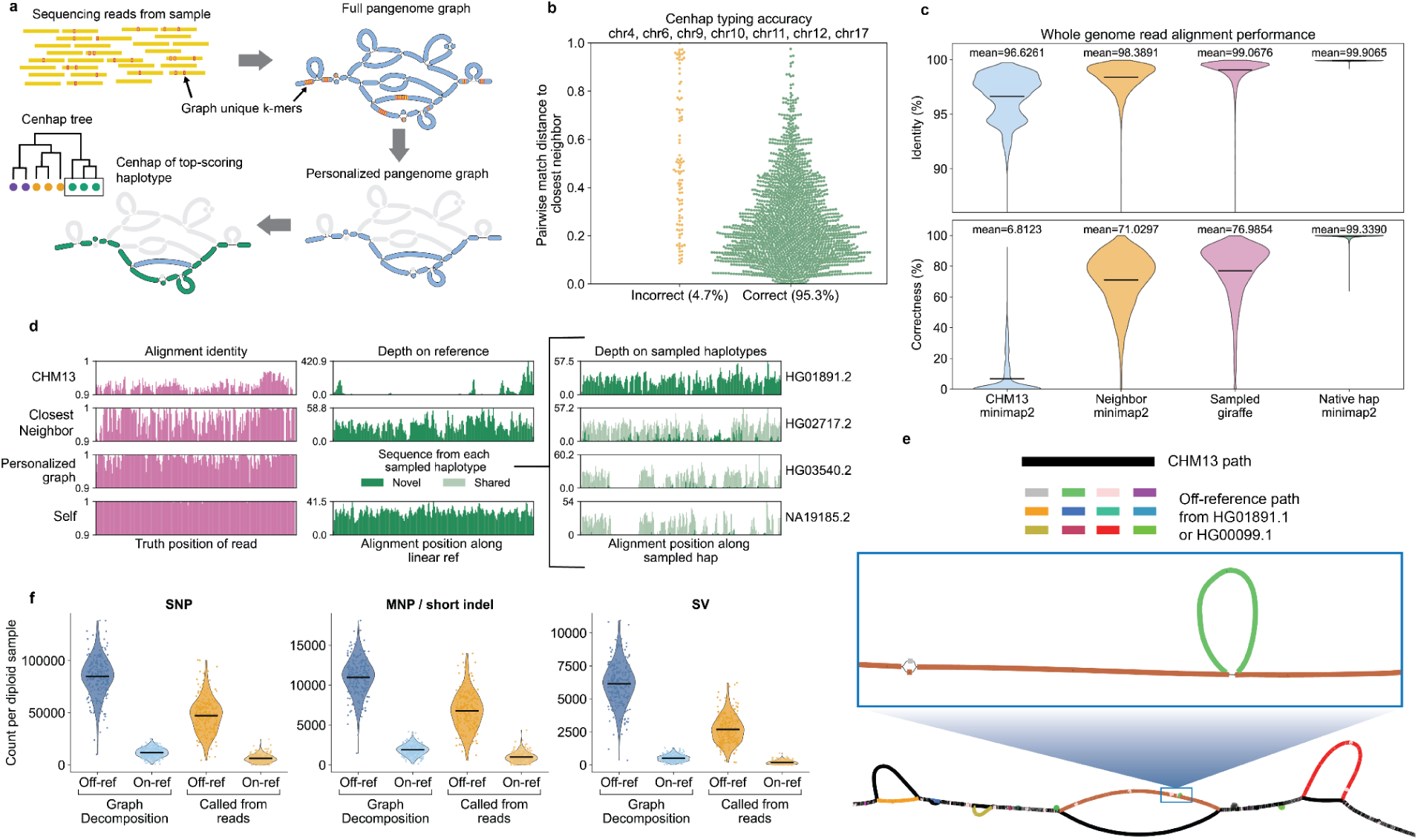
Using Centrolign graphs as a centromere pangenome reference. **a,** Diagram of cenhap typing procedure. Graph-unique k-mers are identified within the read set, matched against haplotypes in the graph, and then used to select a subset of haplotypes; the top such haplotype is used to type the reads. **b**, Cenhap typing results. Left side is haplotypes typed incorrectly, right side is haplotypes typed correctly. Y-axis is minimum pairwise match distance to another haplotype. **c**, Average read alignment identity (top) and average read (bottom) correctness, using PacBio HiFi reads, for alignments to CHM13 with minimap2, the most closely related haplotype (Neighbor minimap2), a personalized pangenome graph reference (Sampled giraffe), and the sample’s own assembly (Native hap minimap2). Black bars indicate means. Alignments to linear references done with minimap2; alignments to graph references done with Giraffe. **d,** Detailed analysis of read alignments for haplotype HG01106.1 (self). Left (pink) column indicates average identity for reads which correspond to certain truth positions, when they are aligned to various references. Middle (dark green) column indicates average read coverage for alignments to certain positions on those references. Right column indicates the same but for the individual haplotypes present in the personalized pangenome graph, ordered from first to last selected during personalization. Depth on nodes that were included in previously sampled haplotypes is shown in light green instead of dark green. **e,** Illustration of off-reference variation as represented with pangenome coordinates on a chromosome 12 centromere subgraph. CHM13 is represented in black, and the additional pangenome coordinate intervals in other colors. Visualized with Bandage. **f,** Total number of variants per haplotype over all chromosome graphs, divided into those that can be expressed relative CHM13 (on-reference) and those that require additional pangenome coordinates (off-reference). Variants decomposed from the graph with vg-deconstruct are in the two leftmost violin plots, while variants genotyped from PacBio HiFi reads with vg-call are the two rightmost violin plots.

Because the complexity of the full HPRC2 centromere graphs made indexing them for read alignment computationally infeasible, we instead used haplotypes identified from each read set to construct personalized subgraphs for alignment with vg Giraffe^53^. For comparison, we also aligned reads with Minimap2^54^ to either CHM13^8^ or the nearest-neighbor haplotype by pairwise match distance. Both personalized approaches (aligning to the personalized graph or the nearest-neighbor) substantially increased alignment identity relative to CHM13, demonstrating the benefit of using centromere sequences more closely related to the sample (**Fig. 5c**, **Supplementary Table 11; Supplementary Fig. 53 and 54**). The personalized graph also improved alignment fidelity over the nearest-neighbor linear reference across chromosomes (**Supplementary Fig. 55**) and haplotypes (**Supplementary Fig. 56)**. These benefits persisted with diploid read sets (**Supplementary Fig. 53 and Supplementary Table 12**). Unlike a single nearest-neighbor reference, personalized graphs can represent sequence diversity distributed across multiple related haplotypes. Reads from regions poorly represented by any single neighbor could instead align to alternative sequences in the personalized graph, improving both alignment identity and concordance with truth positions at individual loci (**Fig. 5d and Supplementary Fig. 57**). The large collection of diverse centromere assemblies in HPRC2 increases the likelihood that sequence related to a new sample is represented in the pangenome, enabling personalized graphs to be constructed directly from unaligned reads without first assembling the centromere.

Mapping reads to personalized centromere graphs addresses reference divergence, but comparing and calling variation across samples also requires a common coordinate system. Variation nested within large haplotype-specific insertions cannot be consistently represented relative to a linear reference such as CHM13, which lacks the sequence needed to anchor these variants. We therefore developed Graph Reference (GRef) coordinates^14^, which use a set of haplotype sequences spanning the pangenome graph to assign canonical positions across its sequence diversity. On chromosome 12, for example, variants nested within large insertions relative to CHM13 are assigned consistent off-reference positions that can be shared across samples (**Fig. 5e**). GRef coordinates retain compatibility with standard variant formats such as VCF^55^. GRef coordinates represented approximately an order of magnitude more sites of variation than CHM13 coordinates (**Fig. 5f, Supplementary Fig. 60**), largely by assigning positions to variation nested within non-reference sequence. GRef coordinates also supported variant calling directly from mapped long reads, with vg call recovering similar numbers of variants to direct variants implied by the haplotypes’ paths through the graph (**Fig. 5f, Supplementary Fig. 60**). Centromere variation can therefore be mapped, called, and compared across individuals without requiring either a pre-existing centromere assembly or a single universal reference sequence.

## Discussion

Complete assemblies spanning human centromeric and pericentromeric satellites are providing increasingly detailed catalogs of sequence, structural and epigenetic variation^7,13,16^. Here, analysis of more than 6,000 complete HPRC2 centromeric assemblies^14^ enables us to characterize this variation across scales, from individual bases and repeat organization to megabase-scale structural differences. Our analyses identify recurrent structural configurations extending across pericentromeric and centromeric regions, where megabase-scale rearrangements reorganize satellite arrays together with intervening sequence. These structural differences are associated with differences in gene content and, in some cases, gene expression. Understanding how this variation arises and evolves depends on relating differences among haplotypes to their shared evolutionary history. Despite substantial turnover, satellite sequence and HOR structure remain associated with cenhap ancestry, providing an evolutionary context for understanding their change over time. Recombination and interchromosomal exchange introduce departures from these lineage-associated patterns^7,13^, connecting otherwise distinct evolutionary histories. Using Centrolign, we establish multiple sequence alignments that define correspondence among these diverse centromeric haplotypes. These alignments bring regions that have remained largely outside pangenomic analysis^56^ into a consistent representation of human genetic variation and provide a route for their integration with the broader human pangenome. Establishing this correspondence addresses a longstanding gap in human genomics, allowing centromeric variation to move from assembly-by-assembly description toward systematic identification, representation and genotyping across individuals. In turn, centromere biology can move beyond comparisons among finished genomes toward genetic analyses across cohorts, enabling direct tests of how satellite variation shapes centromere identity, function, and chromosome inheritance.

Aligning related αSat arrays makes it possible to directly compare how αSats diverge over evolutionary time. We find that SVs are strongly enriched relative to SNVs, with insertions and deletions frequently occurring in complete HOR units. This excess may in part reflect the recent expansion and contraction of αSat sequence, which generates young repeat blocks with relatively little time to accumulate nucleotide substitutions. The tendency for indels to involve complete HORs may reflect exchange between highly similar repeat copies, with homology between HORs naturally favoring expansions and contractions in complete repeat units^57^. The nucleotide substitutions that accumulate are also unusual, with a Ti/Tv ratio of approximately 0.55 compared with ∼2.3 expected from αSat trinucleotide composition under the genome-wide substitution spectrum. Although increased proportions of transversions have been observed in particular genomic and cellular context ^51,58,59^, they are uncommon relative to the genome-wide pattern^60^. Within αSat, this signature is strongest in active arrays and becomes progressively less pronounced in older αSat sequences. The processes responsible remain unclear. Properties of active αSat arrays, including their high and regular CpG methylation and late replication timing, could contribute to this distinct substitution spectrum^9,51^. Gene conversion could alter the observed spectrum through repeated sequence homogenization^57^, while replication or repair processes operating differently in these regions could also contribute^61^. Because our substitution analyses are restricted to αSat, whether similar patterns extend to other centromeric and pericentromeric satellites remains unknown.

Distinct evolutionary patterns are also associated with where the functional centromere is positioned within the αSat array. Across thousands of haplotypes, CDRs preferentially occupy young, homogeneous αSat enriched for intact CENP-B boxes, extending previously observed associations between centromere function and young αSat across diverse human haplotypes^9,13,23^. Despite substantial variation in CDR position and organization, their summed extent remains centered near 90 kbp, suggesting that the amount of hypomethylated centromeric chromatin may be more constrained than its location within the array. In addition, with Centrolign we find significant enrichment for all types of variation within and surrounding the CDR, indicating the sequence environment favored by the functional kinetochore is among the most rapidly remodeled portions of the array. While the CDR tends to associate with the most homogeneous portions of the array, these regions separately display significant depletion of SNVs and enrichment of SVs. Together, these observations support a dynamic view of αSat evolution^7^ in which the youngest, most homogeneous sequence is not simply a record of recent array expansion but continues to undergo structural remodeling while associated with the functional centromere. This relationship is consistent with models proposing that centromere function and satellite renewal are coupled, but does not distinguish whether centromere activity promotes variation, preferentially occurs in recently expanded sequence, or both arise from other properties of young αSat^7,32^. By defining genetic and epigenetic signatures of this process across thousands of haplotypes, our results provide a basis for testing how the continual evolution of centromeric DNA interacts with centromere identity and function.

Until now, fine-scale characterization of centromeric satellite variation has largely depended on complete assembly of these regions. The pangenome provides a means to genotype this variation directly from sequencing reads. Long reads from samples outside the pangenome can be mapped to related αSat haplotypes, assigned to cenhap lineages, and used to identify and report fine-scale variation in standard variant formats. This is particularly important in active αSat, where structural differences between haplotypes create sequences that are absent from any single linear reference. Variants within these haplotype-specific sequences therefore cannot be represented using that reference. Graph Reference coordinates represent approximately an order of magnitude more variant sites than CHM13 coordinates alone, including variants nested within haplotype-specific sequences. Extending these approaches across human diversity will depend on increasing the number and diversity of complete centromeric haplotypes and integrating their representation with whole-genome pangenome resources. At present, the Centrolign centromere graphs remain separate from the primary HPRC pangenome graphs, and unifying these representations will be necessary to support routine whole-genome pangenomic analysis of centromeric variation. Our ability to genotype cenhap lineages from short k-mers indicates that existing sequencing datasets already contain retrievable information about centromeric haplotype diversity. An integrated pangenome reference should allow the routine extraction of this information, although the extent to which this improves fine-scale mapping and genotyping remains to be determined. Only 42.5% of active αSat arrays currently have another assembled array within the threshold used for confident whole-array alignment, indicating that many centromeres still lack the closely related representation needed for fine-scale comparison. Ongoing HPRC efforts to increase complete genome representation from hundreds to thousands of haplotypes should progressively fill these gaps and expand the reach of pangenome-based genotyping. As these resources expand and become integrated with the broader human pangenome, centromeric satellite variation can be studied alongside genetic variation across the rest of the genome. By providing shared representations and standardized coordinates for centromeric satellite variation, this work moves these regions beyond assembly-specific description and into an interoperable form of human genetic variation that can be called, reported, and analyzed alongside the rest of the genome.

## Competing interests

UCSC has received cloud compute credits from Amazon Web Services (AWS) to support the HPRC, and the HPRC data receives free hosting from AWS, whose cloud services are used in this work. J.E. is now employed by Roche Sequencing Solutions. All other authors declare that they have no competing interests.

## Methods

### Data availability

Assemblies are available in Genbank Bioproject PRJNA730822. Sequencing data, assemblies, assembly annotations, and pangenome alignments are also available through AnVIL (https://anvilproject.org) and the AWS Open Data Program (https://registry.opendata.aws). Metadata and download URIs available through the HPRC Data Explorer on https://humanpangenome.org/. Assemblies and select annotations are available through a UCSC Genome Browser Hub (https://hgdownload.gi.ucsc.edu/hubs/HPRC/index.html) as well as the Ensembl Genome Browser151 (https://projects.ensembl.org/hprc/). Assembly and epigenomic data is available in the HPRC Epigenome Browser (https://epigenome.humanpangenome.org/).

### Code availability

Centrolign and code used for simulation, benchmarking, guide-tree inference and alignment evaluation are available at https://github.com/jeizenga/centrolign and https://github.com/jeizenga/centromere-scripts. Custom code used for downstream analysis of Centrolign alignments and analyses presented in this study is available at https://github.com/miramastoras/centrolign_analysis and https://github.com/juklucas/hprc2_censat_paper.

Scripts used for pericentromeric structural-variation analysis are available at https://github.com/juklucas/hprc2_censat_paper. Workflows for satellite annotation and HOR-level analyses are available at https://github.com/kmiga/alphaAnnotation (CenSat annotation). Other tooling can be found at https://github.com/fedorrik/stv (HOR

structural-variant annotation), https://github.com/fedorrik/ClusterSourMash (active αSat array clustering), https://github.com/fedorrik/horhap_tool, and https://github.com/fedorrik/annotaligner (Annotaligner). Code for CDR identification is available at https://github.com/jmenendez98/centrodip and https://github.com/jmenendez98/centrodip-reads. The pangenome integration and cenhap-typing pipeline is available at https://github.com/faithokamoto/centromere-haplotype-sampling-pipeline. Code for Graph Reference coordinate assignment and associated variant calling is available at https://github.com/glennhickey/nested-variants.

Software versions, parameters and additional software used in individual analyses are provided in the Methods.

### αSat array selection and quality assessment

HPRC2 assemblies were annotated for centromeric and pericentromeric satellites (CenSat) as well as strand orientation with an automated workflow described elsewhere ^25,62^. Active arrays were then identified using the CenSat BED files. Rows containing “H1L” in the name, indicating active αSat, were selected and annotated intervals separated by ≤ 200 kbp were merged into candidate active arrays. This merging accommodated short interruptions within active arrays caused by features such as transposable elements or αSat sequences not classified as active by the annotation pipeline. Candidate arrays with a span of less than 100 kbp were discarded. We additionally required ≥ 99% of the sequence within each merged interval to be annotated as active αSat or mixedAlpha. To compensate for the large HSat1A arrays found in the active arrays of chromosomes 3 and 4, arrays on these chromosomes were then subjected to another round of merging with a cutoff distance of ≤ 10 Mbp.

Arrays were selected for analysis by requiring that they pass HPRC2 assembly-quality filters^14^ and additional criteria (described below) intended to exclude potentially incomplete arrays. Briefly, candidate arrays were intersected with Flagger HiFi, Flagger ONT, and NucFlag HiFi QC BED files from HPRC2 to identify regions containing potential assembly errors^13,27,63^. Arrays overlapping any region annotated as an “error” were discarded. To exclude potentially incomplete arrays, we applied two additional filters. First, arrays located within 200 kbp of either end of an assembled sequence were discarded because the array could extend beyond the sequence boundary. Second, arrays were discarded when active-array annotations for the same chromosome and haplotype occurred on more than one assembled sequence, indicating that the array may have been fragmented across multiple sequences in the assembly.

### Pericentromeric and centromeric array selection and quality assessment

Pericentromeric arrays were identified from CenSat BED files in a similar manner to alpha satellite arrays. Regions labeled as “ct” (centric transition regions) or “gap” (stretches of N, typically placeholders for rDNA arrays and sites of scaffolding) were removed, after which annotations belonging to the same satellite family and separated by ≤ 100kb were merged by extending each annotation by 100 kbp using bedtools slopv^63^. The resulting merged regions were subsequently extended by 1 Mbp and overlapping regions were merged to generate candidate pericentromeric annotations. When multiple candidate pericentromeric annotations were identified for a given chromosome, they were ranked first by the presence of an active array and then by region size; only the highest-ranking annotation was retained for each haplotype and chromosome.

Annotations on CHM13 were used to define expected large (>500kbp) annotations for all chromosomes. HPRC2 candidate pericentromeric annotations were required to have the same (unordered) within family annotation set as CHM13 with a relaxed cutoff size of 400kbp (or 50kbp for rDNA arrays). Pericentromeric annotations were then required to not be within 200kbp of the end of the sequence with the exception of acrocentric p-arms. Error predictions that fell entirely within rDNA arrays were removed and pericentromeric annotations that had any retained overlapping error predictions were discarded.

### HOR-StV Annotation

HOR structural variants (HOR-StVs) were identified using previously described methods^7^ (https://github.com/fedorrik/stv), extending the approach originally applied to the T2T-CHM13 assembly. Monomer-level αSat annotations generated by the CenSat workflow (https://github.com/kmiga/alphaAnnotation/blob/main/cenSatAnnotation/centromereAnnotation.wdl) were used as input. CenSat assigns each αSat monomer its expected position within the chromosome-specific HOR based on monomer-specific HMM profiles. Using these monomer assignments and their strand orientations, the HOR-StV method partitions αSat arrays into individual HOR repeat units and classifies units according to their arrangement of monomer composition and order. Within each array, the most frequent HOR structure was designated as the canonical HOR, whereas less frequent HOR structures were classified as HOR-StVs. The HOR-StV annotation was performed using the WDL workflow available through the alphaAnnotation repository (https://github.com/kmiga/alphaAnnotation/blob/main/HOR-StV/HOR-StV.wdl).

### Annotation-guided alignment of pericentromeric satellites

To create alignments of pericentromeric regions, we represented each pericentromeric region as an ordered sequence of annotated satellite arrays (called elements below). The arrays were then aligned using a dynamic programming approach, similar to nucleotides in a traditional sequence alignment such as Needleman–Wunsch with affine gap penalties. Quality-controlled pericentromeric assemblies from the HPRC2 were compared against CHM13 v2.0. Chromosomes 1–12, 16–20, and X were analyzed.

Individual satellite arrays (>20kbp) from the CenSat annotation were classified by satellite subtype, including active higher-order repeats, other higher-order repeats, divergent higher-order repeats, human satellites, beta satellite, gamma satellite, monomeric alpha satellite, classical satellites, and centromeric transition sequence (Ct). Sequence similarity between CHM13 and query elements of the same subtype was calculated using Mash (v2.3)^64^ with a k-mer length of 21 and sketch size of 50,000. Each pericentromeric assembly was aligned to the corresponding CHM13 pericentromere using global dynamic programming with affine gap penalties. Match scores were calculated from Mash distance and log difference in length between the two elements. Increases in Mash distance decrease the alignment score, as do increases in size differences between the elements. Forward and reversed alignments were evaluated, and the higher-scoring alignment was retained. Internal inversions were recovered by recursively testing gap-bounded blocks in the reverse orientation. Internal inversions required at least two reverse-oriented element matches. The resulting alignment path, including matched elements, gaps, and element orientations, were represented as PAF files.

An interval was designated a mixing or invasion event when array sequence–active higher-order repeat, other higher-order repeat, or divergent higher-order repeat–was combined with another satellite family, or when a satellite subtype not represented in the flanking array was introduced. Mixing events were kept distinct from insertions and deletions. If both sides contained the same subtype set and their lengths differed by more than 200 kbp, the event was classified as an expansion or contraction according to the direction of the length difference.

Special filtering was applied near interval boundaries. Events composed entirely of Ct sequence were secondary when they occurred at either end of the analyzed region or outside the outermost substantial satellite element, where a substantial satellite was defined as a non-Ct element of at least 200 kbp. These rules prevented incomplete peripheral sequence and variable Ct boundaries from generating structural types.

For the higher-resolution inversion and large-indel summaries, eligible events were required to be strictly larger than 20,000 bp. Inversions were retained when they contained array-associated satellite sequence. Large indels were restricted to events explicitly classified as insertion or deletion; mixing, replacement, expansion, and contraction events were not counted as indels.

Pericentromeric arrays were classified into structural types using haplotype signatures with thresholds of 20, 100, 250, and 500 kbp for event sizes. The greater-than-500-kbp classification was used for the structural configurations found in Supplementary Table 1.

### Cenhap lineages

Initial cenhap trees and assignments were created by Loucks, Langley, Ryabov et al (in preparation). To create serviceable cenhaps^7^ based on the CHM13-mapped high-coverage 1000 Genomes genotypes^65^, we downloaded regions ≥ 1 Mbp contiguous to the CT annotation from chromosome-specific VCFs. Physical phasing information was removed. The genotypes were then phased using akt pedphase^66^. Variants with genotype quality QUAL ≤ 30 for more than 10% of samples or where excess heterozygosity ExcessHet > 30 were filtered. Phased genotypes were imputed using SHAPEIT4.2^67^, with the akt phasing as the scaffolds and the genetic map lifted over from hg38. SNVs with a minor allele count, MAC > 5 were selected for all non-acrocentric autosomes. Then 1000 SNVs directly flanking the active arrays on the P and Q side were used to compute the pairwise Hamming distance. Hierarchical UPGMA clustering based on these pairwise distances yielded the cenhap trees. To capture ‘common’ cenhap lineages represented in the 1000 Genomes dataset while distinguishing rare and/or recombinant lineages the trees were cut to produce 30 cenhaps. Chromosome 12 had 8 cenhaps cut manually to better separate clades.

To determine cenhaps assignments for samples not in the 1000 Genomes Project (CHM13, HG002, HG005, HG06807, and NA21309), and to ensure that cenhap assignments for non-trio samples has phase assignments matching HPRC2 assemblies, we inferred cenhap assignments from trio samples using nearest-neighbor classification. To create callsets with a common set of variants for comparing established 1000 Genomes cenhap labels with haplotypes phased according to the HPRC2 assemblies, we combined Minigraph-Cactus calls (hprc-v2.1-mc-chm13.wave.vcf.gz) from HPRC2^14^ with a phased, CHM13-based, 1000 Genomes variant callset was obtained^68^, (1KGP.CHM13v2.0.whole_genome.recalibrated.snp_indel.pass.phased.native_maps.biallelic.320 2.bcf.gz). For each chromosome, we subset the calls to CHM13 regions surrounding the active arrays (Supplementary Table 2). Variant calls were further subset to biallelic sites represented identically in both data sets. Diploid genotypes were then decomposed into haplotypes and encoded as reference (0), alternate (1), or missing. To allow classification of CHM13, a synthetic CHM13 haplotype with reference calls at all retained sites was added to the query data set. To create a training set, the 1000 Genomes pedigree file^69^ (integrated_call_samples_v3.20250704.ALL.ped) was used to identify and retain children in trio pedigrees. Haplotypes in the training set whose nearest neighbor had a different cenhap label were iteratively removed. We then calculated Hamming distances between training and query haplotypes using variant sites with calls present in both haplotypes. Cenhaps were assigned using the label of the nearest training haplotype (KNN with k = 1). Cenhap counts are shown in **Supplementary Table 3**. Finally, we generated average-linkage UPGMA trees from missingness-aware Hamming distances among MC haplotypes and painted leaves by their inferred cenhap.

### Active ASat array clustering

ClusterSourMash (https://github.com/fedorrik/ClusterSourMash) was used to cluster active αSat arrays. Based on the CenSat annotations, sequences corresponding to active arrays were extracted. Abundance-aware 31-mer sketches were then generated using *sourmash sketch dna*, with all k-mers retained. A pairwise distance matrix was calculated using *sourmash compare* and subsequently clustered using the UPGMA. This analysis was performed across all samples and chromosomes (Supplementary Note 1) and separately on combined datasets comprising chromosome pairs 13 and 21 and chromosomes 14 and 22 (Fig. 1d and Supplementary Note).

### Annotating HORhaps

#### HORHap analysis for interchromosomal analysis and benchmarking Centrolign

The HORhap tool (https://github.com/fedorrik/horhap_tool) was previously used to generate HORhap annotations for one diploid genome^23^ and four diploid genomes from a single pedigree^33^. In this study, we generated HORhap annotations using a dataset representing the broad diversity of human populations.

Each chromosome was analyzed independently, except for chromosomes 13 and 21 and chromosomes 14 and 22, which were analyzed as chromosome pairs. To reduce computational requirements, a representative subset was selected from the complete HPRC dataset. This subset included one or two samples from each cenhap or each major branch of the active αSat-array clustering tree. This procedure reduced the number of samples analyzed per chromosome or chromosome pair from approximately 200–300 to 10–15.

Sequences corresponding to all major HOR structural-variant (HOR-StV) classes, defined as those with frequencies above 1%, were extracted and aligned using MUSCLE v3. When three or more HOR-StV classes were analyzed, sequences belonging to each HOR-StV class were first aligned separately together with one full-length HOR copy. The resulting alignments were then merged using a custom script (merge_alignments.py). Pairwise Hamming distances between aligned sequences were calculated, followed by hierarchical clustering using Ward’s linkage method. At a given tree depth, this procedure partitioned the HOR sequences into *k* clades, referred to as HORhaps. BED files, phylogenetic trees of HORhap consensus sequences, and alignment visualizations were generated for all values of *k* from 2 to 19. Following manual inspection of these outputs, a single optimal value of *k* was selected for downstream analysis, which was chosen to ensure that all major sequence variants were represented while minimizing the mixing of distinct HORhaps.

To extend the annotations to the complete HPRC2 dataset for each chromosome, the alignment corresponding to each HORhap was used to train a profile hidden Markov model (HMM). The resulting set of *k* HMM profiles was then used to annotate all assemblies in the HPRC dataset using script from https://github.com/fedorrik/HumAS-HMMER_for_AnVIL. Each HOR copy was assigned to the HORhap represented by the HMM profile with the highest score, producing a complete HORhap annotation for each assembly.

This analysis was performed for chromosomes 4, 11, 12, X and for the combined chromosome pairs 13/21 and 14/22.

#### HORHaps for cenhap association analysis

For the genome-wide HORhap analysis, each chromosome was analyzed independently. 20 arrays were sampled for HORhap definition. When available, the cenhap tree was used to guide sampling; otherwise, the whole-array Centrolign UPGMA tree was used. Arrays were selected at approximately evenly spaced positions along the ordered leaves of the chromosome-specific tree producing a topology-spaced subset of arrays for each chromosome. HOR copies were then extracted from each selected active array. The three most abundant HOR-StVs which included >2.5% of all extracted HOR copies for the chromosome were retained. HOR copies shorter than 70% or longer than 130% of that StV-specific median were classified as length outliers (likely due to StV misassignment) and excluded from alignment. Retained HOR copies were aligned with HAlign4^70^. Alignment columns occupied in fewer than 1% of sequences were removed. Pairwise sequence dissimilarity was calculated as the proportion of nucleotide differences at aligned positions that were non-gap in both copies. When more than 1,500 HOR copies were available, a subset was used for hierarchical clustering.

HORhaps were evaluated by cutting the Ward dendrogram from k=2 to k=20. For each k evaluated, the mean silhouette coefficient and the relative difference between the Ward merge heights immediately below and above the (k)-cluster cut were calculated. Cluster numbers (k) were ranked using the combined score (s_k+0.15g_k), where (s_k) is the mean silhouette coefficient and (g_k) is the relative Ward merge gap. The highest-ranked partition below the maximum evaluated (k) was selected as the primary resolution. The highest-ranked finer partition was retained as a nested secondary resolution.

Chr4 was additionally rerun for Figure 1 with the following manual changes: additional StVs were aligned and k was set to 12 manually. Briefly, seven HOR-StV classes were initially included. 20 topology-spaced chr4 arrays were selected from the chr4 cenhap tree (assembly-based). In chr4 some HOR-sized units are represented as single oversized StV annotations. In order to annotate their HOR-StV structure and include them in HORhap analysis, all chr4 StV records at least 10 kb long were examined using their underlying monomer annotations. Candidate periods of 4–40 monomers were evaluated by calculating monomer-label identity at each lag. The highest-scoring period was used when its identity was at least 95%. Complete periodic units were extracted as separate rescued HOR copies, while unresolved flanking sequence was excluded. The rescue procedure reconstructed 117 HOR copies from 10 oversized native records. These copies were combined with the 13,022 copies from the seven selected StVs and aligned using HAlign4. One sample (NA18508.2) with a longer branch length in the tree whose HORhaps clustered with the bottom of the tree was removed from the visualization in Figure1 to reduce complexity. The complete (equivalent) set is shown in the genome-wide figure in the supplement (**Supplementary Fig. 16**).

### Chromosome 13/21 recombinant breakpoint analysis

The HG01252.pat.chr21 contig, which carries a chromosome 13-derived centromere, was mapped using minimap2 to a chromosome 21 representative of the most common branch identified by active chromosome 21 array k-mer clustering (HG01192.pat.chr21), and to the nearest chromosome 13 homolog identified by active chromosome 13 array k-mer clustering (HG00344.hap2.chr13). The q-arm was confidently assigned to chromosome 21, whereas the centromere and the centromere-adjacent half of the p-arm were assigned to chromosome 13. The q-arm edge of the centromere lies within a region of high homology between chromosome 13 and chromosome 21, visible as an overlap between the minimap2 tracks (Fig. SN2a).

A 400 kbp window spanning the minimap2 overlap was extracted, and the recombinant sequence within this window was aligned to the corresponding sequences from the chromosome 13 and chromosome 21 references using HAlign4. In the left part of this window, the recombinant showed higher similarity to chromosome 13, whereas in the right part it showed higher similarity to chromosome 21. At the transition between chromosome 13- and chromosome 21-like sequence, a 531-bp interval had no sequence differences among the three sequences and therefore potentially contains the recombination breakpoint. RepeatMasker annotation identified Alu and L1 elements within this 531-bp interval (Fig. SN2b).

### Analysis of genes in CenSat regions

Using the CAT gene annotations from HPRC Release 2, we intersected gene models with pass-QC CenSat arrays (with bedtools) to obtain all gene copies overlapping pericentromeric sequence. Following the procedure in Altemose^7^, we filtered our gene set in the CenSat array boundaries to only include those flanked by peri/centromeric satellite annotations larger than 100 kbp, to maintain consistency in gene sets across samples and focus our analysis on only those genes embedded in the peri/centromere. Genes were retained only if they carried an HGNC-assigned symbol. Placeholder or uncurated names were removed (Ensembl IDs beginning with ENSG, clone/WGS-style accessions such as AC*/AL*/ABBA*, and related placeholders). Small RNA biotypes were excluded; the final callset comprised protein-coding genes, lncRNAs, and named pseudogenes. Per-haplotype gene counts were then tallied and summarized by chromosome.

To relate gene-content differences to local satellite organization, we compared representative haplotype pairs with SVbyEye plots. For each pair, the syntenic CenSat window was extracted from both selected assemblies. CenSat annotations and CAT gene models were projected onto the same coordinates. Contigs were aligned with minimap2 (-x asm5 -c), and alignments were rendered with SVbyEye. Alignments shorter than 10 kbp were discarded; forward and reverse-strand synteny are shown as light and dark ribbons, respectively (**Fig 1b**).

We then quantified long-read RNA-seq expression relative to each individual’s own haplotype assemblies for all the pericentromeric genes using the donor-matched PacBio Kinnex Iso-Seq reads aligned to HPRC Release 2 haplotypes. Per-copy expression obtained from IsoQuant was normalized to TPM (supporting reads / total mapped reads on that haplotype × 10⁶). Copy-level counts were aggregated to haplotype- and gene-level summaries.

### Local identity annotation method

Active arrays were taken from CenSat BED files as entries ending in “H1L”, and adjacent intervals within 200 kbp were merged with bedtools. For each merged array entry, FASTA sequence was extracted with samtools and run through ModDotPlot^24^ (5 kbp window, delta=0). From the resulting all-vs-all identity BED files, a custom Python script (moddotplot_1d.py) averaged each region’s identity across all nearest neighbors (default: two per size). These averaged values were mapped to a color scale set by the 10th percentile of observed values within each merged array and written to BED files with score as the raw identity values^23^.

### Reference-based CDR annotation

Aggregate CDRs were called with centrodip v1.0.2 (https://github.com/jmenendez98/centrodip) on ONT data. For each haplotype-resolved assembly, active αSat arrays were taken from the CenSat annotation by selecting all active_hor records and merging any within 100 kbp of each other (bedtools v2.31.1), and 5mC bedMethyl from modkit pileup (v0.4.2) were restricted to those arrays. Within centrodip, CDR detection was run separately on each chromosome. First, the per-CpG methylation fraction was smoothed by locally weighted linear regression (LOWESS) in a 10 kbp sliding window, weighting each CpG both by its distance from the window center and by its read coverage, so that low-coverage sites contributed less. Local minima in this smoothed profile were identified with scipy.signal.find_peaks, and a minimum was kept only if it fell below the 10th percentile of smoothed methylation and dropped at least 25% of the chromosome’s total smoothed methylation range. Each dip was then extended outward from its minimum until the smoothed signal recovered to within 75% of the background methylation level (the median smoothed methylation outside of candidate dips), and was scored from 0 to 1000 according to how far below background it sat, both on average and at its lowest point, and how many CpGs it spanned. Dips were retained as final CDR calls only if they fell entirely within a single active HOR array, were at least 100 bp long, and scored ≥ 500.

### Read-based CDR annotation

Centromere dip regions (CDRs) were annotated on individual reads using centrodip-reads (https://github.com/jmenendez98/centrodip-reads), which implements a two-state (CDR, non-CDR) hidden Markov model over per-read 5mC signal. Per-read CpG methylation calls were taken from the MM/ML tags of alignment reads to matched centromere assemblies; only primary alignments were retained. Model parameters were estimated in a supervised fashion. Aggregate CDR annotations, generated with centrodip, were intersected with the active-αSat array coordinates from the CenSat annotation to label each methylation observation on a read as CDR or non-CDR. Because 5mC calling accuracy differs between ONT chemistries, two parameter sets were trained independently: one on R9 data (n=157 samples) and one on R10 data (n=71 samples). Training used one centromere per chromosome, selected for having the most clearly resolved dip in the aggregate 5mC signal (Supplementary Fig. 19). Using these trained parameters, CDR state paths were inferred by Viterbi decoding for all reads overlapping active-αSat at every centromere. CDR intervals separated by ≤ 750 bp were merged, and merged intervals < 2.5 kbp were discarded, yielding the final read-based CDR annotations.

### Read-to-read centromere concordance estimate of read-based CDR annotations

Read-to-read centromere concordance was calculated using a pooled-Jaccard index on the read-based CDR annotations. Across each active-αSat HOR array, reads and their CDR annotations were binarized in 1 kbp windows: a read covers every window it spans and marks every window its CDR patches overlap. Only informative reads and windows were scored. Informative reads had at least one patch and were called across ≥ 95% of the read. Informative patches had a minimum coverage of 5. Within each scored window we counted the read pairs marking it jointly (intersection) and the read pairs in which at least one read marks it (union), then summed both counts across all windows in the array. We report read-to-read centromere concordance as the ratio of these pooled sums, one value per centromere.

### Weighted sub-CDR size sum estimation

Within each active-αSat we calculated the fraction of read-based CDR support along every base. This fraction is the base’s weight: a base called by all of its reads adds a full base pair, a base called by half of them adds half. This calculation was performed and summed across all bases called as non-background. To measure background, each centromere’s own CDR call rate was measured outside of its aggregate CDR calls (padded by 50 kbp). Every base was tested against that rate with a binomial test and counted only if it passed at a 5% false discovery rate, contributing its support minus the background rate. Bases with coverage below 5 were skipped. We verified the background by shuffling each read’s CDR patches to random positions within that same read, where the true answer is zero, and it returned 0.0 kbp at the median centromere and 0.4 kbp at the 90th percentile.

### The Centrolign algorithm

#### Querying exact matches

Sequences as large as active arrays are too large to use optimal, exact alignment algorithms in a feasible amount of time. The standard solution to this problem is to begin by partially specifying the alignment using only exact matches as “anchors”. These exact matches can be identified using data structures that index matches in the sequences. There exist such data structures that remain efficient even for large input sequences.

Most existing match indexes are designed for strings, not sequence graphs. There are indexes for certain acyclic sequence graphs, but they are unsuitable for long tandem repeats since their size can scale exponentially with the longest repeated sequence^71^. In addition, to determine the uniqueness of a match, Centrolign requires the index to support count queries, which return the number of times a match occurs in each of the two graphs. For these reasons, we implemented a hybrid index that combines multiple existing indexing techniques.

The core of Centrolign’s match index is a generalized suffix tree (GST)^72^ over all of the input sequences contained in the MSA subproblem. The GST is implemented implicitly as an enhanced suffix array (ESA)^73^ in which the suffix array (SA) is computed with the SA-IS algorithm^74^ and the longest common prefix (LCP) array is constructed with Kasai et al.’s algorithm^75^.

On its own, this GST is not well-suited to performing count queries for the sequence graph. If a match is present in multiple input sequences, this GST represents it duplicatively, which will inflate the count relative to the graph. Accordingly, we apply another data structure to deduplicate these occurrences. In the ESA, a match corresponds to a certain interval over the SA. Alongside the SA, we prepare another array that contains the graph’s node IDs corresponding to the beginning of each suffix in the SA. To compute the count of a match in the graph (as opposed to in the GST), it is sufficient to compute the number of unique node IDs in the interval of this array that corresponds to that match’s SA interval. This operation can be performed by Hui’s ^76^ color set size (CSS) data structure, with a small modification. We create two of these data structures, each counting only occurrences of the match in one of the two graphs being aligned. This lets us query a match’s count within each graph separately.

Following Bzikadze and Pevzner^42^, we query minimal (*n*, *m*)-rare matches using the GST-CSS index. These matches satisfy two conditions:

1. (*n*, *m*)-rarity: The match has a count of *n* in the first graph and *m* in the second.
2. Minimality: If the match were shortened at all, the count would increase in at least one of the graphs.

Our algorithm to compute minimal (*n*, *m*)-rare matches corrects some technical inaccuracies in Bzikadze and Pevzner’s algorithm and generalizes it to the sequence graph setting. Given the interval corresponding to a match *M* in the GST, we can compute the counts *n* and *m* on the two input graphs using the CSS index. In the **Supplementary Note 2 Section 1**, we establish simple necessary and sufficient conditions for a match to be minimal and (*n*, *m*)-rare, which can be checked using the GST and CSS. This leads to a straightforward algorithm that scans the edges of the GST and selects those that meet these conditions. Each iteration performs a constant number of basic suffix tree operations and CSS queries, each of which takes *O*(1) time. This yields a total run time of *O*(*N*), where *N* is the sum of the lengths of the sequences. By default, Centrolign also restricts to matches for which both *n* · *m* is at most 3,000 to limit downstream computation time. Pseudocode is provided in **Supplementary Note 2 Section 1**. **Supplementary Note 2, Figure 1** shows a worked example.

#### Identifying a chain of exact match anchors

After querying matches, Centrolign identifies the highest-scoring co-linear chain of matches to use to anchor the alignment. Each anchor consists of two walks (one through the MSA graph of each intermediate subproblem) with matching sequences. To be co-linear, the beginning of each anchor must be reachable from the end of the previous one in both graphs.

#### Uniqueness anchor scoring

The anchoring stage is the step of the algorithm in which Centrolign prioritizes uniqueness in the alignment. This is accomplished through a specialized weight function that down-weights matches by the geometric mean of their count on the two graphs. In particular, for an (*n*, *m*)-rare match of length *l* can be given a weight *w* as

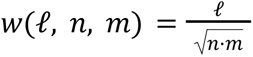

As currently stated, this function fails to reflect some statistical properties of minimal matches. To identify a parsimonious alignment, it makes sense to assign greater weight to matches of greater length. However, due to the minimality criterion, greater length also suggests lower confidence. This is clearer when stated in the contrapositive: the match cannot achieve (*n*, *m*) -rarity with any length smaller than *l*. For example, if a minimal unique (i.e. (1, 1)-rare) match has a length of 1,000, then both of the length 999 subsequences it contains are non-unique. For such long matches to be non-unique suggests a high level of repetitiveness. In contrast, a minimal unique match of length 10 suggests a much lower level of repetitiveness because it would be less surprising that the length 9 sequences it contains were non-unique in the array by chance. To capture the ambivalent value of length for parsimony and confidence, we introduce an additional term to the weight function that provides decreasing weight for longer matches:

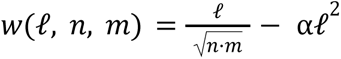

where α is an additional scoring parameter.

#### Chaining with sparse dynamic programming

Centrolign computes the optimal chain of anchors with sparse dynamic programming algorithms. These algorithms are based on previously described sequence-to-graph sparse chaining algorithms^77,78^. At a high level, these previous methods generalize sequence-to-sequence chaining algorithms by applying them to a path cover: a set of paths in which every node of the graph is included in at least one path. Each of these paths is essentially a sequence, and accordingly, the sequence-to-sequence sparse dynamic programming algorithms can be applied to them. To generalize to graphs, the dynamic programming for all of the path cover’s paths is interleaved with each other according to a topological order of the graph. In addition, the dynamic programming communicates between paths at particular points that are strategically chosen to ensure that all chains that cross between paths are considered.

Centrolign generalizes these techniques for graph-to-graph alignment using a path cover for both graphs. We interleave a dynamic programming problem for all pairs of paths (one path from each graph). There are two such chaining algorithms implemented in Centrolign. The first one is based on an algorithm by Mäkinen, et al. (2019) and has no penalty for insertions and deletions. The second is based on an algorithm by Chanda and Jain (2023) that implements affine gap penalties. The affine gap algorithm is considerably slower than the free gap algorithm. For *N* matches and path covers of size *k* and *k*, the free gap algorithm takes time *O*(*k*_1_ *k*_2_ *N* log *N*) and the affine gap algorithm takes *O*(*k*_1_ *k*_2_ *N* log^2^ *N*) Detailed analysis and pseudocode is presented in **Supplementary Note 2 Section 2**. The supplementary information also includes a technique to speed up the affine gap algorithm (as well as Chandra and Jain’s (2023) algorithm) by a factor of *O*(log log *N*), which is not implemented in Centrolign but may be of theoretical interest.

#### Parameterizing affine gap penalties

In general, we prefer the affine gap chaining algorithm, since it encourages parsimonious alignments around large insertions and deletions, which are common in active arrays. However, parameterizing the gap penalties is challenging. Different levels of sequence divergence in the alignment lead to different levels of uniqueness for the matches, in turn leading to different average weights for anchors. If the gap penalties are of a much greater scale than the anchor weights, the chain can be trapped within a single high-scoring diagonal. If they are of a much lesser scale, they do not produce parsimonious chains. To address this challenge, before executing the affine gap algorithm, Centrolign first calibrates the gap penalties to the scale of the anchor weights using the faster free gap algorithm. The scale is estimated as the total weight of the optimal free-gap chain normalized by its length (including the space between anchors). The gap penalties are then scaled proportionately to this value.

In reality, Centrolign uses a piecewise affine gap penalty rather than a simple affine gap penalty. This is accomplished through the typical technique (due to Gotoh^79^) of adding additional dynamic programming structures for each set of penalties. By default, Centrolign uses three sets of penalties with successively larger gap open penalties and smaller gap extend penalties. This helps improve parsimony when there are insertions and deletions of widely varying sizes.

#### Practical limitations and heuristics

In practice, it is possible to query many more matches than can feasibly be used in the chaining algorithms. Centrolign uses multiple strategies to limit this computational burden. First, there is a hard maximum on the number of matches that it will consider, which is set to 1.25 million by default. Next, because more distantly related sequences have fewer alignable regions, this limit is reduced proportionately to the estimated scale of the anchor scores (which was computed when calibrating the gap penalties). Centrolign prioritizes the anchors for use in the chaining algorithm based on the weight function used to score them. Finally, after computing the optimal anchor chain, Centrolign performs a “fill-in” anchoring problem between each pair of adjacent anchors in the chain. In these problems, all of the matches that fall strictly between the two anchors are included. In practice, this greatly reduces the number of matches considered. The algorithmic implementation of the fill-in subproblems is identical to the full anchoring algorithm.

#### Identifying confidently alignable regions

Because active arrays can have rapid sequence evolution, it is common that some portions of the array have nearly completely turned over. In these regions, finding a meaningful base-level alignment is not possible. Accordingly, the alignment algorithm needs to be able to differentiate between alignable and unalignable regions. Centrolign identifies unalignable sequences using the information contained in the optimal anchor chain. When the sequence turns over, it is accompanied by a loss of the uniqueness signal in the anchors, since the only remaining matches are non-specific ones scattered throughout the array. Accordingly, the unalignable regions are often identifiable as extended regions of the anchor chain with a low score. We formalize the algorithmic problem of identifying these regions as one of optimally partitioning the anchor chain into alignable and unalignable intervals according to an objective function and certain constraints. Detailed analysis and pseudocode for this algorithm is presented in **Supplementary Note 2 Section 3**.

The constraints used in the partitioning algorithm include a minimum on the score density of an alignable region: its total anchor weight divided by its length. This minimum threshold is sensitive to the overall scale of the weight function used to score the anchors, which could make it challenging to parameterize. We address this challenge by calibrating the threshold to the scale of the weight function before executing the MSA. At startup, Centrolign computes the optimal chain for each input sequence against itself. The average score density of these self-self chains is used to quantify the overall scale of the weight function on these sequences, and the partitioning algorithm’s threshold is scaled proportionately to this value.

#### Stitching anchors into a base-level alignment

After computing an optimal chain and identifying the alignable regions within it, Centrolign uses alignment algorithms to fill in the gaps between anchors in the alignment, forming a complete base-level alignment. Multiple alignment algorithms are used, depending on the size of the graphs that are being aligned. If the graphs are small enough, we use base-level POPOA^43^. If both graphs are larger than a threshold, we instead use a generalization of the graph wavefront alignment algorithm^80^ that performs graph-graph alignment instead of sequence-graph alignment. We refer to the algorithm as graph-graph wavefront alignment (GGWFA). Since active arrays are enriched for large insertions and deletions, we have also implemented a specialized variant of GGWFA for these cases. It performs GGWFA inward from both ends of the alignment until it can infer a single, long insertion or deletion to join the two sides together. This simplification greatly reduces the computation required to align across these mutations. In all alignment algorithms, we use a piecewise-affine gap penalty with three pieces, similar to those used in the chaining algorithm. Algorithmic details and pseudocode for the GGWFA algorithms are presented in **Supplementary Note 2 Section 4**.

### Applying Centrolign to the HPRC release 2 assemblies

#### Calculating Centrolign pairwise match distance and generating input neighbor-joining trees

To perform an MSA, Centrolign requires as input a guide tree representing the genetic distance between the input assemblies. We took the αSat arrays for each chromosome passing our QC filters, and performed pairwise alignments of all combinations of assemblies with default parameters. We parsed the resulting CIGAR string with a custom Python script (https://github.com/jeizenga/centromere-scripts/blob/main/data_exploration/infer_tree.py) and used the formula 1.0 - ((2.0 * matches) / (ref_len + query_len)) to calculate a distance metric for all pairs of assemblies. It was essential to include insertion and deletion bases in this metric (rather than just using mismatches / mismatches+matches) because the majority of centromere arrays are completely diverged from each other, and have very little alignable bases, and yet we still need a genetic distance measure for them to place them in the tree. We refer to this distance metric in the following sections as the “pairwise match distance”. We next implemented the neighbor-joining algorithm ^44^ in the same Python script to infer a tree from the pairwise match distances.

#### Constructing MSAs for all chromosomes

Because no alignment can be created in the MSA between completely diverged clades, we decided to split the guide trees into 2-3 subgroups per chromosome before passing to Centrolign, in order to save on runtime. Subgroup divisions were created manually by inspecting the tree heatmaps for each chromosome and optimizing tree divisions that kept alignable clades together. The number of samples in each MSA subgroup run and the runtime can be found at Supplementary Table 7.

We found that, while the default Centrolign parameters worked well for most chromosomes, they led to poor consistency between direct and induced pairwise alignments for chromosome Y. This resulted from deceptively high-scoring alignments of rapidly evolving, highly homogenized repeats, which were not accurately identified as unalignable. To remedy this situation, we developed a separate set of scoring parameters for chromosome Y that increased uniqueness penalties on the corresponding matches.

### Benchmarking Centrolign

#### Simulating centromere sequences with sim_centromere

We developed a C++ tool for simulating centromere sequences with a known truth alignment, sim_centromere. This script was designed only to recapitulate certain qualitative features that we observed in real centromere comparisons. In particular, it is not intended to be a mechanistically realistic model of centromere sequence evolution, although some features are motivated by hypotheses of centromere evolution.

To generate a simulated sequence, sim_centromere starts with an annotated active HOR array and repeatedly modifies it over some number of iterations. To generate a pair of sequences with a truth alignment, this process is executed twice from the same initial sequence. Before each execution, each base of the initial sequence is assigned a unique identifier, and during the execution, every new base that is created by the modifications is considered to arise from one of the existing bases (for example, by copying it in a duplication) and therefore inherits its identifier. In a given alignment of the two resulting sequences, a pair of aligned bases is considered correct if both bases have the same identifier, indicating that they originate from the same base in the initial sequence. Moreover, this simulation process makes it possible to compute the maximum number of correctly aligned bases achievable in a colinear alignment. To do so, we treat each identifier as a distinct character and construct a string of identifiers for each sequence. The maximum can then be computed as the longest common subsequence of the two identifier strings. The maximum number of correct bases provides the denominator that is necessary to compute precision and recall for an alignment.

Small extensions of this model also allow sim_centromere to simulate collections of sequences for MSAs. In this mode, sim_centromere takes a tree as input in Newick format. The initial sequence is then treated as the root of the tree, and each branch indicates a new sequence that is simulated from its parent sequence in the manner described above. In the end, the sequences corresponding to the leaf nodes are emitted. For MSAs, it is no longer possible to compute the optimal number of correctly aligned bases, since the multiple longest common subsequence problem is NP-Hard. However, the optimal pairwise alignment provides an upper bound on the number of correctly aligned bases in the pairwise alignment induced by the optimal multiple sequence alignment, which we use to compute approximate precision and recall.

In the sim_centromere model, each iteration of modifying a sequence consists of five phases, corresponding to different types of modification: substitutions, base indels, monomer indels, light-tailed HOR indels, and heavy-tailed HOR indels. In all cases, the modifications are applied independently at each position in the sequence, and each event is chosen to be an insertion or deletion with equal probability. Base insertions add random sequence, but monomer and HOR indels both duplicate the adjacent sequence in tandem.

For monomer and HOR indels, integer numbers of their respective unit are added and removed, and (when possible) the two breakpoints are selected at equivalent positions in the two monomers or HORs to maintain their register. To facilitate doing so, before starting the simulation, sim_centromere aligns every HumAS-HMMER-annotated monomer in the array to the consensus alpha satellite unit. This alignment is used to assign a canonical position within the consensus monomer to every base in the input sequence. During modifications, these canonical positions are inherited in a similar manner to the unique identifiers. The breakpoints of monomer and HOR indels match the canonical positions as closely as possible to ensure that the monomer register is maintained. For HOR indels, the HOR register is also maintained by matching the monomer class within the HOR, which is provided by the input annotations.

All indel types except for heavy-tailed HOR indels use a geometric distribution (formulated with strictly positive support) to model the indels’ lengths. The integer length generated by this distribution is interpreted as a number of bases, monomers, or HORs, depending on the indel type. The heavy-tailed HOR indels use a discrete Pareto distribution^81^. The default parameters are described in Supplementary Table 4. They were tuned by visually comparing the dotplots of the resulting sequences to the dotplots of pairs of real sequences and adjusting parameters to recapitulate qualitative features.

#### MSA simulations

Because sim_centromere does not currently model structurally variant HORs, we selected only chromosomes from CHM13 whose HOR arrays have a single dominant HORs^82^ (chromosomes 1, 5, 8, 9, 13, 18, and 19 were removed). We obtained the CHM13 CenSat annotations from https://s3-us-west-2.amazonaws.com/human-pangenomics/T2T/CHM13/assemblies/annotation/chm13v2.0_censat_v2.1.bed and used bedtools to extract the coordinates of the active array, and subset the CHM13 fasta to these coordinates.

For 30 repetitions for each chromosome, we used msprime^83^ to simulate a tree from CHM13 with 8 sequences and an expected 200 generations to the root of the tree. We then used sim_centromere to simulate FASTA sequences according to the tree. We ran Centrolign with the input tree and sequences to produce an MSA, adding parameter -A to additionally produce pairwise CIGAR alignments induced from the MSA. Using a custom script (https://github.com/jeizenga/centrolign/blob/main/src/scripts/compare_truth_aln.cpp), we compared “truth” CIGAR string alignments produced by sim_centromere with the CIGAR string alignments induced from the simulated MSA, calculating completeness (Recall) as # aln_mismatches / # truth matches, and accuracy (Precision) as aln_matches / (aln_matches + aln_mismatches).

#### Comparison with other pairwise tandem repeat aligners

Using the same regions from CHM13 described in “MSA simulations”, we ran 60 repetitions using sim_centromere to simulate 6 sequences with 25, 50, 100, 150, 200, and 300 generations of evolution. For all 15 possible pairwise combinations, we ran Centrolign in direct alignment mode, producing a CIGAR string. We ran UniAligner v0.1^42^ and Rama v1.2.0^84^ on the same sequences using both program’s default parameters. As in “MSA simulations”, we used (compare_truth_aln.cpp) to calculate precision and recall metrics from comparing against the “truth” CIGAR string produced by sim_centromere.

#### Tree-building evaluation

We tested whether the Centrolign MSA can recapitulate the structure of its input guide tree. We took the MSAs produced on the simulated sequences (see MSA simulations), and for each we calculated pairwise match distances using the formula 1.0 - ((2.0 * matches) / (ref_len + query_len)) on pairwise CIGAR strings induced from the MSA. Using these distances we inferred a neighbor-joining tree with the custom Python script https://github.com/jeizenga/centromere-scripts/blob/main/data_exploration/infer_tree.py. We compared each tree inferred from the MSA to the “true” simulated tree using the bipartition concept of the Robinson-Foulds distance, implemented in the script https://github.com/jeizenga/centrolign/blob/main/src/scripts/tree_compare.cpp. This script directly compares the bipartitions induced by the simulated tree and the inferred tree while also recording the tree height of the edge corresponding to each bipartition.

#### Induced and direct pairwise consistency

To compare the consistency between Centrolign’s direct pairwise alignment mode and the MSA mode, we ran Centrolign in direct pairwise mode on the same simulated sequences from “MSA simulations.” We took the same custom script, (compare_truth_aln.cpp), used to calculate Precision and Recall against the “true” simulated alignment, and ran it on the direct pairwise alignments to compare against the induced ones. We found nearly identical results, indicating high concordance between the induced and direct alignments on simulated data. We also applied the Centrolign pairwise match distance formula to the induced alignments from all chromosomes on real data to assess robustness to artifacts based on progressive alignment.

#### Centrolign SV size comparison with StV annotations

To establish expected size ranges for HOR structural variants (StVs) in each chromosome, we aggregated per-assembly HOR-StV annotation BED files generated by the CenSat annotation pipeline. For each chromosome and HOR repeat name, we calculated the observed size range and relative frequency across all HPRC2 samples. We kept only HOR StVs at greater than 5% frequency per chromosome in the HPRC2 dataset, and plotted their size ranges alongside the size distribution of Centrolign SVs.

#### Aligning HORhaps with annotaligner

We generated HORhap annotations for chromosomes 11, 12, and X of all HPRC2 contigs passing QC (See above section “Annotating HORhaps”). To get HORhap annotation-based SV calls, we used a custom implementation of the Needleman–Wunsch global alignment algorithm (https://github.com/fedorrik/annotaligner). The alignment “alphabet” consisted of strings encoding the HOR name, its StV structure, and HORhap class ID (e.g. S2C9H1L.1-7::C3). This algorithm was applied to pairs of samples with Centrolign pairwise match distance less than 0.2. From the resulting alignments, indels were collected into a set of BEDPE files for comparison with Centrolign SV.

#### Comparison of Centrolign SV and annotaligner SV concordance

To compare annotaligner-derived SV calls to Centrolign’s SV calls, we developed a window-based approach allowing for variation in exact indel placement within a reasonable distance of sequence, as well as split SV calls. First, we created windows around every Centrolign SV that were approximately 2× the observed StV size for that chromosome (2500 bp for chromosome 11, 3000 bp for chromosome 12, and 4500 bp for chromosome X). Within each expanded window, we find the contiguous subset of annotaligner SVs that is closest to the size of the Centrolign SV (to account for split SV calls). If the contiguous subset is within 50% of the size of the Centrolign SV, we label both the Centrolign SVs and annotaligner SVs contained in the subset as “matched”. We repeat the same procedure for the reverse (checking all annotaligner SVs for matches in the Centrolign SVs). We report the fraction of Centrolign SVs covered or matching annotaligner SVs, and vice versa, as well as the harmonic mean of the two metrics. This comparison approach is implemented in the custom Python script (https://github.com/miramastoras/centrolign_analysis/blob/main/scripts/sv_comparev2.py).

### Visualizing Centrolign

#### Graph visualization in Bandage

To aid in visualization, chromosome-level Centrolign GFA files were combined and simplified to reduce the number of segments (nodes) and links (edges) present in the graph. When multiple GFA files were available for a chromosome (due to separate runs), they were merged into a single graph. Dummy start and end segments were added to represent the termini of all paths. The graphs were then simplified iteratively by collapsing bubbles with all alleles ≤50 bp and unbranched chains. Segment lengths, links, and path traversals were updated after each iteration. The resulting filtered GFAs represented simplified segment lengths with LN tags and the node sequence replaced with a placeholder (*).

We further simplified the graph by subsampling haplotypes to aid in visualization of diverse cenhaps. Paths in each chromosome graph were grouped by their cenhap label and up to 20 paths were selected per cenhap (random selection with a fixed seed). A subset GFA was then constructed from the selected paths. Only segments and links traversed by at least one of the selected paths were retained.

The resulting graphs were visualized with BandageNG^46^ using a linear layout. Colors were assigned using the color of the cenhap which had the most paths traversing the segment. Segments with zero assigned support or an exact tie between CenHap labels were colored gray.

### Characterizing alpha satellite variation from induced pairwise alignments

#### Deriving SNV, short indel, and SV mutation rates from induced pairwise CIGAR strings

In order to derive SNV, short indel, and SV mutation rates for each sample pair, we wrote a custom Python script (https://github.com/miramastoras/centrolign_analysis/blob/main/scripts/per_cigar_mutation_rate.py) that parses each input alignment CIGAR string to derive this information. We compute the per-sample pair SV (defined as an indel of size ≥ 50 bp) rate as the # of SVs / the average length of the two arrays. We only include SVs in this calculation that are either flanked by aligned sequence, or those that are flanked by an adjacent insertion or deletion that is > 10% different in size. We implemented this filter to prevent mistaking unaligned sequence (represented in the CIGAR string as adjacent insertions and deletions) as structural variation, despite the fact that the alignment does not directly attest to a variant. Adjacent insertions and deletions that are > 10% different in length imply at least one structural event, if not multiple, so including them leads to a conservative estimate of mutation rate. For short indels (≤ 49 bp), we also use this filtering strategy to exclude counting any unaligned sequence, and define the short indel mutation rate as # short indels / # aligned bases. For SNVs, we define the mutation rate as # SNVs / # of aligned bases. We also divided these rates by the patristic distance between each sample in the Centrolign guide trees (Supplementary Table 9). We derived the patristic distances using a custom script, https://github.com/jeizenga/centrolign/blob/main/src/scripts/tree_pair_dist.cpp, which calculates the distance between any two nodes in the tree as the sum of the branch lengths on the path between them.

To study how these per-sample pair mutation rates vary within chromosomes and among related clades, we implemented a Python script that groups samples into clades based on an input phylogenetic tree and Centrolign pairwise match distances (https://github.com/miramastoras/centrolign_analysis/blob/main/scripts/identify_low_divergence_clades.py). The script finds clade groupings, requiring that 95% of samples in a clade have at least a distance of 0.8. Where inferred cenhap labels were available, we used those instead of the clade labels. For all analysis using pairwise CIGAR strings, unless otherwise specified, we utilized induced pairwise alignments induced from the MSA, rather than direct pairwise alignments, because they are evolution-informed.

#### Comparison of mutation rates in the active array, flanking regions, and genome wide

We derived mutation rate estimates in the regions flanking the active arrays from the HPRC2 v2.0 minigraph-cactus (MC) CHM13 graph. To obtain variant calls against CHM13, we subset the MC VCF (<u>hprc-v2.0-mc-chm13.wave.vcf.gz</u>) to 50 kbp up and downstream of the CHM13 active arrays using bcftools view with flags -r -S and --force-samples ^85^. We also created BED files using bedtools flank -b 50000 to represent these flanking regions on CHM13. To obtain the regions in each assembly aligned to CHM13 in the graph, we converted the HPRC graph in HAL format to MAF format with hal2maf and flags --refGenome CHM13 --noAncestors. We also used flag --refTargets to subset the resulting MAF file to the 50 kbp flanks. We implemented a custom Python script, https://github.com/miramastoras/centrolign_analysis/blob/main/scripts/maf_to_bed_per_sample.py, that parses the MAF file and produces a BED file for each sample containing the aligned bases to CHM13 (Supplementary Figure 35). In a custom IPython notebook, (https://github.com/miramastoras/centrolign_analysis/blob/main/analysis_notes/release2_QC_v2 /notebooks/mutation_comparison_to_flanks.ipynb), we parsed the subsetted MC VCF, calculating pairwise short indel and SNV mutation rates for all combinations of samples. Rates were calculated separately for the p-arm and q-arm flanks, and separately for SNVs and short indels (< 50bp) as (# of differing alleles / # of shared aligned bases to CHM13). We excluded any alleles where one or both samples had a missing genotype. We removed the rate for any sample pairs where the union of their shared aligned bases to CHM13 was less than 50% of the total CHM13 bases for that side of the flank (50 kbp). For the acrocentric chromosomes (chromosomes 13, 14, 15, 21, and 22) we removed the p-arm rate values. For each sample pair we took the average of the p- and q-arm rates after applying these filtering approaches. To obtain the per-sample pair mutation rates in the active arrays, we parsed the Centrolign CIGAR strings induced from the MSA as described in the section above.

To calculate the average number of SNVs, short indels, and SVs genome-wide, we took the HPRC2 v2.0 minigraph-cactus VCF and first ran bcftools norm -m -any to split multi-allelic variants. We next ran bcftools view with flag --regions-file CHM13_notinalldifficultregions.bed in order to subset the variant calls to confident regions in the MC VCF. This BED file was obtained from the GIAB FTP site, and excludes CHM13 segmental duplications, low mappability regions, high/low GC regions, tandem repeats, and difficult XY regions, representing 74% of CHM13. We then used a custom Bash script (https://github.com/miramastoras/centrolign_analysis/blob/main/analysis_notes/release2_QC_v2/mutation_flank_comparison.md) to count the number of variants of each class in the resulting VCF.

#### Visualizing mutation rate as a function of CDR distance and local identity

To visualize the differences in SNV rate as a function of distance to the CDR, we created a BED file containing non-overlapping 10 kbp windows across the active satellite array for every sample. We then took our pairwise SNV calls in BED format for sample pairs with pairwise match distance < 0.2, and used bedtools coverage to calculate the number of SNVs overlapping each 10 kbp window for each sample in the pair. We then used bedtools closest to get the distance from that sample’s centrodip CDR BED intervals for every window. If a window overlaps the CDR at all, it is assigned a distance of 0 by bedtools. We implemented a custom Python script, get_aligned_bases_bed.py, that parses the original CIGAR alignment for each sample pair to produce the number of aligned bases in each window. This allowed us to calculate the SNV rate (# of SNVs / aligned base) in each window. To generate the main panel plots, we binned windows by +/- 1.5 Mbp away from the CDR, and computed a chromosome-weighted average for each bin, requiring that chromosomes have at least 10 windows in a bin to be considered in the average. We repeated this same procedure for short indels and SVs, using window sizes of 10 kbp and 100 kbp respectively. For the SVs, the denominator for the rate calculation was window size, rather than aligned bases.

#### Permutation tests for mutation rate as a function of CDR distance and local identity

Upon observing a subtle increase in mutation rate surrounding the CDR, we developed a per-chromosome permutation test to check whether mutation rates inside the CDR were statistically different than expected by chance in the rest of the array. The test statistic for this test was (mean rate of variation in CDR − mean rate of variation outside CDR). For SNVs and short indels (< 50 bp), the rate was calculated as (# of SNVs / # of aligned bases). For SVs, the rate was calculated as (# SVs / window size). To approximate the null distribution, we performed N = 10,000 permutation replicates per chromosome. Within each replicate, for both samples in every sample pair, we randomly placed intervals the same size as the real CDR across the array and re-computed the test statistic. While permuting, we did not allow overlap between sub-CDRs if multiple sub-CDRs existed for the sample for each sample.

Only haplotype pairs with a Centrolign pairwise match distance < 0.2 were included in the permutation analysis. Pairs where either haplotype lacked CDR annotations or where CDRs spanned more than 20% of the total αSat array size were excluded. All samples but 4 from chromosome Y failed to pass this filter, so chromosome Y was excluded from analysis. An empirical two-tailed p-value was computed as the proportion of null replicates with an absolute test statistic greater than or equal to the observed absolute test statistic. If no null replicates exceeded the observed statistic, the p-value was set to 1/N. We used an α value of 0.01 for the test. We repeated the same permutation experiment using the top 10% of local identity windows per each sample instead of the CDR intervals.

Because of the observation that CDR placement tends to associate with the most homogenized parts of the array, we sought to untangle the contributions of these two sequence contexts. We repeated similar permutation tests. The test statistic was the same as in the previous tests, but the focal intervals were computed by subtracting the CDR intervals from the top 10% local identity windows, and vice versa. The comparison regions consisted of the intervals of the active array that did not fall in either the CDR or the top 10% local identity windows. In each permutation replicate, (N=10,000), for every sample in every sample pair, we randomized the position of the two sets of intervals, and then recomputed the interval arithmetic (i.e. the subtracted intervals and the complement). The test statistic from resulting intervals were used to approximate the null distribution.We computed the p-values and as described above, with the same α value for the test.

### Ti/Tv analysis

#### Region definition and sequence statistics

Analyses were performed using CHM13 v2.0 coordinates on chromosomes 1–22 and X. CenSat intervals at least 20 kb long were extracted for monomeric α-satellite, divergent higher-order repeat (dHOR) α-satellite, and active α-satellite arrays. Non-active HOR annotations were not included as Centrolign variant calls (alignments) were not produced for these regions and Minigraph-Cactus did not align them consistently.

In order to define background regions, genomic intervals were randomly sampled from CHM13 outside CenSat and segmental-duplication annotations. CenSat and flattened segmental-duplication annotations were combined, sorted, and merged using Bedtools. Bedtools complement was used to define regions outside of the merged CenSat/SegDup annotations. Candidate lengths were sampled uniformly from 100–200 kb, and bedtools shuffle was used to place 1,250 non-overlapping candidates genome-wide with a random seed. Candidate regions were evaluated using the minigraph-cactus VCF and retaining biallelic SNVs. Regions were required to have at least 95% of SNV sites with genotype calls in at least 90% of the HPRC samples. 1000 regions were retained.

In order to calculate sequence statistics for selected regions, CHM13 sequence was extracted for each region. GC content was calculated as the percentage of A, C, G, and T bases that were G or C. CpG density was calculated as the number of CG dinucleotides divided by region length. Overlapping occurrences of each of the 64 trinucleotides were counted. Trinucleotide counts were divided by region length to obtain frequencies per base in each region.

#### Ti/Tv variant calling with Minigraph-Cactus and Centrolign

Minigraph-cactus variants were obtained from the CHM13-based HPRC2 VCF for background, monomeric α-satellite, and dHOR α-satellite regions. Variants were filtered to biallelic SNVs using bcftools (bcftools view --min-alleles 2 --max-alleles 2 --types snps). An allele matrix was constructed with variant sites as rows and haplotypes as columns. CHM13 was included as an additional reference haplotype. For each unordered haplotype pair, only sites with an allele call for both haplotypes were compared. A↔G and C↔T differences were classified as transitions; all other single-nucleotide differences were classified as transversions. Ti/Tv was calculated as the number of transitions divided by the number of transversions. In order to reduce artifacts from sampling at low numbers, comparisons were retained when they contained at least 50 pairwise SNV differences. Active-array Ti/Tv values were obtained from Centrolign for haplotype pairs with a pairwise alignment distance <0.2. Values were calculated from SNVs further than 10-bp from an InDel and below the 95th-percentile for that array comparison to remove potential false positives from misalignment artifacts. Comparisons were required to contain at least 50 SNVs.

Minigraph-cactus observations for background, monomeric α-satellite, and dHOR α-satellite regions were combined with Centrolign observations for active α-satellite arrays. To limit plotting density, no more than 5,000 pair-region observations were selected per chromosome and sequence class. Violin plots were generated both genome-wide and separately by chromosome.

#### Trinucleotide correction estimate for Ti/Tv

Genome-wide active-array Centrolign Ti/Tv was corrected using the minigraph-cactus background mutation spectrum. Biallelic SNVs in the 1,000 background regions and pass-filter Centrolign SNVs from the filtered active-array comparisons were assigned to 96 substitution channels. Variants and their sequence contexts were reverse-complemented so that all substitutions were represented as C- or T-centered events. Trinucleotide composition was calculated by pooling trinucleotide counts across all bases in the background and active-array region sets and combining reverse-complementary contexts. For each substitution channel, the background SNV count was divided by the corresponding background-context frequency to estimate a composition-normalized mutation rate. This rate was then weighted by the frequency of the same context in active α-satellite arrays to calculate the Ti/Tv expected from the background mutation spectrum at active-array sequence composition.

#### Comparing Ti/Tv ratio across orthogonal aligners

We validated our Ti/Tv ratio findings by implementing orthogonal alignment approaches for all sets of sample pairs. We generated alignments on the same set of HPRC2 samples that we used for Centrolign with Rama and Unialigner using default parameters, and with minimap2 using parameters -x asm20 --eqx -c -f 0.05 --secondary=no -r 10000. We implemented a custom Python script, (https://github.com/miramastoras/centrolign_analysis/blob/main/scripts/pairwise_titv_from_cigars.py), that parses a set of pairwise CIGAR strings and calculates per-sample-pair Ti/Tv ratio, and overall Ti/Tv ratio across all sample pairs, as # transitions / # transversions. The script also implements an indel distance filter, removing any SNVs within 20 bp of an indel from the calculation.

### Pangenome integration

We developed a pangenome-based strategy for assigning samples to cenhap lineages based on the k-mer content of unaligned reads, taking advantage of the graph-based MSAs produced by Centrolign. It re-purposes a previously developed method for personalizing pangenome graphs in advance of read mapping^52^. This method, implemented in vg’s haplotypes subcommand, sub-samples a pangenome graphs’s haplotypes based on their k-mer similarity to sequencing reads. Since it attempts to select haplotypes that are closely-related to the sample’s own, we reason that they will also often belong to the same cenhap lineage. Accordingly, we can infer the sample’s cenhap lineage based on cenhap annotations of the pangenome’s haplotypes.

We also performed read mapping experiments for the pangenome graphs created by Centrolign. Pangenome personalization removes extraneous sequences from the pangenome before mapping, leading to more accurate read alignment. This is likely to be especially important in centromeres, where abundant similar sequences create bait for misalignment, so we also used vg’s personalization strategy for read mapping. Since the cenhap typing also utilizes the same personalization algorithm, we combine both of these two analyses into a single execution of haplotype sampling.

Code for the pangenome integration is available on GitHub at: https://github.com/faithokamoto/centromere-haplotype-sampling-pipeline. Reads were aligned with vg version v1.74.

#### Pangenome creation

To facilitate pangenome analyses in vg, we created a single pangenome reference graph per chromosome from the separate graphs that were constructed from clades of haplotypes (See “Constructing MSAs for all chromosomes”). In particular, vg’s haplotype sampling algorithm requires all haplotypes to participate in a single, nearly linear graph structure; This requires all haplotypes to pass through common nodes. Thus, we joined the graphs by connecting sentinel source and sink nodes (sequence “N”) to all haplotypes’ start and end, respectively. Finally, we converted the resulting graph to GBZ format and indexed for haplotype sampling. We refer to this combined graph as the “Centrolign graph”.

We ran typing experiments on chromosomes with assembly-based cenhap assignments (Langley and Loucks, manuscript in preparation) available chromosomes 4, 6, 9, 10, 11, 12, and 17 (excluding HG00272.1, which was not included in the cenhap resource). However, for read alignment experiments, we used all haplotypes which had reads easily accessible from the HPRC; this excluded CHM13 and HG002.

#### Input reads

For each centromeric haplotype of interest, we downloaded PacBio high-fidelity (“HiFi”) long reads from HPRC and extracted reads with alignments contained within the alpha-satillites. In addition, for each haplotype we simulated reads from haplotype with vg sim. The simulated reads had a length, count, and error profile similar to the real reads, and due to being simulated, they have known truth alignment positions. This is useful for assessing alignment correctness.

#### Pangenome personalization

For each haplotype we generated a personalized pangenome using the haplotype sampling implemented in vg haplotypes. In each case, we used the --ban-sample option to prevent the sample’s own haplotypes from being selected. This approximated a leave-one-out experiment. Haplotypes were dropped from the results if they had a score further than 500 (for haploid read sets) or 2000 (for diploid read sets) below the top-scoring haplotype. In addition, different parameters were used for haploid and diploid read sets. For haploid, we used --num-haplotypes 5 (vs. --num-haplotypes 10 for diploids) and --haploid-sampling.

We also found that, for the Centrolign graphs, it was necessary to adjust the --absent-score parameter, which adds score to haplotypes that exclude k-mers that are absent in the reads, from its default value. Otherwise, we observed that certain haplotypes had so few k-mers that were unique in the graph that, if even a few of these k-mers matched the reads, the haplotype would score highly based on the absent k-mer scoring. In the highly repetitive and variable active arrays, this could easily occur by chance without a close relationship to the haplotype, leading to frequent mis-calls. Accordingly, we decreased --absent-score parameter to 0.05 from its default value of 0.8. This parameter value was optimized on chromosome 4 (Supplementary Figure 51).

#### Cenhap typing

For each haploid read set, we assigned the haplotype to the cenhap lineage of the top-scoring sampled haplotype. We assigned the diplotype to the cenhap lineages of the first two sampled haplotypes from distinct cenhap lineages. If all sampled cenhap lineages were the same, we assigned a homozygous diplotype. We ran diploid typing experiments for all samples where both haplotypes were available in a chromosome’s Centrolign graph.

#### Haploid read alignment

We aligned each haploid read set to one pangenome reference and three linear references using Giraffe^53^ and minimap2^54^. The linear references were: CHM13, the haplotype’s own assembly, and the assembly with minimum pairwise match distance from the Centrolign alignments, which we refer to as its neighbor. The pangenome reference was the previously described personalized pangenome. To create each linear reference, we extracted a single haplotype path from the larger Centrolign graph, and then converted that single-haplotype graph into a FASTA. We ran Giraffe in “hifi” mode on the single-haplotype graph and minimap2 in “map-hifi” mode on the FASTA. Because Giraffe was consistently worse than minimap2 at aligning reads to linear references (Supplementary Figure 58), we only used minimap2’s linear alignments for further comparisons. We aligned to the pangenome using Giraffe’s “hifi” mode.

#### Diploid read alignment

We aligned each diploid read set to the CHM13 linear reference using minimap2’s “map-hifi” mode, and to three graph references using Giraffe’s “hifi” mode. The graph references were analogous to the non-CHM13 haploid conditions, except now with two haplotypes. Specifically, one consisted of the sample’s own haplotypes, one consisted of the neighbor haplotypes to the sample’s haplotypes, and one was a personalized pangenome (see “Pangenome personalization”).

#### Alignment correctness

Given high divergence in centromeres, many reads cannot project onto a single linear reference. This makes it impossible to use a matching reference position to identify a correct alignment. Thus, we used a correctness metric based on the path in the graph that generated the read, which is included in the output of vg sim. This metric takes into account that some nodes on the reads’ paths are private to each haplotype path and therefore unavailable as an alignment target in the leave-one-out experiment. We define a read’s correctness as the percent of non-private truth nodes which the test alignment overlaps, ignoring orientation. Reads which failed to align automatically have correctness of 0. For alignments to the reads’ own haplotype, we used a variation of the same formula, but including private nodes that are now available as alignment targets.

We defined truth positions for real reads as their original alignment positions in the downloaded HPRC BAM files, which are aligned against the samples’ own assemblies. Corresponding correctness calculations match up well with alignments of simulated reads that have known truth positions (Supplementary Figure 59), suggesting that the HPRC alignment positions are a good truth.

#### Computational resource usage

We ran all alignments using 20 threads on a high-performance computing cluster with 16-core AMD processors. We profiled CPU-second runtime and memory usage of each aligner using /usr/bin/time -v. Each combination of haplotype, aligner, and reference only ran once.

#### GRef Coordinates

We annotated each chromosomal centromere graph with a graph reference coordinate system using vg paths -u (vg commit 0fd8859b). This procedure covers the graph with path intervals, beginning with CHM13 and greedily extending it with uncovered path intervals. These coordinates allow variation to be represented in a consistent manner across samples, even if it has no position on CHM13. We also used vg deconstruct to generate a VCF representing the variation of the haplotypes in the graph, referenced on the pangenome coordinate system for each graph. This enabled the variation on and off CHM13 to be compared. With diploid read alignments mapped to the GRef annotated Centrolign graphs using the same methodologies described above, we used vg pack -Q 5 and vg call -zaA (on the original, not sampled Centrolign graph) to run the genotyping. bcftools merge was used to construct a single multisample VCF for each chromosome, and bcftools view was used to filter to only PASS variants. The SnakeMake workflow can be found here: https://github.com/glennhickey/nested-variants/tree/3fbd7a46792ca1db7fa8f10b3093e2ed5508c9e5/centrolign-all.

## Acknowledgements

Grants supporting this work are from the US National Institutes of Health, awarded to KHM NIH/NHGRI R01 R01HG011274, NIH/NHGRI UM1 HG010971, and BP R01HG014490, U01HG010961, U24HG010262, OT2OD026682, and U24HG011853. F.O. was supported by the NHGRI, fellowship 5T32HG012344-05. FR was supported by the HSE basic research program. NA is a HHMI Hanna H. Gray Fellow, Pew Biomedical Scholar, and Biohub Investigator. KHM was supported in part by the Searle Scholars Program.

## Author contributions

J.E., M.M. and J.K.L. contributed equally to this work. K.H.M. and B.P. jointly conceived and supervised the study. J.K.L. created the active αSat and pericentromeric array lists used throughout the study and led pericentromeric structural typing and cenhap association analyses. J.E. designed and implemented the Centrolign algorithm. J.E. and M.M. developed the simulation framework used for benchmarking. J.E. and M.M. performed Centrolign benchmarking. M.M. ran Centrolign for the HPRC2. M.M., J.K.L, and J.E. analyzed centrolign results. F.R. developed the HOR-StV annotation pipeline, HORhap clustering methods, and the Annotaligner tool. F.R. and I.A.A. analyzed interchromosomal mixing. J.M. developed the aggregate and read-based CDR annotation methods and analysis. G.H. performed Graph Reference (GRef) analysis. F.O. developed the pangenome haplotype-sampling pipeline for cenhap typing and as well as read alignment. J.K.L, H.L., S.A.L., and C.H.L. led cenhap lineage construction and assignment with contributions from Y.Z. F.R., J.K.L, J.M., N.A., J.M.F., and I.A.A. contributed to satellite annotation methodology and interpretation of centromere evolution. P.H. led the gene analysis. K.H.M. and B.P. acquired funding and supervised the project. J.E., M.M., J.K.L., J.M., F.O., P.H., K.H.M., and B.P. wrote the manuscript with input from all authors. All authors reviewed and approved the final manuscript.

