## Supplement Note 1 for "Pangenome alignment reveals global diversity and evolution of human centromeric regions"

### Supplementary Note 1. Interchromosomal exchange in centromeres

Active α-satellite arrays in humans evolve from 4 distinct ancestral repeats (designated by SFs 1-4), and each SF comprises repeat families organized into chromosome-specific lineages[1,2]. However, the active arrays of some nonhomologous chromosomes are highly similar, particularly those of the acrocentric chromosome pairs 13/21 and 14/22 and 1/5/19 triplet (**Supplementary Fig. 18**) suggesting some kind of sequence exchange among these chromosomes, as previously proposed [3–5]. Across primate species, rDNA-bearing chromosomes always have closely related centromeres sharing SF or subSF assignments, and if they change their SF identity, it occurs in a coordinated manner, which was dubbed the "rDNA-centromere coordination model"[6]. Human acrocentric chromosomes all have SF2 centromres and rDNA gene clusters in their short arms. The latter are known to associate within the nucleolus and undergo recurrent recombination between non-homologs in their p-arms[7].

#### Chromosome 13/21 recombinations

Centromeres of chromosomes 13 and 21 are highly similar and share the same active HOR, S2C13/21H1L, and therefore cannot be distinguished at the monomer level. However, chromosome-specific differences at the HOR level were identified in CHM13[4]. Human acrocentric chromosomes undergo recurrent interchromosomal exchange within their rDNA-bearing short arms[7]. In marmoset, all rDNA-bearing chromosomes were shown to have similar centromeres belonging to the same suprachromosomal family, suggesting ongoing exchange among these chromosomes which includes centromeres[6]. However, clear evidence for exchange extending into acrocentric centromeres, including between human chromosomes 13 and 21, has not previously been demonstrated.

The active-array k-mer clustering tree (Methods) constructed from the combined HPRC2 chr13 and chr21 dataset did not separate the arrays into two chromosome-specific clades. Instead, the tree contained intermittent chr13- and chr21-associated deep branches, suggesting a shared evolutionary history (**Fig. SN1a**). Most arrays nevertheless fell within two major branches associated with their chromosomes of origin, but both major branches included several exceptions indicating more recent interchromosomal exchange.

Three chr21 active arrays (HG01252.1, HG01884.2, and NA20282.2), together with HG01261.pat, which failed HPRC QC but was confirmed in the Verkko dataset, clustered within the major chr13-associated orange branch and were closely related to typical chr13 arrays (Fig. SN1a). This pattern suggests relatively recent recombination events in which chr21 acquired a centromere derived from chr13. HORhap analysis independently supported this interpretation: whereas typical chr21 arrays are dominated by the blue HORhap, these four arrays were represented by the orange/red HORhaps characteristic of chr13 (**Fig. SN1b,e**).

Reciprocal cases, in which chr13 carried a chr21-like centromere, were observed in an orange outgroup branch adjacent to the major blue chr21 branch (**Fig. SN1a**), containing eight chr13 arrays. HORhap analysis showed that these arrays were dominated by the blue chr21-associated

HORhap, independently supporting exchange with chr21. Their q-arm edges retained some orange chr13-associated HORhap, suggesting that part of the original chr13 array remained and that the recombination breakpoint may lie within the active array (**Fig. SN1c**, top). In four of the eight arrays, an additional recombination event introduced normal chr13-associated red/orange HORhaps, producing active arrays composed of three distinct segments (**Fig. SN1c**, bottom). Importantly, only the first event represents exchange between chromosomes 13 and 21; the second occurred between two chr13 arrays – one normal orange and another chr21-derived blue.

One additional case, HG01960.mat.chr13, showed a particularly clear transition within the active αSat array. Although this sample failed HPRC QC, the same structure was confirmed in the Verkko assembly. The q-arm half of the array contained the olive HORhap characteristic of a minor chr13 branch (**Fig. SN1e**), whereas the p-arm half contained the blue HORhap characteristic of the major chr21 branch, revealing a recombination breakpoint in the center of the active array (**Fig. SN1d**).

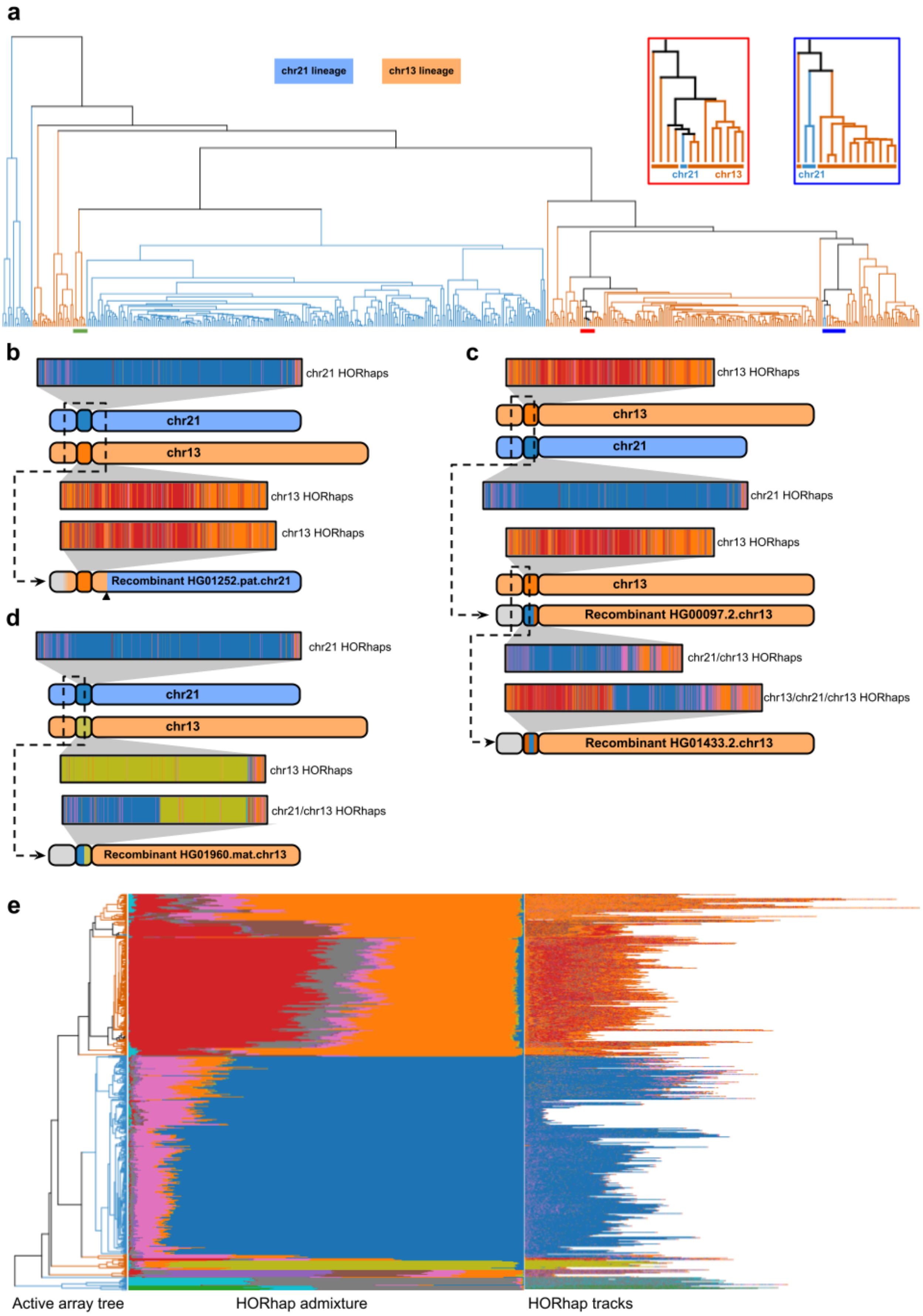
a
chr21 lineage
chr13 lineage
chr21
chr13
chr21
b
chr21 HORhaps
chr21
chr13
chr13 HORhaps
chr13 HORhaps
Recombinant HG01252.pat.chr21
c
chr13 HORhaps
chr13
chr21
chr21 HORhaps
chr13 HORhaps
chr13
Recombinant HG00097.2.chr13
chr21/chr13 HORhaps
chr13/chr21/chr13 HORhaps
Recombinant HG01433.2.chr13
d
chr21 HORhaps
chr21
chr13
chr13 HORhaps
chr21/chr13 HORhaps
Recombinant HG01960.mat.chr13
e
Active array tree
HORhap admixture
HORhap tracks

**Fig. SN1. Chr13/chr21 recombinant centromeres. a.** Active-array k-mer clustering tree of chr13 and chr21 assemblies. The close-ups of red and blue underlined regions show position of chr21 samples with recombined centromeres from chr13 and their close relation to normal chr13 arrays. Green underline shows location of chr13 samples with chr21-like centromeres. **b-c-d.** Cartoons showing recombination events. **e.** HORhap tracks of the assemblies (right) and admixture graphs (center) sorted according to the same tree as in SN1a (left). Admixture graphs show the proportion of each HORhap in a given array. The panel shows that typical chr22 arrays are dominated by blue HORhap, while typical chr13 arrays are mostly formed by red/orange HORhaps.

To obtain an independent confirmation and further investigate the recombination breakpoint, one chr21 haplotype carrying a chr13-like centromere, HG01252.1.chr21, was mapped with minimap2 simultaneously to representative chr13 and chr21 haplotypes. HG01192.2.chr21 was selected as a representative of the major chr21 branch in the active-array clustering tree, whereas HG00344.2.chr13 was selected as the chr13 array most closely related to HG01252.1.chr21 in the same tree. Minimap2 revealed different chromosome identities across the recombinant chromosome (**Supplementary Fig. 17a**). The q-arm was confidently assigned to chr21. In contrast, the p-arm flank adjacent to the centromere corresponded to a region of high sequence similarity between chr13 and chr21 visible as overlapping minimap2 tracks. The active αSat array (brown), HSat array (green), and the right half of the p-arm showed stronger homology to chr13, whereas the left half of the p-arm showed stronger homology to chr21. These p-arm assignments should be interpreted cautiously because of the highly repetitive structure and extensive sequence similarity between chr13 and chr21 in this region.

A 400-kb window spanning the overlap between the chr13 and chr21 mappings was therefore extracted and aligned to the corresponding sequences from both references. Overall, all 3 sequences were 98.9-99.5% identical in this interval, therefore it should be technically described as a segmental duplication (SD). In the left part of this window, the recombinant showed slightly higher similarity to chr13, whereas in the right part it showed slightly higher similarity to chr21. At the transition between chr13- and chr21-like sequence, a 531-bp interval displays no differences among the three sequences and therefore presumably contains the recombination breakpoint. RepeatMasker annotation identified Alu and L1 elements within this interval (**Supplementary Fig. 17b**).

#### Chr14/22 recombinations

A similar analysis was performed for chromosomes 14 and 22, which also have highly similar active arrays with HORs that are indistinguishable at the monomer level (S2C14/22H1L). The active-array k-mer clustering tree initially showed no clear cases of arrays from one chromosome falling within a clade characteristic of the other. However, such cases emerged when samples excluded during QC were included in the analysis (**Fig. SN3a**).

One case, supported by HORhap analysis in both the HPRC and Verkko assemblies, involved HG02984.pat.chr14, whose active array clustered with chromosome 22 arrays and showed a chromosome 22-like HORhap composition (blue close-up in **Fig. SN3a**; **Fig. SN3b**). Two reciprocal cases involved chromosome 22 arrays within the chromosome 14 clade (red close-up in **Fig. SN3a**). HG02841.mat.chr22 centromere was composed predominantly of the light-green HORhap characteristic of chromosome 14, consistent with interchromosomal exchange (**Fig. SN3c**). In the HPRC assembly, this array was split between two contigs, whereas the Verkko assembly contained it in a single contig.

HG01258.pat.chr22 was more ambiguous. Its αSat array was split across two (HPRC) or three (Verkko) contigs. One contig showed the orange/violet/pink HORhap composition typical of chromosome 22, whereas another contig (HPRC) or two contigs (Verkko) showed the light-green HORhap composition typical of chromosome 14. This pattern could represent a hybrid active array, similar to HG01960.mat.chr13 (above), but an assembly error cannot be excluded, for example if the chromosome 14 contig was incorrectly assigned to chromosome 22.

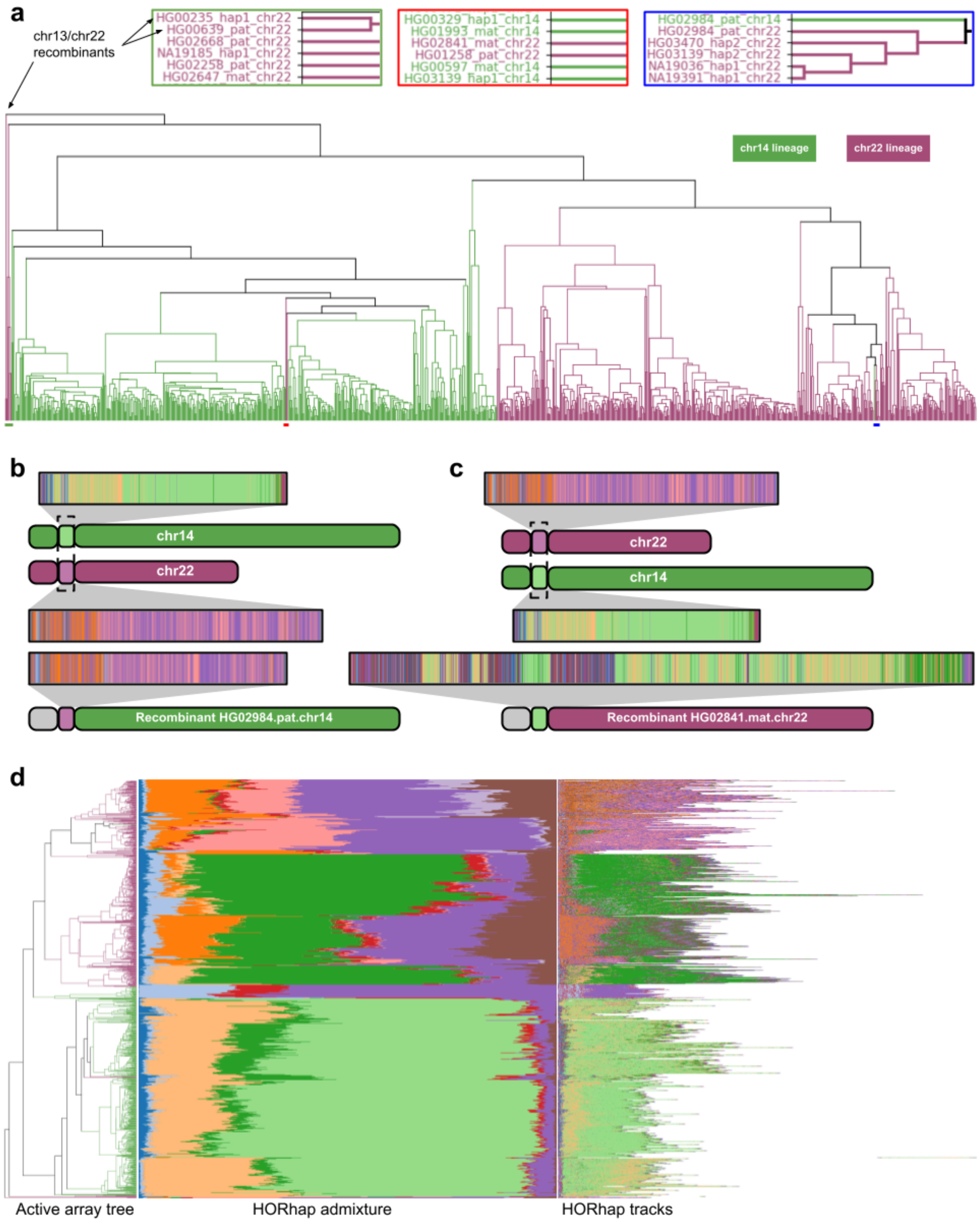


**Fig. SN3. Chr14/22 recombination. a.** Active-array k-mer clustering tree of chr14 and chr22 assemblies. QC-failed arrays are also included. Close-ups show suspected recombinants. **b-c.** Cartoons showing recombination events. **e.** HORhap tracks (left) and admixture graphs (center) are sorted according to the same tree as in SN1a (left).

#### Chr13/22 recombination

The deepest branch of the chr14/chr22 array tree (green close-up in Fig. **SN3a**), containing HG00235.1.chr22 and HG00639.1.chr22, represents previously reported recombinants between acrocentric chromosomes with dissimilar αSat arrays — chromosomes 13 and 22[8]. A third such recombinant reported in Gao 2026[8] was not present in either the HPRC or Verkko datasets. HOR/CenSat annotation of both arrays clearly showed a hybrid structure, with chromosome 13-associated αSat (S2C13/21H1L) at the p-arm edge and chromosome 22-associated αSat (S2C14/22H1L) at the q-arm edge and in the central part of the array.

Both arrays were analyzed using HORhap HMMs generated for the chr13/21 and chr14/22 chromosome pairs (Fig. **SN4**). This analysis confirmed previous observations and showed that the chromosome 13-derived portion, represented by the orange HORhap, is closely related to typical chromosome 13 arrays (Fig. **SN2e**), whereas the chromosome 22-derived portion, represented by the violet/pink HORhaps, is closely related to typical chromosome 22 arrays (**Fig. SN3d**).

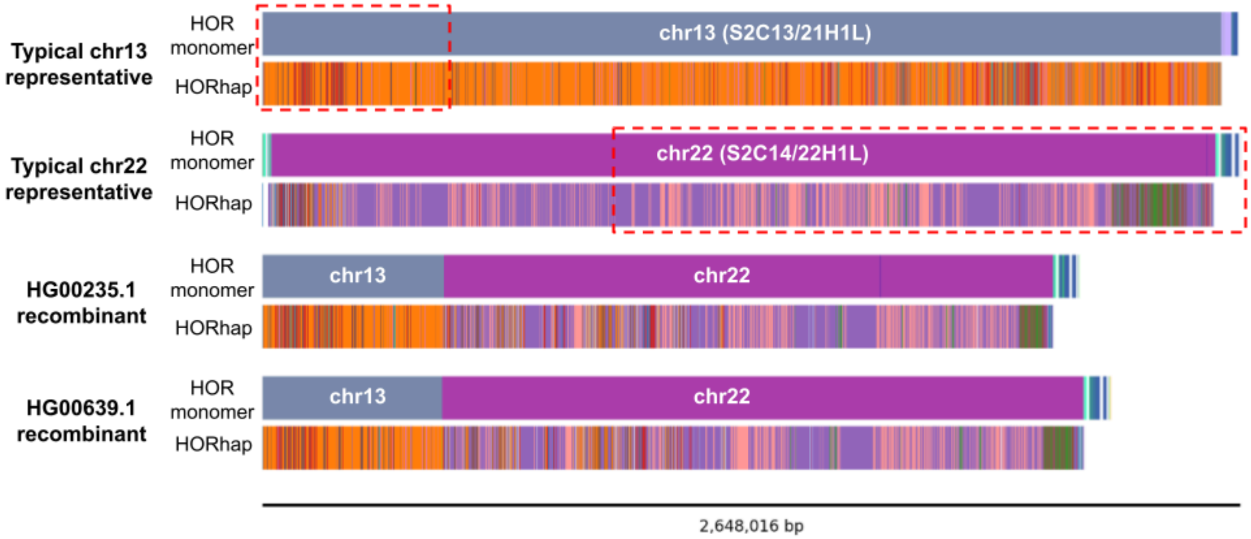


**Fig. SN4. Chr13/22 recombinants.** The top panels show HORhap annotations of chr22 and chr13 array representatives (HG02738.2.chr13 and NA19909.1.chr22). Red boxes indicate the regions that, when combined, can form the recombinant structures similar to the ones shown in the bottom panels.

#### Chr19 active-array insertion originating from chr10

The cases described in the sections above represent interchromosomal αSat mixing resulting from recombination between nonhomologous acrocentric chromosomes which are brought together in a nucleolus by virtue of rDNA gene clusters contained within their short arms. In this section, we consider a different case which features insertion of a fragment from the active array of one chromosome into the active array of another. Specifically, an approximately 250-kb fragment from the chromosome 10 centromere, including the active HOR S1C10H1L and neighboring HORs from the q-arm edge of the centromere (S1C10H1-B and S1C10H2), was inserted into the chromosome 19 active array (HOR S1C1/5/19H1L). The insertion was observed in three samples: HG01109.2, HG02055.2, and HG03209.1. All three failed HPRC QC because of either a sequence gap within the centromere (black in Fig. **SN5a**) or fragmentation of the centromeric region across multiple contigs. However, the corresponding Verkko assemblies for all three samples contained the same insertion. In addition, 60-mers spanning the insertion breakpoints were supported by short-read data from all three samples (**Table SN1**). These breakpoint k-mers were absent from the chromosome 19 active arrays of all other HPRC assemblies and from two tested short-read datasets obtained for individuals without insertion.

It is unclear whether these insertions occurred independently or were inherited from a common ancestor. On one hand, the samples represent different populations and geographical locations: one is from the Americas (PUR), whereas the other two are from Africa (ACB and MSL). On the other hand, the insertions occur in the same sequence context, with identical 60-bp windows surrounding the breakpoints in all three arrays. The inserted sequences themselves are also highly identical (over 99.988%), with their differences explainable by a small number of simple SNVs and indels. Moreover, a k-mer-based clustering tree of chromosome 19 active arrays, generated after excluding the chromosome 10 insertion, placed the American and one African sample as the two closest arrays, with the second African sample in the same orange branch (**Fig. SN5c**). Also note that the American PUR population is known for its high proportion of African ancestry. Together, these observations are more consistent with a shared origin of the insertion, although independent events cannot be excluded.

This event may represent an example of inter-array seeding, previously proposed as a mechanism of αSat evolution[1,3]. Under this “interchromosomal exchange/amplification model”, αSat sequence from one chromosome can be inserted into the centromere of another chromosome, and under favorable conditions it may subsequently expand, displace the old array and eventually emerge as a new centromere with only the remnants of the old array on the flanks. Such a mechanism would explain spreading of the “new SF” (SF 1-3) αSat across most African apes’ centromeres, with each SF presumably originating on a single chromosome, but subsequently each spreading onto a group of chromosomes[1,3]. It was further proposed by the “kinetochore selection hypothesis” that the circumstances favorable for expansion occur if the introduced sequence is capable of anchoring the kinetochore formation which presumably drives or directs the expansion process[4,9].

The chromosome 10 insertion into chromosome 19 may therefore represent an early-stage example of such inter-array seeding. If the introduced sequence were better suited for kinetochore, it could in principle expand further and eventually occupy a larger part of the chromosome 19 centromere. However, both chromosome 10 and 19 HORs belong to SF1, so the functional differences between them may be too small to provide for a strong kinetochore preference. Nevertheless, this event provides direct evidence that relatively large αSat fragments can be transferred between active arrays of different chromosomes, consistent with the type of seeding event proposed in the models of centromere evolution. Such alien insertions in active arrays were also recently reported in marmosets[6].

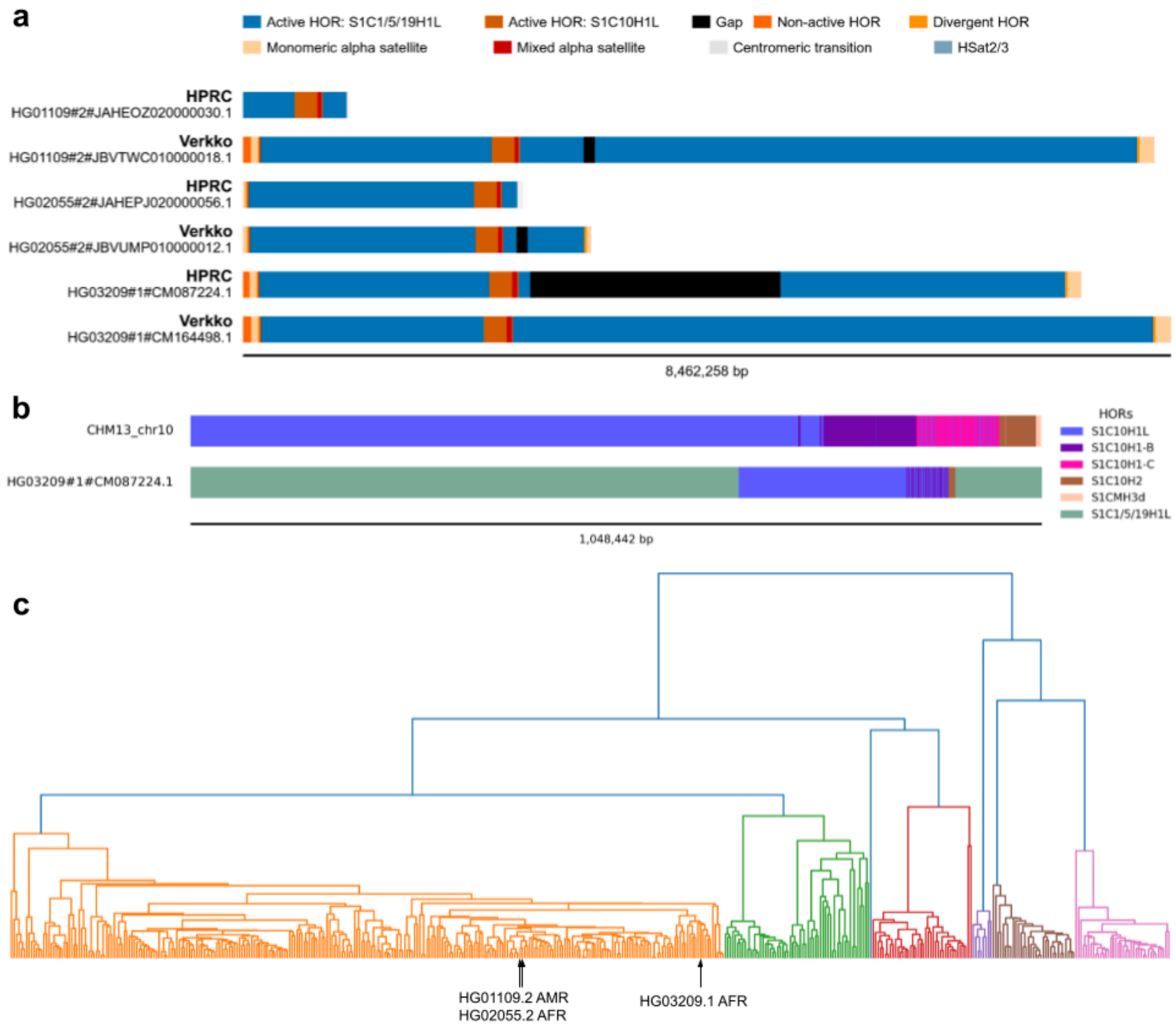

**Fig. SN5. Insertion of cen10 fragment to cen19. a.** CenSat tracks for contigs of chr19 active arrays with insertions from chr10 centromere. Active arrays of chr19 and chr10 are colored differently. **b.** HOR monomer track close-ups of CHM13 chr10 centromere q-edge and one of the contigs from above show that inserted fragments likely belong to q-edge of chr10 centromere. **c.** A k-mer-based clustering tree of chromosome 19 active arrays generated after excluding the chromosome 10 insertion from the three arrays which are shown with black arrows.

**Table SN1.** Counts of chr19/chr10 breakpoint-targeting 60-mers in assemblies and short read samples.

| sample name | chr10 insertion | Left_breakpoint | Right_breakpoint |
|---|---|---|---|
| HG01109.2 assembly | present | 1 | 1 |
| HG02055.2 assembly | present | 1 | 1 |
| HG03209.1 assembly | present | 1 | 1 |
| all other HPRC assemblies * | absent | 0 | 0 |
| HG01109 (ERR3988842) | present | 8 | 13 |
| HG02055 (ERR3988979) | present | 5 | 10 |
| HG03209 (ERR3242539) | present | 8 | 17 |
| HG00096 (ERR3240114) | absent | 0 | 0 |
| HG03015 (ERR3242636) | absent | 0 | 0 |

* — to reduce computations, only chr19 active array sequences were checked, not entire assemblies.
Left_breakpoint
CTTCGTTGGAAGCGGGATTTCTTCATATTCTGCTAGACAGAAGAATTCTCAGTCACTTCT
Right_breakpoint
CAGAGCAGTTTTGAAACACTCTTTTTGTGGAATTTGCAAGTGGATATTTCAGCCGCTTTG

Summary on interchromosomal exchange in centromeres:

1. All non-homolog exchange cases found so far involve arrays that belong to the same SFs: SF2 in acrocentrics and SF1 in the chr10/19 case.
2. Same-HOR exchanges within acrocentric 13/21 and 14/22 paired arrays seem to be recurrent and feature both recent independently generated cases and some historical events which gave birth to particular centromere haplotype clades that persist in population.
3. Different-HOR exchanges observed here may represent progeny of anecdotal single events, although further investigation is necessary.
4. Sequence exchange in acrocentrics is not limited to short arms, it can also involve centromeres with breakpoints either in centromeric satellite arrays (yielding hybrid arrays) or in SDs of the q-arm (then entire centromere belongs to a different chromosome).
