## Supplement Note 2 for "Pangenome alignment reveals global diversity and evolution of human centromeric regions"

### 1 Identifying minimal rare matches

Here we describe an algorithm that identifies minimal $(n, m)$-rare matches between two sequence graphs using a generalized suffix tree (GST) over all of the sequences used to construct the graphs. The GST is augmented by Hui's (1) color set size (CSS) data structure, which can query the number of unique nodes (from the input graphs) that can serve as the start of the suffixes represented by a suffix array interval. Our algorithm uses two of these structures, one returning the count for nodes from each of the two input graphs.

In order for an $(n, m)$-rare match to minimal, it must not be possible to shorten the match without introducing additional occurrences. Our algorithm ensures this condition in different ways for the left and right ends of the match. We first consider the right end.

Consider the path from the root of the GST walked by a minimal $(n, m)$-rare match. The match will end along some edge in the graph pointing from $v$ to $c$. We claim that it must end after matching exactly one character of this edge. This follows from the fact that every match (not necessarily minimal) ending on $(v, c)$ occurs at the same positions (which are the leaves of $c$'s subtree) and therefore occurs the same number of times. If a match ends at any other position on $(v, c)$, it can be shortened on the right end without changing its count and is therefore not minimal.

For querying minimal rare matches between two strings, this necessary condition for minimality on the right end is also sufficient. In strings, $v$'s other children indicate additional occurrences of the match if the rightmost character were removed. In a graph, they may instead indicate different neighbors of the same match. Thus, we must directly check the right end minimality condition by comparing the number of occurrences of $c$ and $v$ using the CSS indexes. The match is minimal on the right end if $v$ occurs more times than $c$ on at least one of the graphs.

Next we consider verifying the minimality of the match on the left end. In a suffix tree, the suffix link from a non-root node $v$ points to the node whose root-to-node path has the leftmost character removed, relative to $v$'s root-to-node path. We know that a minimal match must end on some edge from $v$ to $c$, which it overlaps by one character. To verify minimality, we can follow $v$'s suffix link to $w$ and then follow the edge whose initial character matches that of $(v, c)$. Let $d$ be the node this edge points to. Using the CSS indexes, we can then compare the number of occurrences that $c$ and $d$ have. The match is minimal on left end if and only if at least one of $d$'s counts is higher than that of $c$. Figure 1 shows a worked example of a minimal rare match.

Thus, we have necessary and sufficient conditions for minimality on both the left and right sides of a match. Our algorithm iterates over the internal nodes of the GST and considers the matches that extend one character onto each of their downward edges. If these matches meet both the left and right end minimality conditions, then they are minimal $(n, m)$-rare matches. Since, the CSS indexes perform count queries in $O(1)$ time, there is $O(1)$ work per edge, which gives $O(N)$ total run time, where $N$ is the size of the GST. Pseudocode is presented in Algorithm 1.

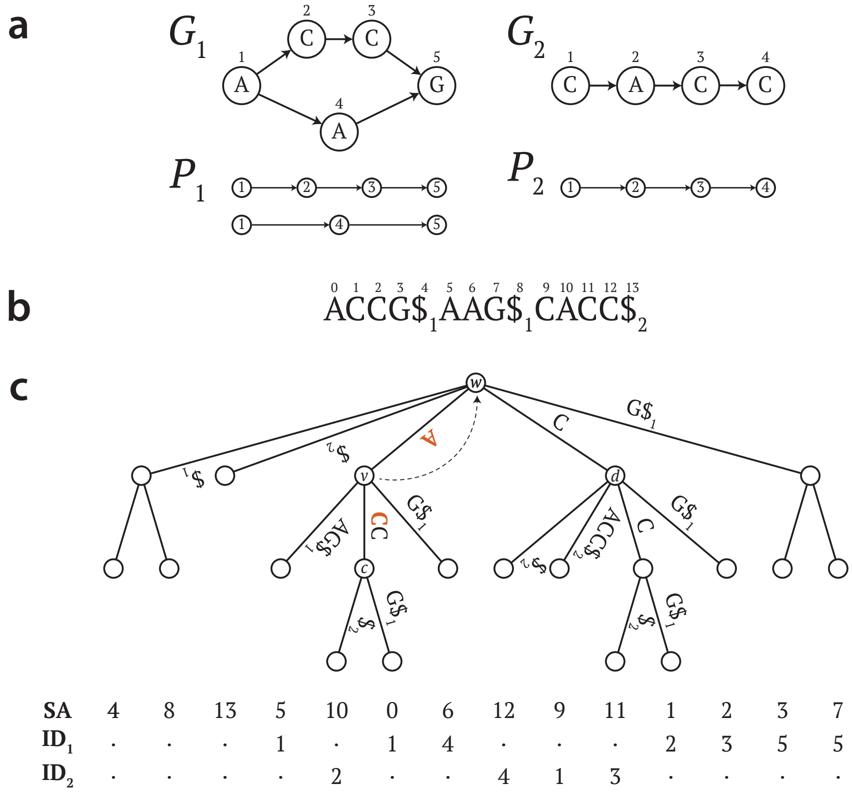


Figure 1: **Illustration of a minimal rare match**. **a** Two graphs $G_1$ and $G_2$ with path covers $P_1$ and $P_2$. **b** The concatenated path sequences, separated by sentinels, which is used to construct the generalized suffix tree. **c** The generalized suffix tree with a minimal rare match highlighted in red. The match extends by one character onto the edge $(v, c)$. The suffix link from $v$ to $w$ is shown as a dotted arrow (other suffix links not shown). $d$ is the child of $w$ that is reached by following the edge of the same label as $(v, c)$, which is C. $c$'s subtree includes 1 occurrence in $G_1$ and 1 in $G_2$ (shown in $\text{ID}_1$ and $\text{ID}_2$). $d$'s subtree includes 2 occurrences in $G_1$ and 3 in $G_2$. $v$'s subtree includes 2 occurrences in $G_1$ and 1 in $G_2$ Since both $d$ and $v$ have additional occurrences compared to $c$, AC is a $(1, 1)$-rare match.

**Algorithm 1:** Identify minimal rare matches

```
Input: Generalized suffix tree T (implemented as an enhanced suffix array), color set size
       indexes C1 and C2 for the two input graphs, and maximum count k
Output: Suffix array intervals and match lengths corresponding to minimal (n, m)-rare matches
Function MinimalRareMatches(T, C1, C2, k):
    M ← ∅ // set of matches
    foreach internal node v in T do
        p ← C1.colorCount(v)
        q ← C2.colorCount(v)
        foreach c ∈ T.children(v) do
            n ← C1.colorCount(c)
            m ← C2.colorCount(c)
            if n = 0 or m = 0 or n · m > k or (n = p and m = q) then
                // No match, count is above max, or not minimal on right end
                continue
            end
            if v is not the root then
                w ← T.suffixLink(v)
                l ← T.edgeLabel(v, c)
                d ← T.childWithLabel(w, l)
                if n = C1.colorCount(d) and m = C2.colorCount(d) then
                    // Not minimal on left end
                    continue
                end
            end
            // Record a minimal (n, m)-rare match
            M.add((T.SAInterval(c), T.depth(v) + 1))
        end
    end
    return M
```

#### 2 Chaining algorithms for graph-graph alignment

Here we describe the sparse dynamic programming algorithms used to chain matches together in Centrolign. There are two such algorithms: one with no gap penalties, and one with affine gap penalties. Both are based on the path cover technique originally published in (2).

A path cover is a set of paths through the graph such that every node is included in at least one path. We assume that a path cover has been computed for both graph $G_1 = (V_1, E_1)$ and $G_2 = (V_2, E_2)$, which are being aligned—in Centrolign, we use the set of input sequences as the path cover. We denote the path covers as $P_1$ and $P_2$, and we denote their sizes as $k_1 = |P_1|$ and $k_2 = |P_2|$. We drop the subscripts when writing generically about either graph.

The chaining problem takes as input a set of matches $M$. Each match consists of two walks $w_1$ and $w_2$ in $G_1$ and $G_2$, respectively, which have the same sequence. We denote the number of matches as $N = |M|$. Each match is associated with a numerical value by a function `weight`. We also use convenience functions `begin` and `end` to return the first or last node in a walk, respectively.

##### 2.1 Support data structures

Both chaining algorithms use a set of preprocessed data structures. Algorithm 2 shows these preprocessing steps. Two of these structures have non-trivial construction algorithms, which are cited but not reproduced. The first is *LastToReach*, which was also previously studied under the name *ChainMerge* (3). It is defined by

$$LastToReach[v, p] = \max \{k \mid p[k] \text{ can reach } v \text{ in } G\}, \tag{1}$$

where $v \in V$ and $p \in P$. The second of data structure, $D$, is only used in the affine gap algorithm(4). It is defined by

$$D[v,p] = \text{length of the shortest walk in } G \text{ from } p[\mathit{LastToReach}[v,p]] \text{ to } v. \quad (2)$$

Both data structures can be constructed in $O(k|E|)$ time using previously published algorithms(2; 4).

#### 2.2 Chaining with free alignment gaps

This algorithm is based on Section 6.2 of (2). Psuedocode is shown in Algorithm 3 .The sparsity in the dynamic programming is based on a dynamic RangeMaxQuery (RMQ) data structure for key-value pairs. This structure must be able to insert key-value pairs and return the greatest value (if any) within an arbitrary range of keys. To provide this interface, we use a balanced binary search tree whose nodes are augmented with pointers to their subtree's maximum value. This data structure can perform the `insert` and `rangeMax` queries in $O(\log N)$ time.

The overall run time of the algorithm is dominated by the loop that contains the `rangeMax` queries. By the definition of the *LastToReach* structure, there are $k_1$ forward edges to each node. Thus, the execution will enter the innermost loop $k_1$ times for each match. In this loop, it will execute `rangeMax` $k_2$ times with an RMQ containing at most $N$ entries. Thus, the total run time is $O(k_1 k_2 N \log N)$.

As noted in Lemma 3.3 of (2), it is possible to improve the runtime to $O(k_1 k_2 N \log\log N)$ using van Emde Boas trees instead of balanced binary trees. However, this is not part of our implementation. Centrolign achieves a constant factor improvement by noting that this algorithm remains correct even if dynamic programming values are not inserted into every RMQ but rather a single RMQ corresponding to an arbitrary pair of a path $p_1 \in P_1$ containing $v$ and $p_2 \in P_2$ containing $\text{end}(M[i].w_2)$. This reduces the time spent in the first inner loop to $O(N \log N)$, although the overall worst case run time is not affected.

#### 2.3 Chaining with affine gap penalties

This algorithm is based on Algorithm 4 of (4), which employs additional techniques compared to the free gap algorithm. First, the length of gaps is determined by the minimum difference in lengths between the two anchors along a particular set of paths in the graph. Each $p \in P$ can generate one of these paths. The paths are chosen so that the lengths 1) are efficient to compute, and 2) consist of two additive terms, one that depends only on the earlier anchor and one that depends only on the later one. In particular, between an anchor ending on $u$ and some $p \in P$ such that $u = p[k]$, the distance to an anchor starting at $v$ is given by

$$\delta_p(u,v) = \underbrace{\mathit{LastToReach}[p,v] + D[p,v]}_{\text{depends only on } v} - \underbrace{k}_{\text{depends only on } u}. \quad (3)$$

The length of a gap in the chain is the absolute difference between the lengths on two of these paths. That is, for an anchor ending at $u_1$ in $G_1$ and $u_2$ in $G_2$ and another anchor starting at $v_1$ in $G_1$ and $v_2$ in $G_2$, the gap length between them corresponding to paths $p_1 \in P_1$ and $p_2 \in P_2$ is given by $|\delta_{p_1}(u_1,v_1) - \delta_{p_2}(u_2,v_2)|$.

The value of having two additive terms is that it allows functions of the distance to be split up between the index and the query. For example, the sign of the term inside the absolute value of the distance formula is determined by:

$$\begin{aligned}
\delta_{p_1}(u_1,v_1) &\lesseqqgtr \delta_{p_2}(u_2,v_2) \\
\mathit{LastToReach}_1[p_1,v_1] + D_1[p_1,v_1] - k_1 &\lesseqqgtr \mathit{LastToReach}_2[p_2,v_2] + D_2[p_2,v_2] - k_2 \\
\underbrace{k_2 - k_1}_{\substack{\text{depends only on } u_1 \text{ and } u_2,\\ \text{entered into index}}} &\lesseqqgtr \underbrace{\mathit{LastToReach}_2[p_2,v_2] - \mathit{LastToReach}_1[p_1,v_1] + \mathrm{D}_2[p_2,v_2] - D_1[p_1,v_1]}_{\substack{\text{depends only on } v_1 \text{ and } v_2,\\ \text{provided by query}}}
\end{aligned} \quad (4)$$

Conditioned on this sign, the absolute value in the gap penalty formula can be removed, and the gap penalty becomes affine function of the two distances. Like the difference in distances, this function can split into additive terms that each only depend on one of the two anchors. For example, if we have $\delta_{p_1}(u_1, v_1) < \delta_{p_2}(u_2, v_2)$, then the gap penalty is given by

$$
\begin{aligned}
\text{gap}_{p_1,p_2}(u_1, u_2, v_1, v_2) &= g + e \cdot (\delta_{p_2}(u_2, v_2) - \delta_{p_1}(u_1, v_1)) \\
&= \underbrace{e \cdot (k_1 - k_2)}_{\substack{\text{depends only on } u_1 \text{ and } u_2, \\ \text{entered into index}}} \underbrace{+ g + e \cdot (LastToReach_2[p_2, v_2] - LastToReach_1[p_1, v_1] + \mathrm{D}_2[p_2, v_2] - D_1[p_1, v_1])}_{\substack{\text{depends only on } v_1 \text{ and } v_2, \\ \text{added after querying}}}
\end{aligned}
\tag{5}
$$

Accordingly, the earlier anchors' values are added to the index, and the later anchors' values are computed at query time.

The dynamic programming values are entered into two-dimensional OrthogonalRangeMaxQuery (OMRQ) data structures. The OMRQ uses two keys for each value, and the `rangeMax` queries take two intervals, one for each key. The queries then return the maximum value among entries (if there are any) for which both keys fall into their respective interval. Using Equation 4, the chaining algorithm uses the additional key dimension to constrain the query to values in which the gap takes a certain sign using Equation 4. There is a separate OMRQ for each of these cases, and the gap penalties are entered into each one assuming the corresponding signedness as in Equation 5.

We provide the OMRQ interface using a two-dimensional layered range tree (5). Once again, we augment the data structure with subtree maximum pointers. It then provides `insert` and `rangeMax` queries in $O(\log^2 N)$.

The run time of this chaining algorithm is dominated by the two inner loops in the dynamic programming. In each of these loops, the body of the innermost loop executes at most once on every match per pair of paths from the path covers (one from $P_1$ and one from $P_2$). In both cases, the body of the loop performs a constant number of ORMQ operations. Accordingly, the total run time is $O(k_1 k_2 N \log^2 N)$

##### 2.3.1 Theoretical improvements

Although it is not implemented in Centrolign, we note for the sake of theoretical interest that it is also possible to reduce this run time by a factor of $O(\log \log N)$ in the word RAM model of computation. Moreover, the technique applies equally to the algorithm of (4), improving its run time by the same factor. The speed-up results from a more specialized data structure to perform two-dimensional ORMQ.

First, we note that the keys that are entered into the ORMQs depend only on the match values in $M$. As such, they can be computed before executing dynamic programming and used to initialize the ORMQ, with the initial values set to $-\infty$. This allows the ORMQ to have static keys. The `insert` operations can then be replaced with `update` operations, which change the value associated with a key pair but do not introduce a new key pair.

Our proposed data structure has a similar structure to a layered range tree. It includes an outer balanced binary search tree that is sorted by the first of the two keys. Unlike the typical layered range tree, the internal nodes are not associated with a one-dimensional range tree sorted by the second of the keys. Rather, they are associated with a sorted array, and the sorted arrays are connected via fractional cascading links (6). At the root, we can find the interval over the second key that corresponds to the query using binary search, after which the interval at each subsequent child can be determined in $O(1)$ time by following the links. Thus, this has the effect of reducing queries from an RMQ over key-value pairs to queries to an RMQ over an array of values.

Finally, we propose to use the data structure of (7) to perform RMQ operations for each array. In the word RAM model, it is capable of performing array `rangeMax` queries in $O(1)$ time and `update` operations in $O(\log N / \log \log N)$ time. Since the layered range tree must perform $O(\log N)$ such operations per `rangeMax` and `update` call to the ORMQ, the overall time of these operations becomes $O(\log N)$ and $O(\log^2 N / \log \log N)$, respectively. Thus, the runtime for the full chaining algorithm becomes dominated by the `update` calls, which take a total of $O(k_1 k_2 N \log^2 N / \log \log N)$ time.

**Algorithm 2:** Initialize data structures for sparse dynamic programming

**Input:** Graphs $G_1$ and $G_2$, path covers $P_1$ and $P_2$, matches $M$ where each match consists of a walk from each graph: $M[i] = \{(w_1, w_2)_i\}$.
**Output:** Data structures that are used in sparse dynamic programming

**Function** `InitializeSparseDP`($G_1$, $G_2$, $P_1$, $P_2$, $M$):
  Construct $LastToReach_1$ and $LastToReach_2$ from $G_1$ and $G_2$ using Lemma 3.1 of Mäkinen, et al. (2019)
  `// Invert one` $LastToReach$ `table`
  **foreach** $v \in V_1$ **do**
    $Forward[v] \leftarrow \{(u, p) \mid LastToReach_1[u, p] = v\}$
  **end**
  `// Record the beginning and end of each match`
  **foreach** $v \in V_1$ **do**
    $B[v] \leftarrow \{i \mid \text{begin}(M[i].w_1) = v\}$
    $E[v] \leftarrow \{i \mid \text{end}(M[i].w_1) = v\}$
  **end**
  Construct $D_1$ and $D_2$ from $LastToReach_1$ and $LastToReach_2$ using Lemma 3 of Chandra & Jain (2023)

**Algorithm 3:** Compute the heaviest weight co-linear chain

**Input:** $G_1$, $G_2$, $P_1$, $P_2$, and $M$ as in Algorithm 2.
**Output:** Co-linear chain of matches that maximizes total weight

**Function** `MaxWeightChain`($G_1$, $G_2$, $P_1$, $P_2$, $M$):
  `// Initialize` $LastToReach_1$, $LastToReach_2$, $Forward$, $B$, `and` $E$
  `InitializeSparseDP`($G_1$, $G_2$, $P_1$, $P_2$, $M$)
  `// Initialize sparse query data structures`
  **foreach** $p_1 \in P_1$, $p_2 \in P_2$ **do**
    $Tree[p_1, p_2] \leftarrow$ empty RMQ
  **end**
  `// Initialize dynamic programming table`
  **foreach** $M[i] \in M$ **do**
    $DP[i] \leftarrow \text{weight}(M[i])$
  **end**
  `// Do sparse dynamic programming`
  **foreach** $v \in V_1$ in topological order **do**
    `// Enter matches ending here into sparse query structures`
    **foreach** $i \in E[v]$ **do**
      **foreach** $p_1 \in P_2$ such that $v \in p_1$ **do**
        **foreach** $p_2 \in P_2, k$ such that $p_2[k] = \text{end}(M[i].w2)$ **do**
          $Tree[p_1, p_2].\text{insert}(k \rightarrow DP[i])$
        **end**
      **end**
    **end**
    `// Use DP entries up to this point to affect later DP entries`
    **foreach** $u, p_1 \in Forward[v]$ **do**
      **foreach** $i \in B[u]$ **do**
        **foreach** $p_2 \in P_2$ **do**
          $k \leftarrow LastToReach_2[\text{end}(M[i].w_2), p_2]$
          $DP[i] \leftarrow \max(DP[i], \text{weight}(M[i]) + Tree[p_1, p_2].\text{rangeMax}((-\infty, k])$
        **end**
      **end**
    **end**
  **end**
  **return** Anchors from traceback starting at $\arg\max DP$

**Algorithm 4:** Compute the heaviest weight co-linear chain with affine gaps

**Input:** $G_1$, $G_2$, $P_1$, $P_2$, and $M$ as in Algorithm 2, gap open penalty $g$ and extend penalty $e$.
**Output:** Co-linear chain of matches that maximizes total weight

```
Function MaxWeightChainAffine(G1, G2, P1, P2, M):
    // Initialize LastToReach1, LastToReach2, Forward, D1, D2, B, and E
    InitializeSparseDP(G1, G2, P1, P2, M)
    // Initialize sparse query data structures
    foreach p1 ∈ P1, p2 ∈ P2 do
        Initialize 3 empty 2-D ORMQs in Tree[p1,p2] named M, I, and D
    end
    // Initialize dynamic programming table
    foreach M[i] ∈ M do
        DP[i] ← weight(M[i])
    end
    // Do sparse dynamic programming
    foreach v ∈ V1 in topological order do
        // Enter matches ending here into sparse query structures
        foreach i ∈ E[v] do
            foreach p1 ∈ P2, k1 such that p1[k1] = v do
                foreach p2 ∈ P2, k2 such that p2[k2] = end(M[i].w2) do
                    Tree[p1,p2].M.insert((k2, k1 − k2) → DP[i])
                    Tree[p1,p2].I.insert((k2, k1 − k2) → DP[i] + e · (k1 − k2))
                    Tree[p1,p2].D.insert((k2, k1 − k2) → DP[i] + e · (k2 − k1))
                end
            end
        end
        // Use DP entries up to this point to affect later DP entries
        foreach u1, p1 ∈ Forward[v] do
            foreach i ∈ B[u1] do
                w ← weight(M[i])
                u2 ← begin(M[i].w2)
                foreach p2 ∈ P2, k2 such that p2[k2] = u2 do
                    q ← LastToReach1[u1,p1] − LastToReach2[u2,p2] + D1[u1,p1] − D2[u2,p2]
                    DP[i] ← max(DP[i], w + Tree[p1,p2].M.rangeMax((−∞, k2], [q, q]))
                    DP[i] ←
                      max(DP[i], w − g − e · q + Tree[p1,p2].I.rangeMax((−∞, k2], (−∞, q − 1]))
                    DP[i] ← max(DP[i], w − g + e · q + Tree[p1,p2].D.rangeMax((−∞, k2], [q + 1, ∞))

                end
            end
        end
    end
    return Anchors from traceback starting at arg max DP
```

### 3 Identifying alignable regions in an anchor chain

We formalize the process of identifying unalignable regions from an anchor chain through a set of problems that we call optimal partition problems. They take as input two arrays of equal length, $W = W_1 W_2 \dots W_N$ and $L = L_1 L_2 \dots L_N$, which are interpreted as weights and lengths, respectively. In our use case, the entries in these arrays refer to anchors in the chain and the gaps between them. The weights in $W$ are the corresponding anchor's score or 0 for a gap between anchors. The $L$ values are the length of the anchor or, for a gap, the shortest path through through the gap in the graph. The output of optimal partition problems is a set of ordered, nonoverlapping intervals $[i_1, j_1], [i_2, j_2], \dots [i_m, j_m]$ contained in $[1, n]$ that maximize an objective function subject to some constraints. We will present three variants of optimal partition with increasing complexity, building up to the final variant that is used in Centrolign.

#### 3.1 Maximum weight partition

In this variant, the goal is to produce a set intervals with maximum total weight with a cost $c$ for each interval. We solve this problem with a dynamic programming formulation that includes two tables $I$ and $X$. $I_i$ and $X_i$ contains the optimal value for a length $i$ prefix of the input if the $i$-th entry is included or excluded, respectively. The base cases are given by

$$\begin{aligned} X_0 &= 0 \\ I_0 &= 0, \end{aligned} \tag{6}$$

and the dynamic programming recursions are

$$\begin{aligned} X_i &= \max \{X_{i-1}, I_{i-1}\} \\ I_i &= \max_{0 \le j < i} \left( X_j - c + \sum_{k=j+1}^{i} W_k \right). \end{aligned} \tag{7}$$

In the recursion for $I$, the sum indicates the total weight of the final interval in the partition. Every new interval must be preceded by a gap. Thus, the $X_j$ term is sufficient to summarize the contribution of the prefix of the solution that precedes $j + 1$.

This recursion yields an obvious $O(N^2)$ dynamic programming implementation, but this can be improved to $O(N)$ with some additional observations. First, the sums over $W$ can each be computed in $O(1)$ time by preprocessing $W$ into a prefix sum array $P$ in $O(N)$ time. The recursion can be stated

$$I_i = \max_{0 \le j < i} \left(X_j - c + P_i - P_j\right). \tag{8}$$

Next, observe that this maximum always occurs at the $j < i$ that maximizes $X_j - P_j$, since the other terms do not depend on $j$. Accordingly, the algorithm can simply maintain a pointer to this $j$ and update it over the course of iteration. With this modification, it is no longer necessary to explicitly compute the maximum, which reduces the overall complexity to $O(N)$. Pseudocode is provided in Algorithm 5.

#### 3.2 Average constrained maximum weight partition

In this next variant of optimal partitioning, we add a constraint that the average weight of each interval is greater than some parameter $m$. By average, we mean that is averaged over the lengths in $L$, so that in every output interval $[i, j]$ we have $(W_i + \ldots + W_j)/(L_i + \ldots + L_j) \ge m$. Nearly the same recursions apply to this problem, except that this constraint is added to the max term.

$$I_i = \max_{\substack{0 \le j < i \\ \frac{W_{j+1} + \cdots + W_i}{L_{j+1} + \cdots + L_i} \ge m}} \left(X_j - c + P_i - P_j\right). \tag{9}$$

To help evaluate this constraint efficiently, we borrow a technique from (8). We define an auxiliary array $F$ where $F_i = W_i - mL_i$, and we construct the prefix sum array of $F$, which we name $A$. Simple algebraic manipulations show that an interval $[i, j]$ satisfies the average weight constraint if and only if $A_j \ge A_i$. Accordingly, we can rearrange the recursion for $I$ as

$$I_i = -c + P_i + \max_{\substack{0 \le j < i \\ A_{j+1} \le A_i}} \left(X_j - P_j\right). \tag{10}$$

This max operation can performed with an RMQ over key-value pairs, where the keys are $A_i$ and the values are $X_i - P_i$. We use the same augmented balanced binary search tree as in Algorithm 3 to provide this interface. Each dynamic programming iteration performs one `insert` and one `rangeMax` operation, which each take $O(\log N)$ time. Thus, the total run time for this algorithm is $O(N \log N)$. Pseudocode is provided in Algorithm 6.

##### 3.3 Window-average constrained maximum weight partition

This variant of optimal partitioning is similar to the previous, except that the constraint applies to a windowed average rather than an average over the full interval. Specifically, it takes a window length parameter $d$, and every $d$-length subsegment of an interval must have average greater than $m$. If we conceptualize $L$ to be a series of entries with length $L_i$ laid end-to-end, then the subsegment of the window may be positioned freely over these entries (Figure 2). In particular, it is possible that it only partially overlaps one of the entries. In this case, we consider the weight of that entry to scale proportionately to the length of the overlap for the purposes of computing the windowed average. That is, if a window overlaps entry $i$ by some length $\ell < L_i$, then this entry contributes $W_i\ell/L_i$ to the numerator of the average and $\ell$ to the denominator.

In general, there are infinitely many windows of length $d$ that fit inside an interval of entries in $L$. However, it is not necessary to check all possible windows to verify that an interval satisfies the windowed average constraint. It is straightforward to verify that an interval $[i, j]$ with $\sum_{k=i}^{j} L_k \geq d$ satisfied the constraint if and only if the constraint is satisfied for windows that are wholly contained within $[i, j]$ and either 1) begin at the beginning of an entry in $L$, or 2) end at the end of an entry in $L$. We refer to these cases as *left-adjusted* and *right-adjusted*, respectively (Figure 2). If $\sum_{k=i}^{j} L_k < d$, then the windowed average for the interval is equal to its full, un-windowed average.

Our algorithm exploits these facts to efficiently check the windowed average condition. For intervals of total length less than $d$, we use the same technique as in Algorithm 6. We construct the $A$ array and an RMQ. Over the course of iteration, entries in the RMQ are erased once their distance exceeds $d$.

For intervals of length at least $d$, we first construct arrays $V^l$ and $V^r$ that contain 1 in the $i$-th position if average constraint is violated by the left- or right-adjusted windows of the $i$-th entry, respectively, or else they contain 0. These arrays can be constructed in $O(N)$ time by a sweep over the input array in the forward or reverse direction. We construct prefix sum arrays for $V^l$ and $V^r$, which we denote $C^l$ and $C^r$. Let $i'$ be the lowest index such the right-adjusted window of $L_{i'}$ is contained in $[i, j]$, and let $j'$ be the maximum such index for a left-adjusted window. We can then check that interval $[i, j]$ of length at least $d$ satisfies the windowed average constraint by checking that $C^l_i = C^l_{j'}$ and $C^r_{i'} = C^r_j$. We store this $i'$ in an array entries $T^r_j$ and likewise $j'$ in $T^l_i$ (Figure 2). These values can be precomputed in a linear sweep over $L$.

We also note that, as in Algorithm 5, among $j$ that satisfy these conditions, the maximum will occur at the $j$ that maximizes $X_j - P_j$. Accordingly, we can use the same technique as in Algorithm 6 of maintaining a pointer to this $j$ and updating it during iteration. However, there are a few additional complications. First, since we are restricting to $j$ such that $\sum_{k=j}^{i} L_k \geq d$, we only consider $j$ as a candidate for this pointer when iteration reaches a length at least $d$ away from $j$. Next, if the minimum windowed average constraint is violated, it is possible that no feasible $j$ exists. Thus, we set $j$ to *Null* when it is interrupted by a window that fails the minimum average constraint.

The time required to construct all of the arrays involved in this algorithm is $O(N)$. Checking whether a candidate $j$ is feasible and maximizes $X_j - P_j$ takes $O(1)$ time for each entry for a total of $O(N)$ time. Each entry also generates three RMQ operations that requires $O(\log N)$ time each. Thus, the RMQ queries dominate the run time with a total of $O(N \log N)$ for the entire algorithm. Pseudocode is presented in Algorithm 7.

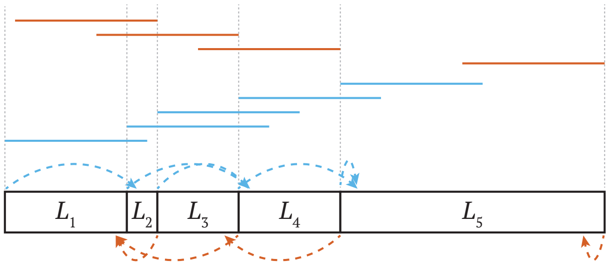


Figure 2: **Window-average constrained partition**. The fixed-length windows (solid lines) are arranged as if intervals whose lengths are given by $L$ are laid end to end. Left-adjusted windows are shown in blue and right-adjusted windows in orange. The values in $T^l$ (blue dashed lines) and $T^r$ (orange dashed lines) point to the interval that contains the end of the left- and right-adjusted windows, respectively.

**Algorithm 5:** Maximum weight partition into intervals

**Input:** Array of weights $W$ and interval cost $c$
**Output:** A partition of intervals that maximizes total weight with a cost applied to each interval

**Function** `MaxWeightPartition`($W$, $c$):
    $P \leftarrow$prefix sum array of $W$
    $I_0 \leftarrow 0$
    $X_0 \leftarrow 0$
    $j \leftarrow 0$
    **foreach** $i = 1, 2, \ldots, N$ **do**
        $X_i \leftarrow \max\{X_{i-1}, I_{i-1}\}$
        $I_i \leftarrow X_j + P_i - P_j - c$
        **if** $X_i - P_i > X_j - P_j$ **then**
            $j \leftarrow i$
        **end**
    **end**
    **return** Intervals from traceback starting at $\arg\max I$

**Algorithm 6:** Maximum weight partition into intervals subject to a constraint on average weight

**Input:** Array of weights $W$, array of lengths $L$, minimum average weight $m$, and interval cost $c$
**Output:** A partition of intervals that maximizes total weight with a cost applied to each interval and with each interval having average value above a minimum threshold

**Function** `AvgConstrainedMaxWeightPartition`($W$, $L$, $m$, $c$):
    $P \leftarrow$ prefix sum array of $W$
    $A \leftarrow$ prefix sum array of $W - m \cdot L$
    $R \leftarrow$ empty RMQ
    $I_0 \leftarrow 0$
    $X_0 \leftarrow 0$
    **foreach** $i = 1, 2, \ldots, N$ **do**
        $X_i \leftarrow \max\{X_{i-1}, I_{i-1}\}$
        $I_i \leftarrow -c + P_i + R.\text{rangeMax}((-\infty, A_i])$
        $R.\text{insert}(A_i \rightarrow X_i - P_i)$
    **end**
    **return** Intervals from traceback starting at $\arg\max I$

**Algorithm 7:** Maximum weight partition into intervals subject to a constraint on average weight

**Input:** Array of weights $W$, array of lengths $L$, window length $d$, minimum windowed-average weight $m$, and interval cost $c$

**Output:** A partition of intervals that maximizes total weight with a cost applied to each interval and with each interval having average value above a minimum threshold for all windows contained in the interval

**Function** `WindowAvgConstrainedMaxWeightPartition(`$W$, $L$, $d$, $m$, $c$`):`
  `// Scan the array to assess the window constraint on left-adjusted windows`
  $ww \leftarrow 0, wl \leftarrow 0$, j = 1
  **foreach** $i = 1, 2, \ldots, N$ **do**
    **while** $j \leq N$ and $wl \leq d$ **do**
      $ww \leftarrow ww + W_j$, $wl \leftarrow wl + L_j$, $j \leftarrow j + 1$
    **end**
    $T_i^l \leftarrow j - 1$
    **if** $j \leq N$ **then**
      $V_i^l \leftarrow 1$ if $ww - W_j + (d - wl + L_j)W_j/L_j \geq d \cdot m$ else 0
    **else**
      $V_i^l \leftarrow V_{i-1}^l$
    **end**
    $ww \leftarrow ww - W_i$, $wl \leftarrow wl - L_i$
  **end**
  Repeat the above loop in reverse to construct $V^r$ and $T^r$
  `// Arrays of left- and right-adjusted window constraint satisfaction`
  $C^l, C^r \leftarrow$ prefix sum arrays of $V^l, V^r$
  $P \leftarrow$ prefix sum array of $W$
  $A \leftarrow$ prefix sum array of $W - m \cdot L$
  $R \leftarrow$ empty RMQ
  $j \leftarrow Null$, $i' \leftarrow 0$ `// arg max of` $X_j - P_j$ `and index of full window closest to` $i$
  $wl \leftarrow 0$, $wi \leftarrow 0$ `// length and index of window adjacent to` $i$
  $I_0 \leftarrow 0$, $X_0 \leftarrow 0$
  **foreach** $i = 1, 2, \ldots, N$ **do**
    $X_i \leftarrow \max\{X_{i-1}, I_{i-1}\}$
    `// Determine bound of left-adjusted windows inside interval ending at` $i$
    **while** $T_{i'+1}^l \leq i$ **do**
      $i' \leftarrow i' + 1$
    **end**
    `// Check if interval starting at` $j$ `is now interrupted by infeasible window`
    **if** $j \neq Null$ and $(C_j^l \neq C_{i'}^l$ or $C_{T_j^r}^r \neq C_i^r)$ **then**
      $j \leftarrow Null$;
    **end**
    $wl \leftarrow wl + L_i$
    **while** $wl - L_{wi} \geq d$ **do**
      `// Entry` $wi$ `has fallen out of the window adjacent to` $i$
      **if** $C_{wi}^l = C_{i'}^l$ and $C_{T_{wi}^r}^r = C_i^r$ and $(j = Null$ or $X_{wi} - P_{wi} > X_j - P_j)$ **then**
        $j = wi$
      **end**
      $R$.erase$(A_{wi} \to X_{wi} - P_{wi})$
      $wl \leftarrow wl - L_{wi}$, $wi \leftarrow wi + 1$
    **end**
    `// Find max of intervals longer than` $d$ `and shorter than` $d$
    $I_i \leftarrow -c + P_i + \max\{X_j - P_j, R.\text{rangeMax}((-\infty, A_i])\}$ `// or` $-\infty$ `if neither exist`
    $R$.insert$(A_i \to X_i - P_i)$
  **end**
  **return** Intervals from traceback starting at $\arg\max I$

### 4 Graph-graph wavefront alignment

Our graph-graph wavefront alignment (GGWFA) algorithm generalizes the graph wavefront alignment (GWFA) algorithm(9), which aligns sequences to a graph. GGWFA requires that at least one of the graphs being aligned is acyclic (in Centrolign, both are acyclic). Like the prior wavefront alignment (WFA) algorithm(10; 11), GGWFA requires alignment parameters in which the match score is 0. Accordingly, we convert the conventional alignment parameters used for POPOA using the formulas of (12).

Unlike the original WFA algorithm, the run time for GGWFA is not linear when parameterized by the score of the alignment. As noted by (9), this parameterized run time is not possible under the Strong Exponential Time Hypothesis (SETH). This follows from the previous result that exact matching between a sequence and an acyclic sequence graph (which corresponds to an alignment with score 0) is not possible in strongly subquadratic time under SETH(13).

GGWFA essentially consists of a method to compute the interior of the dynamic programming matrix for POPOA in order of increasing score. The alignment completes when it reaches the final position for the first time, which necessarily occurs with the minimum possible score. This is accomplished through use of a bucket queue containing dynamic programming entries, bucketed by score.

The core iteration is presented in pseudocode in Algorithm 8. To begin, the lowest scoring dynamic programming entry is popped from the bucket queue. Next, the dynamic programming entries that depend on the popped entry are added to the queue with the score that results from extending from this entry. Algorithm 9 shows the outer flow that directs this core iteration. This process is equivalent to Dijkstra's algorithm through a graph whose nodes are dynamic programming entries and whose edges are dependencies between entries. The edge lengths are the score parameter associated to the dependency's edit operation. Thus, the length of the path is the sum of the edit penalties, which is the alignment score. Accordingly, the correctness of GGWFA follows from the correctness of Dijkstra's algorithm.

Since the filter at the beginning of Algorithm 8 prevents it from executing twice for the same dynamic programming entry, the main body is executed once for each of the $O(|V_1||V_2|)$ entries. In an execution for $v_1 \in V_1$ and $v_2 \in V_2$, there are $O(\deg(v_1) \cdot \deg(v_2))$ queuing operations. Thus, the total execution time is $O(\sum_{1\in V_1, v_2\in V_2} \deg(v_1) \cdot \deg(v_2)) = O(|E_1||E_2|)$. Note that the number of queueing operations also bounds the number of calls to Algorithm 8 that fail the initial filter. Finally, we assume that $x + g + e = O(1)$, in which case it is only necessary to check $O(1)$ buckets in the bucket queue before finding a nonempty one, which makes each queue operation $O(1)$ time. Thus, the total run time is $O(|E_1||E_2|)$.

#### 4.1 A long deletion variant of the graph-graph wavefront algorithm

When one of the graphs supplied to GGWFA is much longer than the other, it is parsimonious to assume that their alignment contains a single long deletion. We have developed an additional variant of GGWFA that uses this assumption to dramatically reduce the run time at the cost of guaranteed optimality. The core strategy is to execute GGWFA simultaneously in the forward and reverse directions until they meet on the short graph. A long deletion can then be inferred for the long graph.

This algorithm needs to efficiently identify whether two nodes in the long graph can reach each other, which is necessary to determine if they can be connected by a deletion. To choose the best position for the deletion, it is also helpful to be able to efficiently determine the distance between these nodes. We use a previously published distance oracle based on superbubble graph motifs to provide these queries(14). To initially compute the superbubbles, we use Algorithm 2 of (15).

**Algorithm 8:** Compute optimal alignment of paths between two graphs

**Input:** Acyclic sequence graphs $G_1$ and $G_2$, bucket queue $Q$, score at top of queue $s$, direction of iteration $d$, mismatch penalty $x$, gap open penalty $g$, and gap extend penalty $e$

**Output:** Node pair from $V_1 \times V_2$ that was removed from $Q$ in this iteration

```
Function GGWFAIteration(G_1, G_2, Q, s, d, x, g, e):
    u_1, u_2, c ← Q.popNext()
    if u_1, u_2, c has already been popped in an earlier call then
        return Null
    end
    if d = forward then
        Next_1 ← G_1.successors(u_1)
        Next_2 ← G_2.successors(u_2)
    else
        Next_1 ← G_1.predecessors(u_1)
        Next_2 ← G_2.predecessors(u_2)
    end
    if c = M then
        // Match or mismatch
        foreach v_1 ∈ Next_1, v_2 ∈ Next_2 do
            if v_1.label() = v_2.label() then
                Q[s].insert((v_1, v_2, M))
            else
                Q[s + x].insert((v_1, v_2, M))
            end
        end
        // Gap open
        foreach v_1 ∈ Next_1 do
            Q[s + g + e].insert((v_1, u_2, I))
        end
        foreach v_2 ∈ Next_2 do
            Q[s + g + e].insert((u_1, v_2, D))
        end
    else
        // Gap close
        Q[s].insert((u_1, u_2, M))
        // Gap extend
        if c = I then
            foreach v_1 ∈ Next_1 do
                Q[s + e].insert((v_1, u_2, I))
            end
        else
            foreach v_2 ∈ Next_2 do
                Q[s + e].insert((u_1, v_2, D))
            end
        end
    end
    return u_1, u_2
```

**Algorithm 9:** Compute optimal alignment of paths between two graphs

**Input:** Acyclic sequence graphs $G_1$ and $G_2$, alignment source $(s_1, s_2) \in V_1 \times V_2$, alignment sink $(t_1, t_2) \in V_1 \times V_2$, mismatch penalty $x$, gap open penalty $g$, and gap extend penalty $e$

**Output:** The optimal alignment of a paths in $G_1$ and $G_2$ that connect $s_1$ to $t_1$ and $s_2$ to $t_2$

**Function** `GGWFA`($G_1$, $G_2$, $s_1$, $s_2$, $t_1$, $t_2$, $x$, $g$, $e$):
  $Q \leftarrow$ empty bucket queue
  $Q[0]$.insert$((s_1, s_2, M))$
  $s \leftarrow 0$
  **while** $Q[s]$.next() $\neq (t_1, t_2, M)$ **do**
    `GGWFAIteration` ($G_1$, $G_2$, $Q$, $s$, forward, $x$, $g$, $e$)
    Increase $s$ to the index of lowest non-empty bucket in $Q$
  **end**
  **return** Alignment from traceback

**Algorithm 10:** Compute an alignment between paths from two graphs with a single large deletion

**Input:** Acyclic sequence graphs $G_1$ (long) and $G_2$ (short), alignment source $(s_1, s_2) \in V_1 \times V_2$, alignment sink $(t_1, t_2) \in V_1 \times V_2$, mismatch penalty $x$, gap open penalty $g$, and gap extend penalty $e$

**Output:** An alignment of paths in $G_1$ and $G_2$ that connect $s_1$ to $t_1$ and $s_2$ to $t_2$, containing one large deletion along $G_1$

**Function** `DeletionGGWFA(`$G_1$, $G_2$, $s_1$, $s_2$, $t_1$, $t_2$, $x$, $g$, $e$`):`
  `// Initialize forward and reverse queues`
  $Q_f, Q_r \leftarrow$ empty bucket queues
  $Q_f[0]$.insert$((s_1, s_2, M))$
  $Q_r[0]$.insert$((t_1, t_2, M))$
  `// Maps to record` $G_1$ `nodes aligned to` $G_2$ `nodes and their scores`
  $L_f, L_r \leftarrow$ empty multimaps
  $S_f, S_r \leftarrow$ empty maps
  $D \leftarrow$ superbubble distance oracle for $G_1$
  $s_f, s_r \leftarrow 0, 0$ `// Forward and reverse scores`
  $s_{end} \leftarrow \infty$
  $p \leftarrow \max\{x, o + e\}$ `// Amount that could need to search past first meet score`
  **while** $s_f + s_r < s_{end}$ **do**
    **if** $s_f \leq s_r$ **then**
      `// Iterate in forward direction`
      $u_1, u_2 \leftarrow$ `GGWFAIteration` ($G_1$, $G_2$, $Q_f$, $s_f$, forward, $x$, $g$, $e$)
      **if** $s_{end} = \infty$ and there exists $v_1 \in L_r[u_2]$ such that $D$.distance$(u_1, v_1) < \infty$ **then**
        `// Found a location where a long deletion can be inferred`
        $s_{end} \leftarrow s_f + s_r + p$
      **end**
      $L_f$.insert$(u_2 \rightarrow u_1)$
      $S_f$.insert$(u_1, u_2 \rightarrow s_f)$
      Increase $s_f$ to the index of lowest non-empty bucket in $Q_f$
    **else**
      `// Iterate in reverse direction`
      $u_1, u_2 \leftarrow$ `GGWFAIteration` ($G_1$, $G_2$, $Q_r$, $s_r$, reverse, $x$, $g$, $e$)
      **if** $s_{end} = \infty$ and there exists $v_1 \in L_f[u_2]$ such that $D$.distance$(v_1, u_1) < \infty$ **then**
        `// Found a location where a long deletion can be inferred`
        $s_{end} \leftarrow s_f + s_r + p$
      **end**
      $L_r$.insert$(u_2 \rightarrow u_1)$
      $S_r$.insert$(u_1, u_2 \rightarrow s_r)$
      Increase $s_r$ to the index of lowest non-empty bucket in $Q_r$
    **end**
  **end**
  Iterate $L_f$ and $L_f$ to find the nodes $u_1, v_1 \in V_1$ and $u_2 \in V_2$ that minimize $S_f[u_1, u_2] + S_r[v_1, u_2] + g + e \cdot D$.distance$(u_1, v_1)$
  **return** Alignment from traceback

### 5 Inducing cycles at tandem duplications

We identify tandem duplications as segments of anchor chains between a sequence and itself in which matches between the exact same position have been removed as options. The tandem duplications are identified by having high score relative to the optimal chain that aligns the sequence trivially to itself. We also require that the segment be separated from the main diagonal by roughly its own length. This is a requirement for the repeat to be in tandem. We allow a small deviation from this criterion that scales with the square root of of this length, which is motivated by the scaling behavior of random walks.

We formulate this problem as an optimal partition problem, and we apply the same dynamic programming technique as in Section 3. However, unlike the algorithms in Section 3, we are not aware of a sparse formulation to compute the maximum, and therefore we use the trivial quadratic implementation.

The objective function of this dynamic programming algorithm is the total length of the duplications. We also include a cost for each interval in the objective function, which additionally functions as the minimum length for any chosen interval. The restrictions on score and position are treated as constraints on feasibility. Pseudocode is provided in Algorithm 11.

#### 5.1 Identifying inconsistent representations of variation

Centrolign employs two heuristics to identify regions of the graph where variation has been inconsistently represented in tandem duplication alignments, both based on the cactus graph of the pangenonme graph(16). The first uses the cactus graph to induce a snarl decomposition, represented in a snarl tree(17). Short cycles are identified directly using Kahn's algorithm on net graphs of the snarls. The length of the cycle is considered to be the maximum length of any subsequence of the input sequences that traverse the snarl. The second looks for short intervals of a chain in which the same sequence traverses the interval multiple times, taking partially disjoint nodes each time. This is a characteristic pattern of an indel being positioned differently across different tandem duplication alignments.

#### 5.2 Creating a guide tree for realignment

Algorithm 13 presents the method that Centrolign uses to create a guide tree for the realignment problems. The strategy is to retain the topology of the guide tree from the full problem, but to duplicate subtrees to accommodate multiple traversals of the subgraph by the same sample's sequence. First, the algorithm identifies subtrees in which every sample has the same number of traversals using an upward pass over the tree. Next, a downward pass over the tree finds the highest nodes whose subtrees have a consistent number of traversals and duplicates them at that point. When the subtrees are duplicated, the $n$-th duplicate receives the $n$-th subsequence from each of the samples.

**Algorithm 11:** Identify tandem duplications in a chain of matches from a sequence to itself

**Input:** A colinear chain $C = \{(x_i, y_i, \ell_i)\}_{i=1}^{N}$ of self-matches in a sequence, which begin at position $x_i$ and $y_i$ and have length $\ell_i$, and three parameters: $\alpha$, which controls the scaling of the lenience in the separation condition, $\beta$, which controls the lenience of the score requirement, and $c$, which is the cost of an interval

**Output:** A set of intervals that correspond to tandem duplications

**Function** `FindTandemDuplications`($C$, $\alpha$, $\beta$, $c$):
- $I_0 \leftarrow 0$
- $X_0 \leftarrow 0$
- **foreach** $i = 1, 2, \ldots, N$ **do**
  - $X_i \leftarrow \max\{X_{i-1}, I_{i-1}\}$
  - $s \leftarrow 0$
  - $\ell \leftarrow 0$
  - **foreach** $j = i, i-1, \ldots, 1$ **do**
    - $s \leftarrow s + \text{score}(C[j])$
    - $\ell \leftarrow \ell + C[j].\ell$
    - **if** $j \neq i$ **then** $\ell \leftarrow \ell + \min\{C[j+1].x - C[j].x, C[j+1].y - C[j].y\} - C[j].\ell$
    - **if** $|C[j].x - C[j].y| > \ell - \alpha\sqrt{\ell}$
      and $\beta \cdot s >$ the corresponding score on the trivial chain
      and $X_{j-1} + \ell - c > I_i$ **then**
      - $I_i \leftarrow X_{j-1} + \ell - c$
    - **end**
  - **end**
- **end**
- **return** Intervals from traceback starting at $\arg\max I$

**Algorithm 12:** Identify partially disjoint multiple traversals of chains, which can indicate inconsistently represented indels

**Input:** Graph $G$, snarl tree $S$, window size $w$, minimum disjoint length $d$
**Output:** List of pairs of boundary nodes for snarls that contain short cycles

**Function** `FindInconsistentIndels(`$G$, $S$, $w$, $d$`):`
    $Q \leftarrow$ queue containing the root chain of $S$
    **while** $Q$ is not empty **do**
        Pop a chain from $Q$
        **foreach** Window of length $w$ of snarls in the chain **do**
            **foreach** Sequence path $p$ in $G$ that traverses $c$ multiple times **do**
                **foreach** Pair of traversals $(t_1, t_2)$ of $p$ through the window **do**
                    $\ell_1 \leftarrow$ length of sequence in $t_1$ that doesn't overlap $t_2$
                    $\ell_2 \leftarrow$ length of sequence in $t_2$ that doesn't overlap $t_1$
                    `// Sequence that overlaps neither traversal can indicate a partially overlapping deletion`
                    $\ell_. \leftarrow$ length of sequence in the window that overlaps neither $t_1$ nor $t_2$
                    **if** $\ell_1 + \ell_. > d$ and $\ell_2 + \ell_. > d$ **then**
                        Record an inconsistency in this window
                **end**
            **end**
        **end**
        Add child chains to $Q$ for all snarls that were not in an inconsistent window
    **end**
    **return** Boundary nodes for merged windows

**Algorithm 13:** Identify partially disjoint multiple traversals of chains, which can indicate inconsistently represented indels

```
Input: Guide tree for full MSA T, subsequences for realignment S
Output: A guide tree for a realignment problem in the
Function MakeRealignmentGuideTree(T, S):
    P ← samples whose sequences are represented in S
    T′ ← T, pruned to include only ancestors of P up through their lowest common ancestor,
      with all degree-2 nodes except possibly the root compacted into single edges
    // Identify subtrees with consistent copy count
    C ← empty map
    foreach n ∈ T′ in postorder do
        if n is a leaf then
            C[n] ← the number of sequences in S from n.sample()
        else if C[c] has the same values for all c ∈ n.children() then
            C[n] ← C[c], where c ∈ n.children()
        else
            C[n] ← Inconsistent
        end
    end
    T″ ← empty tree
    // Q will contain records of (Node from T′, Action, Parent in T″)
    Q ← empty queue
    Push (T′.root(), Traverse, Null) onto Q
    // Traverse downward, duplicating subtrees with consistent copy count
    while Q is not empty do
        n, a, p ← Q.pop()
        u ← new node in T″ with parent p
        if a = Traverse and C[n] ≠ Inconsistent then
            foreach i = 1, 2, ..., C[n] do
                // This subtree will receive the i-th subsequence from each sample
                   among its leaves
                Push (n, Copy, u) onto Q
            end
        else
            foreach c ∈ n.children() do
                Push (c, a, u) onto Q;
            end
        end
    end
    return T″
```
