## Supplementary Materials for "Pangenome alignment reveals global diversity and evolution of human centromeric regions"

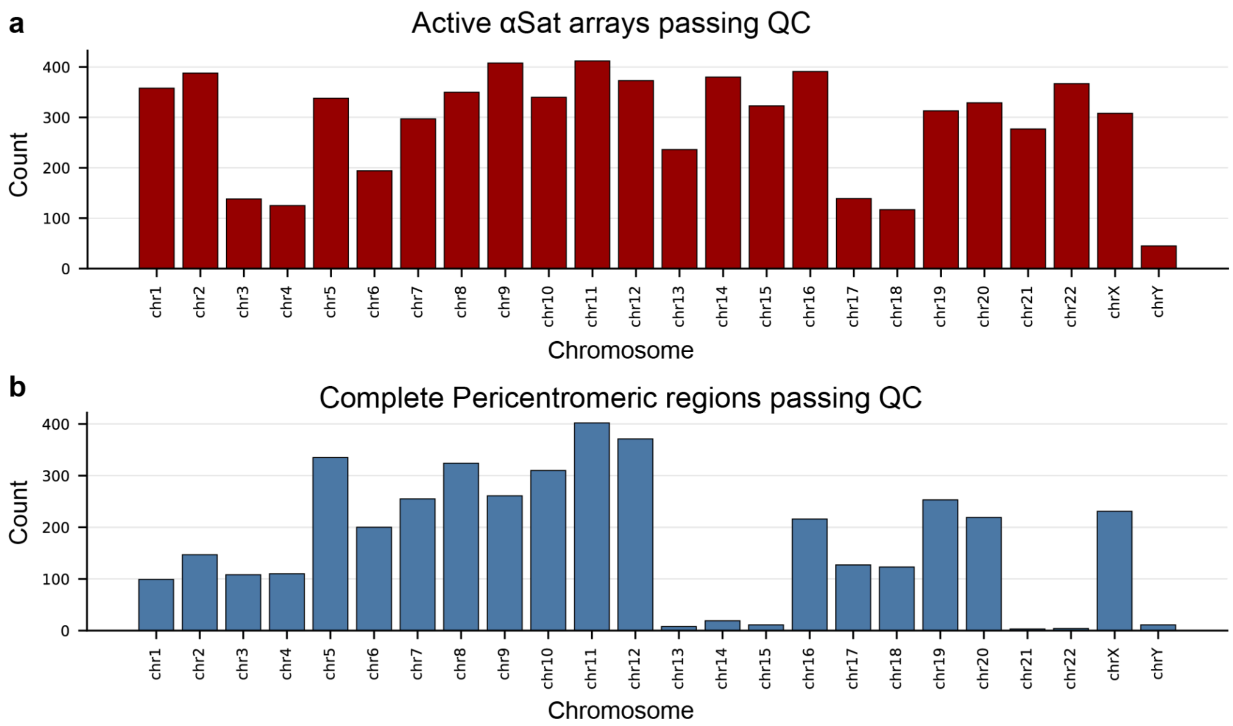


**Supplementary Figure 1. Chromosome-level count of active arrays and pericentromeric regions passing QC**

**a**, Count of active arrays passing quality control (QC). **b**, Count of complete pericentromeric regions passing QC for each chromosome. QC required the entire active array (**a**) or pericentromeric region (**b**) to be assembled in one sequence without gaps or predicted errors (Methods). Autosomes have a theoretical maximum of 460 haplotypes and chromosomes X and Y have a theoretical maximum of 345 and 115, respectively. Fewer complete pericentromeric regions passed QC in acrocentric chromosomes and chromosome Y due to their large, and complex, pericentromeres. Chromosome Y is only expected to be present in one haplotype in male samples.

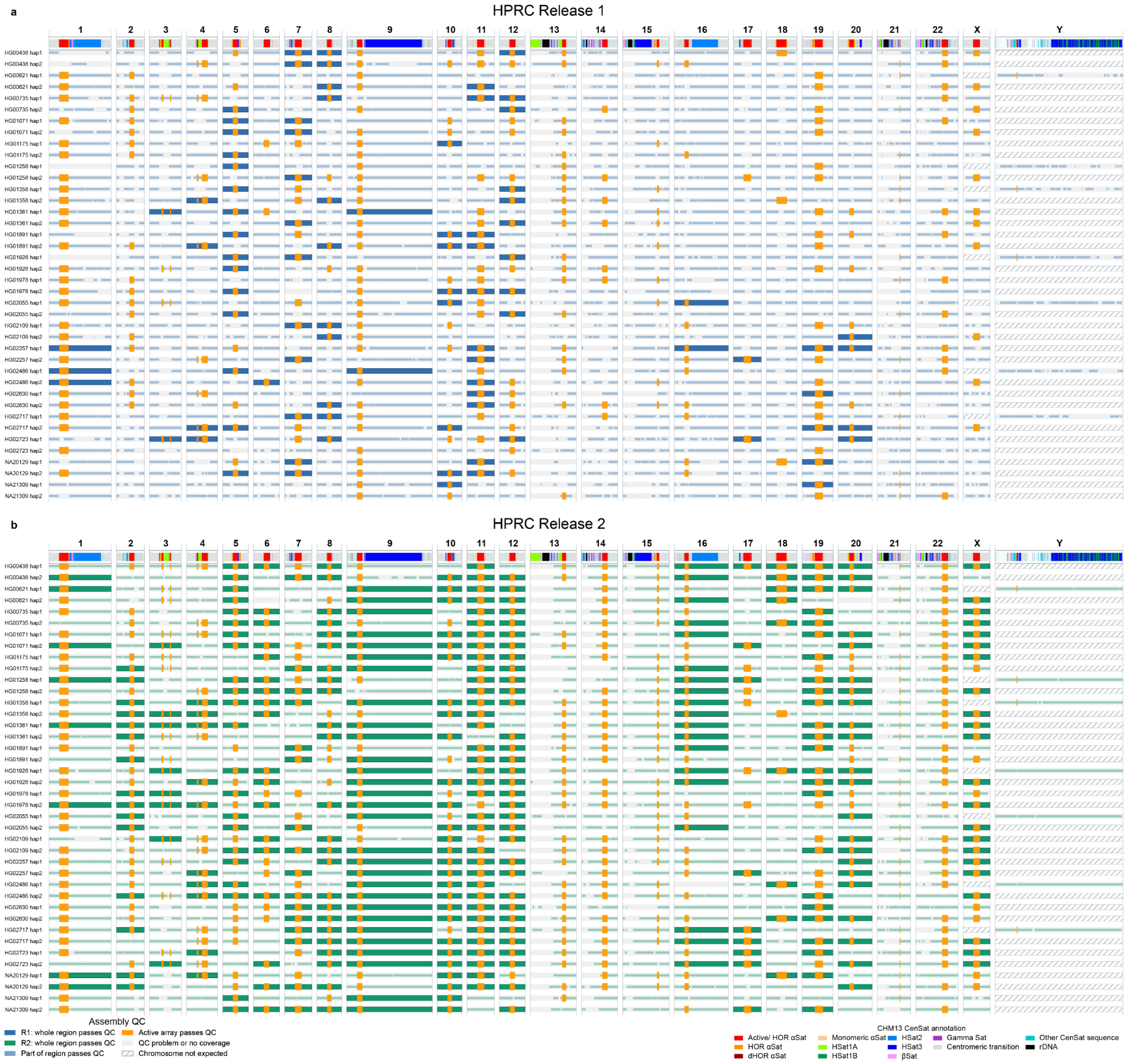


**Supplementary Figure 2. Improved assembly of peri/centromeres from HPRC Release 1 to Release 2.**

Shown are 40 haplotype assemblies (20 samples) present in both Release 1 and Release 2 of the HPRC. Rows are the assemblies aligned to CHM13 (methods). CenSat annotations for CHM13 are shown at the top. Dark blue (R1) and green (R2) indicate pericentromeric regions where the assembly's alignment spans the entirety of CHM13's CenSat annotations without predicted QC errors. Lighter colors indicate assembled regions without predicted QC errors, but without complete coverage. Orange indicates an active array where the assembly's alignment spans the entirety of CHM13's active array without predicted QC errors. Pale gray indicates QC-problem or no-coverage sequence.

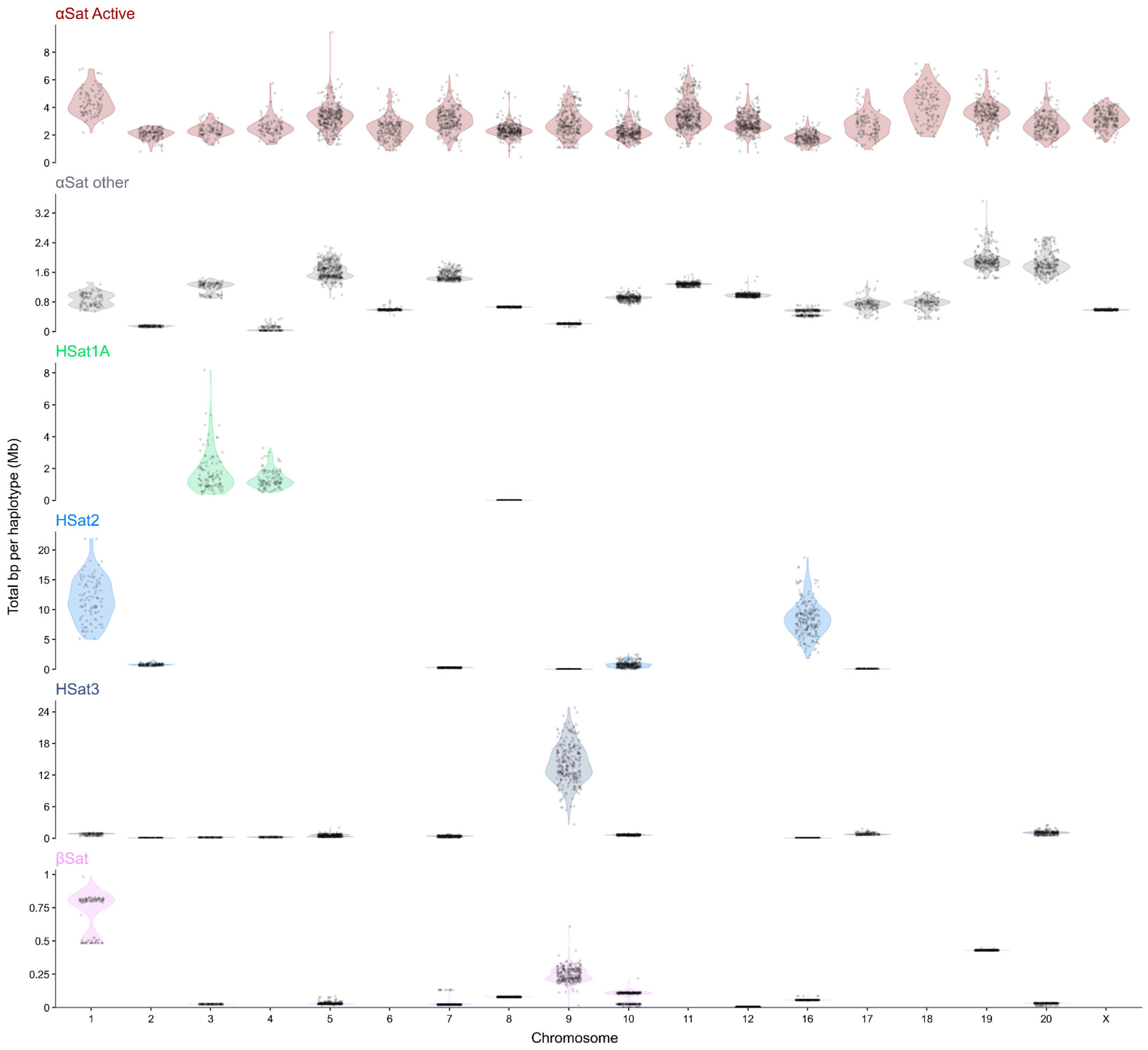


**Supplementary Figure 3. Satellite Load Across Chromosomes**

Points show the total assembled length of each satellite class per chromosome, per haplotype. Violins are kernel density estimates of the array totals across haplotypes, trimmed to the observed data range and scaled to a common maximum width. Satellite assignments are from CenSat annotations. Only haplotypes that passed QC for pericentromeric regions in that chromosome are shown. Acrocentrics and chrY were omitted due to the low number of entire pericentromeres assembled. Panels show active α-satellite arrays (αSat Active), other α-satellite sequence (αSat other), HSat1A, HSat2, HSat3, and βSat. ASat other combines HOR, divergent HOR, and monomeric α-satellite sequence. Values are shown in megabases (Mb).

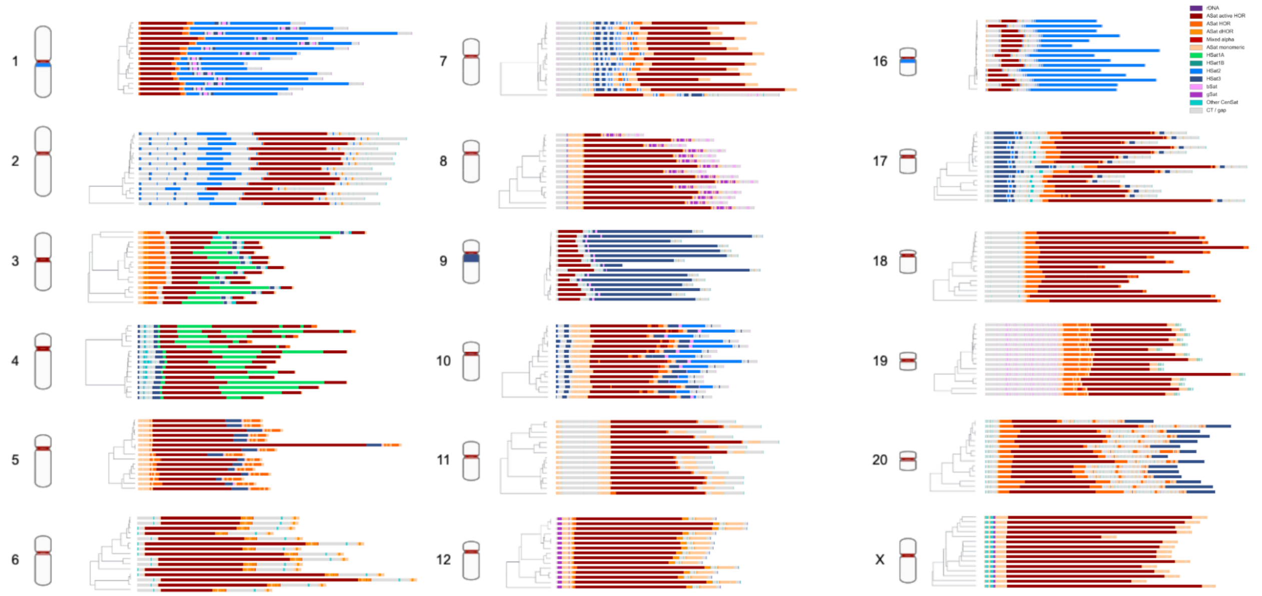


**Supplementary Figure 4. Arrays In HPRC2 show pericentromeric variability**
Chromosome cartoons (left) showing CenSat annotations from CHM13 are next to select haplotypes (right) organized by cenhaps. Chromosome cartoons are schematic: centromeric constrictions are drawn at the largest annotated active array and CenSat annotation band thickness is enlarged for visibility. Chromosomes shown are included in structural analysis of pericentromeric regions; acrocentric and chromosome Y are omitted. For (randomly) selected haplotypes, CenSat annotations (horizontal tracks) are shown alongside cenhap trees.

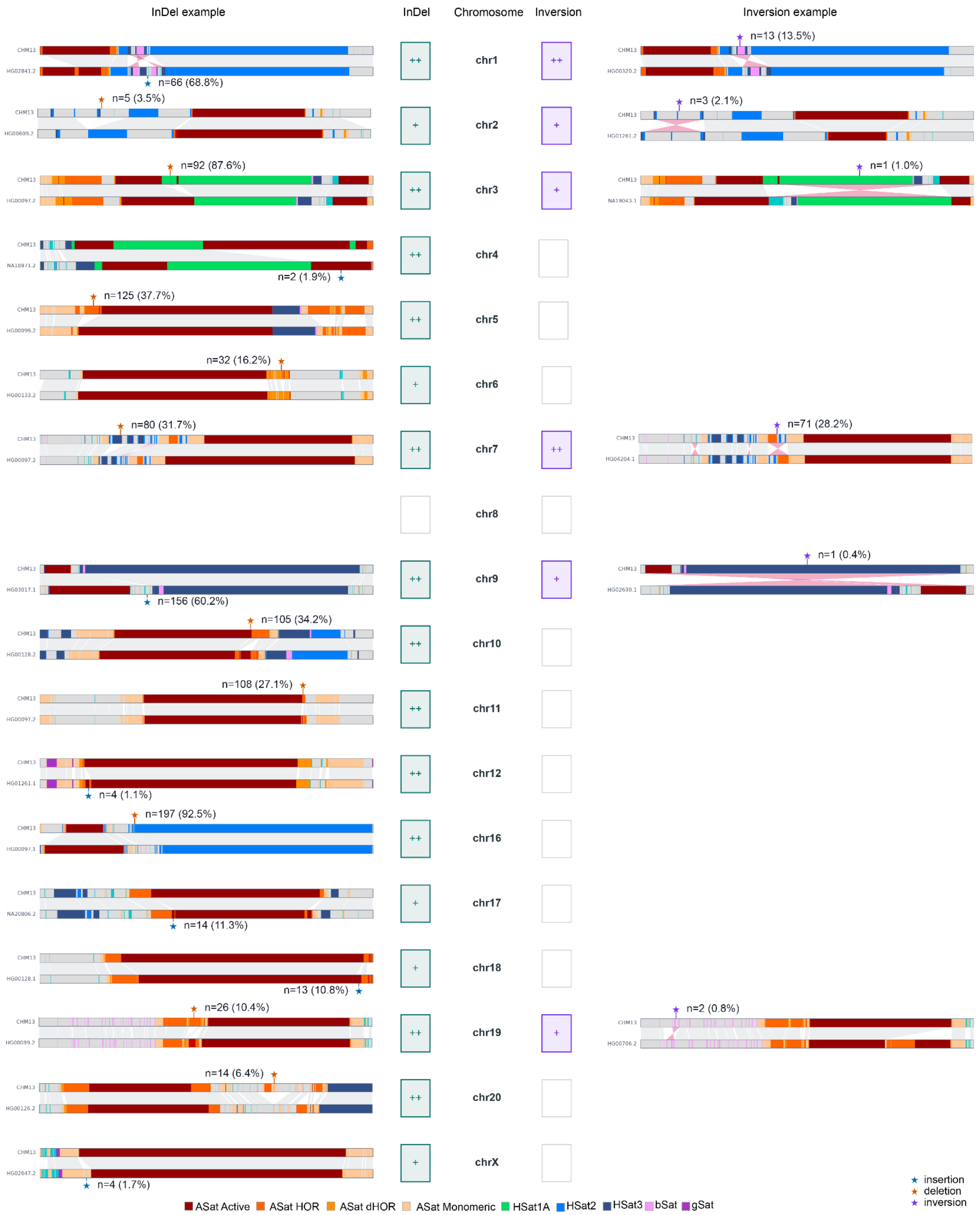


**Supplementary Figure 5. Select structural events in pericentromeres**

Large insertions and deletions combined as InDels are shown on the left and inversions shown on the right for all chromosomes analyzed for large structural changes in pericentromeric sequences. Status boxes are shown for each chromosome to indicate no large changes were detected (blank), at least one event is detected (+), or at least one event is found in over 20% of analyzed haplotypes (++) as compared to CHM13. Example events and haplotypes are shown for each chromosome with centromeric satellite annotations for CHM13 and a representative haplotype carrying the variant shown. Ribbons represent syntenic regions identified by satellite array-level alignment (Methods). Stars represent the variant position. The number and percentage of non-CHM13 haplotypes carrying the variant are shown. At most one example is displayed for each chromosome (and type).

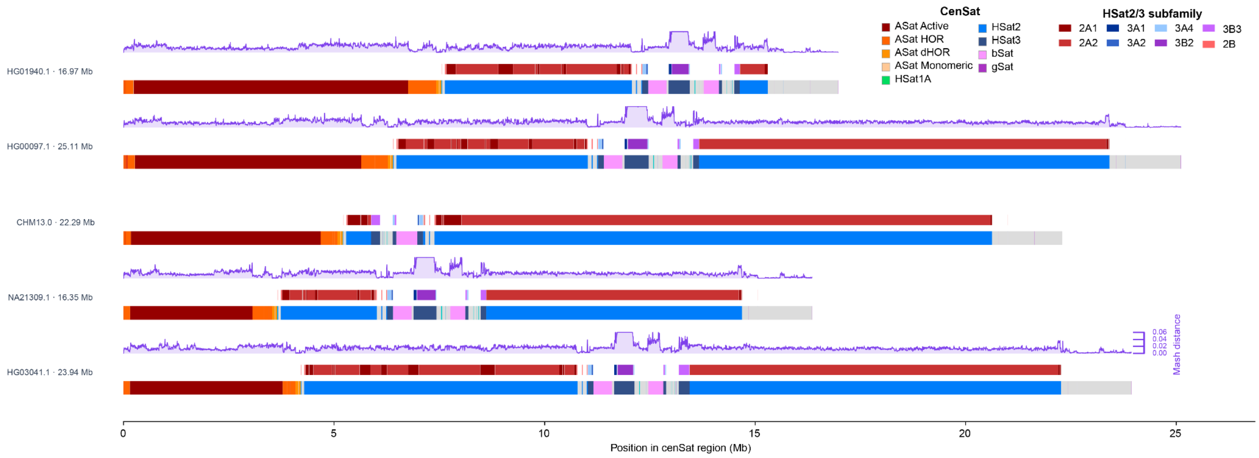


**Supplementary Figure 6. Inserted sequences in the pericentromere of chr1 are not found in CHM13**

Representative haplotypes with a large insertion as compared to CHM13 are shown. Centromere Satellite annotations are overlaid with k-mer based distances for 5 kbp windows against CHM13. The plotted distance is the minimum distance found for the window against all windows in CHM13 (genome wide).

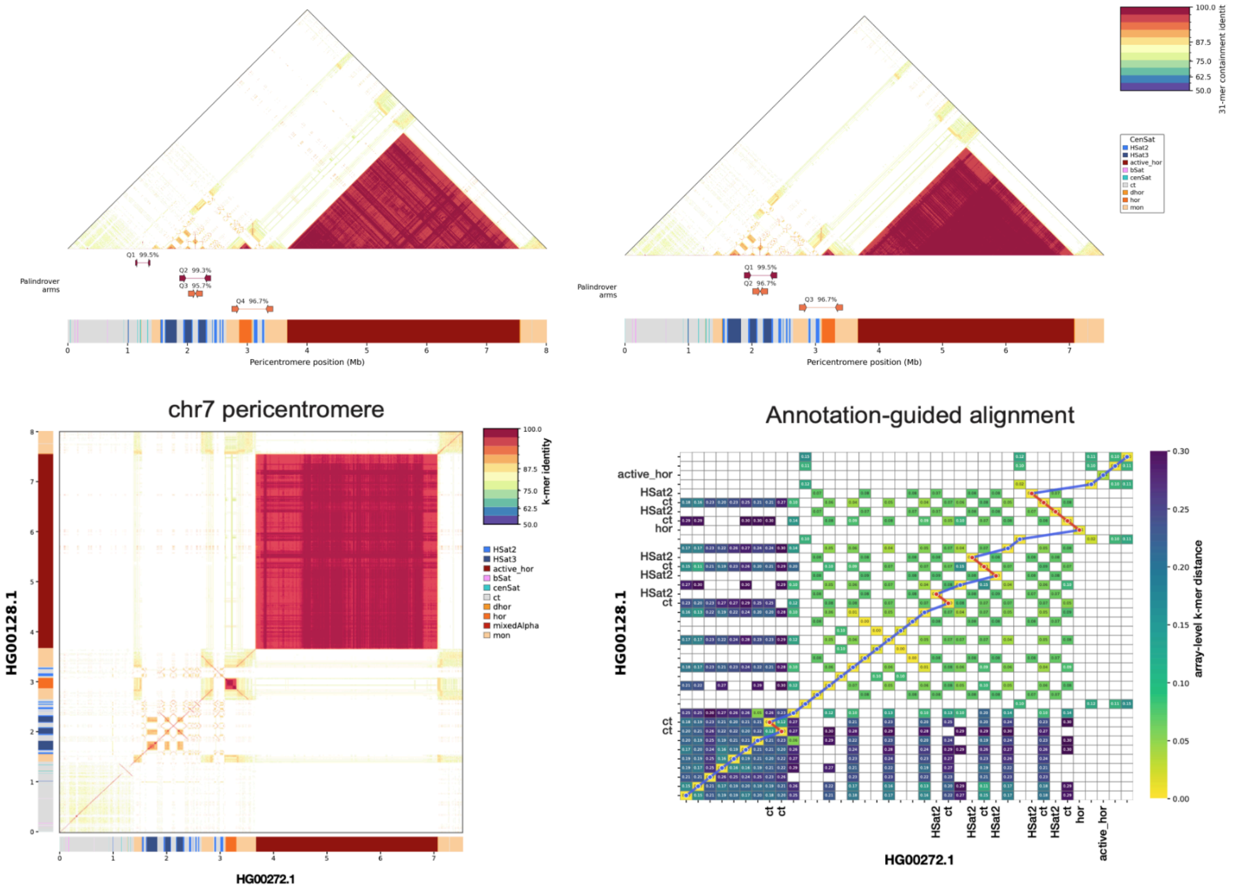


**Supplementary Figure 7. Inversions in chromosome 7 are found in palindromes**
a, Representative pericentromeres from two haplotypes of chromosome 7. CenSat annotations (bottom) are overlaid with palindromes called with Palindrover (see Methods) and k-mer self-identity dot plot. Vertical lines in dot plot correspond to inverted repeats b, Dot plot comparison of two pericentromeres shown in a. c, Pericentromere elements (arrays) shown with k-mer based distance and element-wise alignment (see Methods).

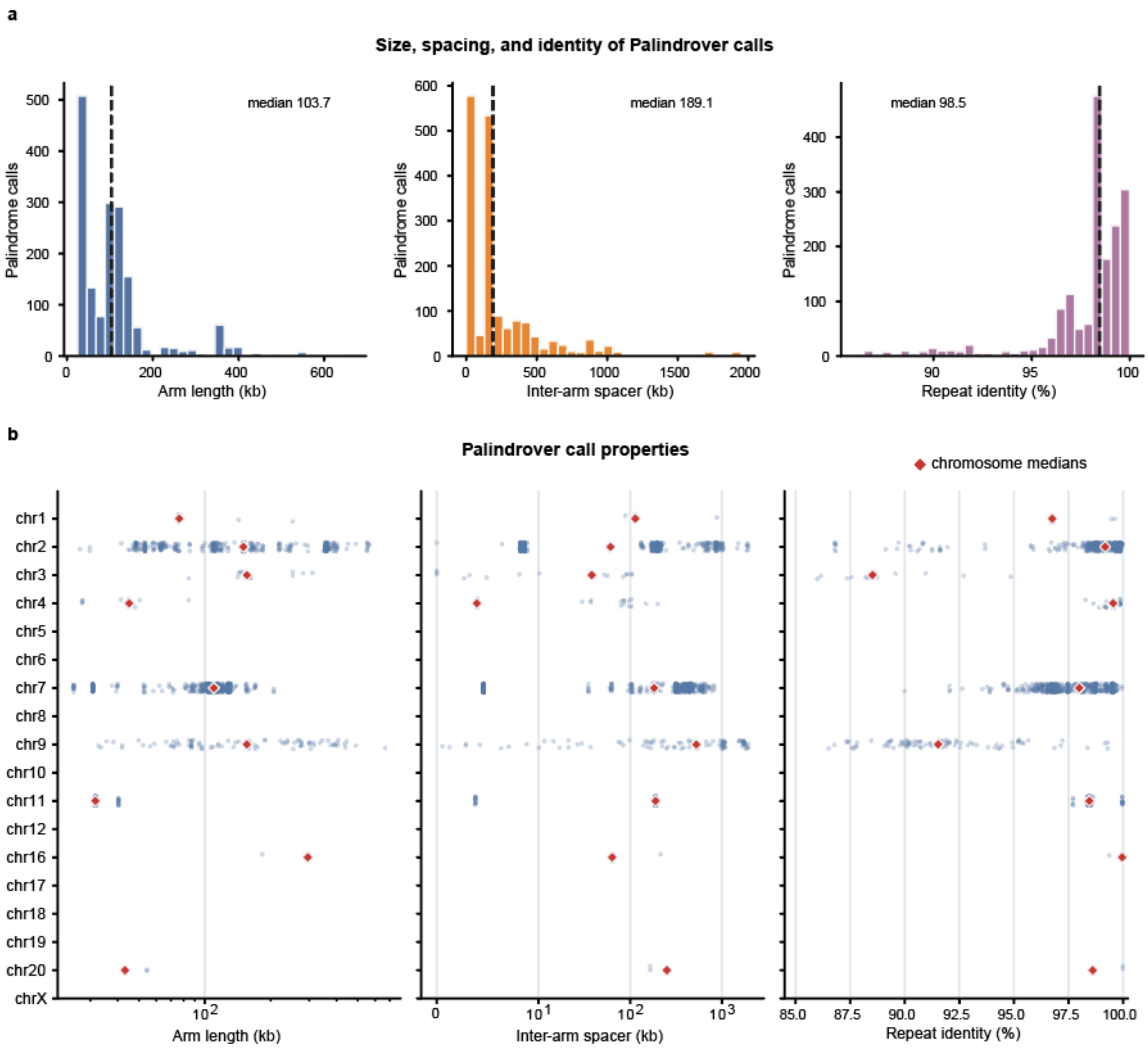


**Supplementary Figure 8. Palindrome properties in pericentromeres**

Properties of palindromes called by Palidrover genome wide (a) and broken down by chromosome (b).

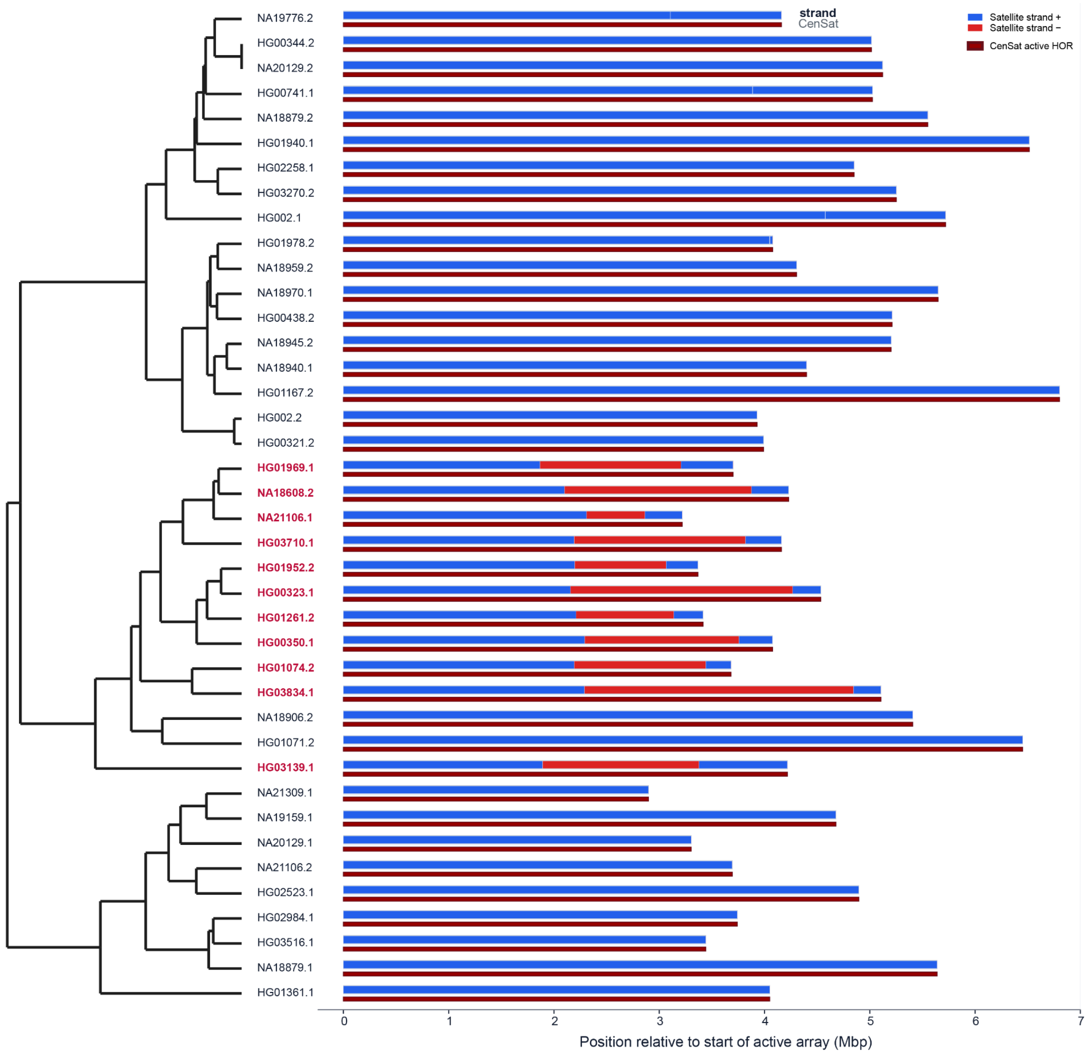


**Supplementary Figure 9. Chromosome 1 active-array inversion is restricted to a single cenhap clade**

Chromosome 1 cenhap tree (left) pruned to 40 randomly sampled haplotypes. Red haplotype names identify haplotypes carrying the internal inversion of the active array. On the right, satellite-strand orientation across the chr1 active array with plus-strand (blue) and minus-strand (red) intervals shown above CenSat tracks.

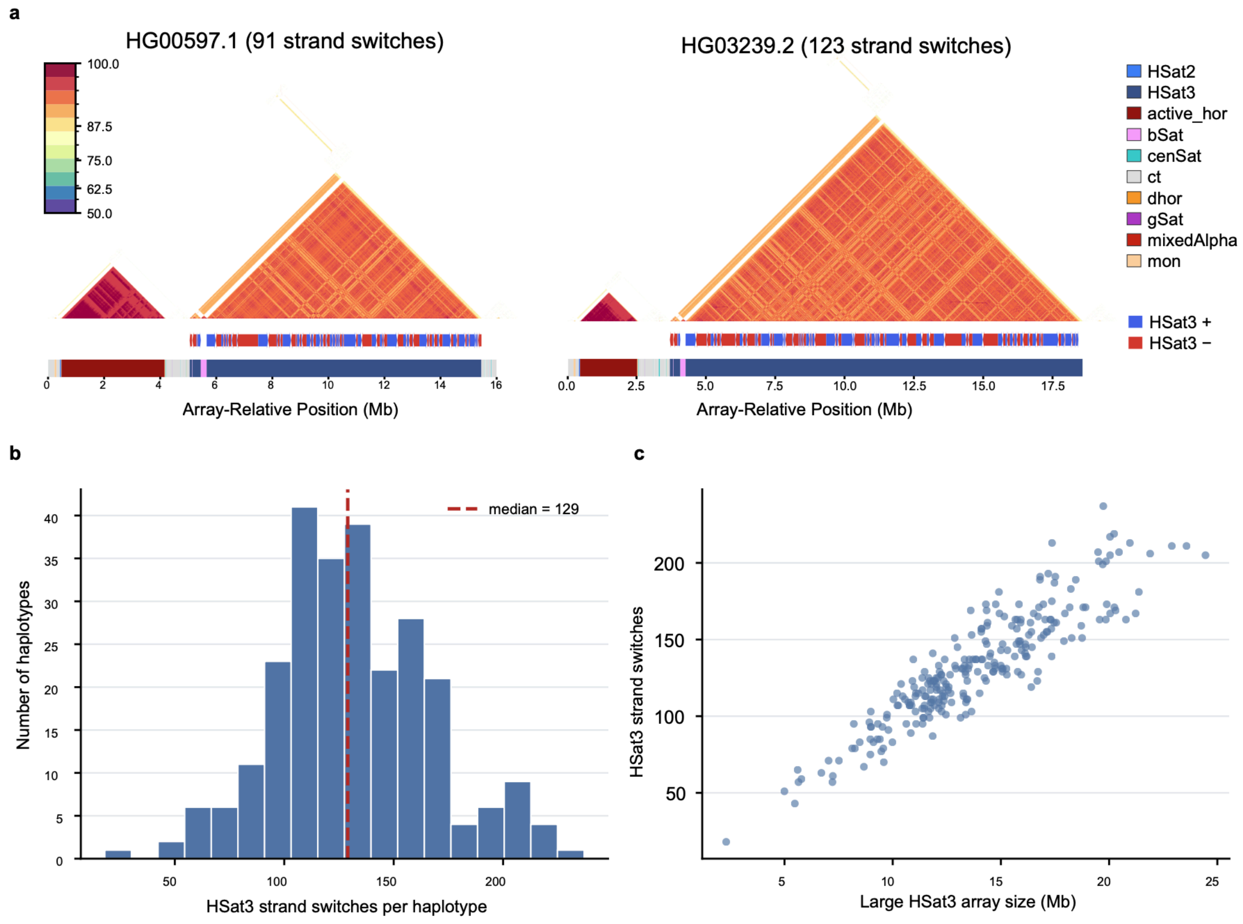


**Supplementary Figure 10. Strand switching in HSat3 array on chromosome 9.**
a, Representative pericentromeres from two haplotypes. CenSat annotations (bottom) are overlaid with annotated strand orientation (see Methods) and k-mer self-identity dot plot. Dot plot used canonicalized k-mers to avoid strand-based differences in identity. b, Histogram of the number of strand switches identified per haplotype. c, Size of (large) HSat3 array against the number of strand switches.

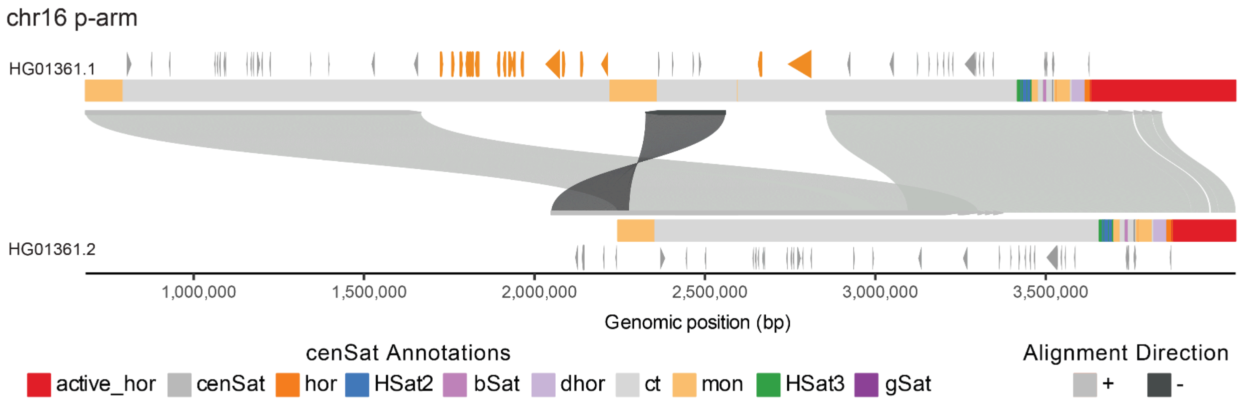


**Supplementary Figure 11. SVbyEye showing structural rearrangement on chromosome 16 p-arm between two haplotypes that involves the DUSP22 gene.**
Example of two haplotypes with different structural arrangements in the chromosome 16 p-arm pericentromeric region. Ribbons connect aligned sequences with light and dark grey indicating forward (+) and reverse (-) alignment orientations respectively. Centromeric satellite annotations are colored according to legend. Genes are shown along each haplotype, with genes that are inserted into one haplotype highlighted in orange and other genes shown in grey.

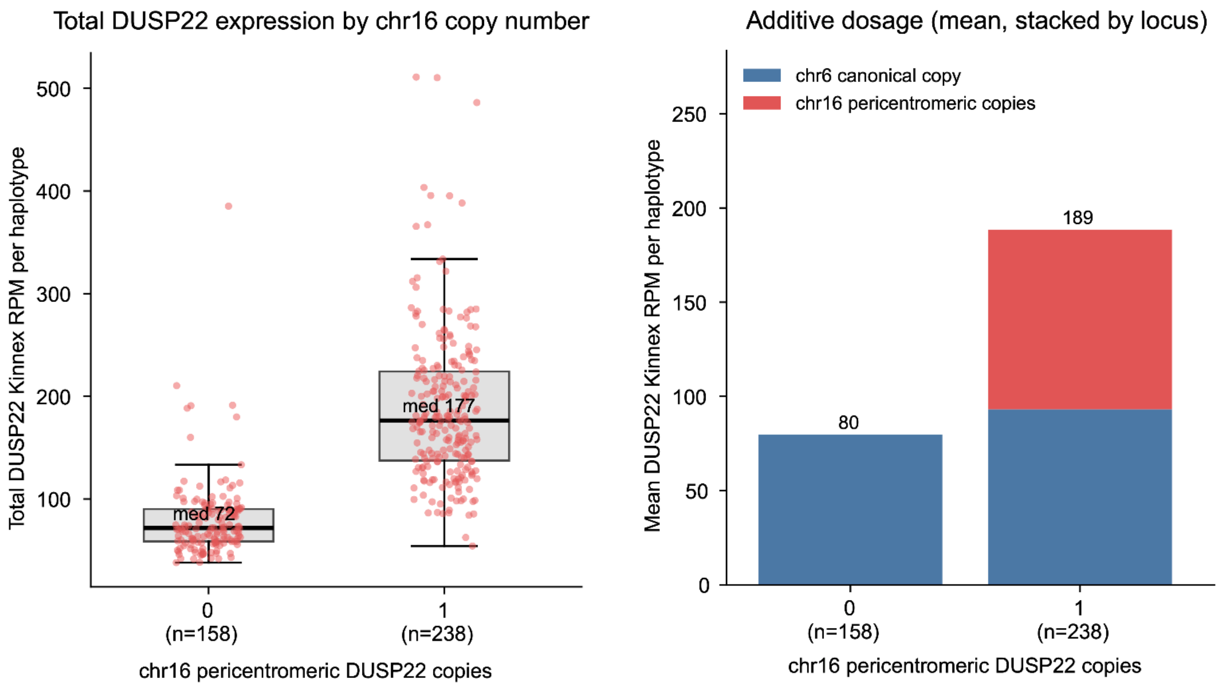


**Supplementary Figure 12. DUSP22 Expression in HPRC2.**
Left figure shows total DUSP22 RPM per haplotype, by whether 0 or 1 copy is present in the chromosome 16 pericentromere. All haplotypes have one copy of DUSP22 on chromosome 6.

Right figure shows the additive dosage of the DUSP22 expression when an additional chromosome 16 pericentromeric copy is present.

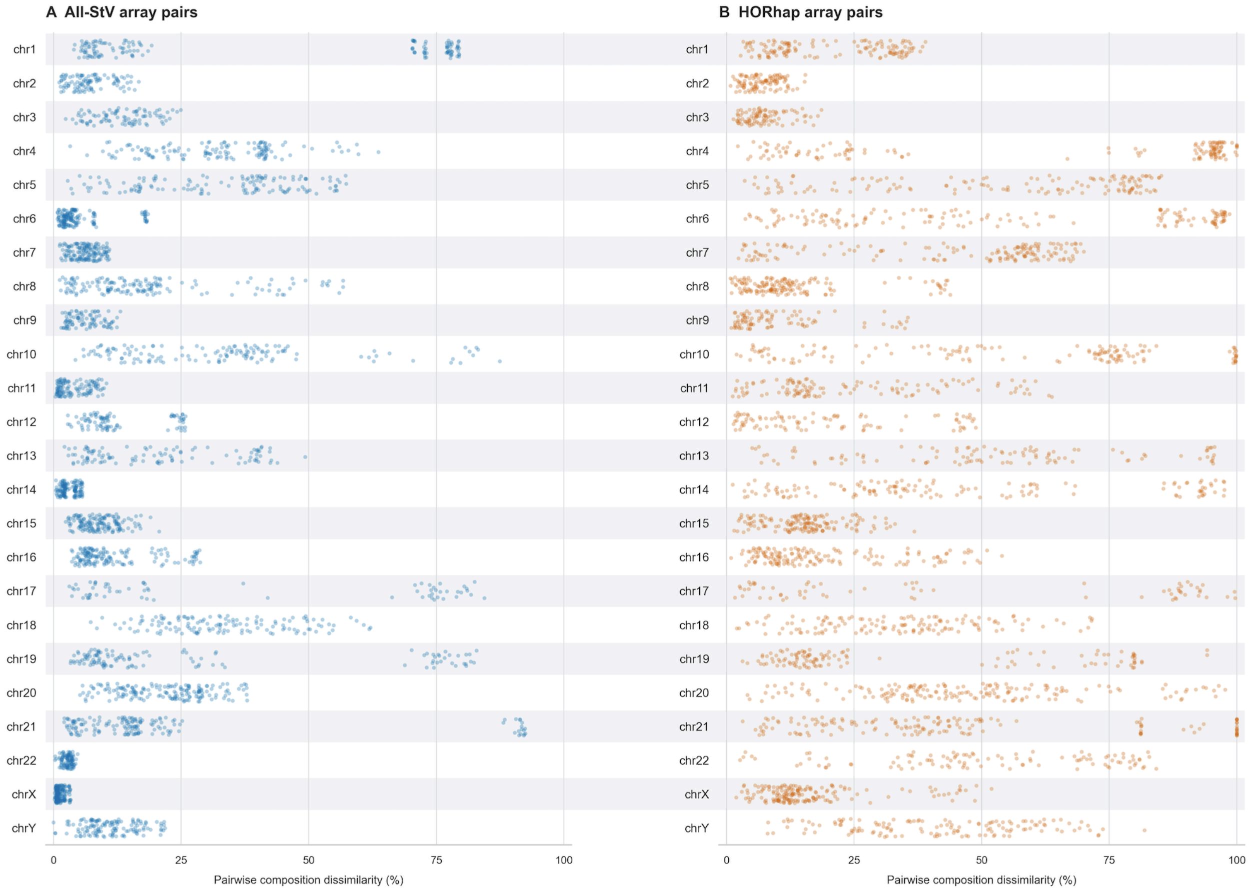


**Supplementary Figure 13. Pairwise StV and HORhap composition dissimilarity**
Dots represent Bray-Curtis dissimilarity for HOR-StVs (a) and HORhaps (b) from all pairs of 20 randomly selected haplotypes for each chromosome. Dissimilarity is calculated with normalized abundances.

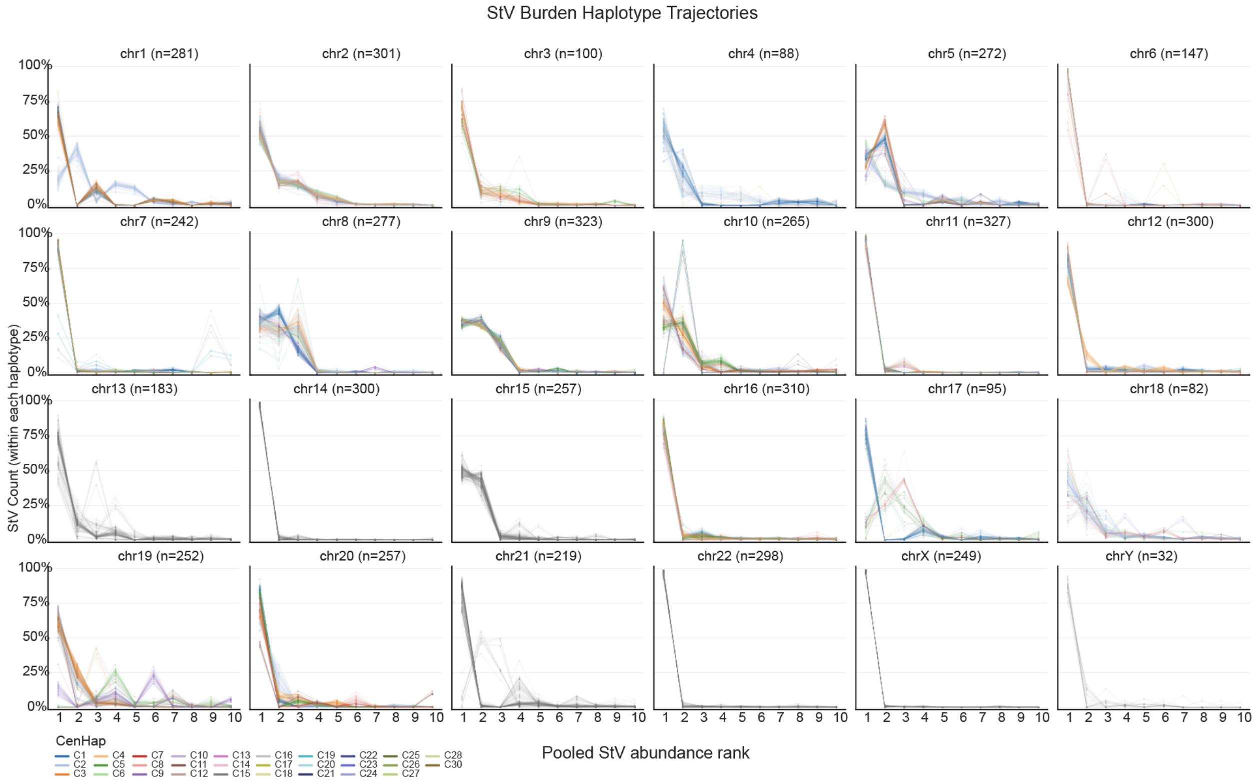


**Supplementary Figure 14. Active array structural variant (StV) composition per haplotype**
Per-haplotype StV burden for the top 10 StVs for each chromosome. Haplotypes are colored by cenhap assignments to distinguish cenhap-specific trends. Chromosomes without cenhap assignments are plotted in grey.

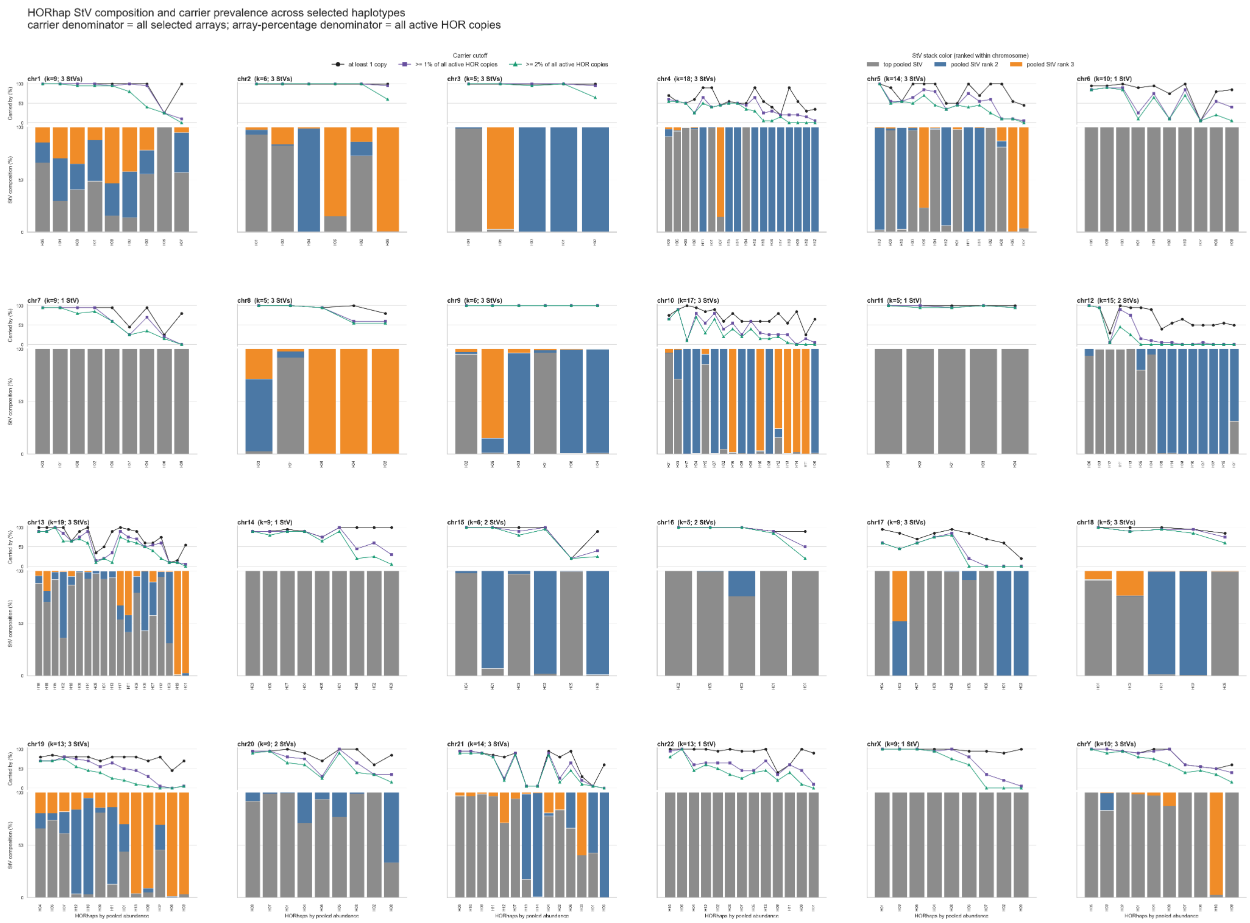


**Supplementary Figure 15. StV composition of HORhaps**

Stacked bar panels for each chromosome show HORhaps colored by HOR-StV composition. HORhaps are ordered by abundance. Upper plots show the percentage of the selected haplotypes is the percent of the selected arrays that have each HORhap at greater than 0%, 1%, 2%.

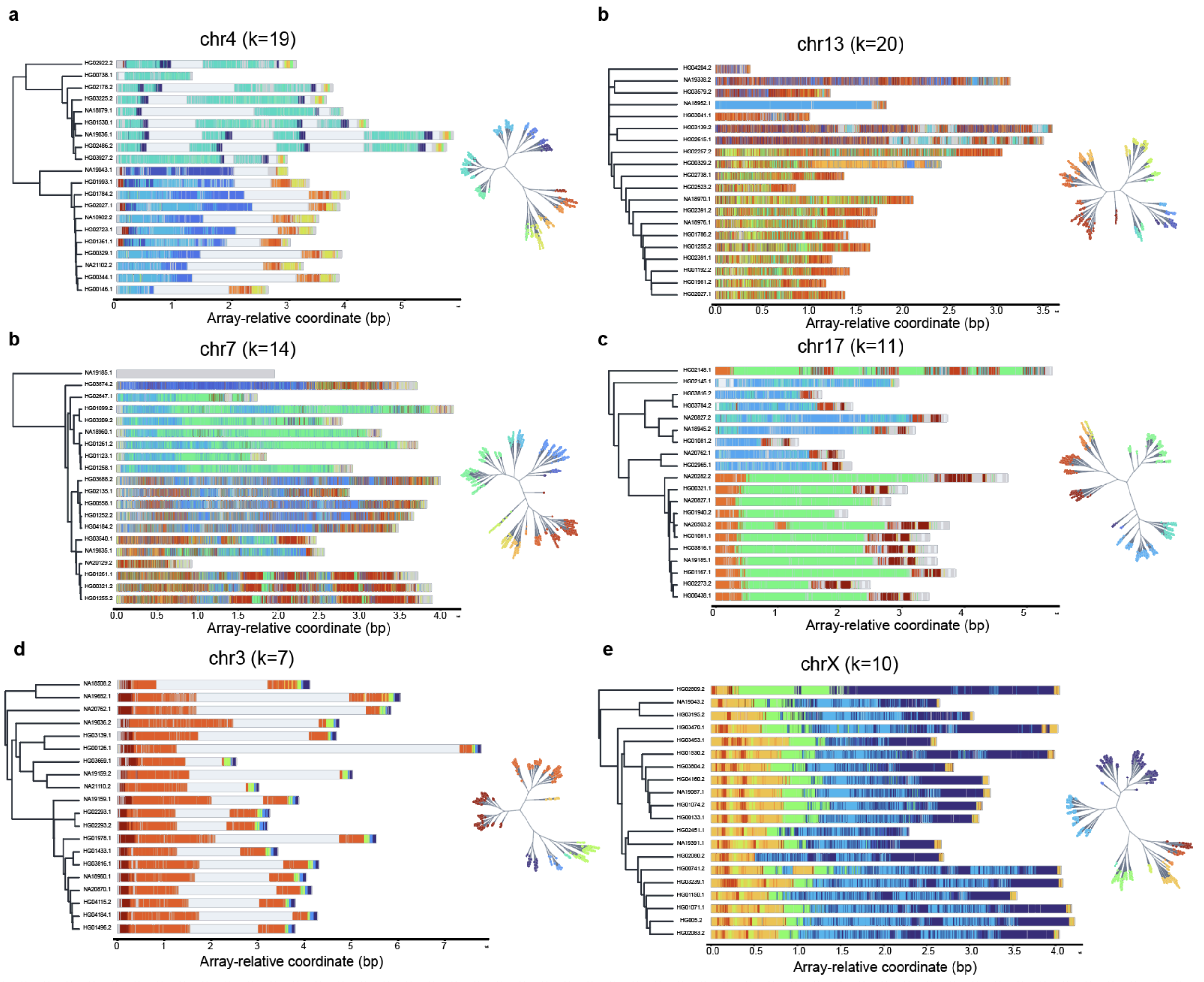


**Supplementary Figure 16. HORhap association with cenhap lineages**

Select chromosomes with (top four) and without (bottom two) visible HORhap association with cenhap lineages. Cenhap trees are shown to the left when available. When cenhap trees were not available, a UPGMA version of the Centrolign guide tree is shown (chr13). 20 haplotypes sampled across the tree were selected for analysis. To the right of the tree, the active array span is shown with individual HOR copies colored by their HORhap assignment. The top 3 HOR-StVs were used; other StVs are shown in grey, and non-active-HOR insertions are shown in pale grey. The top haplotype in chr7 (NA19185.1) is inverted reversing its StVs assignments and is shown in all grey. HORhap cluster numbers are shown in parentheses.  Unrooted ward trees (inset) are shown for each chromosome. Each terminal point represents one HOR copy and is colored according to its HORhap assignment.

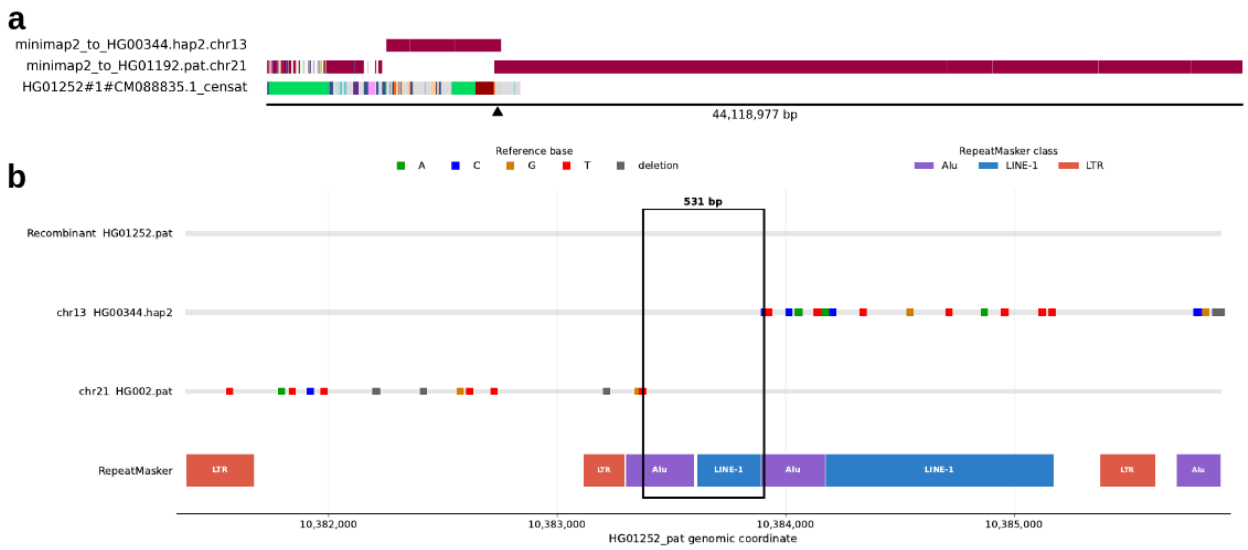


**Supplementary Figure. 17. Recombination breakpoint between acrocentric chromosomes 13/21.** a. Regions of homology to chr13 (top track) and chr21 (middle track) in a recombinant chr21 carrying a chr13-derived centromere, together with its CenSat annotation (bottom track). The q-arm is confidently assigned to chr21. The p-arm flank of the centromere corresponds to a region of high homology between chr13 and chr21, visible as an overlap between the minimap tracks. Therefore the region is technically an inter-chromosomal segmental duplication. The active αSat array (brown), HSat array (green), and right half of the p-arm show homology to chr13, whereas the left half of the p-arm shows homology to chr21. However, the p-arm results should be interpreted cautiously because this region is highly repetitive and highly similar between chr13 and chr21. A 400-kbp window spanning the minimap overlap was extracted and aligned. In the left part of this window, the recombinant showed higher similarity to chr13, whereas in the right part it showed higher similarity to chr21. b. A 4,500-bp alignment window containing a 531-bp mutation-free interval spanning Alu and LINE-1 elements, which likely contains the recombination breakpoint.

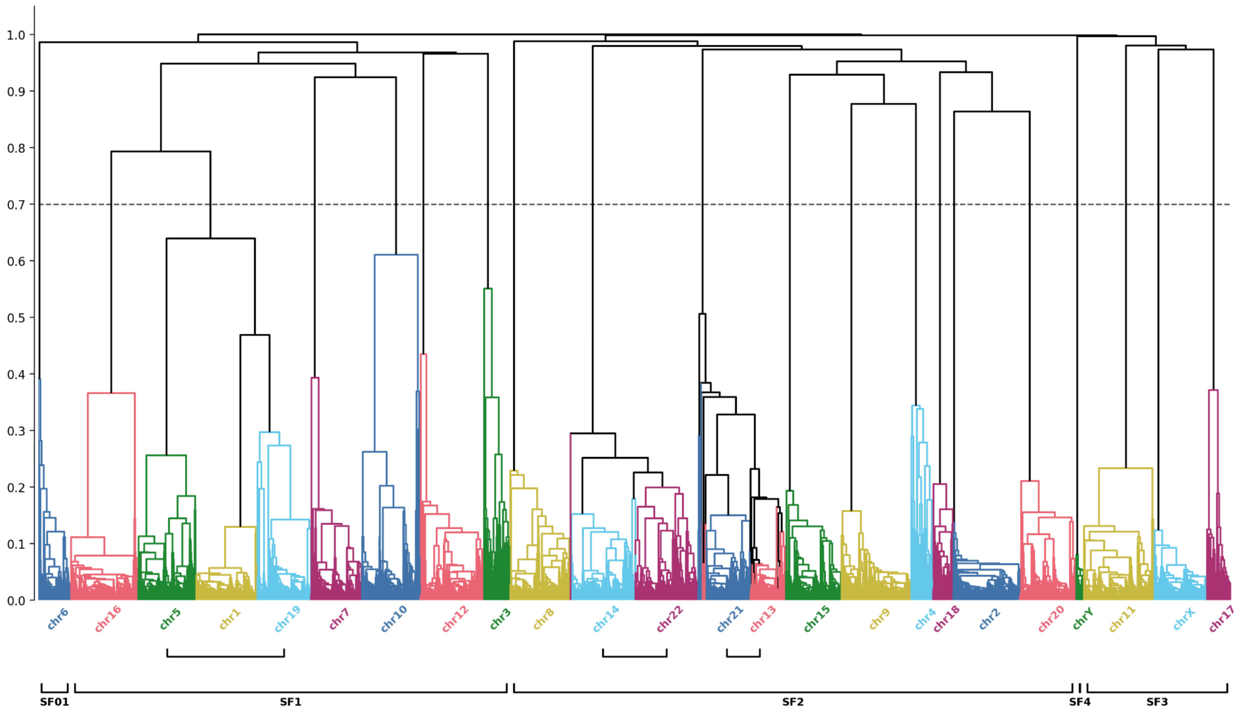


**Supplementary Figure 18. Genome Wide Active Array K-Mer Distance Tree.** UPGMA clustering of pairwise k-mer based distance matrix between active ASat arrays across all samples and chromosomes. The tree has 5 major clades which correspond to ASat SuperFamilies (SF01, SF1, SF2, SF3, SF4). Every chromosome forms its own branch if the tree is split at depth 0.7, except for chromosome pairs 13,21 and 14,22 and triple 1,5,19 which are known to be closely related. Some mixing between branches of chromosome pairs 13,21 and 14,22 is observed.

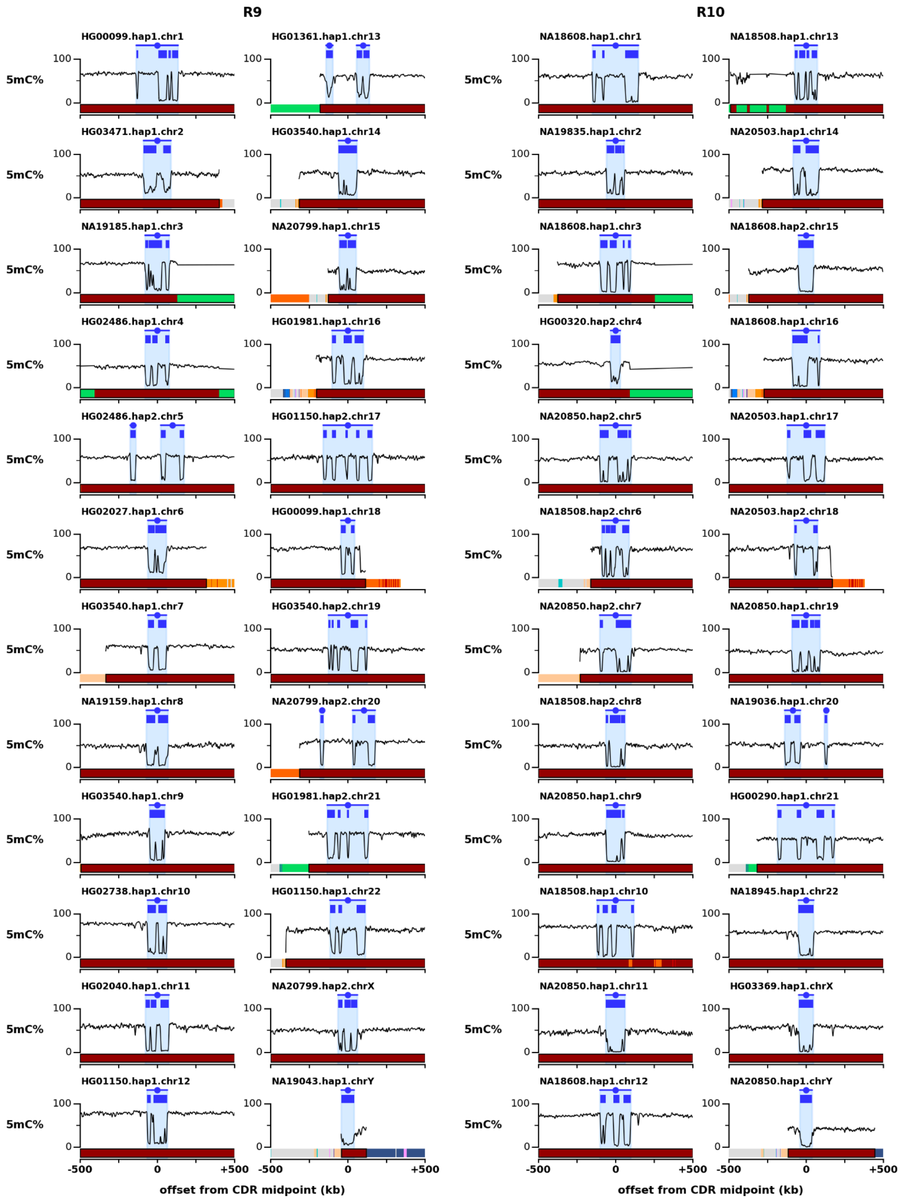


**Supplementary Figure 19. CDR call-set for HMM training centromeres.** Overviews of the CDRs and active αSat arrays used to train the CDR read-calling. One array for each chromosome and both R9 and R10 were selected. These arrays were hand-selected for their pronounced and discrete sub-CDRs. Plots are centered on the CDR-midpoint, shown on top in blue, a 5mC % line-plot is shown in the middle showing smoothed methylation values from centrodip, and CenSat track is shown on bottom.

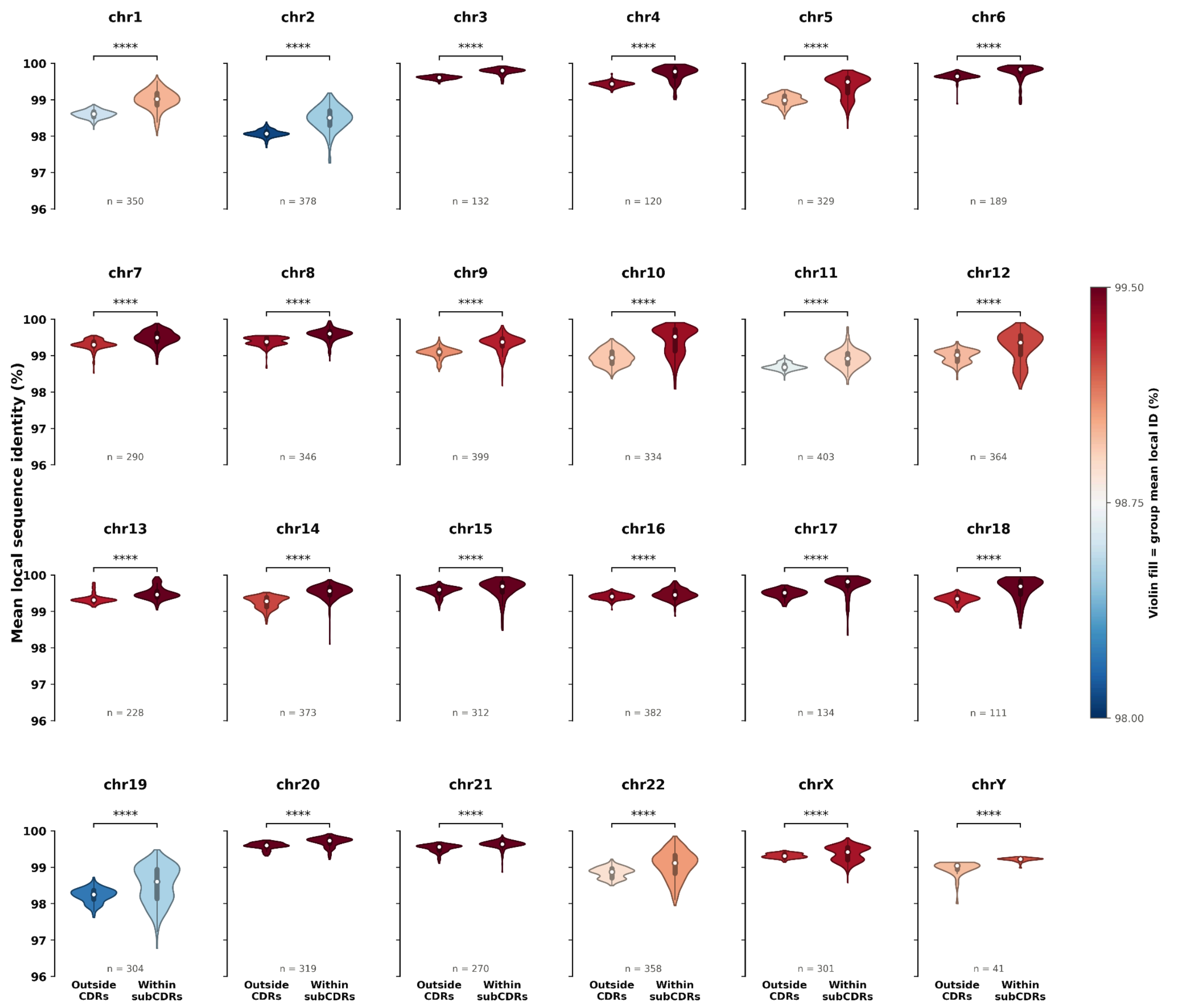


**Supplementary Figure 20. Chromosome-specific sub-CDR sequence-associations local identity percent.** Violin plots showing the haplotype-averaged local identity distributions of the active αSat and CDR for each chromosome. Per-chromosome violins are colored by their average local identity. Within the violins a white dot marks the median and boxplots show the IQR. All 24 chromosomes show a significantly elevated local identity within the CDR on average (one-sided Wilcoxon signed-rank test, $P < 10^{-4}$).

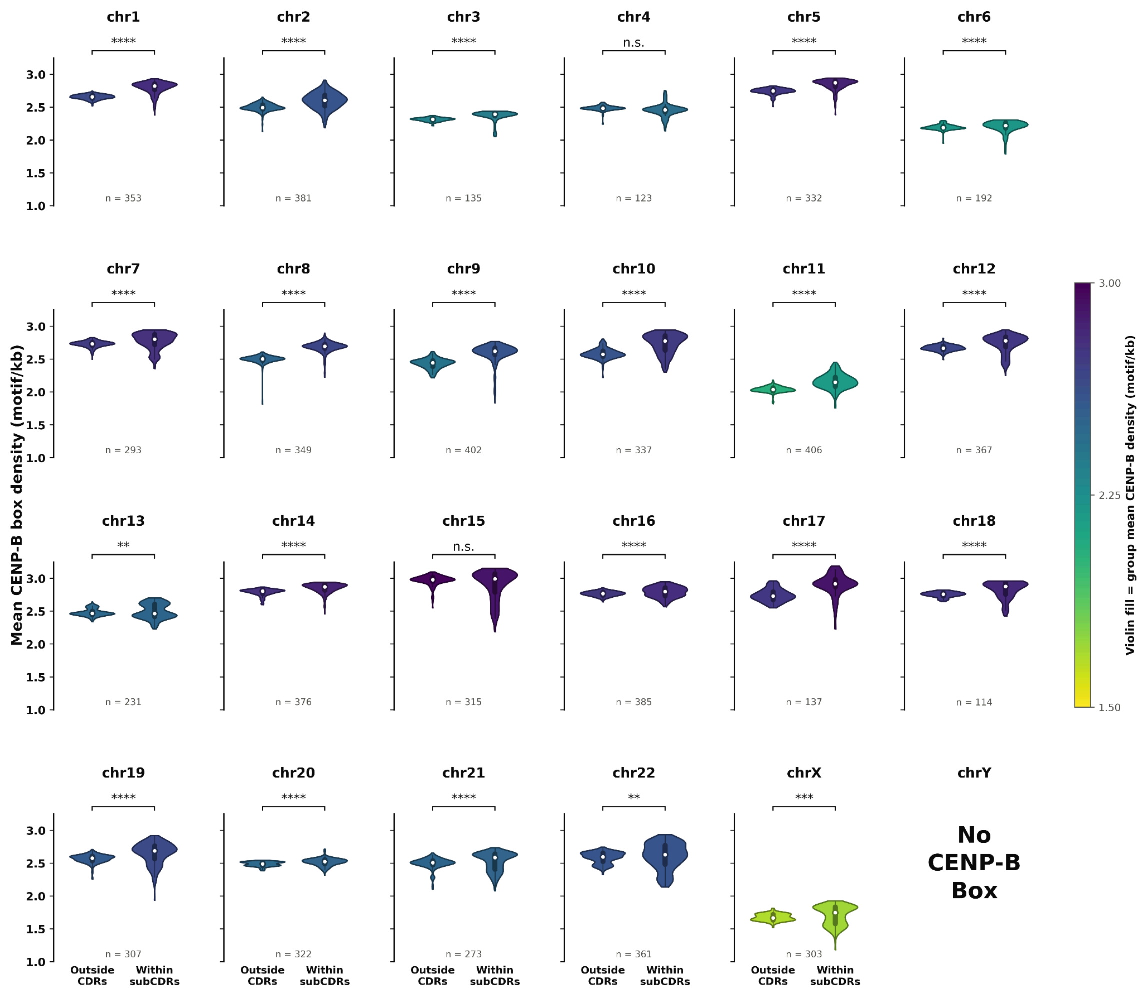


**Supplementary Figure 21. Chromosome-specific sub-CDR sequence-associations CENP-B box density.** Violin plots showing the haplotype-averaged CENP-B box density distributions of the active αSat and CDR for each chromosome. Per-chromosome violins are colored by their average CENP-B box density. Within the violins a white dot marks the median and boxplots show the IQR. 21/23 chromosomes represented here show a significantly elevated local identity within the CDR on average (one-sided Wilcoxon signed-rank test, $P < 10^{-2}$).

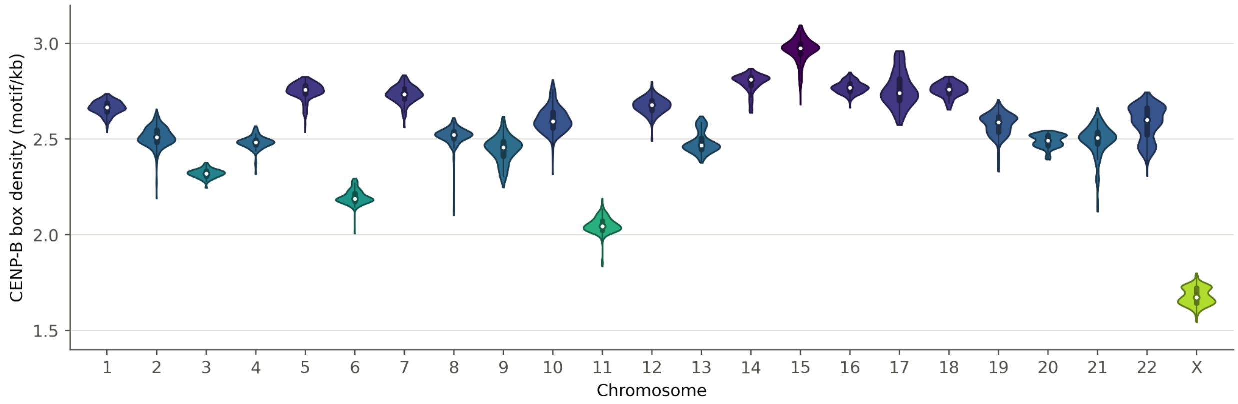


**Supplementary Figure 22. Chromosome-specific CENP-B box density.** CENP-B box density averaged per active αSat, shown as per chromosome violins. Each chromosome has its own density patterns, driven by StV content and change in existing CENP-B boxes. Chromosomes 6, 11, and X display uniquely low CENP-B box densities.

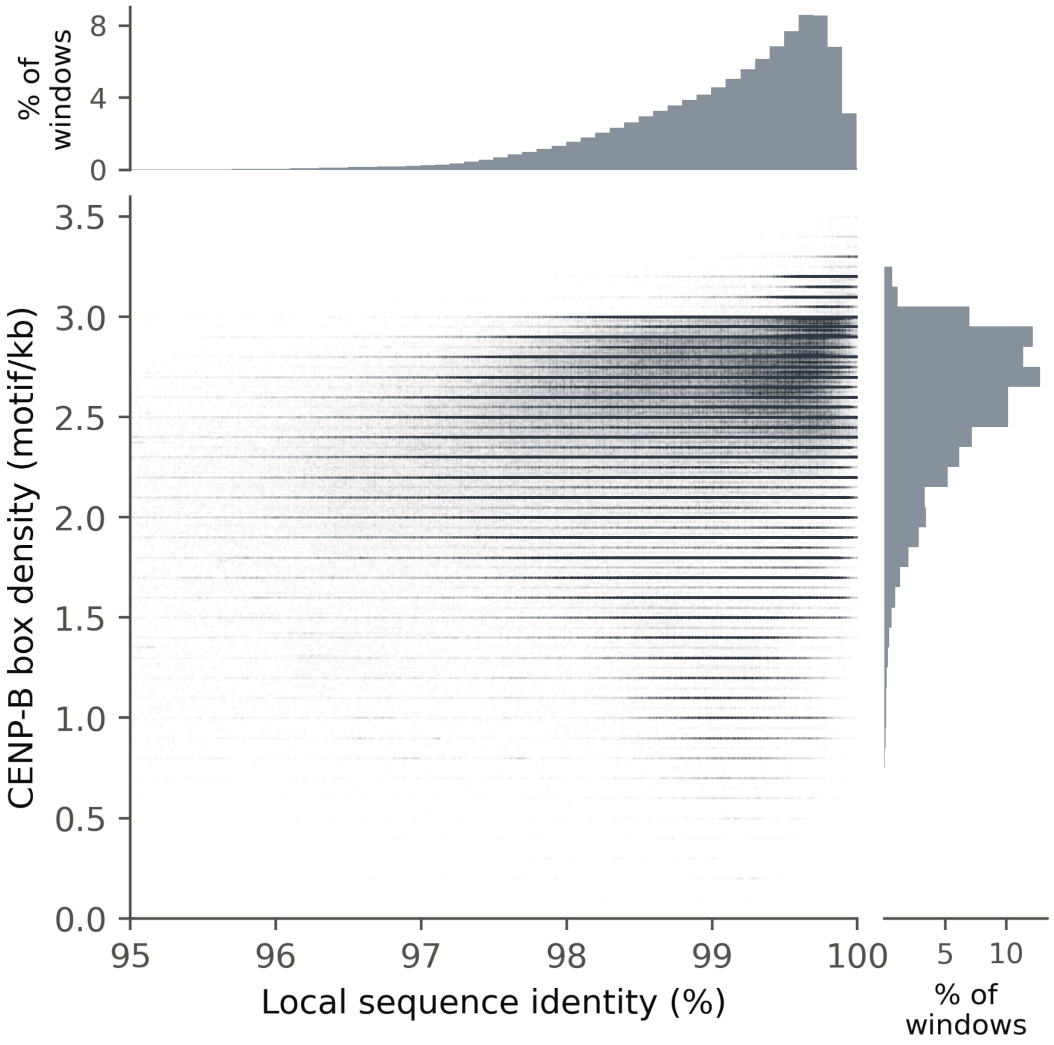


**Supplementary Figure 23. CENP-B box density vs local identity window values.** CENP-B box density and local identity are covariate. Regions within the active αSat with higher CENP-B box density often have higher local identity, and vice versa.

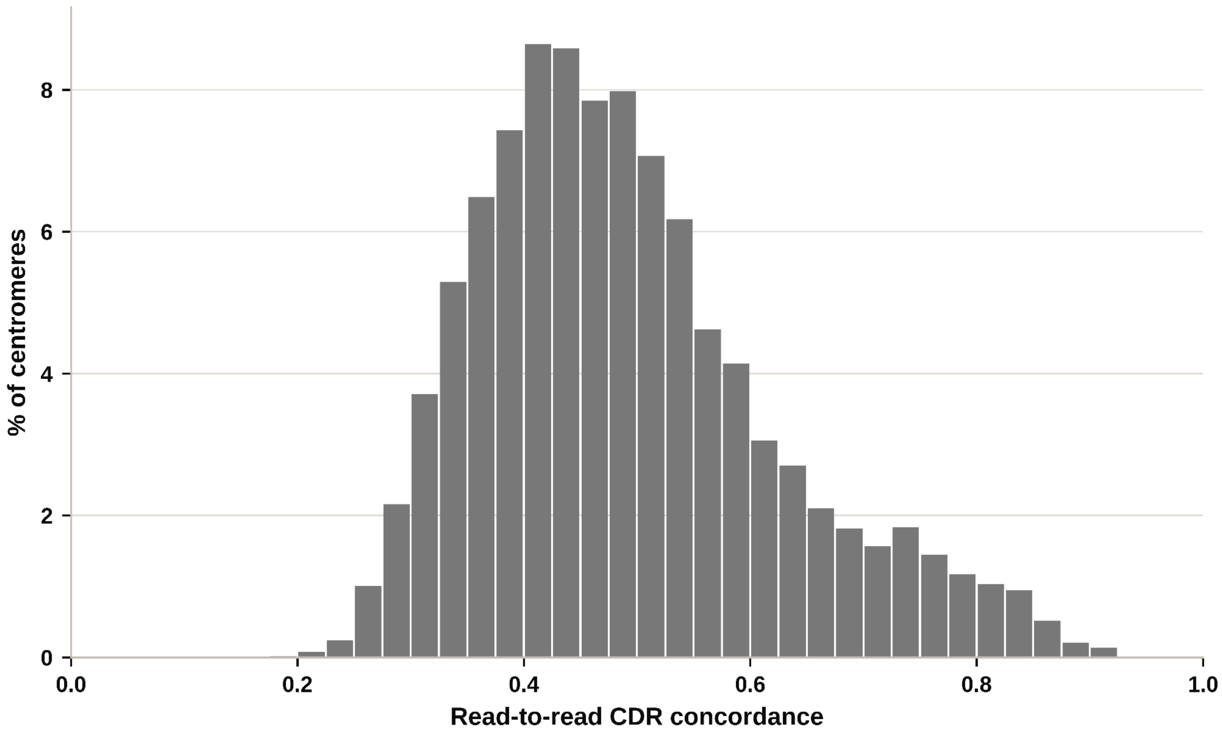


**Supplementary Figure 24. Read-to-read concordance distribution.** Histogram of the read-to-read concordance values (Methods) from 6,718 centromeres. All centromeres used to train read-based CDR predictions were excluded.

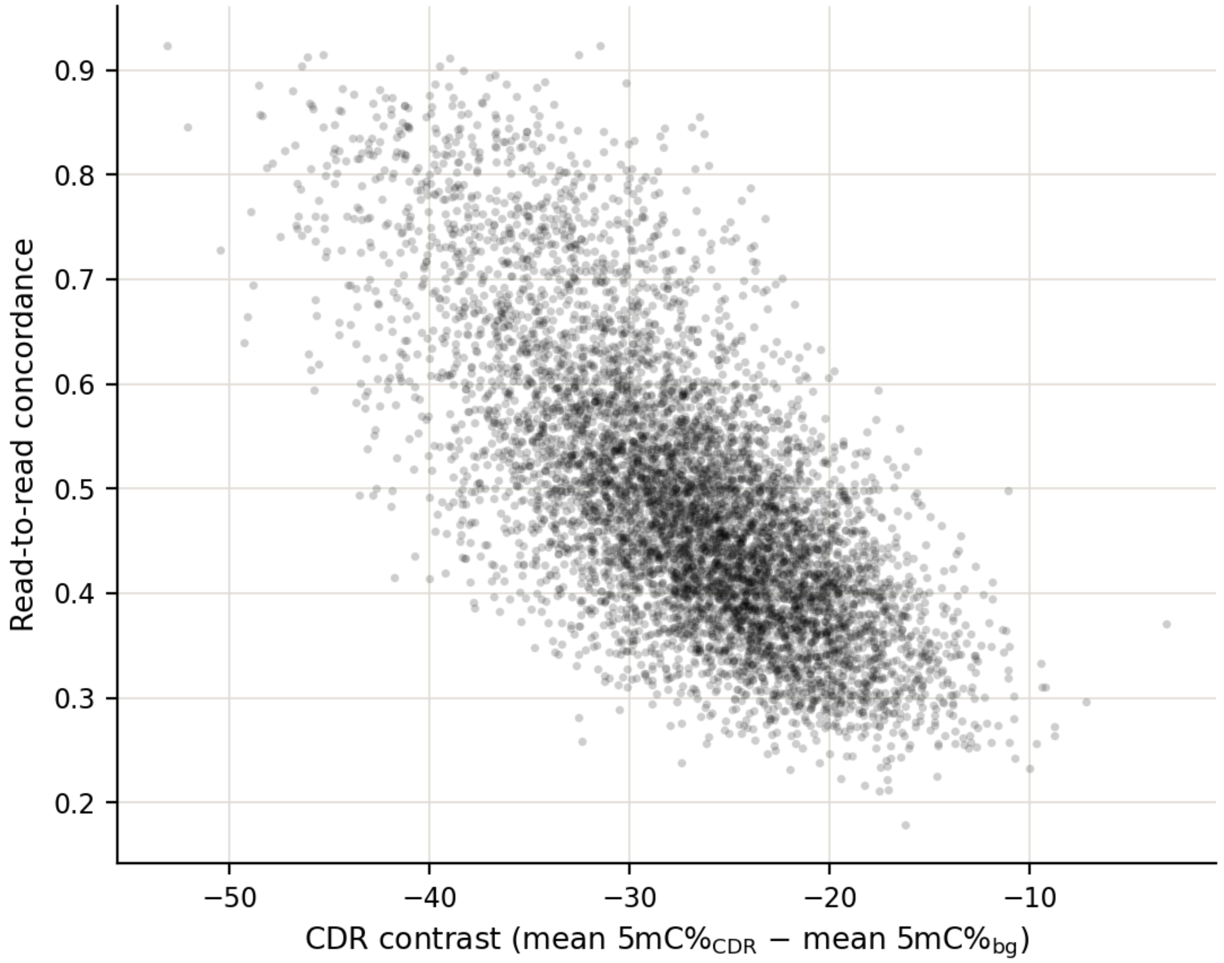


**Supplementary Figure 25. CDR contrast vs pooled array Jaccard.** Read-to-read concordance within a centromere was negatively correlated with CDR contrast. CDR contrast was calculated as the average 5mC% within the CDR minus the average 5mC% within the rest of the active αSat. Suggesting that shallower aggregate CDR signals may be masking read-to-read heterogeneity.

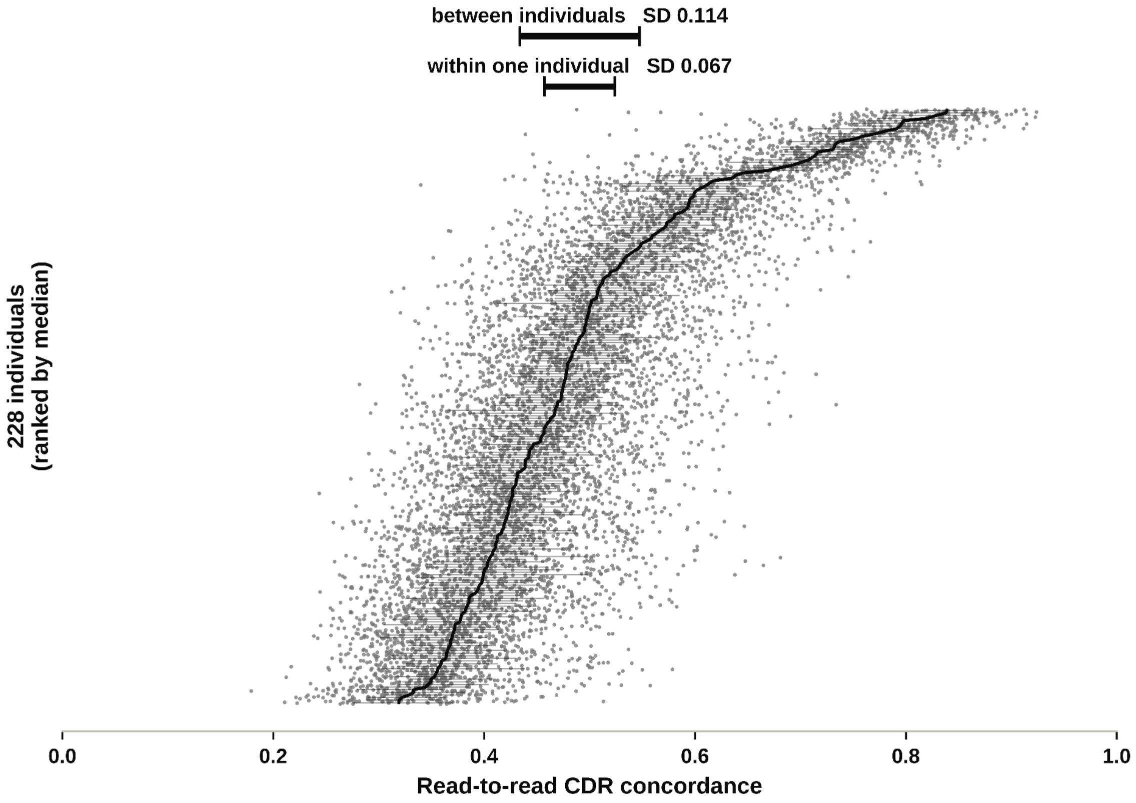


**Supplementary Figure 26. Pooled Jaccard within-sample vs between sample variation.** Scatter plot plot of read-to-read concordances along the x-axis, and HPRC2 individuals along the y-axis. Individuals are ranked along the y-axis by their mean read-to-read concordance. Standard deviation within one individual is nearly half of what it is between individuals. Suggesting that the read discordances we see are largely sample-driven.

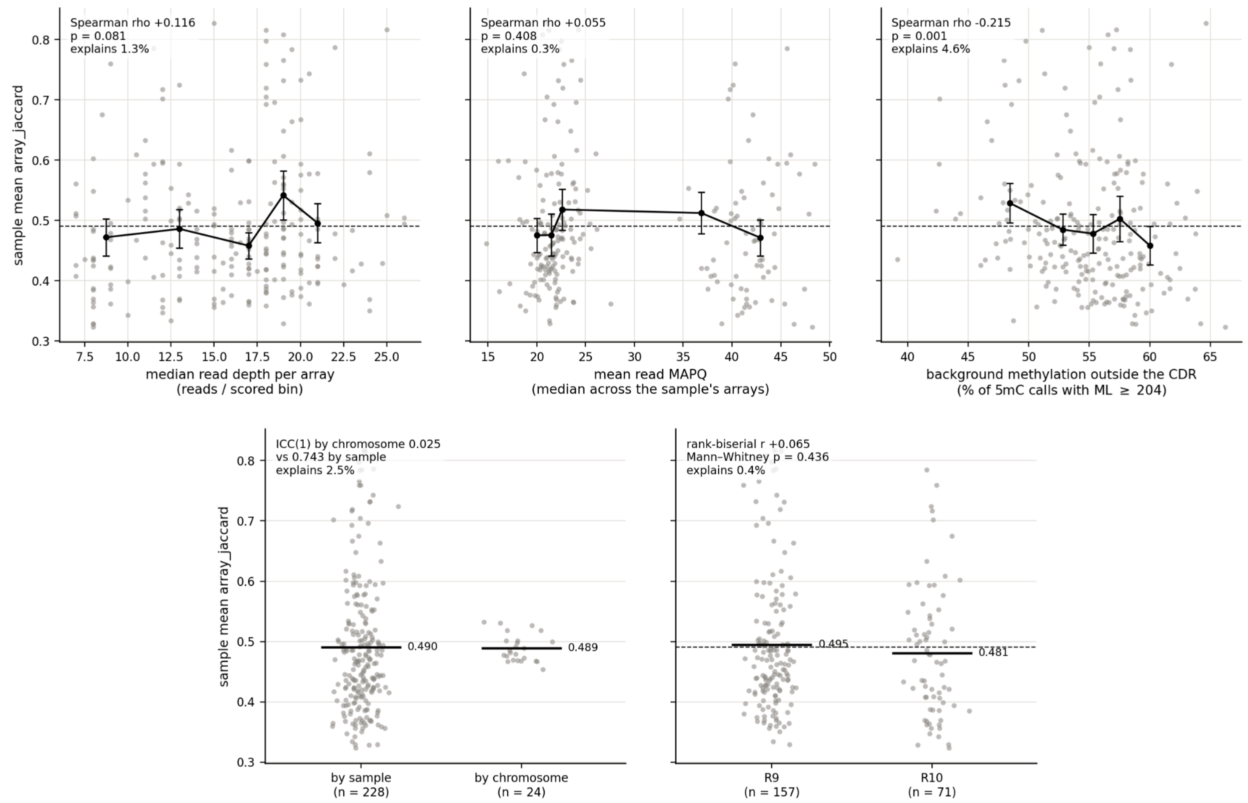


**Supplementary Figure 27. Per-centromere read concordance vs confounding variables.** Five potential confounding variables we ruled out as being the cause for the low read-to-read concordance values we identified. We examined read depth, MAPQ, background 5mC%, chromosome, and ONT sequencing chemistry. Of these 5 the only one that shows statistical significance was the background 5mC%. However, despite the statistical significance, the effect was rather subtle with a Spearman ρ of -0.215.

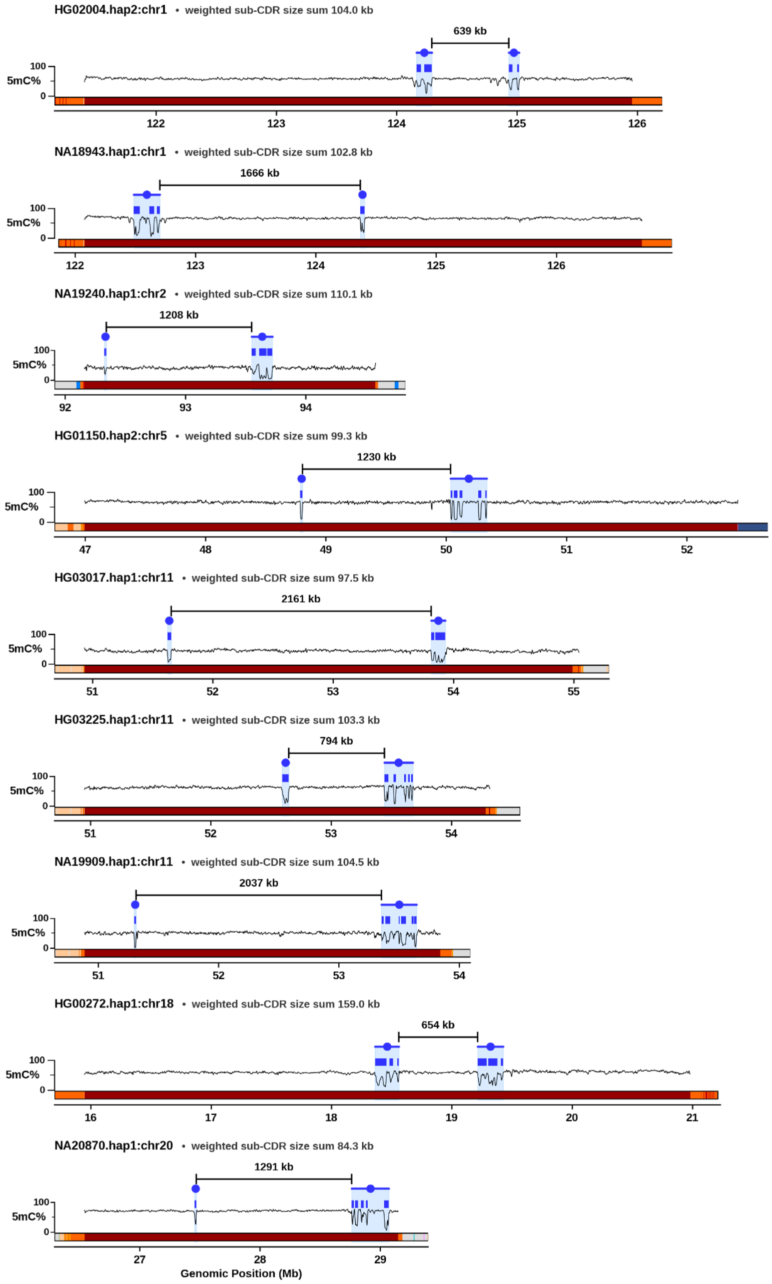


**Supplementary Figure 28. 9 centromeres with sub-CDR gaps over 500kbp.** Active αSat overviews for the 9 centromeres we identified in HPRC2 that had sub-CDR gaps over 500 kbp. Tracks show CDRs in blue on top, 5mC% as a line-plot, and CenSat tracks on the bottom. Centromere identities, sub-CDR gaps over 500 kbp, and weighted sub-CDR size sums are labelled within the plot.

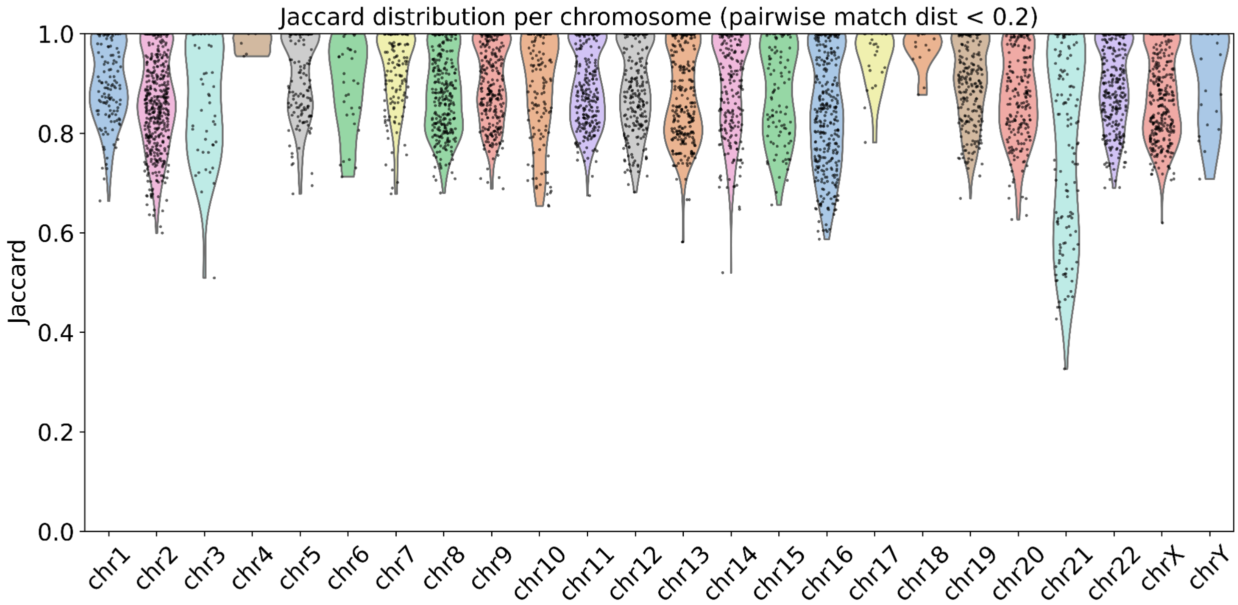


**Supplementary Figure 29. Jaccard similarity of induced and direct pairwise alignments**
Jaccard similarity between induced (MSA) and direct Centrolign pairwise alignments (subset to pairwise match distance < 0.2).

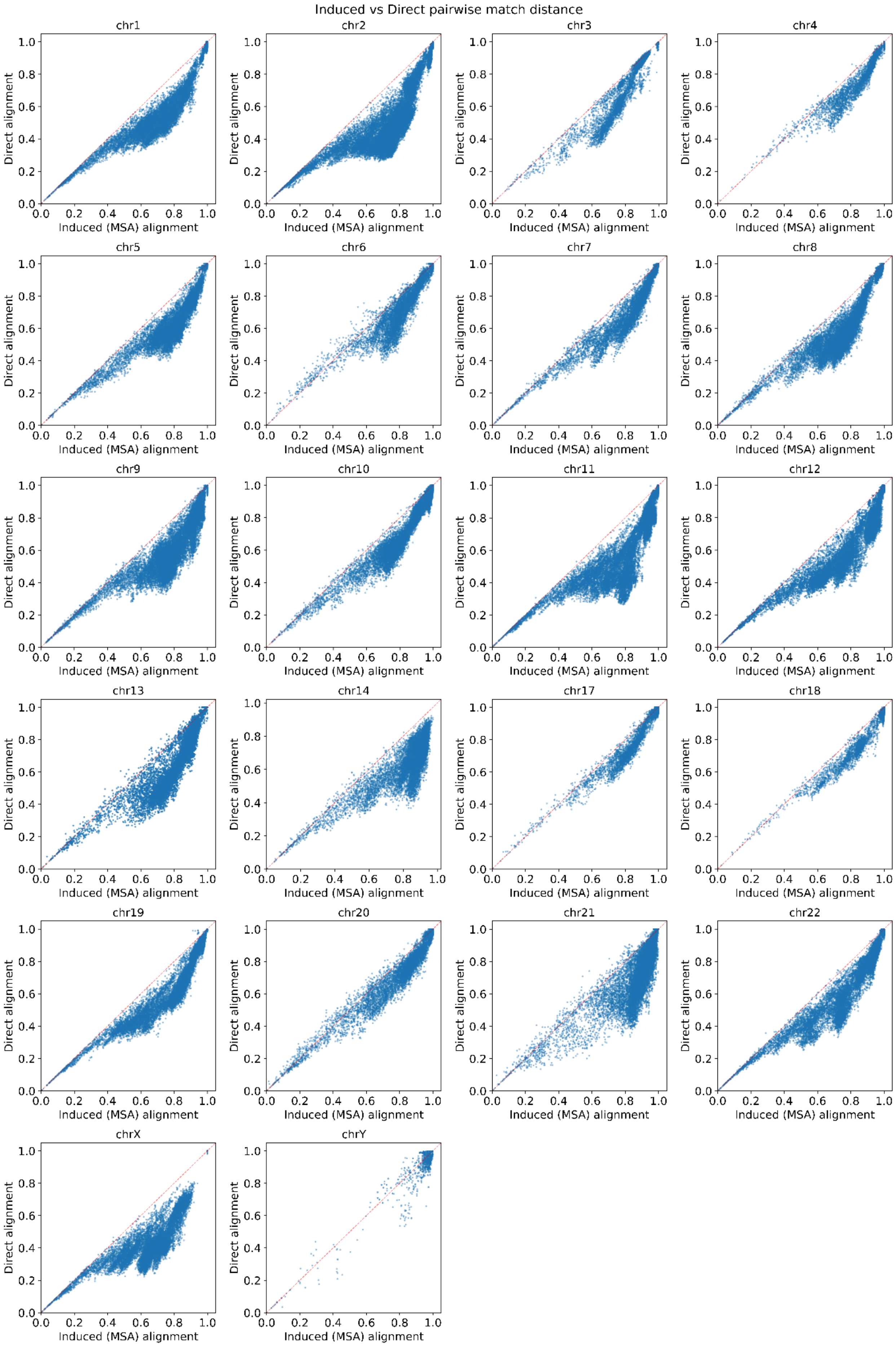

Induced vs Direct pairwise match distance
chr1
chr2
chr3
chr4
chr5
chr6
chr7
chr8
chr9
chr10
chr11
chr12
chr13
chr14
chr17
chr18
chr19
chr20
chr21
chr22
chrX
chrY
Direct alignment
Induced (MSA) alignment
0.0
0.2
0.4
0.6
0.8
1.0

**Supplementary Figure 30. Centrolign induced vs. direct pairwise match distance**
Pairwise match distance ((2 * # mismatches) / (query_len+ref_len)) calculated from pairwise alignments induced from the Centrolign MSA (x axis) and Centrolign direct pairwise alignments (y axis).

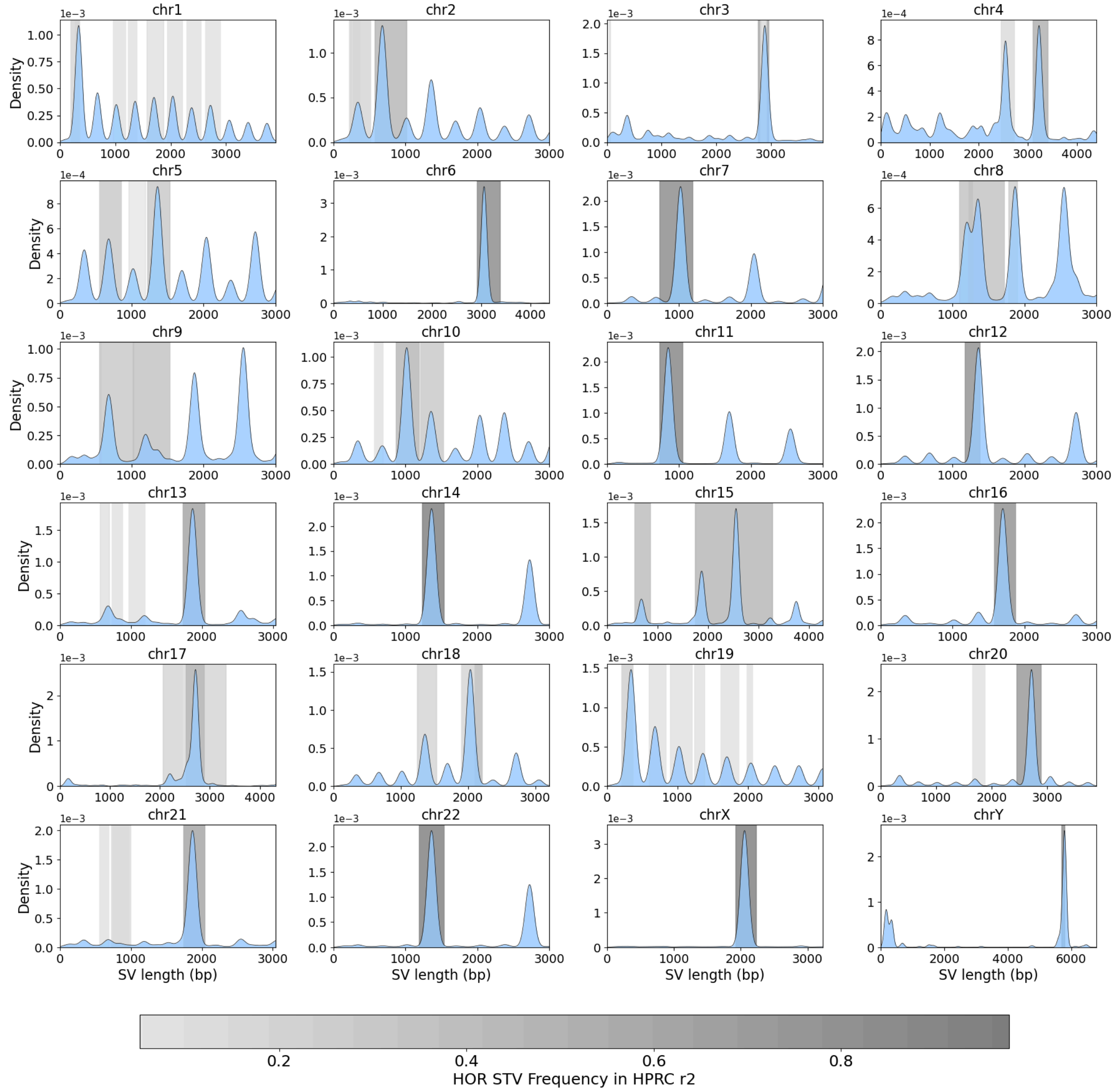


**Supplementary Figure 31. Centrolign SV sizes compared with observed HOR-StV sizes**
Histogram of SV sizes derived from Centrolign induced pairwise alignments (blue). The plot is showing sample pairs where pairwise match distance < 0.2. Grey bars indicate the size of

expected STV counts derived from HPRC release 2 annotations, with shading indicating their frequency in HPRC release2.

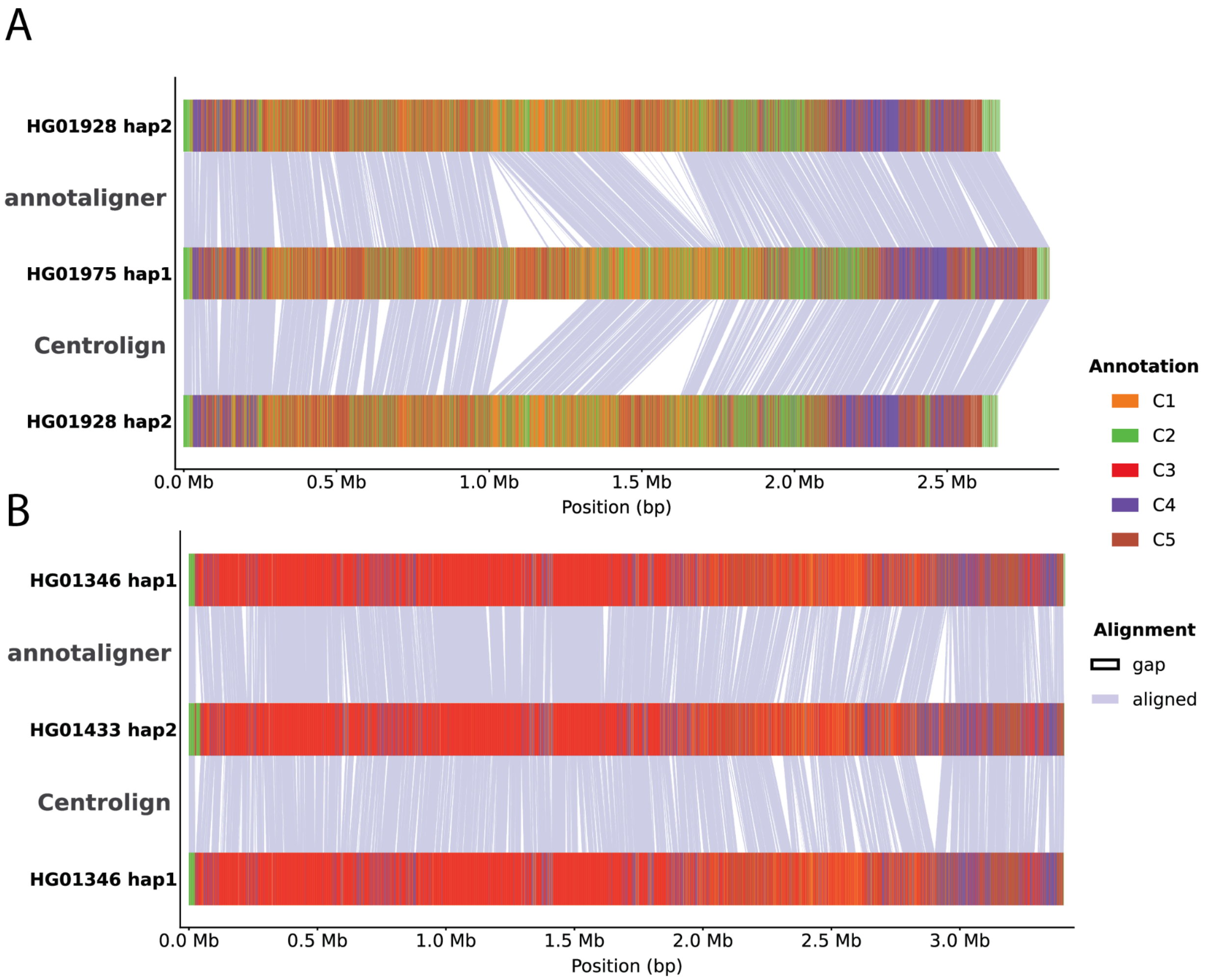


**Supplementary Figure 32. Example of Centrolign and annotaligner alignments**
Comparison of alignments produced by Centrolign and annotaligner on the same sample pairs. Both sample pairs (A and B) are from chromosome 11. Bed tracks are colored by HORHap annotation using k=5. Aligned sequence is shaded in grey, white represents gaps.

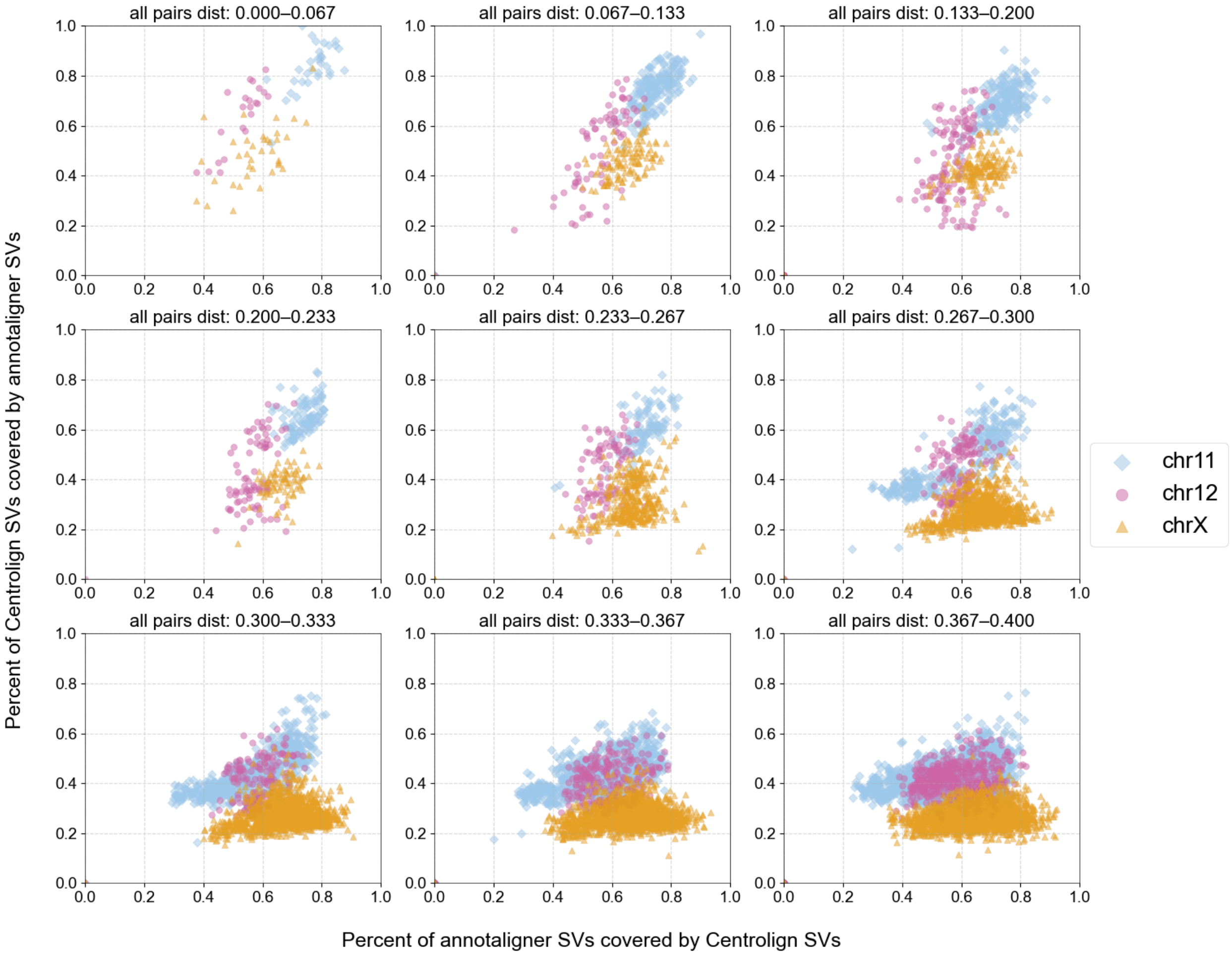


**Supplementary Figure 33. SV concordance between annotaligner SVs and Centrolign SVs.** The y axis represents, per sample pair, the percent of centrolign SVs covered by annotaligner SVs. The x axis represents the inverse. Sample pairs are colored and shaped by chromosome Each grid represents a different subset of pairwise match distances.

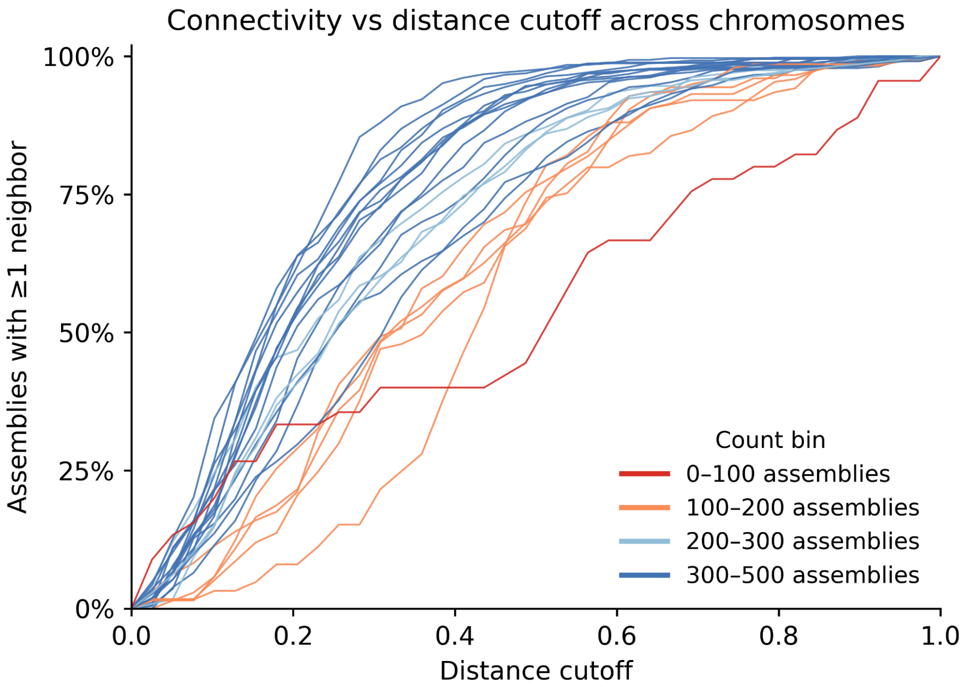


**Supplementary Figure 34. Connectivity of haplotypes in Centrolign graphs**

Pairwise match distance (x axis) versus the percent of haplotypes with at least one neighbor haplotype at or below the pairwise match distance value (y axis). Each line represents one chromosome, and chromosomes are colored by the number of assemblies for that chromosome present in the analysis.

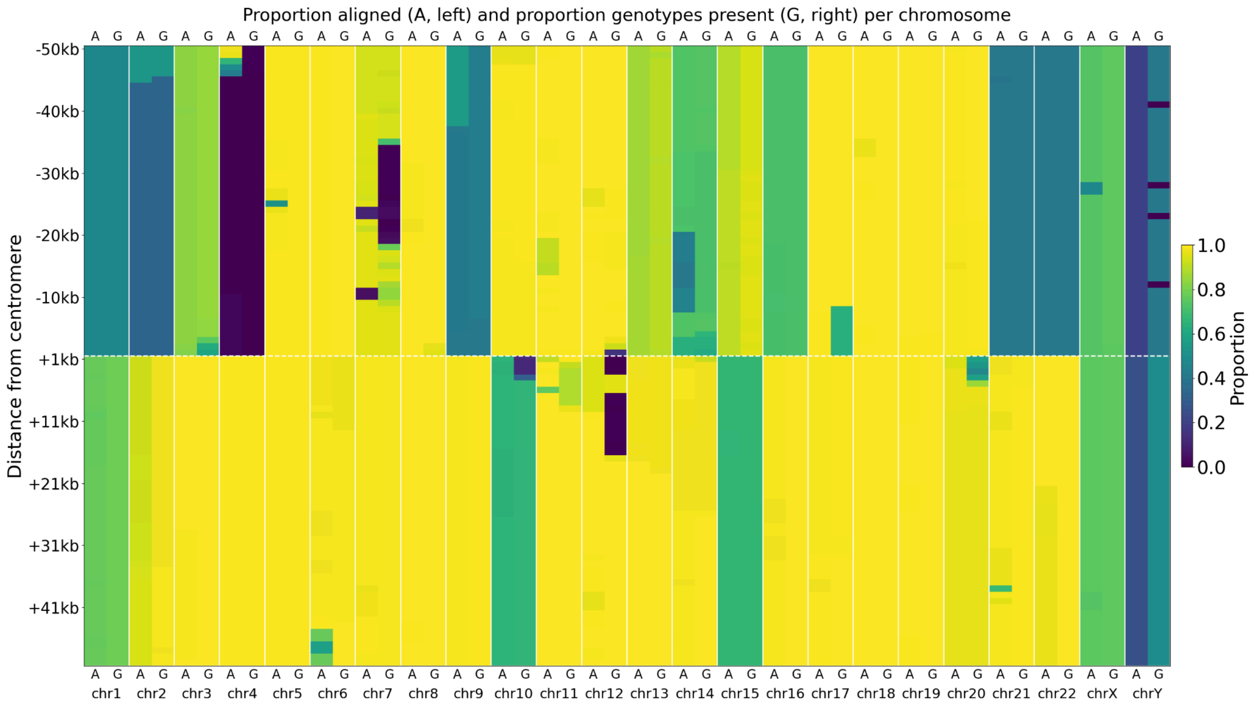


**Supplementary Figure 35. Genotype missingness of HPRC2 MC vcf in regions flanking the active alpha satellite arrays.**

Heatmap indicating the coverage and genotype missingness of the HPRC2 minigraph-cactus (MC) CHM13 graph for 50 kbp upstream (negative) and downstream (positive) of the CHM13 alpha satellite arrays. Columns labelled “A” represent the proportion of haplotypes in the graph aligned to CHM13 in each 1 kbp window. Columns labelled “G” represent the proportion of genotypes from the MC vcf that are present in each 1 kbp window. Each chromosome’s “A” and “G” columns are separated by a white line and labelled.

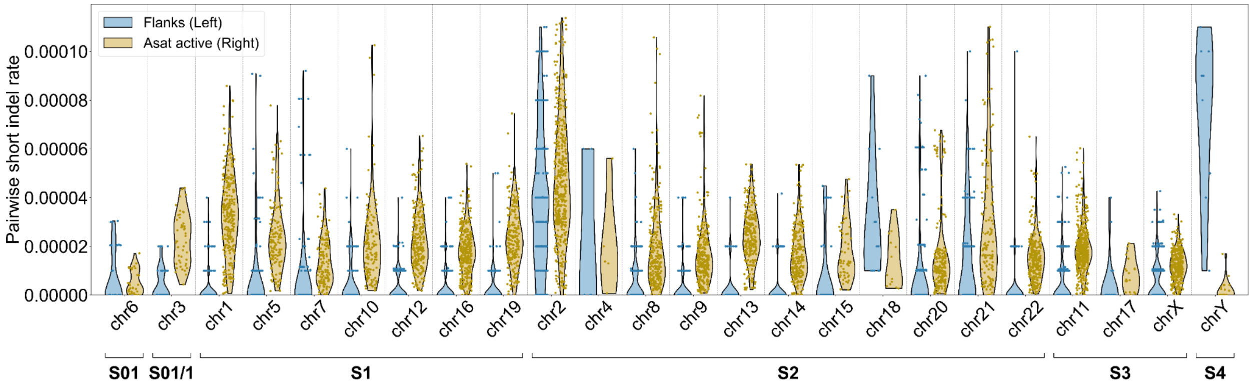


**Supplementary Figure 36. Pairwise short indel rate in active alpha vs immediate flanks**

Pairwise short indel rate in the 50 kbp flanking regions (blue, leftmost violin) and in the alpha satellite arrays (yellow, rightmost violin). Chromosomes are grouped by their suprachromosal family.

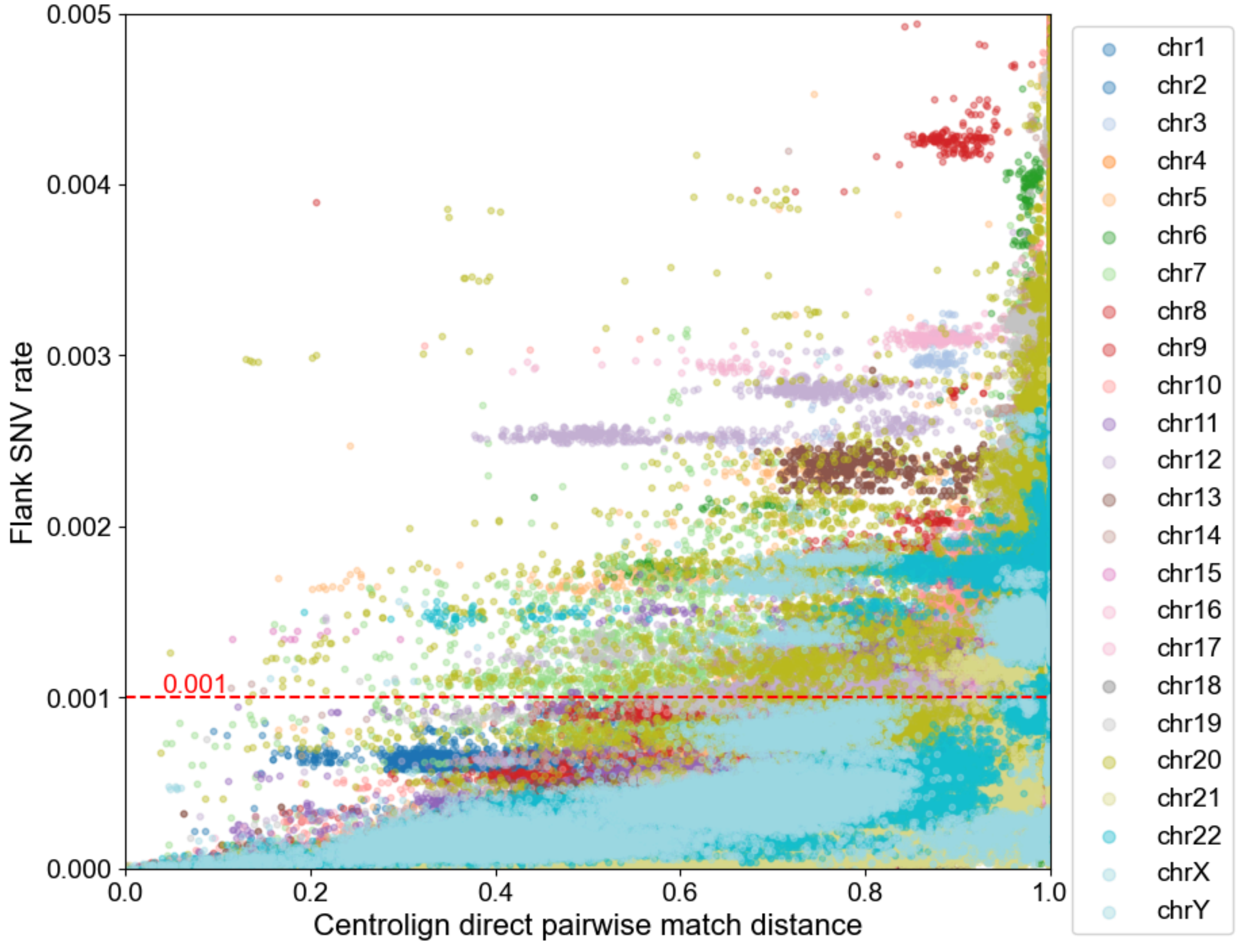


**Supplementary Figure 37. SNV rate in regions flanking alpha satellites for all samples in HPRC2**

Comparison of the SNV rate from the 50 kbp regions flanking the alpha satellite arrays derived from the MC vcf (y axis) and the Centrolign direct pairwise match distances for the alpha satellite array. The average rate value for each sample pair is plotted and colored by chromosome. The red dashed line indicates the expected genome wide SNV rate of 1/1000 bp.

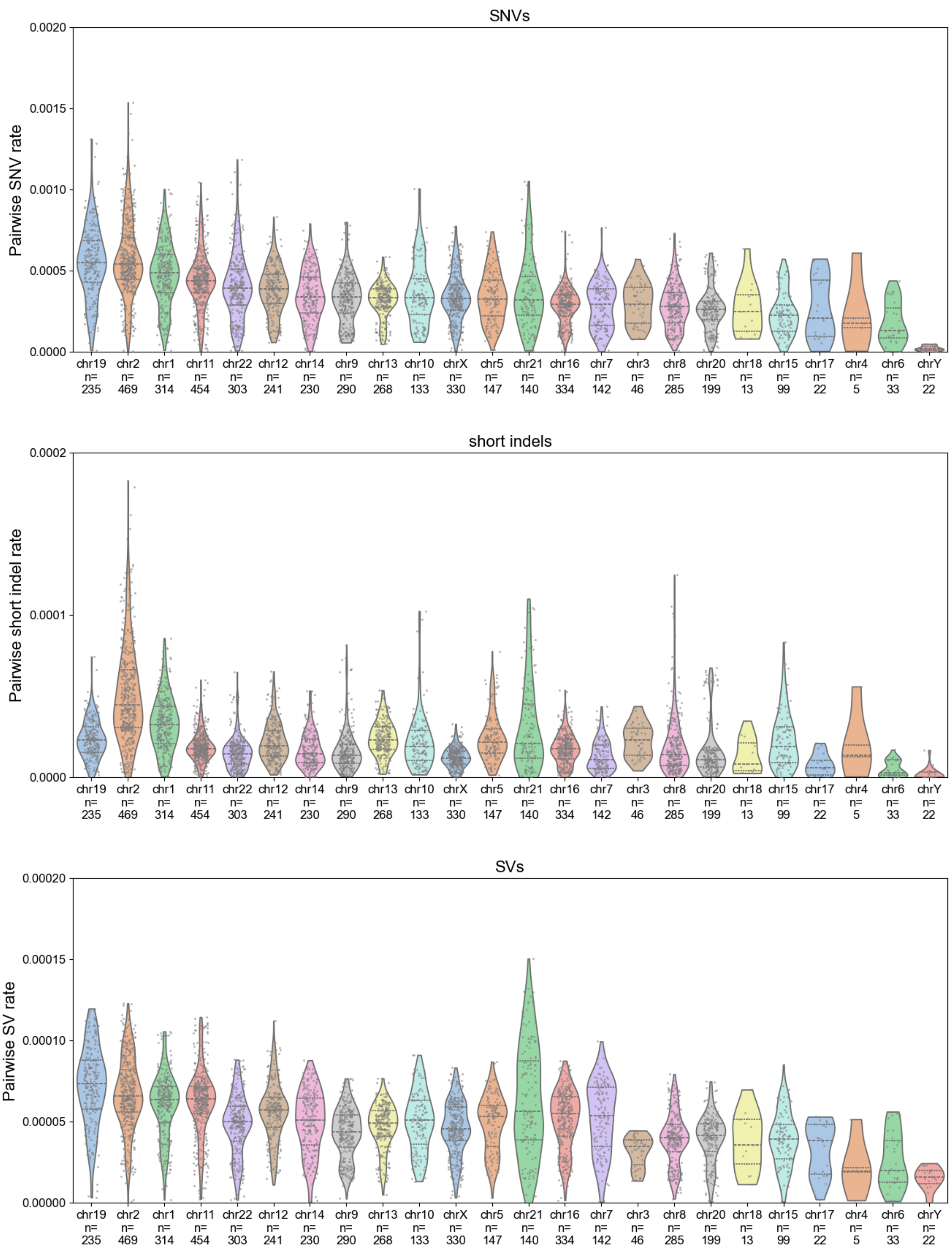
SNVs
Pairwise SNV rate
0.0020
0.0015
0.0010
0.0005
0.0000
short indels
Pairwise short indel rate
0.0002
0.0001
0.0000
SVs
Pairwise SV rate
0.00020
0.00015
0.00010
0.00005
0.00000
chr19 n= 235
chr2 n= 469
chr1 n= 314
chr11 n= 454
chr22 n= 303
chr12 n= 241
chr14 n= 230
chr9 n= 290
chr13 n= 268
chr10 n= 133
chrX n= 330
chr5 n= 147
chr21 n= 140
chr16 n= 334
chr7 n= 142
chr3 n= 46
chr8 n= 285
chr20 n= 199
chr18 n= 13
chr15 n= 99
chr17 n= 22
chr4 n= 5
chr6 n= 33
chrY n= 22

**Supplementary Figure 38. Per-sample pair variation rate distributions across chromosomes.**

All panels are sorted by the SNV variation rate. The number of sample pairs per chromosome is labelled on the x axis. SNV variation rate is calculated as (# SNVs / aligned bases). Short indel variation rate is calculated as (# short indels / aligned bases). SV variation rate is calculated as (# SVs / average length of the two arrays).

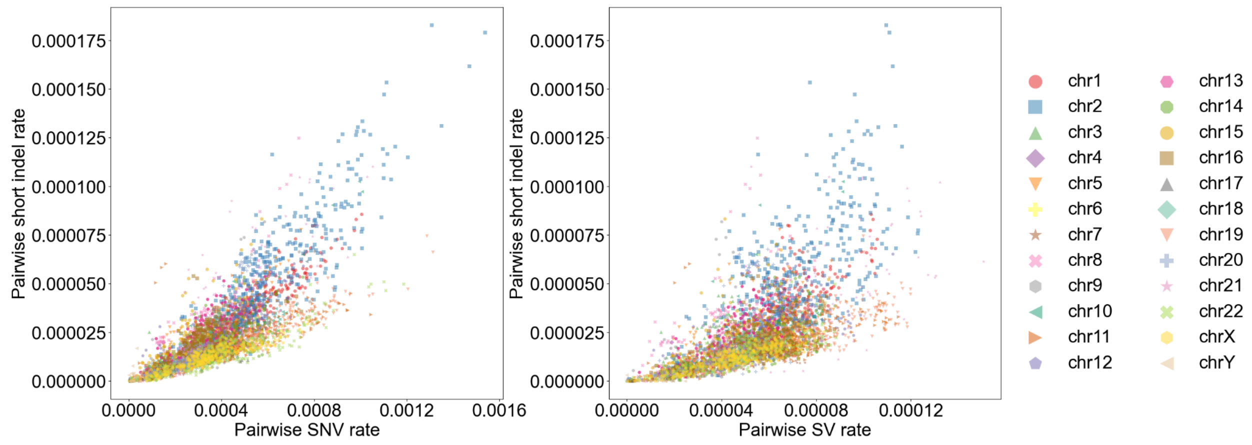


**Supplementary Figure 39. Centrolign pairwise short indel rate vs SV rate**

Pairwise short indel rate (y axes) vs SNV rate (x axis, left) and SV rate (x axis, right) for all chromosomes. Plot only shows sample pairs with < 0.2 pairwise match distance.

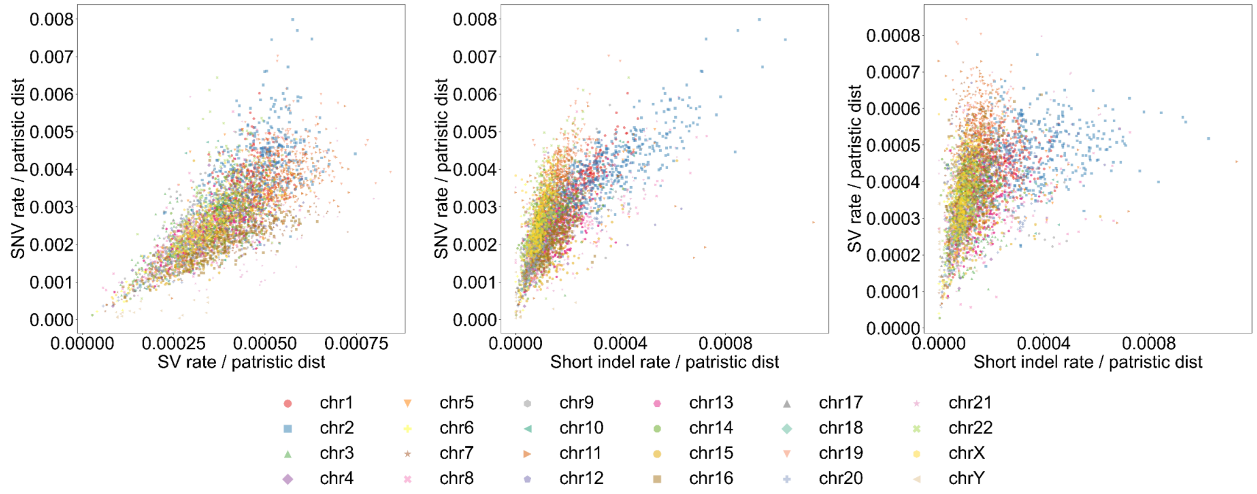


**Supplementary Figure 40. Linear relationship between variation rates corrected by patristic distance**

Pairwise short indel rate (y axes) vs SNV rate (x axis, left) and SV rate (x axis, right) for all chromosomes. Plot only shows sample pairs with < 0.2 pairwise match distance.

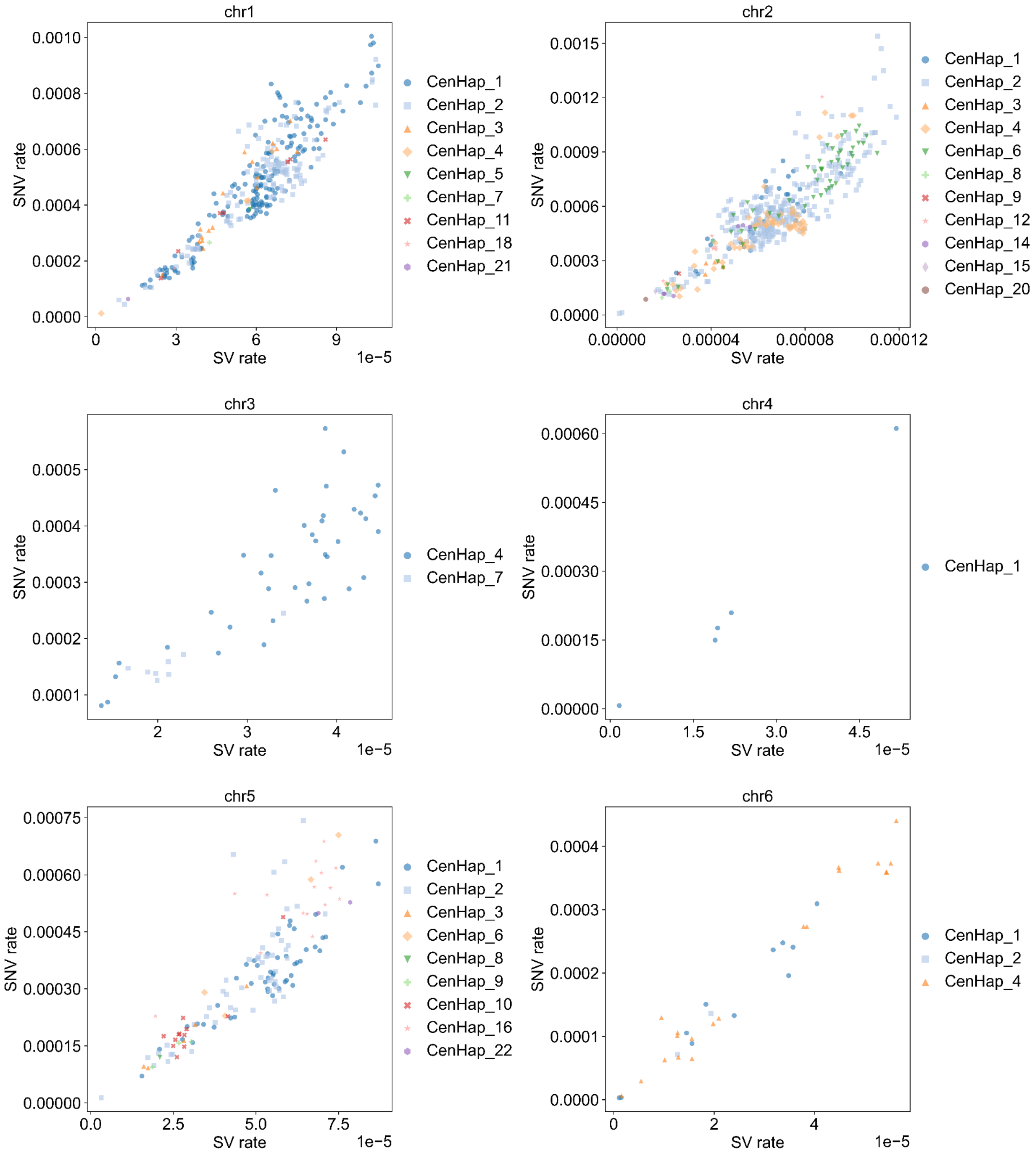
chr1
SNV rate
0.0010
0.0008
0.0006
0.0004
0.0002
0.0000
0
3
6
9
SV rate
1e−5
CenHap_1
CenHap_2
CenHap_3
CenHap_4
CenHap_5
CenHap_7
CenHap_11
CenHap_18
CenHap_21
chr2
SNV rate
0.0015
0.0012
0.0009
0.0006
0.0003
0.0000
0.00000
0.00004
0.00008
0.00012
SV rate
CenHap_1
CenHap_2
CenHap_3
CenHap_4
CenHap_6
CenHap_8
CenHap_9
CenHap_12
CenHap_14
CenHap_15
CenHap_20
chr3
SNV rate
0.0005
0.0004
0.0003
0.0002
0.0001
2
3
4
SV rate
1e−5
CenHap_4
CenHap_7
chr4
SNV rate
0.00060
0.00045
0.00030
0.00015
0.00000
0.0
1.5
3.0
4.5
SV rate
1e−5
CenHap_1
chr5
SNV rate
0.00075
0.00060
0.00045
0.00030
0.00015
0.00000
0.0
2.5
5.0
7.5
SV rate
1e−5
CenHap_1
CenHap_2
CenHap_3
CenHap_6
CenHap_8
CenHap_9
CenHap_10
CenHap_16
CenHap_22
chr6
SNV rate
0.0004
0.0003
0.0002
0.0001
0.0000
0
2
4
SV rate
1e−5
CenHap_1
CenHap_2
CenHap_4

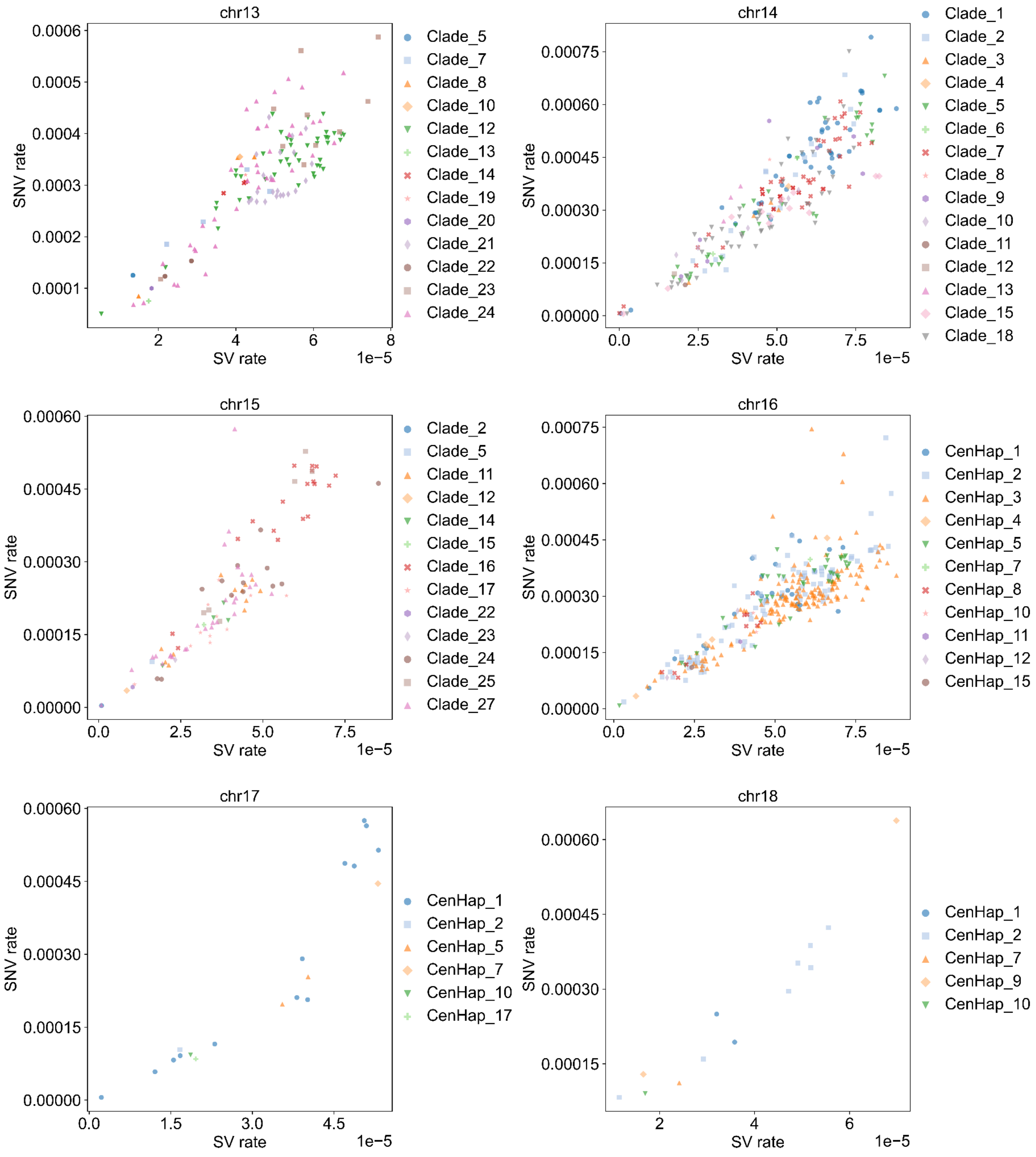
chr13
SNV rate
SV rate
1e−5
0.0001
0.0002
0.0003
0.0004
0.0005
0.0006
2
4
6
8
Clade_5
Clade_7
Clade_8
Clade_10
Clade_12
Clade_13
Clade_14
Clade_19
Clade_20
Clade_21
Clade_22
Clade_23
Clade_24
chr14
SNV rate
SV rate
1e−5
0.00000
0.00015
0.00030
0.00045
0.00060
0.00075
0.0
2.5
5.0
7.5
Clade_1
Clade_2
Clade_3
Clade_4
Clade_5
Clade_6
Clade_7
Clade_8
Clade_9
Clade_10
Clade_11
Clade_12
Clade_13
Clade_15
Clade_18
chr15
SNV rate
SV rate
1e−5
0.00000
0.00015
0.00030
0.00045
0.00060
0.0
2.5
5.0
7.5
Clade_2
Clade_5
Clade_11
Clade_12
Clade_14
Clade_15
Clade_16
Clade_17
Clade_22
Clade_23
Clade_24
Clade_25
Clade_27
chr16
SNV rate
SV rate
1e−5
0.00000
0.00015
0.00030
0.00045
0.00060
0.00075
0.0
2.5
5.0
7.5
CenHap_1
CenHap_2
CenHap_3
CenHap_4
CenHap_5
CenHap_7
CenHap_8
CenHap_10
CenHap_11
CenHap_12
CenHap_15
chr17
SNV rate
SV rate
1e−5
0.00000
0.00015
0.00030
0.00045
0.00060
0.0
1.5
3.0
4.5
CenHap_1
CenHap_2
CenHap_5
CenHap_7
CenHap_10
CenHap_17
chr18
SNV rate
SV rate
1e−5
0.00015
0.00030
0.00045
0.00060
2
4
6
CenHap_1
CenHap_2
CenHap_7
CenHap_9
CenHap_10

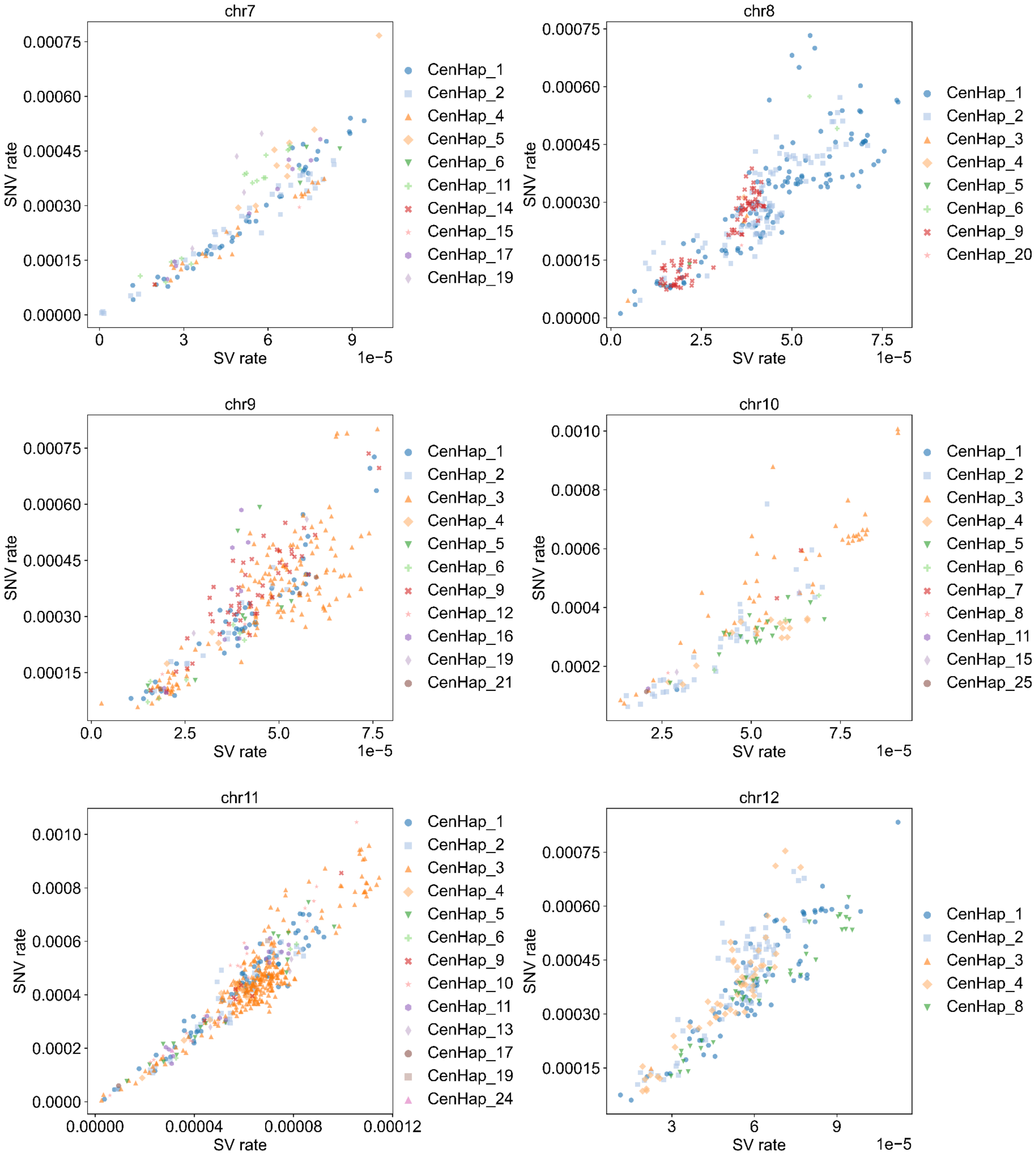

chr7
SNV rate
0.00000
0.00015
0.00030
0.00045
0.00060
0.00075
SV rate
0
3
6
9
1e−5
CenHap_1
CenHap_2
CenHap_4
CenHap_5
CenHap_6
CenHap_11
CenHap_14
CenHap_15
CenHap_17
CenHap_19
chr8
SNV rate
0.00000
0.00015
0.00030
0.00045
0.00060
0.00075
SV rate
0.0
2.5
5.0
7.5
1e−5
CenHap_1
CenHap_2
CenHap_3
CenHap_4
CenHap_5
CenHap_6
CenHap_9
CenHap_20
chr9
SNV rate
0.00015
0.00030
0.00045
0.00060
0.00075
SV rate
0.0
2.5
5.0
7.5
1e−5
CenHap_1
CenHap_2
CenHap_3
CenHap_4
CenHap_5
CenHap_6
CenHap_9
CenHap_12
CenHap_16
CenHap_19
CenHap_21
chr10
SNV rate
0.0002
0.0004
0.0006
0.0008
0.0010
SV rate
2.5
5.0
7.5
1e−5
CenHap_1
CenHap_2
CenHap_3
CenHap_4
CenHap_5
CenHap_6
CenHap_7
CenHap_8
CenHap_11
CenHap_15
CenHap_25
chr11
SNV rate
0.0000
0.0002
0.0004
0.0006
0.0008
0.0010
SV rate
0.00000
0.00004
0.00008
0.00012
CenHap_1
CenHap_2
CenHap_3
CenHap_4
CenHap_5
CenHap_6
CenHap_9
CenHap_10
CenHap_11
CenHap_13
CenHap_17
CenHap_19
CenHap_24
chr12
SNV rate
0.00015
0.00030
0.00045
0.00060
0.00075
SV rate
3
6
9
1e−5
CenHap_1
CenHap_2
CenHap_3
CenHap_4
CenHap_8

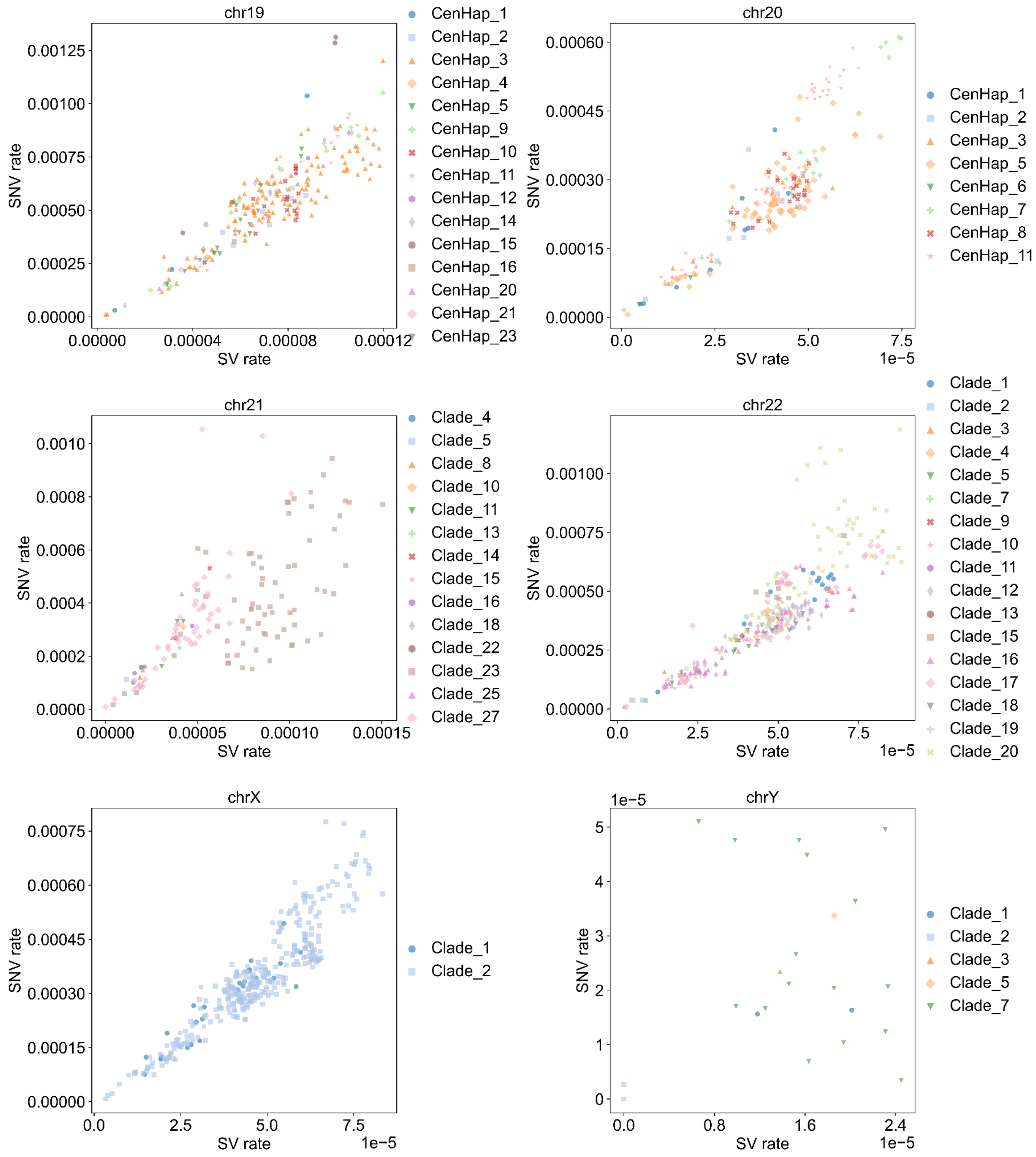


**Supplementary Figure 41. SNV rate vs SV rate colored by cenhap clade**

Per sample pair SNV (y axis) vs SV (x axis) variation rates. Each sample pair is colored and shaped by clade or cenhap (where available). Clades are defined from the input guide trees to Centrolign, where 95% of sample pairs in a clade must have at least an pairwise match distance of 0.8.

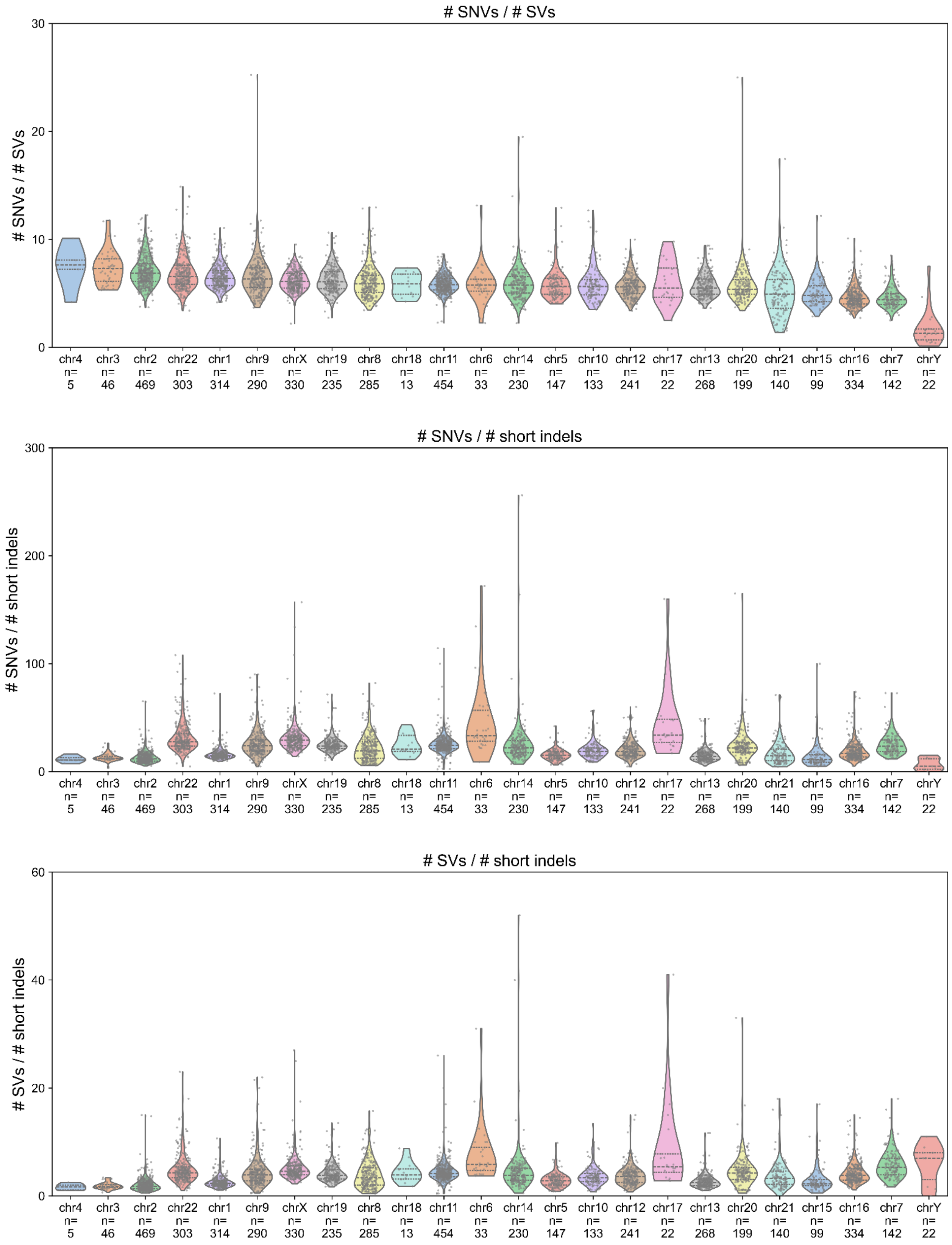
### SNVs / # SVs
### SNVs / # SVs
0
10
20
30
chr4 n= 5
chr3 n= 46
chr2 n= 469
chr22 n= 303
chr1 n= 314
chr9 n= 290
chrX n= 330
chr19 n= 235
chr8 n= 285
chr18 n= 13
chr11 n= 454
chr6 n= 33
chr14 n= 230
chr5 n= 147
chr10 n= 133
chr12 n= 241
chr17 n= 22
chr13 n= 268
chr20 n= 199
chr21 n= 140
chr15 n= 99
chr16 n= 334
chr7 n= 142
chrY n= 22
### SNVs / # short indels
### SNVs / # short indels
0
100
200
300
chr4 n= 5
chr3 n= 46
chr2 n= 469
chr22 n= 303
chr1 n= 314
chr9 n= 290
chrX n= 330
chr19 n= 235
chr8 n= 285
chr18 n= 13
chr11 n= 454
chr6 n= 33
chr14 n= 230
chr5 n= 147
chr10 n= 133
chr12 n= 241
chr17 n= 22
chr13 n= 268
chr20 n= 199
chr21 n= 140
chr15 n= 99
chr16 n= 334
chr7 n= 142
chrY n= 22
### SVs / # short indels
### SVs / # short indels
0
20
40
60
chr4 n= 5
chr3 n= 46
chr2 n= 469
chr22 n= 303
chr1 n= 314
chr9 n= 290
chrX n= 330
chr19 n= 235
chr8 n= 285
chr18 n= 13
chr11 n= 454
chr6 n= 33
chr14 n= 230
chr5 n= 147
chr10 n= 133
chr12 n= 241
chr17 n= 22
chr13 n= 268
chr20 n= 199
chr21 n= 140
chr15 n= 99
chr16 n= 334
chr7 n= 142
chrY n= 22

**Supplementary Figure 42. Balance of SVs, short indels, and SNVs.**
Per-sample pairwise # of SNVs / # SVs, # of SNVs / # short indels (bottom panel), and # of SVs / # short indels, indicating the balance between mutation types across chromosomes.

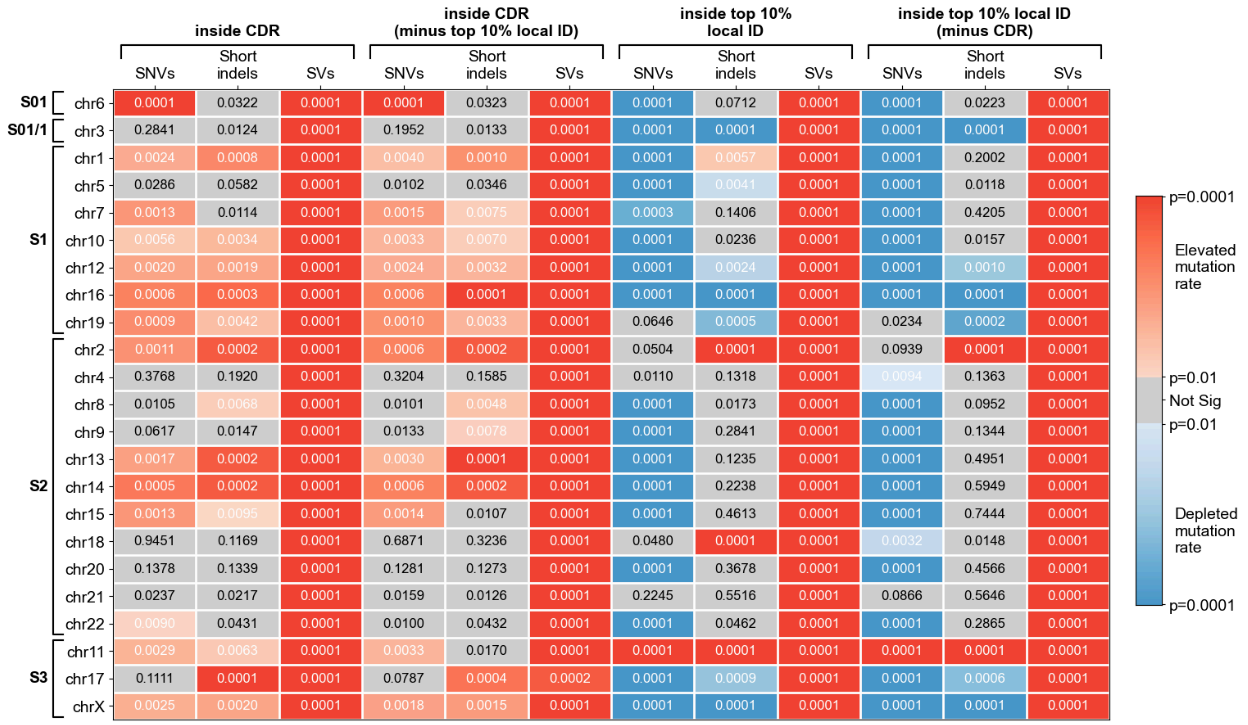


**Supplementary Figure 43. Summary of permutation test results for variation rate against spatial features of the array**
Summary of permutation test results. The left three columns (labelled "inside CDR") indicate permutation tests for comparing the variation rate inside the CDR against the rest of the array. The next three columns ("inside CDR minus top 10% local ID") indicate permutation tests for variation rate inside the CDR subtracting out any regions from the top 10% of local identity that fall inside the CDR. The columns labelled "inside top 10% local ID" indicate permutation tests run comparing the variation rate inside the highest 10% of local identity windows with the rest of the array. The rightmost columns labelled "inside top 10% local ID minus CDR" indicate a permutation test comparing the variation rate for the top 10% of local identity windows, but subtracting out any overlapping CDR regions. Each box indicates the p-value for a given mutation type and chromosome, shaded by the level of significance relative to the ceiling of .001 for N=10,000 reps.

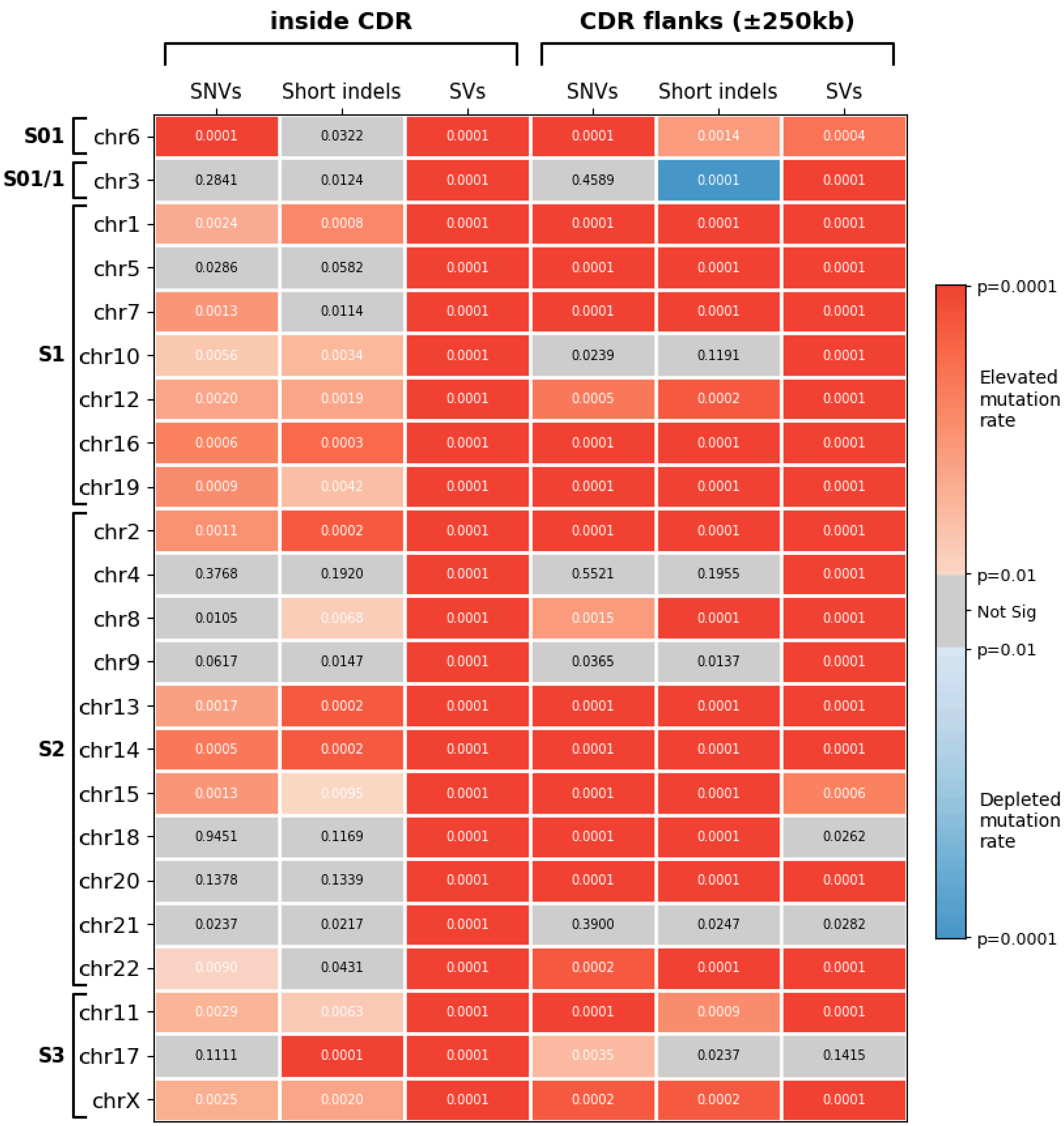


**Supplementary Figure 44. Summary of permutation results comparing inside the CDR with CDR flanks**

Summary of permutation results for variation rates inside the CDR (left three) and 250 kbp flanking the CDR (right three). Each box indicates the p-value for a given mutation type and chromosome, shaded by the level of significance relative to the ceiling of .001 for N=10,000 reps.

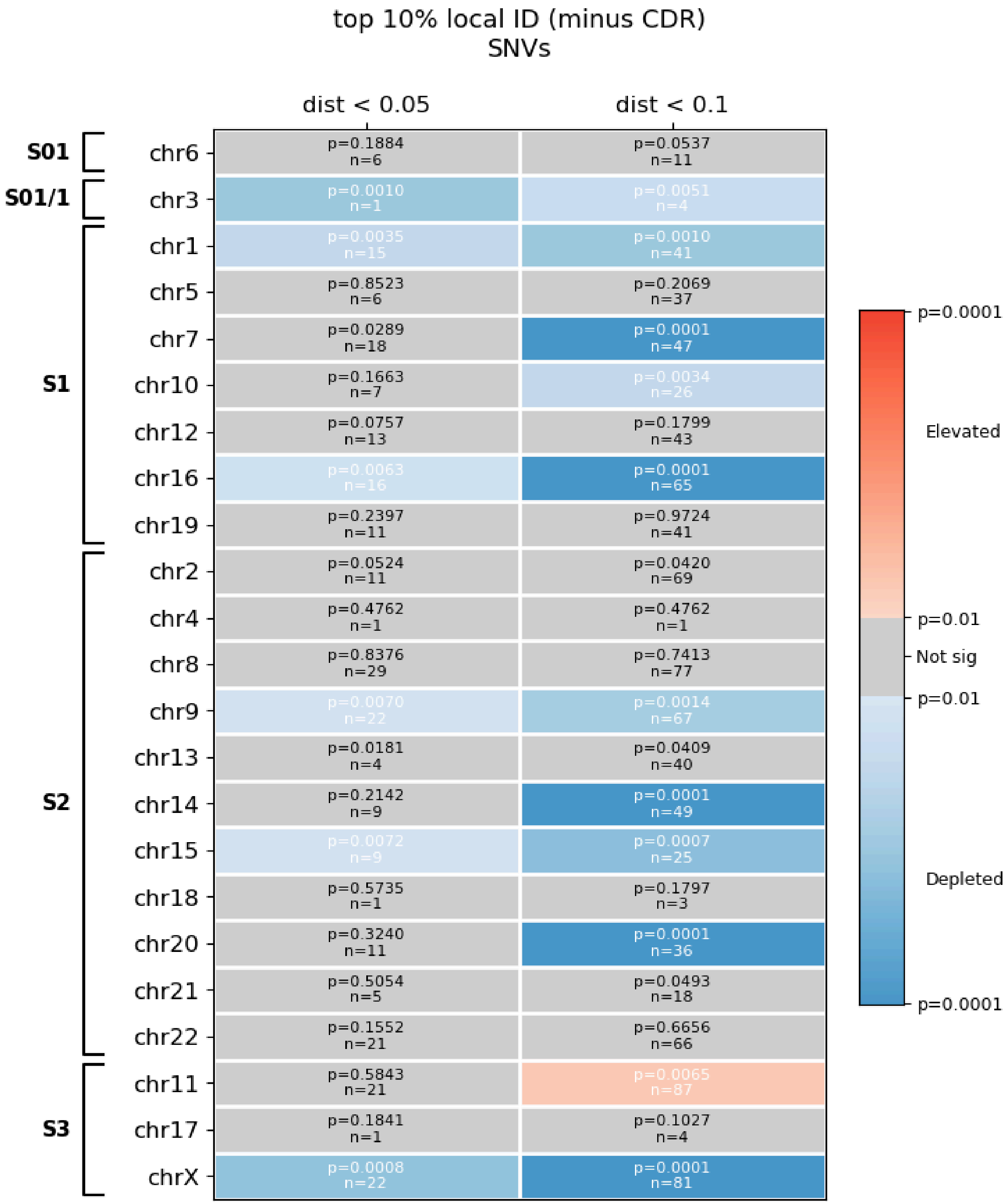


**Supplementary Figure 45. Summary of permutation test results for local identity with close distances**

Summary of permutation test results for local identity with close distances (pairwise match distance < 0.05 on the left, < 0.1 on the right) with n= number of sample pairs in each comparison passing that distance threshold. Each box indicates the p-value for a given mutation type and chromosome, shaded by the level of significance relative to the ceiling of .001 for N=10,000 reps.

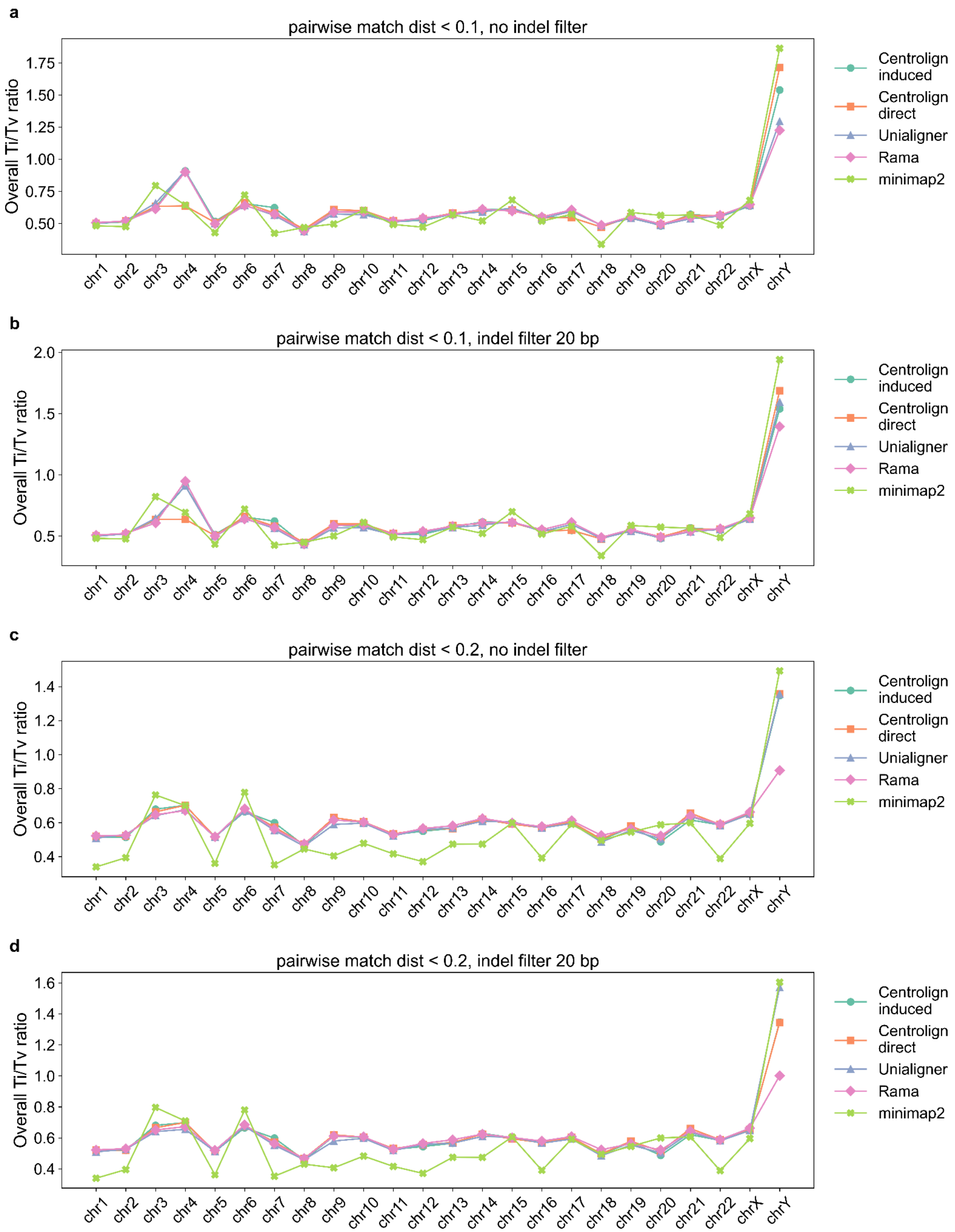
a
pairwise match dist < 0.1, no indel filter
b
pairwise match dist < 0.1, indel filter 20 bp
c
pairwise match dist < 0.2, no indel filter
d
pairwise match dist < 0.2, indel filter 20 bp
Overall Ti/Tv ratio
chr1 chr2 chr3 chr4 chr5 chr6 chr7 chr8 chr9 chr10 chr11 chr12 chr13 chr14 chr15 chr16 chr17 chr18 chr19 chr20 chr21 chr22 chrX chrY
Centrolign induced
Centrolign direct
Unialigner
Rama
minimap2

**Supplementary Figure 46. Ti/Tv ratio from orthogonal aligners**
Comparison of pairwise Ti/Tv ratio calculated by Centrolign induced from the MSA (green, circle), Centrolign direct alignment mode (orange, square), Unialigner (purple, triangle), Rama (pink, diamond) and minimap2 (light green, cross). Each panel represents different filter criteria: **a,** Only sample pairs with pairwise match distance < 0.1. **b**, pairs with pairwise match distance < 0.1 AND removing SNVs within 20 bp of an indel. **c**, Only sample pairs with pairwise match distance < 0.2. **d,** Only sample pairs with pairwise match distance < 0.2 AND removing SNVs within 20 bp of an indel.

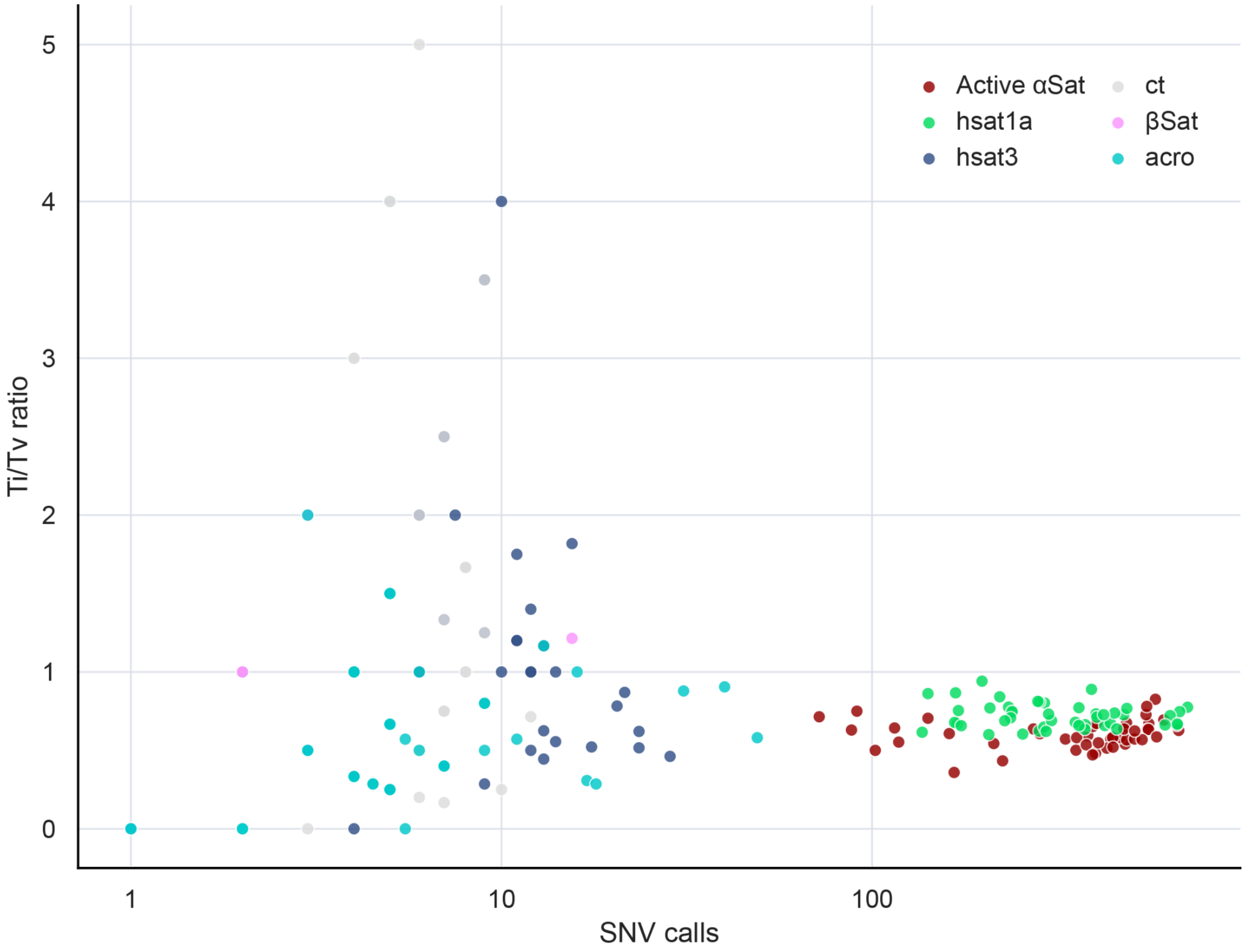


**Supplementary Figure 47. Ti/Tv ratio across satellite classes in chromosome 3's centromere**
Each point represents the transition-to-transversion (Ti/Tv) ratio for pairwise comparisons of satellite classes in between (and including) the active array on chromosome 3. Points are colored using colors from the CenSat satellite annotation. Arrays with Centrolign pairwise match distance < 0.2 for the entire region are shown. Only satellite classes that had arrays larger than 50 kbp are included. HSat3, ct, βSat, and acro regions have low variant counts and highly variable Ti/Tv ratios.

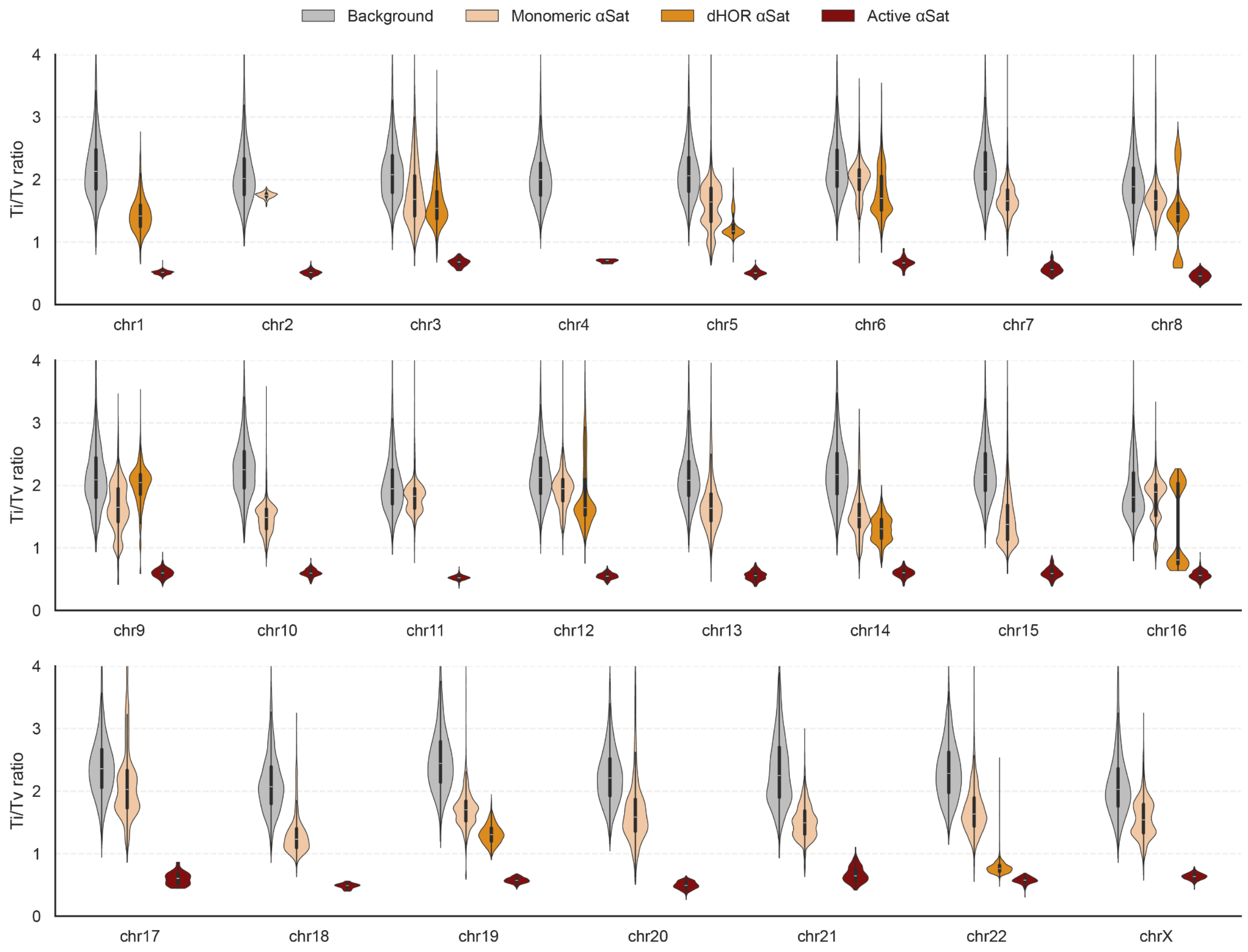


**Supplementary Figure 48. Ti/Tv ratios across chromosomes and sequence classes**

Violin plots show the distributions of pairwise transition-to-transversion ratios for background regions, monomeric α-satellite (αSat), divergent higher-order repeat (dHOR) αSat, and active αSat arrays. Active αSat values were calculated from Centrolign SNVs for closely related sequence pairs (distance <0.2). Background, monomeric αSat, and dHOR αSat values were calculated from biallelic SNVs from minigraph-cactus. For Minigraph-Cactus, sampling from all possible comparisons are shown in order to avoid the loss of comparisons with fewer than 50 differing SNVs. Violins are shown only where a given sequence class is present (some chromosomes lack monomeric αSat or dHOR αSat annotations meeting the region-size criteria). Comparisons with fewer than 50 SNV differences were excluded, and up to 5,000 observations were sampled per chromosome and sequence class. The bimodal dHOR distributions on chromosomes 8 and 16 may reflect comparisons against divergent haplotypes, and in the case of chromosome 16 different distributions for two different dHOR intervals. Internal boxes indicate the median and interquartile range. Chromosome Y was excluded.

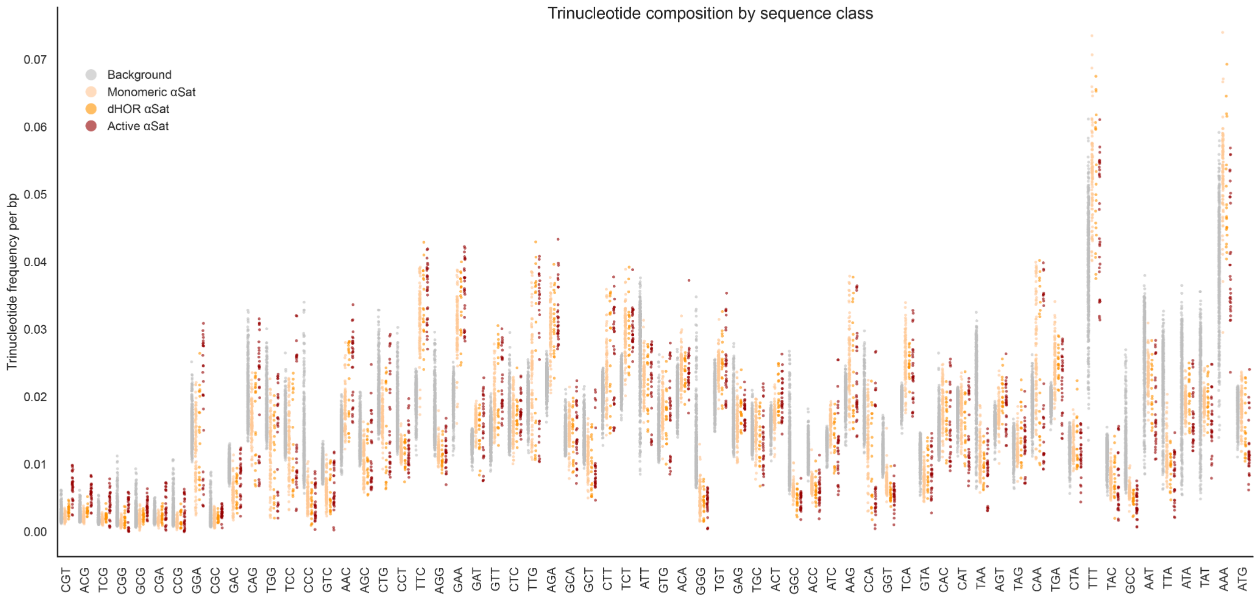


**Supplementary Figure 49. Triplet composition of active arrays differs from diverged αSat as well as background regions**

Frequencies of trinucleotides (64 total) were calculated from CHM13 sequence for background regions, monomeric α-Sat, divergent higher-order repeat (dHOR) αSat, and active αSat arrays. Each point represents the frequency (count divided by region length) of a trinucleotide within one genomic region. Up to 500 regions per sequence class and trinucleotide were displayed. The x axis is ordered by the enrichment in active arrays relative to dHOR αSat. Analyses include chromosomes 1–22 and X.

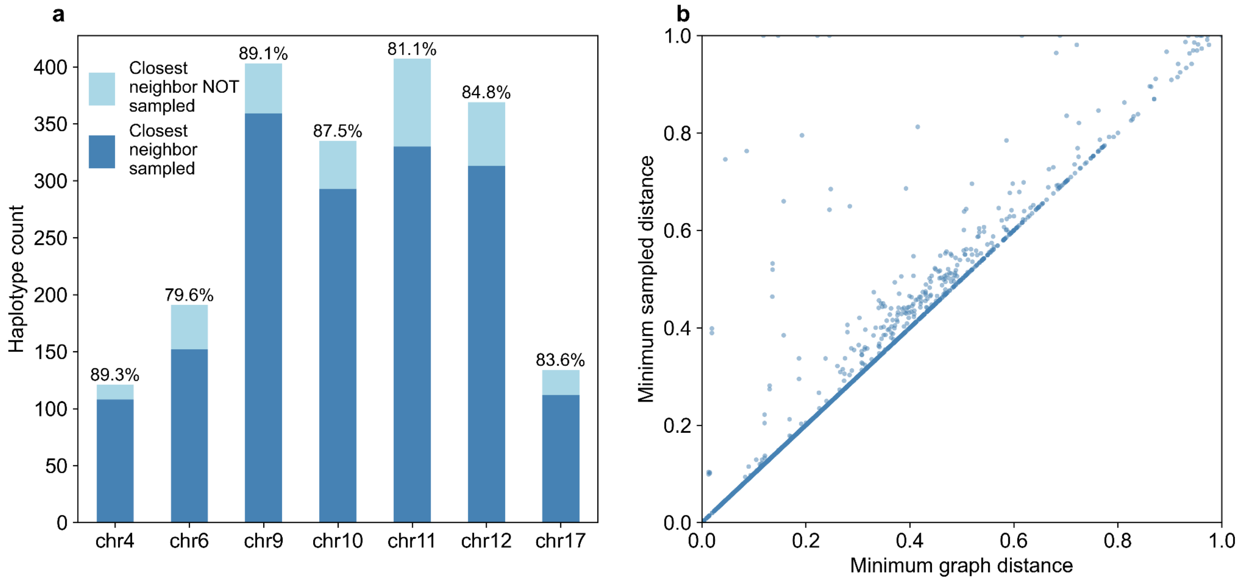


**Supplementary Figure 50: k-mer based detection of most closely related haplotype. a.** Count of haploid samples where haplotype sampling selected the closest neighbor (by pairwise match distance) in the Centrolign graph is shown in dark blue, while the count of haploid samples where the closest neighbor was not selected in the personalized graph is shown in light blue. Percentages on the top of each bar indicate the percent of haploid samples whose closest neighbor was selected. **b**. For each read set, minimum pairwise match distance to any haplotype in the graph is on X axis, and minimum pairwise match distance to a haplotype in the personalized graph is on Y axis.

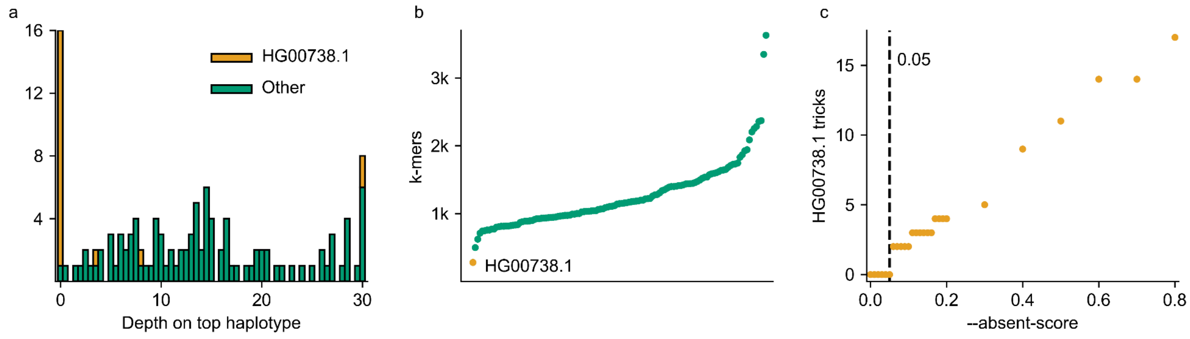


**Supplementary Figure 51: Adjusting the --absent-score parameter**. **a,** Read alignment depth on nodes private to the top haplotype selected by the sampler. This first haplotype is the one used for cenhap typing and is thus of particular interest. Average depth in bins of 0.5 on the X axis, with depths over 30 combined into the depth 30 bin. Count of tested Chromosome 4 haplotypes on the Y axis. Histogram bins split by whether the top haplotype is HG00738.1. **b,** Number of graph-unique k-mers present in each Chromosome 4 haplotype. Haplotypes sorted by count on X axis, k-mer count on Y axis. HG00738.1 is the large red dot. **c,** How often the haplotype sampler erroneously selects HG00738.1 as the top sampled haplotype for reads from a different cenhap. Varying values of --absent-score on X axis, count of erroneous selections on Y axis.

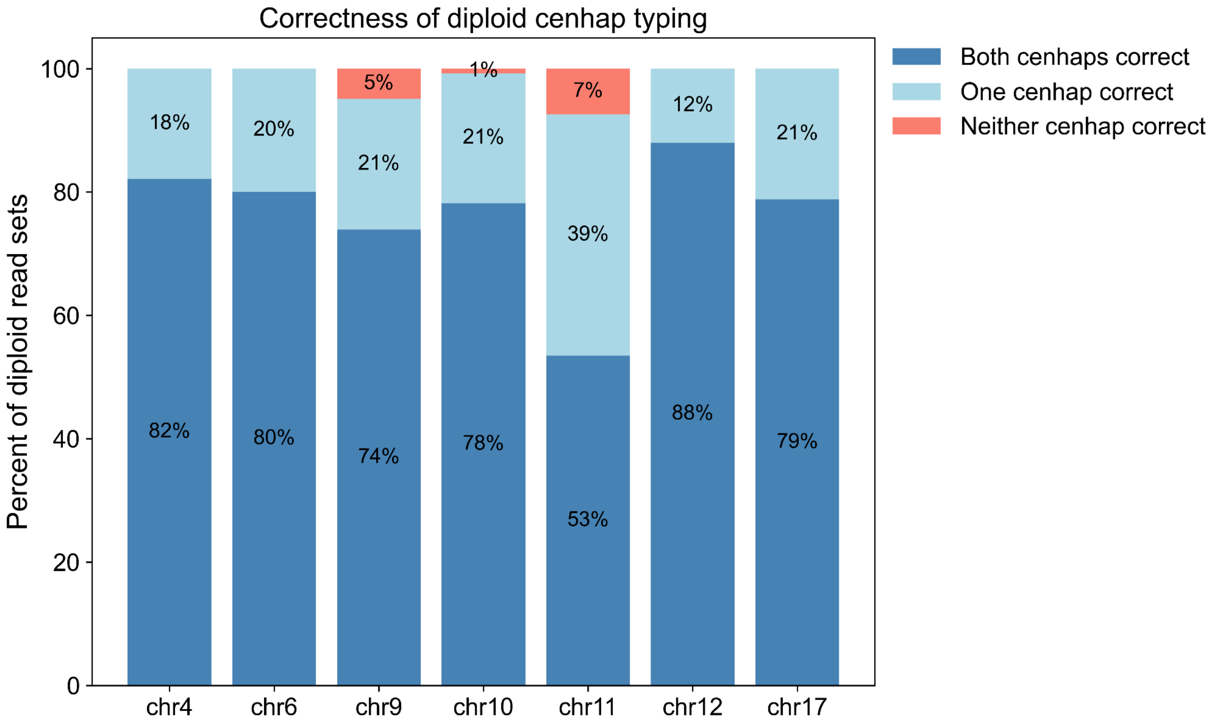


**Supplementary Figure 52: Cenhap typing on diploid read sets**. For each chromosome, the percentage of diploid read sets where both cenhaps were correctly typed (dark blue, bottom), one cenhap was correctly typed (light blue, middle) or neither cenhap was correctly typed (top, pink).

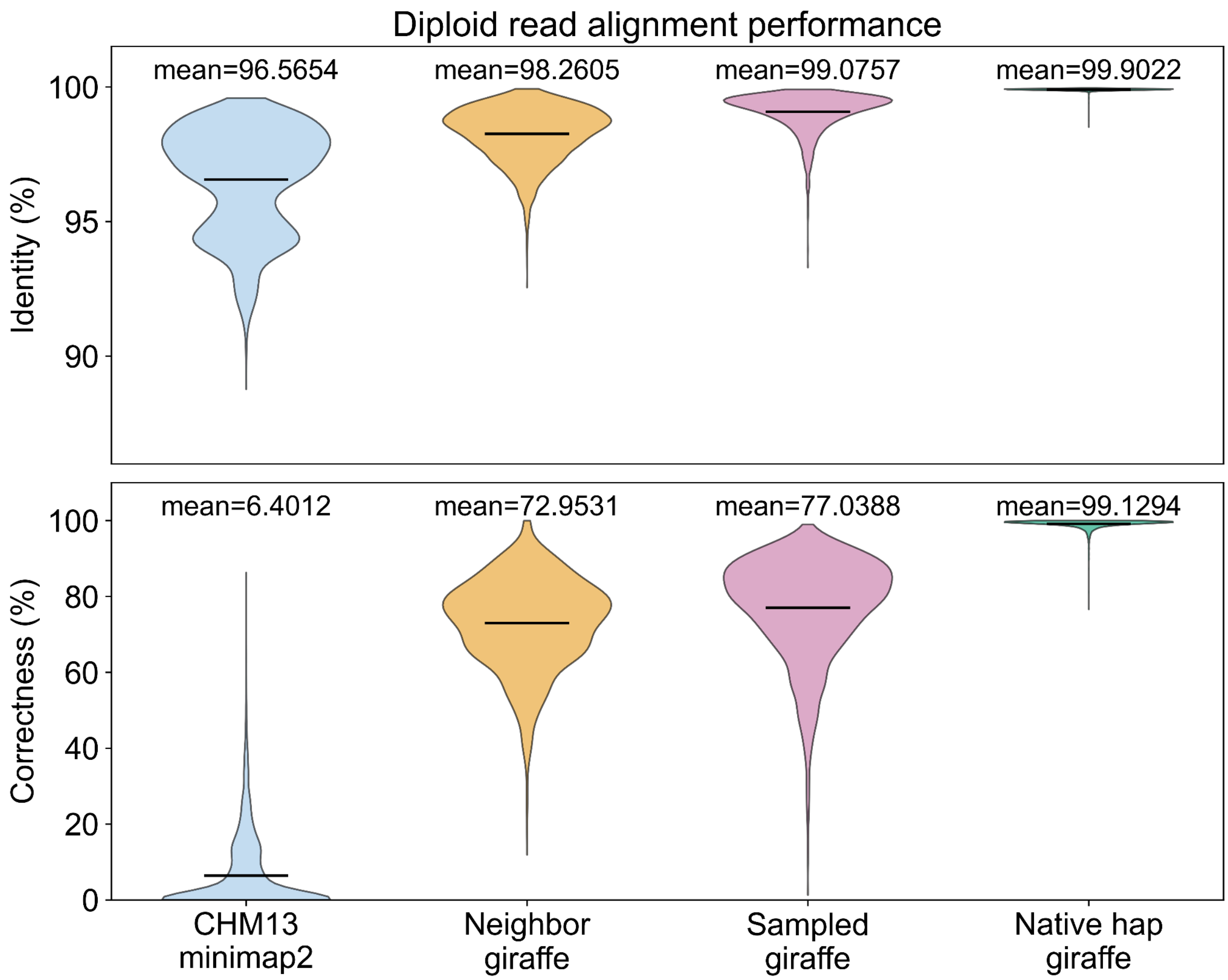


**Supplementary Figure 53: Diploid read alignment results.** Top, average read alignment identity and bottom, average read correctness, using real reads, for alignments to CHM13 (far left), a graph subset to the closest relation for each sample haplotype (middle left), a personalized graph reference (middle right), and a graph subset to the sample's own two haplotypes (far right), split by chromosome. Black bars indicate means. Alignments to linear references done with Minimap2, alignments to graph references done with Giraffe.

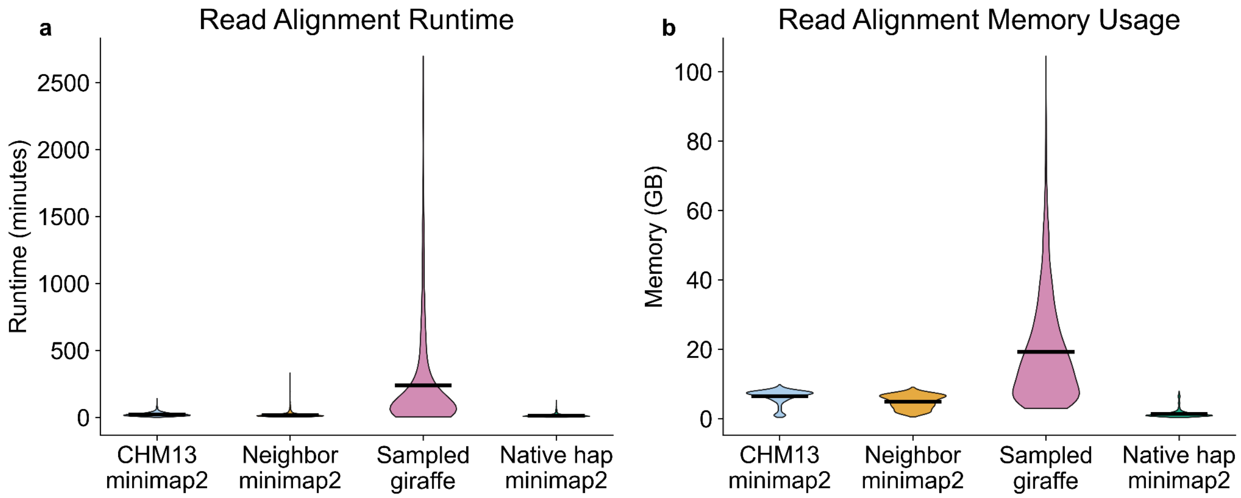


**Supplementary Figure 54: Computational resource usage of read alignments**. **a**, Runtime and **b**, memory usage of the read alignment step for alignments to CHM13, the most closely related haplotype, a personalized graph reference, and the reads' own assembly. Black bars indicate means. Alignments to linear references done with Minimap2. Alignments to graph references done with Giraffe.

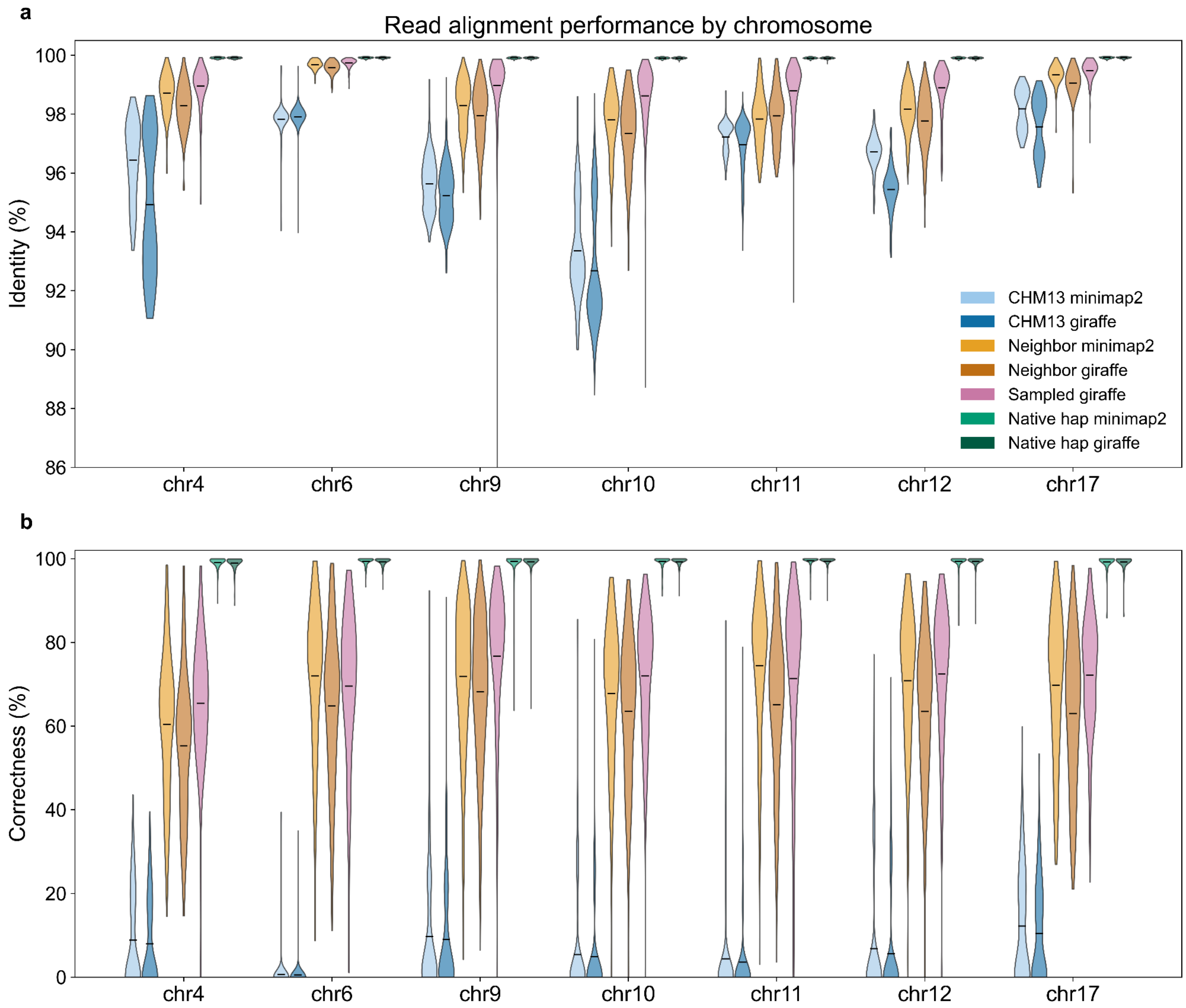


**Supplementary Figure 55: Read alignment results by chromosome. a**, Average read alignment identity and **b**, average read correctness, using real reads, for alignments to CHM13 (far left), the most closely related haplotype (middle left), a personalized graph reference (middle right), and the reads' own assembly (far right), split by chromosome. Black bars indicate means. Alignments to linear references done with Minimap2, alignments to graph references done with Giraffe.

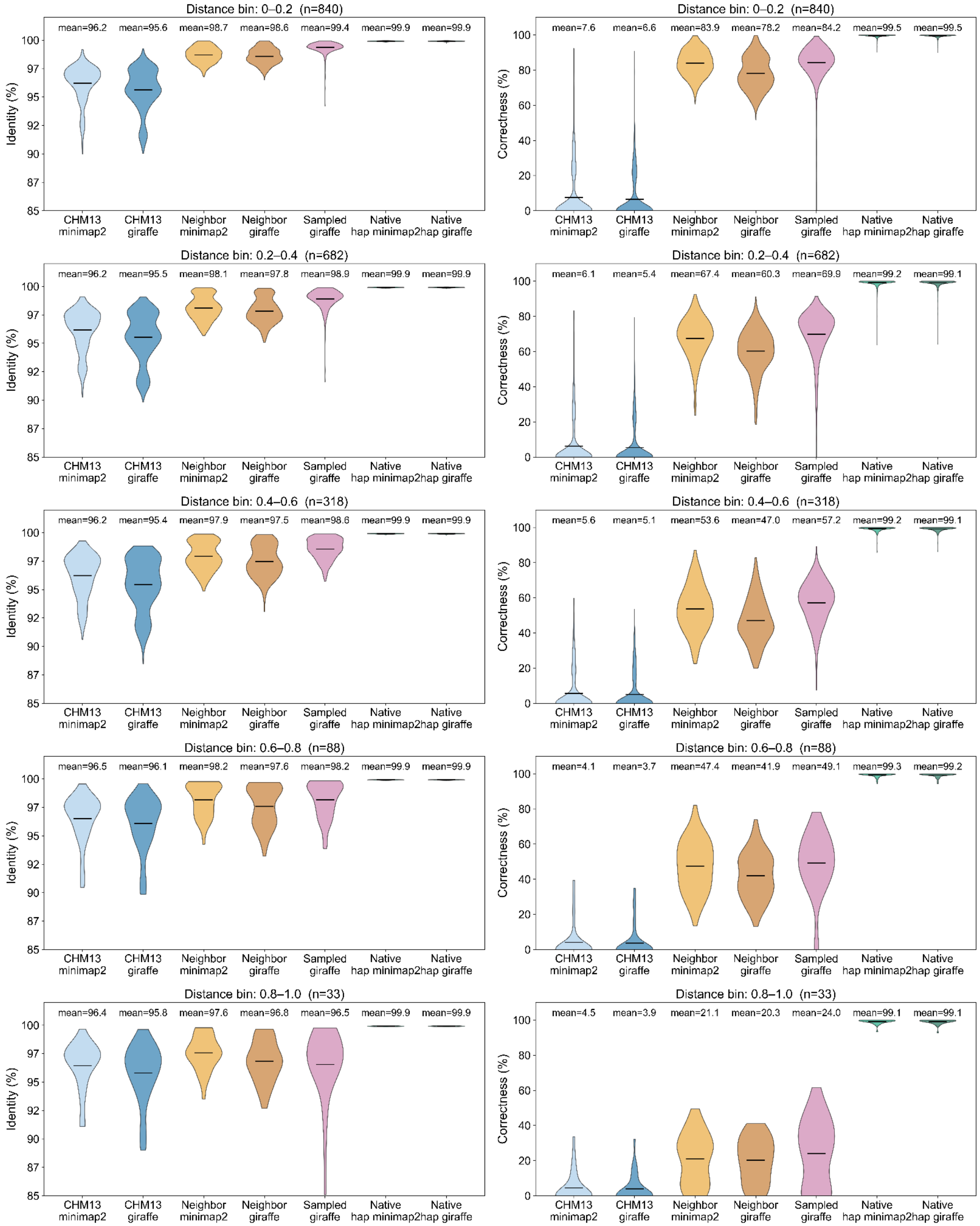


**Supplementary Figure 56: Read alignment results by minimum pairwise match distance.** Average read alignment identity (left column) and average read correctness (right column), using real reads, for alignments to CHM13 (far left), the most closely related haplotype (middle left), a personalized graph reference (middle right), and the reads' own assembly (far right), split by minimum pairwise match distance to a nearest neighbor assembly in the graph. Black bars indicate means. Alignments to linear references done with Minimap2, alignments to graph references done with Giraffe.

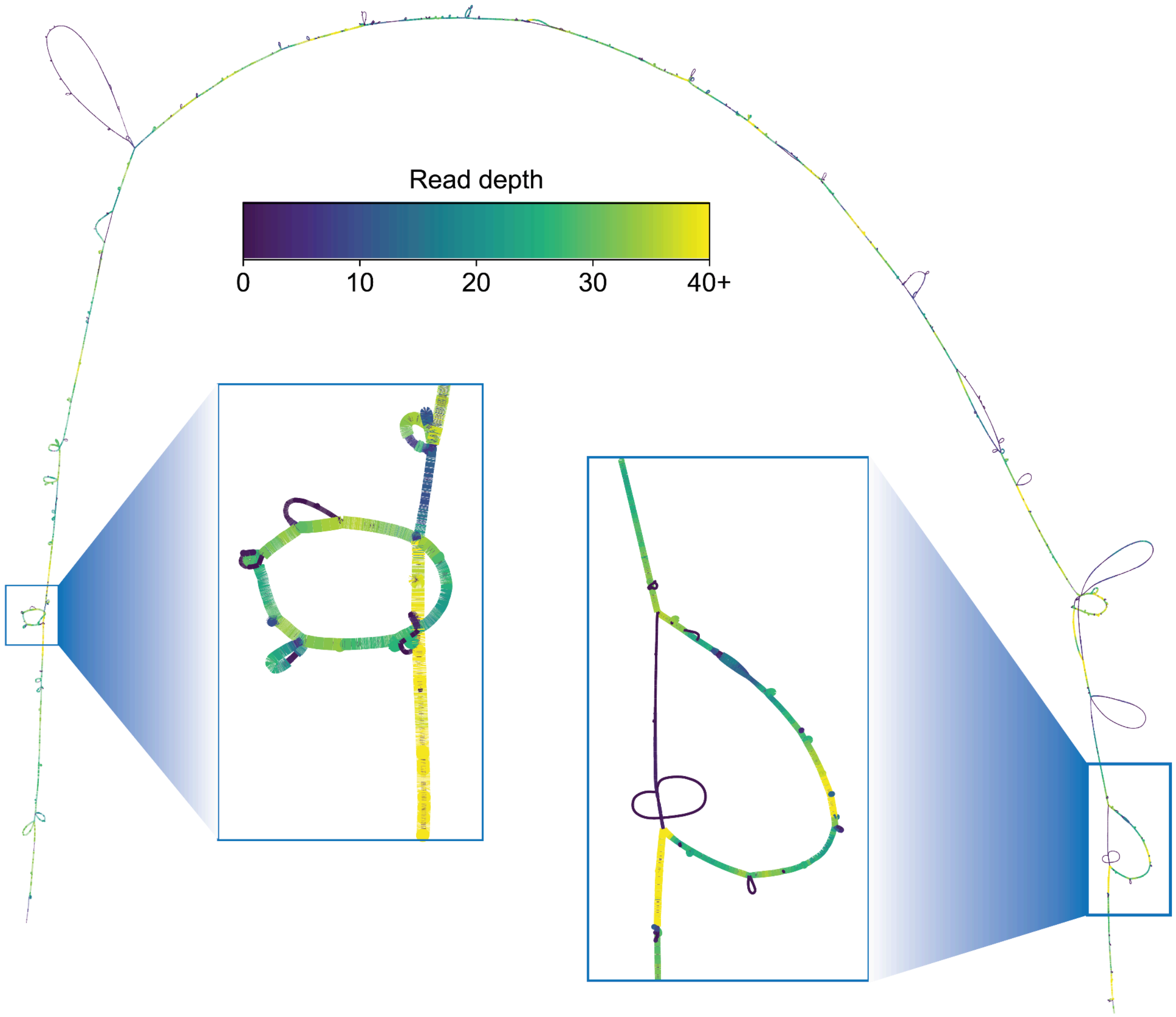


**Supplementary Figure 57: Depth on a personalized pangenome.** BandageNG visualization of the personalized pangenome graph created for haplotype HG01106.1 on Chromosome 10, with nodes colored by average HiFi read coverage.

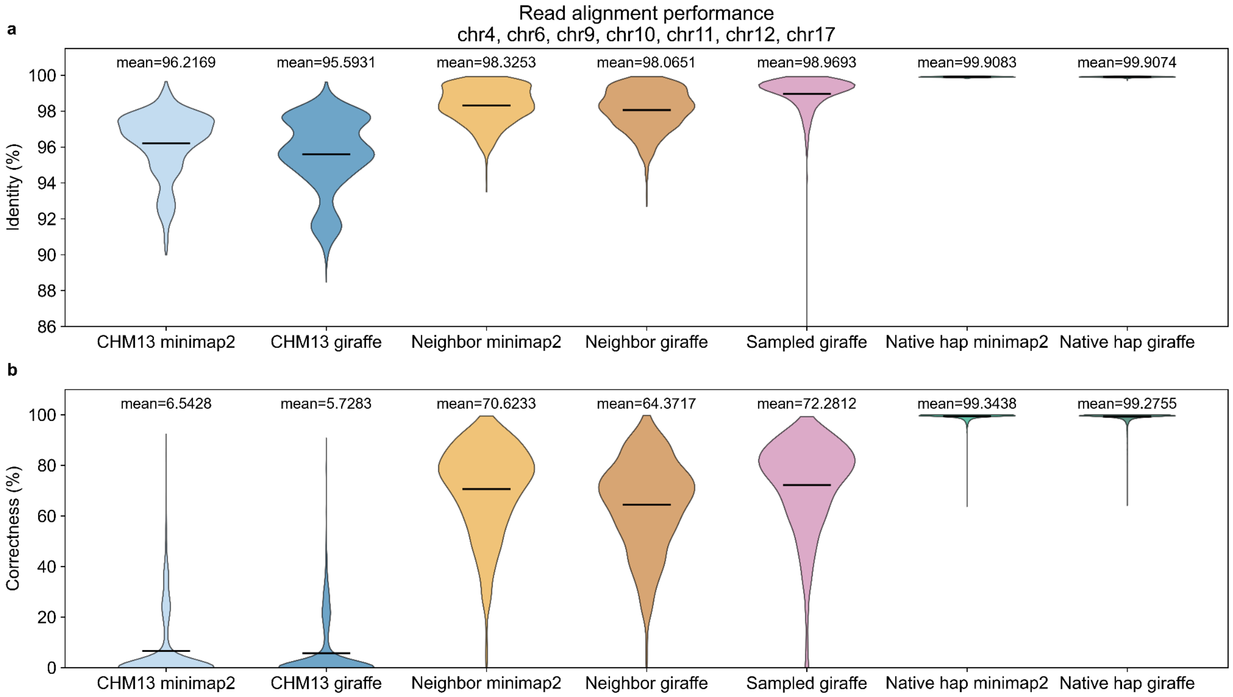


**Supplementary Figure 58: Giraffe alignments to linear references**. **a**, Average read alignment identity and **b**, average read correctness, using real reads, for alignments to CHM13, the most closely related haplotype, and the reads' own assembly. Black bars indicate means. Alignments to linear references done with Minimap2 and Giraffe.

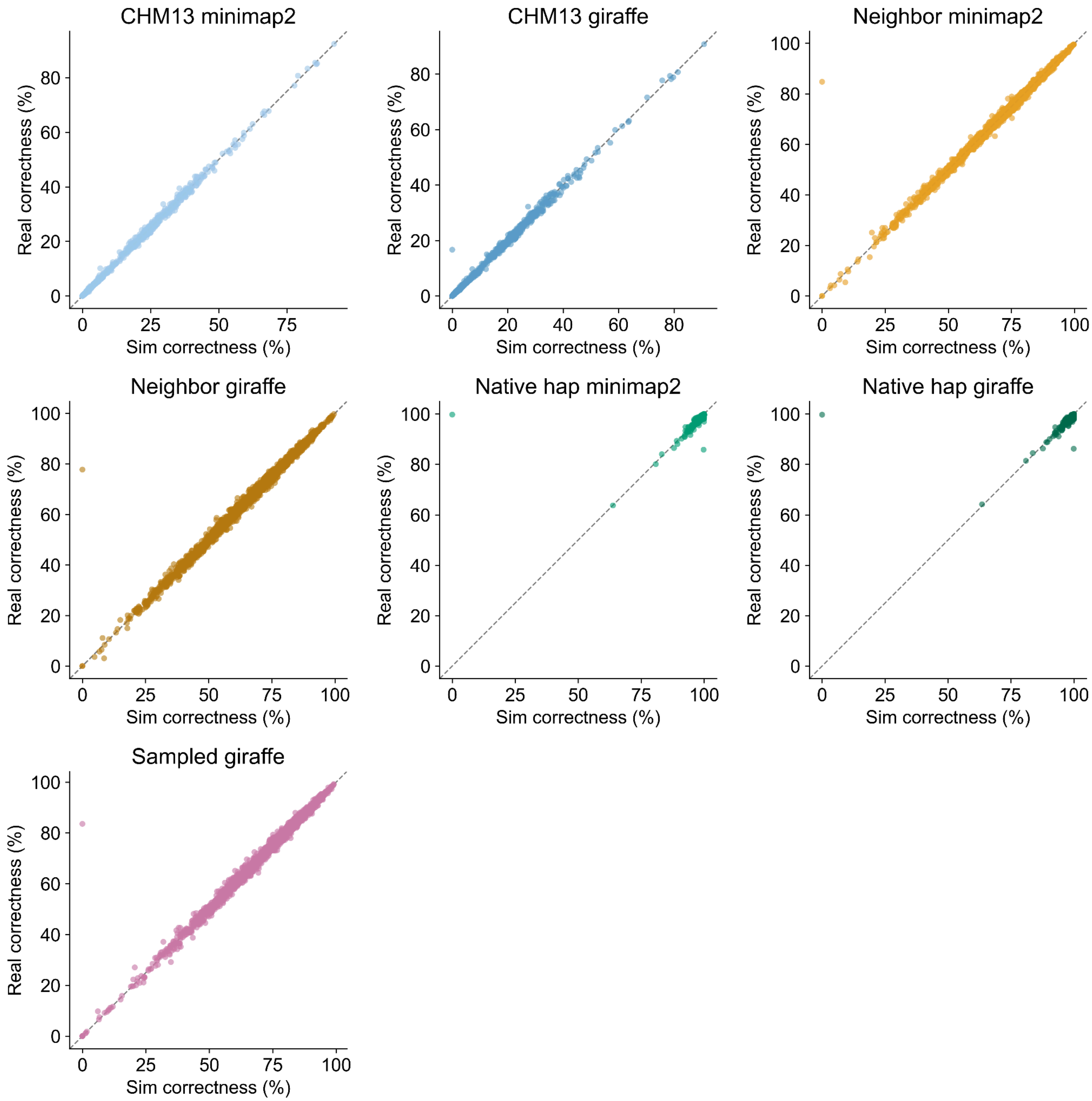


**Supplementary Figure 59: Correctness metrics as applied to real and simulated reads**. We defined a permissive measure of alignment correctness to accommodate high amounts of private sequence within the centromeres. This metric is well-correlated between real and simulated reads, indicating that the real reads' alignments are trustworthy and a good indication of true performance. Only alignments to haplotype-sampled graphs exhibit notable differences in correctness between real and simulated reads; these graphs were sampled separately for each read set.

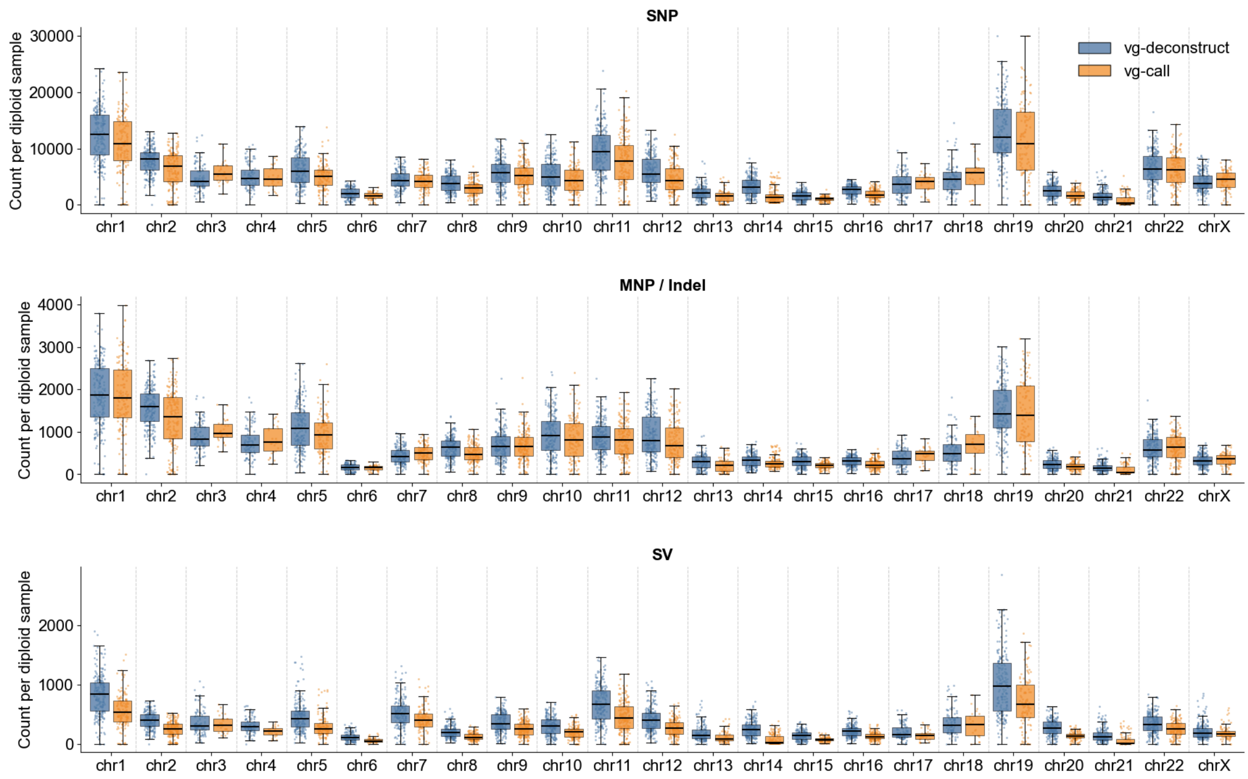


**Supplementary Figure 60: Per-chromosome variant calling results using rGFA.** Y axis represents the count of variant calls per diploid sample. Blue violin plots (left) indicate variant calls derived from variant decomposition of the Centrolign graphs with vg-deconstruct. Orange violin plots (right) indicate variant calls derived from read alignments to the Centrolign graph, using vg-call to call variants. Calls are divided into single nucleotide polymorphisms (SNPs, top), multinucleotide polymorphisms and short insertions/deletions less than 50 bp (middle) and structural variants (SVs, bottom)
